# A field theoretic approach to non-equilibrium population genetics in the strong selection regime

**DOI:** 10.1101/2023.01.16.524324

**Authors:** Daniel J. Balick

**Affiliations:** Department of Biomedical Informatics, Harvard Medical School, Boston, MA; Division of Genetics, Brigham and Women’s Hospital, Harvard Medical School, Boston, MA

**Keywords:** theoretical population genetics, non-equilibrium, natural selection, demography, allele frequency dynamics, equilibration time, stochastic field theory, exponential expansion, population bottleneck, cyclical populations

## Abstract

Natural populations are virtually never observed in equilibrium, yet equilibrium approximations comprise the majority of our understanding of population genetics. Using standard tools from statistical physics, a formalism is presented that re-expresses the stochastic equations describing allelic evolution as a partition functional over all possible allelic trajectories (‘paths’) governed by selection, mutation, and drift. A perturbative field theory is developed for strong additive selection, relevant to disease variation, that facilitates the straightforward computation of closed-form approximations for time-dependent moments of the allele frequency distribution across a wide range of non-equilibrium scenarios; examples are presented for constant population size, exponential growth, bottlenecks, and oscillatory size, all of which align well to simulations and break down just above the drift barrier. Equilibration times are computed and, even for static population size, generically extend beyond the order 1/*s* timescale associated with exponential frequency decay. Though the mutation load is largely robust to variable population size, perturbative drift-based corrections to the deterministic trajectory are readily computed. Under strong selection, the variance of a new mutation’s frequency (related to homozygosity) is dominated by drift-driven dynamics and a transient increase in variance often occurs prior to equilibrating. The excess kurtosis over skew squared is roughly constant (i.e., independent of selection, provided 2*Ns* ≳ 5) for static population size, and thus potentially sensitive to deviation from equilibrium. These insights highlight the value of such closed-form approximations, naturally generated from Feynman diagrams in a perturbative field theory, which can simply and accurately capture the parameter dependences describing a variety of non-equilibrium population genetic phenomena of interest.

## Introduction

We live in a world of non-equilibrium population genetics; nearly all inferences of historical effective population sizes (i.e., demographic history) have described dynamic scenarios with various time-dependent demographic features: bottlenecks (e.g., the ‘Out of Africa’ bottleneck in humans [1–4], barn swallow population history [5]), exponential expansion (e.g., post-agricultural human population growth [1, 4, 6], post-Ice Age growth in megafauna [7], viral transmission in a host population [8]), oscillatory changes (e.g., seasonal oscillations in the allele frequency and various phenotypes in Drosophila [9–11] and other species with short lifespans [12], serial founder effects during range expansion [13–15]), and pulses of migration between isolated populations (or connected demes [16]) and between close species (e.g., Neandertal, Denisovan archaic introgression [17–19]). Even meticulously controlled laboratory evolution experiments tend to require serial bottlenecking, albeit not necessarily severe, to maintain a temporal genetic output of the experiment in the form of a ‘living fossil record’ of frozen yeast or bacteria [20, 21]. Temporally varying phenomena, beyond changes in population size, also occur: mutation rates can change over time due to the presence of mutator[22] or anti-mutator [23] phenotypes, and the selective effects of specific alleles or the entire distribution of fitness effects can meaningfully vary if the environment changes on a relevant timescale and there are sufficient gene by environment interactions [24, 25]. Complicating matters, these temporal shifts can occur simultaneously and interact. Such phenomena are presumably observed in extant species because they are beneficial to long-term survival; dynamic processes can alter the efficacy of natural selection [26, 27] or allow populations to better respond to environmental shifts by exhibiting increased genetic variance and more genotypic variation as a consequence of changes to selection pressures [28], population size [29, 30], or mutation rate [31, 32]).

In the the context of population genetics, one can phrase any explicit time dependence driving the evolution of allele frequencies as non-equilibrium behavior. First, this can take the form of *equilibration*, i.e., the dynamics as a population approaches an equilibrium state of any kind (e.g., mutation-selection balance, mutation-drift balance, steady-state mutation-selection-drift balance [33–35]). Equilibration is characterized by the instability of the initial state, but this can occur without invoking any additional time dependence (e.g., a changing population size). This is of particular importance to understanding the behavior shortly after the emergence of a new allele in a population, which tends towards higher frequencies or extinction over time (except in finely-tuned special cases). If a sufficiently stable equilibrium point in parameter space exists relatively near the population’s current state, it will generally tend towards that equilibrium if possible (see, for example, [36]). Equilibration is restricted to populations that asymptotically approach some class of steady-state distribution of allele frequencies. Considering for a moment a population genetic simulation with constant parameters (population size, mutation rate, etc.), the equilibration process can be thought of as occurring during the ‘burn-in’ period in the first 8*N* generations before the initial conditions are effectively erased (i.e., all alleles in all individuals coalesce at a time after the initial condition has fully equilibrated). Adding a wrinkle, there can exist *transient* behavior that is not well-described by expanding around the initial or final state of the population that can only be identified during the equilibration process or a strong departure from equilibrium. The second class of non-equilibrium behavior a population may experience is characterized by an explicit time dependence in any of the parameters associated with population genetics: time-dependent population size *N*, mutation rate *μ* and back mutation rate *μ_b_*, selection coefficients *s* or the distribution of fitness effects, recombination rate *r*, or any external (i.e., environmental) or internal (e.g., mating preferences) shift that implicitly changes these parameters. In other words, a large-scale change in any of the otherwise ‘constant’ parameters describing the population at a given time will result in non-equilibrium behavior after such a change occurs. A time-dependent population size *N*(*t*) is generally referred to as the *demography* and tends to be the most commonly considered non-equilibrium scenario in the literature. However, all of the other population parameters are known to be time-dependent in nature on some timescale, as discussed above.

Given the ubiquity of non-equilibrium behavior in populations observed in nature, it is natural to hope to describe populations in such states. However, the majority of the mathematics developed to model population genetics is, by construction, restricted to equilibrium or pseudo-equilibrium regimes. This is for both analytic and intuitive simplicity, but, additionally, forays far beyond equilibrium population genetics are notoriously difficult to describe mathematically, particularly when more than one parameter can temporally vary (see, e.g., [37] for a recent paper on co-varying parameters). There are, of course, a number of exceptions (i.e., non-equilibrium models [27, 37–52]), some of which are highlighted in the Discussion. Non-equilibrium descriptions in the literature are often constructed using ad hoc techniques suited to model a specific demography or phenomena of interest (e.g., models presented in [27, 37]). Instead, this manuscript aims to describe a distinct and generalizable formalism that is flexible enough to characterize a wide range of behavior in non-equilibrium populations in a unified framework. That said, much of the present analysis focuses on details specific to strong additive purifying selection (i.e., deleterious genic selection), which, as I will show, emerges as a natural regime in this formulation of population genetics. More importantly, this regime is of practical interest, as it covers the range of selection coefficients where most deleterious variation of interest lies, from disease variants in humans [53] to a variety of phenotypes observed in Drosophila [54]. Strongly deleterious selection is also of theoretical interest, as it counteracts the frequency-increasing forces of mutation and drift (which can also decrease frequency) and is generally expected to eventually result in stable equilibria (or pseudo-equilibria in the absence of back mutations) dependent on the precise parameter values. This will be used as a premise to demonstrate that, even with this naively stable end-state, non-equilibrium behavior emerges en route to equilibrium, examples of which are described analytically in the proceeding text. Given the highly mathematical nature of field theory and its historical relegation to physics and related fields, this manuscript is written to be comprehensive and largely self-contained, serving a secondary purpose as something of a primer on field theory intended to be accessible to mathematical population geneticists. However, various supplemental resources on the topic are mentioned throughout to provide alternative treatments for interested readers, the most pertinent of which is a brief but intuitive overview of the application of field theory to solving stochastic differential equations (aimed at biologists) by Chow and Buice (2015) [55]. I have relied heavily on the latter in my presentation of the necessary background, but, given the varied expertise of population geneticists, I suggest using their review as an accessible first introduction to field theory or for clarification if the description herein is insufficient.

### Solutions to the diffusion equation

The goal of this manuscript is to describe various dynamical behaviors of a population with non-equilibrium demography (or, for example, temporally varying mutation or selection parameters) in the form of closed-form analytic approximations to the time-dependent distribution of frequencies in a population. This can be re-phrased as an attempt to find approximate solutions to the equations describing evolving populations (e.g., Kimura’s backward or forward diffusion equations [33] shown below) without resorting to setting the time derivative to zero (i.e., a steady-state approximation).

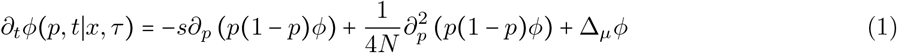

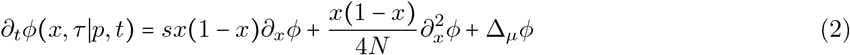

Equation 1 is Kimura’s forward equation (i.e., the forward Kolmogorov or Fokker-Planck equation) describing the dynamics of the probability distribution *ϕ*(*p, t*|*x, τ*) for reaching an allele frequency *p* at time *t*, conditional on a known frequency *x* at an earlier time *τ*, or, expressed in terms of the Dirac delta function, obeying the condition *ϕ*(*p* = *x, t* = *τ*) = *δ*(*p – x*). The three terms on the right hand side, in order, describe additive purifying natural selection with a selection coefficient *-s* (the strength of selection *s* is assumed to be positive throughout this manuscript, with *s* = |-*s*| for deleterious alleles, such that the sign remains explicit), genetic drift in a finite diploid population comprised of 2*N* haploid individuals (which will later become a time-dependent function 2*N*(*t*) describing demography), and a general term Δ*_μ_ϕ* representing the change in the probability density due to mutational influx. The mutational term has been left in general form due to the difficulty of describing new mutations appearing at a finite location *p* = 1/2*N* in the continuous frequency limit 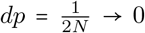, but this is usually dealt with post-hoc (i.e., by applying the appropriate mutational flux to moments of *ϕ* after computing other contributions, following the methods used by Kimura in, e.g., [56, 57]). The backward equation, shown in Equation 2, is the complement of the forward equation and describes the likelihood of having started at frequency *x* at time *τ* conditional on observing frequency *p* at a later time *t* ≥ *τ* (the conditioning for *ϕ*(*x, τ*|*p, t*) is switched to account for the different interpretation of the backward equation). The backward description of evolving allele frequencies is particularly important for inference of the state of the allele frequency distribution at some time in the past *τ*, conditional on current day observations of *p*(*t*), and is extensively used for inferring the demography *N*(*τ*) (see, e.g., [45]).

One approach to solving this partial differential equation (PDE) exactly involves representing the transition density as an infinite series. Kimura (1955, 1964) presented this class of solutions for the dynamics of a neutral allele in a population of constant size in terms of a sum over Gegenbauer polynomials [33, 58]. This is an example of restricting to an extreme limit (e.g., 2*Ns* → 0 for finite nonzero N) to provide a partial solution, which is a commonly used technique, particularly for describing time-dependent behavior. Taking the opposite limit 2*Ns* → ∞ for finite nonzero *s*, one can describe the *deterministic* behavior in an infinite sized population. In this case, the probability density collapses to a delta function at all time points (i.e., the variance goes to zero). The deterministic behavior of *p*(*t*) can then be described by the following ordinary differential equation (ODE).

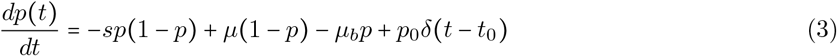

Here, I have replaced *x* and *τ* with *p*_0_ and *t*_0_, respectively, to make clear that the delta function represents an initial condition *p*(*t*_0_) = *p*_0_. I have chosen a frequency-dependent functional form for the mutational influx (i.e., Δ*_μ_p* = *μ*(1 – *p*), where *μ* is the mutation rate per individual per generation) that accounts for recurrent mutations such that the target size (*L* = 1) may be saturated due to loss of unmutated individuals (i.e., Δ*_μ_p* → 0 as *p* → 1). This is counteracted by mutational outflux (i.e., Δ*_μ_b__p* = –*μ_b_p*) with back mutation rate *μ_b_* per site. The initial condition is now specified explicitly, henceforth written as *p*_0_*δ*(*t*) after setting *t*_0_ = 0 for convenience, but can be altered appropriately to model a variety of dynamics. This non-linear equation can be solved exactly. However, the focus of all subsequent analyses is on strong selection, which maintains alleles at low frequencies *p*(*t*) ≪ 1 at all times, resulting in a much simpler deterministic equation; Equation 3 can be approximately linearized and expressed as 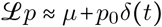, where 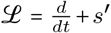 is a homogeneous linear operator dependent on *s*′ ≡ (*s + μ + μ_b_*) (note, I have intentionally excluded the inhomogeneous terms). In fact, in the infinite population size limit 2*Ns* → ∞, this low frequency linearization applies to any non-zero deleterious selection coefficient unless mutation is strong enough to overcome selection (*μ* ≫ *s*, where back mutations are rare or absent). Solving the linearized deterministic equation yields the following.

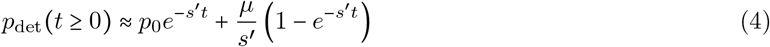

Here, *s*′ is an important constant (see Materials and Methods) that determines the rate of frequency decay over time; if mutation rates are low (i.e., *s* ≫ *μ, μ_b_*), *s*′ ≈ *s* and the timescale of exponential frequency decay is of order 1/*s*, which provides the deterministic intuition for the equilibration time. Equation 4 describes an allele frequency initially set to *p* = *p*_0_ at time *t* = 0, with subsequent exponential decay away from this initial frequency and towards a state of mutation-selection balance that approaches frequency *p* = *μ/s*′ at asymptotically late times *t* → ∞. This approximate deterministic solution under strong selection is relevant to expressions derived later in this manuscript (see Results) because it captures the behavior of one variable of particular interest, the mean frequency of an allele *E*[*p*(*t*)] (this becomes the mutation load 〈*p*〉 = *L* × *E*[*p*(*t*)] for *L* independently evolving sites under strong additive selection and weak mutation 2*N_μ_* ≪ 1, where *μ* ≫ *μ_b_*). Song and Steinrücken (2012) went beyond the extreme limit approach to present an exact solution for the transition density for general diploid selection coefficients (*s*_hom_ = *s, s*_het_ = *hs*) evolving in a population of constant size [42]. Later, Živković et al. (2015) generalized this analysis to produce an exact solution for non-equilibrium population size variation (i.e., where *N* = *N*(*t*) becomes a time-dependent function) [43]. Both of these exact solutions are presented in the form of an infinite series, like Kimura’s solution for neutrality [33, 58], and describe the temporal evolution of transition density *ϕ*(*p, t, s*_het_, *s*_hom_|*p*_0_, *t*_0_).

The existence of such solutions provides proof that the relevant equation describing population genetics is indeed soluble and can be useful for understanding and modeling population dynamics computationally, particularly for non-equilibrium scenarios of interest. And yet, an infinite series is somewhat unsatisfying in that it obscures intuition and requires algorithm-like steps to compute results for a given scenario to the desired accuracy.

An alternative approach is simply to solve the equation in the relevant parameter regime numerically. There are now several software packages that use numerical solutions to infer various population genetic parameters (population sizes and splits, migration patterns, distribution of fitness effects, etc.). Amongst the most well-known computational software, ∂a∂i takes the end-state site frequency spectrum as input and uses that to numerically solve the finite difference equations that correspond to the discretized backward diffusion equation (i.e., the complement to 1) [45]. A newer, alternative strategy, `moments`, makes use of a system of discrete ODEs describing the temporal evolution of the sample frequency spectrum that, when the diffusion approximation is applicable, corresponds to the system of equations describing the evolution of moments of the allele frequency spectrum in a non-canonical basis of polynomials [46]. The authors demonstrated that an approximate solution can be obtained by using jackknife extrapolation to approximately close this system of equations, which was shown to outperform methods like *∂*a*∂*i that numerically solve the diffusion equation directly with respect to computational efficiency and flexibility. As helpful as these numerical solutions are to the inference of various parameters, there is little population genetic intuition gained by doing so, short of a rigorous exploration of the vast parameter space of possible non-equilibrium demographic histories, along with the many other population parameters.

Despite a wide range of empirical applications, the aforementioned techniques to solve for the allele frequency dynamics away from equilibrium have, in my view, left something of a gap our understanding of the provided solutions. To intuitively grasp generic phenomena in non-equilibrium population genetics, it is perhaps of more practical use to derive closed-form analytic solutions to Equation 1, even if approximate. Closed-form approximations augment the current methods by providing clear and concise encapsulations of the relevant parameters that drive the dynamics of interest, usually at the cost of a more limited scope. As this is a common perspective in some areas of physics, an appropriate mathematical framework to naturally generate approximations for non-equilibrium phenomena already exists; stochastic field theory (SFT) and quantum field theory (QFT), related fields that differ in purpose and methodology, together provide a rich set of well established perturbation-based techniques for approximating the behavior of a dynamic system (see [59] for an introduction to path integration in SFT; see [60–62] for in depth introductions to QFT in particle physics and many-particle systems). SFT focuses on using path integration to *average out* stochastic behavior to generate functional dependences for the expectation values of a set of quantities affected by some stochastic process. In addition to many other powerful techniques, QFT provides a straightforward perturbative approach (applied when exact time-dependent results are intractable) to produce a simplified time-dependent description of the interactions between particles (usually referred to as ‘scattering’ when studying the behavior of particle collisions) [60, 61] or the emergent behavior of a large number of particles comprising a macroscopic material [62]. Perturbative QFT has been shown to yield phenomenally accurate predictions in particle physics, in particular (see, e.g., [63, 64] for the most extreme accuracy test to date). While there is no hope of competing with the accuracy of fundamental physics in a biological system, this provides sufficient motivation to attempt to construct a field theoretic description of stochastic population genetics with the aim of using perturbation theory to generate quantitatively accurate predictions that sidestep many of the complications and non-linearities encountered in non-equilibrium systems.

### Path integrals in population genetics

Importantly, some effort has been made in the theoretical population genetics community to use path integration, the construct describing a field theory, for a diverse set of applications. Rouhani and Barton (1987) discussed path integration as a perturbative technique to derive transition rates between fitness optima in their analysis of Wright’s ‘shifting balance’ scheme [47]. Their results were placed in an evolutionary context to assess the plausibility of speciation as a consequence of genetic divergence primarily driven by genetic drift. Barton (1989) later described the evolution of highly polygenic traits, in a somewhat similar context, using path integration over a range of local phenotypic optima and nearby regions to compute, amongst other variables of interest, the allele frequency distribution in the neighborhood of the optima [48]. In a distinct application, Mustonen and Lässig (2010) introduced the notion of fitness flux, a measure that accounts for the adaptive history of a population by integrating over the net selection and rate of adaptation over all accessible genotypes (akin to a path integral defined on genotypic space) in a temporally evolving population [49]. Neher and Shraiman (2012) used path integration to account for fluctuations when computing the rate of Mueller’s ratchet in a completely asexual population [50]. They evaluated the response of the bulk of the population to fluctuations in the fittest class that result in the loss of this least-loaded class, but showed that this is susceptible to a delayed restoring force generated by fluctuations of the mean fitness that modifies the rate of fitness loss. In a totally different context, a recent study by Nourmohammad and Eksin (2021) used a path integral formalism to determine a means of the optimal means of intervention via artificial selection in applications ranging from the genetics of domesticated species to strategies for epidemiological response [65]. There, path integration was used to characterize the consequences of human intervention on the distribution of all possible evolutionary trajectories, which allows for an informed optimization that accounts for the stochastic future of the population. Notably, each of these studies presented path integration as a calculational tool, but did not introduce other techniques commonly used in field theory. Most conspicuously, diagrammatic expansions (i.e., Feynman diagrams), which provide a simplified means of evaluation of a series of contributions to an observable of interest, are absent from (and largely unneeded in) the aforementioned literature. In a notable exception, Lorenz, Park, and Deem (2013) studied a Moran model of asexual evolution by rephrasing the stochastic dynamics as a mean field theory (a field theoretic technique used to simplify systems with a large number of degrees of freedom; see [62]) and used the corresponding Feynman diagrams to compute generic properties of the mean fitness in an arbitrary fitness landscape [66]. While the biological system and methods may differ from those presented here (e.g., mean field theory, saddle-point approximation, and other techniques used therein differ substantially), the authors of [66] presented a robust application of field theory to population dynamics, using a number of relevant techniques beyond path integration (see [67] for an additional example from two of the authors).

The theoretical work described in Schraiber (2014) [51] is specifically relevant to the field theoretic reformulation of allelic dynamics. Schraiber constructed a path integral representation of the transition density for alleles under additive selection in a finite population. To accomplish this, he computed the transition density for drift-driven neutral alleles, derived from Kimura’s work on diffusion theory [33], and used this to normalize the path integral characterizing the transition density for alleles under selection. Treating neutral allele trajectories as a null, paths involving selection were expanded in powers of 2*Ns* to form an infinite series of integrals over neutral paths; this was then treated perturbatively in the weak selection regime (2*Ns* ≪ 1) by truncating the series to approximate the transition density. This approximation corresponds to a diagrammatic representation of an allele propagating under stochastic drift, occasionally interrupted by interactions with a weak potential generated by natural selection; this is complementary to the analysis described herein, which depicts alleles propagating under efficient selection, occasionally interrupted by interactions with stochastic drift or mutation. By loose analogy, the formalism used in [51] follows directly from the diffusion equation of population genetics and describes propagation in a potential generated by selection, which subsequently resembles the path integration techniques used in quantum mechanics (of course, with ‘quantum’ replaced with ‘stochastic’ behavior). In contrast, the present formalism is rooted in the Langevin representation (see Materials and Methods) of the dynamics and shares more in common with techniques from quantum field theory (again, ‘quantum’ → ‘stochastic’). Schraiber’s paper thus plays an important conceptual role in the work presented in the following sections, as it introduced the notion of path integration over stochastic allele frequency trajectories, but the focus on a non-overlapping dynamical regime limits the direct relevance of the analytic results derived therein.

## Materials and Methods

Due to the large number of quantities and parameters referred to in this manuscript, many of which may be unfamiliar to geneticists, Supplemental Material S1 has been provided for the reader’s convenience: Table S1 contains definitions of relevant functions, functionals, and symbols, Table S2 defines general population genetic parameters, and Table S3 defines demography-specific parameters. Throughout this manuscript, I restrict all equations to the simplest possible dynamics (unless otherwise stated) for clarity: deleterious additive selection with a single strength *s* (negative sign explicit), constant mutation rate *μ* and back mutation rate *μ_b_* (allowing for recurrent mutations if 2*N_μ_* ≳ 1), and a diploid population size of *N* with 2*N* haploid individuals. For the beginning of this manuscript, including Materials and Methods, *N* may be assumed to be constant over time until non-equilibrium demography is introduced, after which the population size is generalized to a deterministic time-dependent function *N*(*t*) describing the demographic history of the population. I focus on the dynamics of a single derived allele with frequency *p*(*t*), but all expressions can be trivially generalized to *L* independently evolving (i.e., unlinked) loci if desired (provided mutation remains relatively weak with 2*N_μ_* ≪ 1).

### A field theoretic treatment of the stochastic differential equation for evolving populations

Path integration first arose in the context of stochastic differential equations (SDEs) and is generally attributed to Norbert Wiener, who devised an integration scheme to study Brownian motion and diffusion [59]. Given that the dynamics of population genetics can be described by an analogous diffusion-like equation [33], there must be a path integral representation of finite population dynamics (e.g., the formulation discussed in [51]). However, the present goal is not simply to squeeze population genetics into a specific formalism, but rather to use a particularly well-developed and flexible formalism to address questions relevant to non-equilibrium population dynamics.

#### Langevin equation representation of allelic dynamics

The diffusion model of population genetics, usually represented by the PDEs in Equations 1 and 2, can be equivalently expressed in terms of an explicit stochastic variable (for derivations, see [59, 68]). Focusing on the forward equation, this is the same relationship as between the Fokker-Planck equation and the Langevin equation, which provide alternative descriptions of the same phenomena. The Langevin equation that describes the temporal dynamics of a single allele in a finite population, including the explicit initial condition (dependence on the initial time *t*_0_ is shown in Materials and Methods for clarity; *t*_0_ = 0 is again used in Results), can be written as follows.

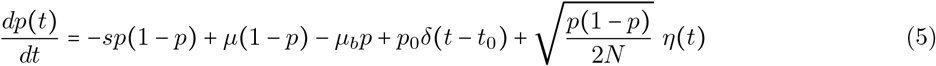

This expression is identical to the deterministic equation shown in Equation 3 other than the *η*(*t*) dependence, which introduces stochastic behavior due to genetic drift. Here, *η*(*t*) represents Gaussian white noise with a mean of zero (i.e., *E*[*η*(*t*)] = 0) and unit variance (i.e., Var[*η*(*t*)] = *E*[*η*^2^(*t*)] = 1) and satisfies the Markov condition *E*[*η*(*t*_1_)*η*(*t*_2_)] = *δ*(*t*_2_ – *t*_1_), which requires that noise contributions from distinct time points remain uncorrelated. This is provides an alternative description of the population genetic behavior expressed in terms of the allele frequency *p*(*t*), directly, which is now stochastic, rather than in terms of the probability distribution *ϕ*(*p, t*); the PDE was simply converted to an SDE. This formulation was previously used for several distinct applications to population genetics in the literature (see, e.g., [50, 69]), as it makes properties of drift-based fluctuations more straightforward to compute. In the present context, this formulation is only an intermediate step en route to a field theoretic representation.

#### Action and partition functional for a population genetic field theory

Following the derivation presented in Chow and Buice [55], which provides an excellent and highly relevant crash course on several methods for solving SDEs using field theory, the Langevin equation in Equation 5 can be re-expressed as an *action* 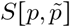 of the following form (see Appendix A1 for full derivation).

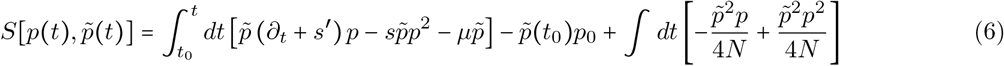

For notational convenience, the total derivative over time 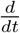 is denoted as *∂_t_* here and henceforth. All terms linear in *p*, and thus bilinear in *p* and 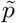, have been collected in the first term (recall that the linear operator 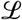 from the deterministic equation is the same as that in the parentheses) which defines the constant *s*′ (usually referred to as the ‘mass’ by analogy to particle physics) as follows.

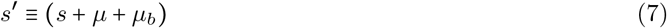

The effective selection coefficient *s*′ is the sum of all allele frequency-reducing effects: deleterious natural selection *s*, saturation of the mutational target (*L* = 1) into the derived allele due to recurrent mutations, and back mutation *μ_b_* away from the derived allele. As discussed in Equation 4, this parameter determines the timescale of exponential frequency decay in the strong selection limit. In the infinite sites limit, defined by the the absence of recurrent mutations (2*N_μ_* ≪ 1) and rare or absent back mutations (*μ_b_* ≪ *μ*), this parameter simply reduces to *s*′ → *s*, the deleterious selection coefficient.

The action in Equation 6 formally treats *p*(*t*) as an *allele frequency field* (i.e., a scalar valued function defined at all time points *t_i_* in the continuum limit *dt* = (*t*_*i*+1_ – *t_i_*) → 0), probabilistic properties of which (e.g., moments of the probability distribution) are the primary topic of interest in this field theory. For the present purposes, 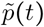 can be considered an abstract mathematical tool used to calculate properties of the allele frequency field *p*(*t*). The conjugate field 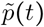, which is an imaginary field (i.e., a complex field with non-zero values only along the imaginary axis in the complex plane), is defined by 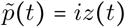, where *z*(*t*) is the Fourier dual field that comes from re-expressing the probability density for *p*(*t*) in Fourier space at every time point (see full derivation in Appendix A1; additional discussion of the field 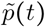 can be found in [55]). In fact, the field 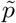 can be entirely removed from the action by explicit path integration, if desired, but it is useful for constructing the diagrammatic representation described in the following sections and has a simple interpretation in that context [55]. More intuitively, 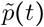 plays a similar role the variable *x* that appears in the diffusion equations; the relationship between the fields 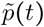 and *p*(*t*) is somewhat analogous to that between the allele frequencies *x* and *p* in Equation 1. The transition density *ϕ*(*p, t*|*x,τ*) defined in the forward diffusion equation is concerned with probabilistically predicting the distribution of allele frequencies *p* at some time *t*, conditional on a value of frequency *x* at time *τ*; time ordering *t* > *τ* is implicit, here, (*t* is in the future relative to *τ*) and transition densities describe probabilities from a past frequency *x* that cause changes to a future frequency *p*, including infinitesimal transitions between adjacent time points *x*(*τ*) → *p*(*τ + dt*). Similarly, the action in Equation 6 assigns probabilities to *paths*, world-lines representing potential allele frequency trajectories between the fields 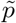 and *p*. Again, there is an implicit time ordering that requires that 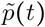 occurs prior to *p*(*t + dt*) due to the kinetic action of the time derivative *∂_t_p*. This defines causality for the allele frequency trajectory connecting the fields, which propagates along paths 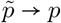. Further interpretation of 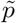 is discussed in the next subsection on allele frequency propagation and, later, in the context of Feynman diagrams.

The action in Equation 6 defines the *partition functional* 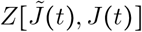, which serves as the moment generating function for expectations of powers of the allele frequency (e.g., mean frequency E[p(*t*)], homozygosity *E*[*p*^2^(*t*)]). Like a moment generating function, the partition functional was created by taking Laplace transforms of the probability distribution with respect to the functions *p*(*t*) and 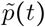 (see Appendix A1), resulting in a dependence on the arbitrary functions 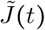 and *J*(*t*), respectively.

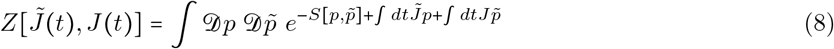

The path integral over 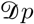 and 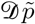 defines probabilities on the space of possible functional forms for *p*(*t*) and 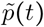, with a probability density for each infinitesimal time slice dictated by the exponentiated action *e^−s^*. In many field theories, the normalization of the path integral 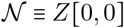 plays an important role in controlling divergences (see Appendix A1.1 for discussion); however, this path integral is finite and well behaved and the normalization is (thankfully) simply 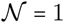. Moments of the allele frequency probability distribution can be obtained by taking functional derivatives of the partition functional with respect to 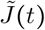 (i.e., 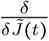) or *J*(*t*) and taking the limit 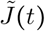, *J*(*t*) → 0 (if needed, see Supplemental Material S2 for a brief note on applying functional derivatives). The procedure to compute general moments of the form 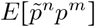 is analogous to computing moments via partial derivatives from a moment generating function.

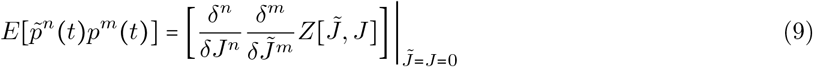

Applying the functional derivatives and taking the limit 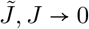, *J* → 0 results in fully time-dependent (i.e., non-equilibrium, unless special conditions are met) solutions for the temporal trajectory of a given moment of the allele frequency probability distribution (e.g., the time-dependent mean frequency *E*[*p*(*t*)]). The moments *E*[*p^n^*] have a straightforward interpretation as moments of the probability distribution, evaluated at time *t*, conditional on the initial conditions (i..e., in the language of Equation 1, moments of *ϕ*(*p*(*t*), *t*|*x, τ*) with (*x, τ*) = (*p*_0_, *t*_0_)). Moments of 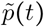 are related to solutions of the backward equation shown in Equation 2, but will not be elaborated on further in this manuscript. At this point, I will refer to this formulation of population genetics a *population genetic field theory* (PGFT), which is a reasonable description given the well-defined action and partition functional. This is technically a complete description of the field theory, having specified the partition functional, but direct evaluation of the path integral (i.e., integrating the infinite set of, in this case, non-Gaussian integrals) defined by the (quite complicated) action in Equation 6 is extremely difficult, if not impossible. Instead, I will describe a set of perturbative techniques, collectively referred to as perturbative field theory, that can be used to generate analytic approximations of the moments *E*[*p^n^*] from the PGFT partition functional.

#### Propagation of the allele frequency

To evaluate any quantity of interest in PGFT, one must first define a *propagator* for the allele frequency field *p*(*t*). This dictates the way in which an allele moves through time (i.e., the infinitesimal transition from 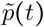 to *p*(*t* + *ϵ*) as defined in the discretized form shown in Appendix A1). The infinitesimal propagator is heuristically defined by the two point function, the expectation 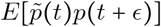, which yields additional insight into the interpretation of 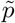: the field 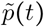 represents a *causal* allele frequency (i.e., the allele frequency at this time point causes a change to the allele frequency in the future), while *p*(*t + ϵ*) represents an *affected* allele frequency (i.e.,the allele frequency at this time point is the result of being affected by past events). Here, the notion of ‘causal’ and ‘affected’ refer to the direction of causality (i.e., the the arrow of time), where 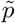 represents a cause and *p* represents an effect; equivalently stated, 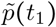 represents an *outgoing* allele frequency, carrying causal information away from *t*_1_, while *p*(*t*_2_) represents an *incoming* allele frequency, incorporating causal information into time *t*_2_. Allele frequency trajectories thus represent flows of causal information (about the probability) resulting in a distribution of time-dependent allele frequency paths. The notions of ‘incoming’ and ‘outgoing’ will become significantly clearer in the context of Feynman diagrams, which explicitly include the arrow of time.

As discussed above, the exact propagator 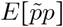 for the particularly complicated action defined in Equation 6 is extremely difficult to evaluate in directly via path integration. However, by isolating the Gaussian component of this action (i.e., the part bilinear in *p* and 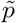), which I will refer to as the *free action* 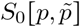, one can evaluate this oversimplified partition functional exactly to obtain an explicit moment generating functional in terms of 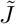 and *J*; this is derived in [55] using nearly the same free action. With this in mind, by treating the bilinear action 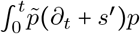 as the dominant term, corrections to the free action due to all other terms can be computed by centering around the Gaussian (i.e., bilinear) approximation; in other words, perturbation theory approximates the action as bilinear in 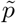 and *p*, moments of which are readily evaluated using an explicit generating function, and treats all remaining terms in the action as perturbations around the bilinear model (including those involving more than two fields, like the first drift term 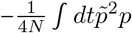, which make the full action insoluble). Importantly, this is not always possible across the entire parameter space (i.e., for all combinations of (2*N, μ, s, p*_0_) that define the strengths of drift, mutation, selection, and the initial condition), so I will at some point sacrifice generality for analytic tractability.

The *free action* 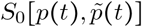 is defined simply as the bilinear (exactly solvable) component of the action 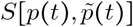, which only contains terms from the homogeneous linear component of the Langevin equation in Equation 5 (again, recall the deterministic homogeneous linear operator 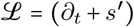 used to approximately solve Equation 3; this procedure is analogous).

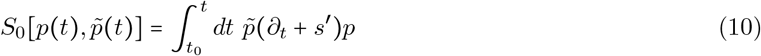

Note that, here, I have excluded terms in Equation 6 that are linear in 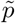, but not in *p* (the mutation term 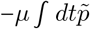 and initial condition 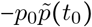 have no dependence on *p*, as written). If desired, these linear terms can be absorbed into the free action via a change of variables referred to as a field redefinition (i.e., changing the full inhomogeneous linear operator to a homogeneous linear operator by completing the square), but I have chosen not to do so because this obscures intuition about the population genetic forces represented by each term in the action. Interestingly, the free action in Equation 10 is strictly deterministic (i.e.,non-stochastic); the linear operator (*∂_t_ + s*′), which happens to be the same as the homogeneous operator 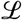 for the linearized deterministic equation under strong selection, is not dependent on the terms representing stochastic drift. 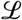 can be used to construct an operator representing the inverse propagator *G*^-1^(*t*_2_ – *t*_1_), which is again strictly deterministic.

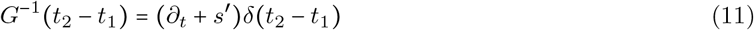

The partition functional for the free action 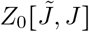 after explicit path integration, i.e., the generating function for this bilinear action, takes the following form in terms of the inverse propagator (this can be derived via Gaussian functional integration and is provided explicitly in [55]).

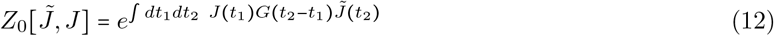

In this expression, *G*(*t*_2_ – *t*_1_) is the propagator, the operator inverse of *G*^-1^, which can be obtained by solving the following Green’s function equation [55].

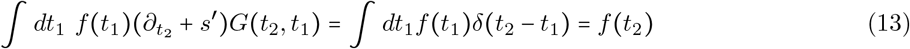

Here, *f*(*t*) is an arbitrary function, indicating that *G*(*t*_2_, *t*_1_) is the inverse operator of (*∂_t_ + s*′). Using this definition it becomes clear that the propagator is similar to a linear response function of the affected (i.e., incoming) allele frequency *p*, defined through convolution with the causal (i.e., outgoing) frequency 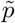. The solution for *G*(*t*_2_, *t*_1_), which must be a function of the combined variable (*t*_2_ – *t*_1_) due to the time dependence of the inverse propagator, is the following (this can be confirmed by substitution into Equation 13).

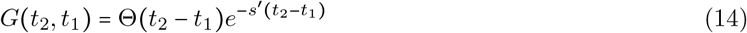

This is the free propagator for PGFT, which is expressed in terms of the Heaviside function Θ(*t – t*′) using the left-continuous definition: Θ(*x* > 0) = 1 and Θ(*x* ≤ 0) = 0 (importantly, Θ(0) = 0). The Heaviside function appearing in the propagator imposes an explicit *time ordering* on the free allele dynamics ([55] noted that this is a direct manifestation of the Itô condition), i.e., *t*_1_ must occur *earlier* than *t*_2_, or there is no propagation of allele frequency (the left-continuous definition of the Heaviside function, with Θ(0) = 0, also enforces that they occur at *different* time points). This makes the causal interpretation of the infinitesimal path 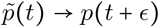 (and the interpretation of 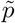 and *p*) clearer; the fields 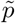 and *p* are intrinsically connected by the propagator and together define the allele frequency trajectory. The probabilistic space is defined directly on the set of paths between 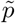 and *p*, which collectively describe the transition density. As an aside, the free propagator shown in Equation 14 is fundamentally different from that of most (free particle) quantum field theories [60, 61], which usually describe wave-like propagation due to second derivatives in time and space (i.e., the free action contains a term of the form 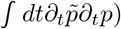, rather than diffusion or advection due to a single time derivative. Instead, the free action in Equation 10 happens to correspond to the free action for Ornstein–Uhlenbeck (O-U) process, and, as a result, the propagator is the same as the free propagator for the O-U process, a fact which may come in handy for future applications. As with the free action, the free propagator is deterministic, with no reference to genetic drift. Thus, the free allele dynamics are simply deterministic exponential decay due to the effective selection *s*′.

Returning to the non-bilinear terms, I will refer to the collection of remaining terms in the action as the *coupling action* 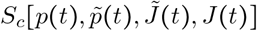 (often referred to as the *interacting action S_I_*, or *interaction terms*), which describes all interactions other than the trivial propagation of allele frequency (i.e., exponential decay over time). In order from left to right, these terms represent interactions corresponding to: non-linear selection at high frequency (due to the fixation boundary), mutational influx, an allele frequency source at time *t*_0_ (the initial condition), and fluctuations due to drift at low frequency (from the extinction boundary) and high frequency (from the fixation boundary), respectively; the source functions 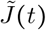 and *J*(*t*) have also been included in the coupling action, which will become important shortly.

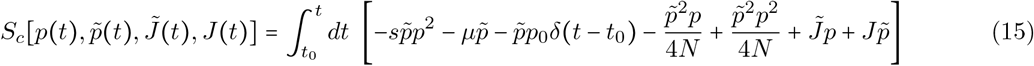

As stated above, further analysis using perturbation theory to take advantage of the free action requires that each term in the coupling action remains subdominant to the bilinear terms in the free action at all times. Generally speaking, this restricts the parameter regime described by the theory, the extent to which depends on how natural this assumption is to the interacting phenomena described by the coupling action. This restriction can be placed on the parameters (*s, μ*, etc.) directly, but the dependence on allele frequencies 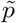 and *p* is also relevant, which can make the parameter regime more permissive; if one restricts to the regime of relatively efficient natural selection 2*Ns* > 1 at all times, the allele frequencies will generally remain low, which aids in treating the interactions as subdominant. For example, analogous to the linearization of the deterministic equation, the high frequency selection interaction 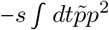 will generally remain smaller than the bilinear term 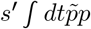 due to the higher order dependence on *p*, provided the allele frequency stays below roughly 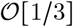, which can be accomplished with modest selection coefficients; the same may be true of the high frequency drift interaction 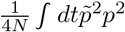, but this now depends on the efficacy of selection 2*Ns* and thus the population size (this is highly relevant to the demographic scenarios discussed below).

With the inclusion of the subdominant interactions defined in the coupling action, the allele frequency propagates freely via exponential decay in time, after the initial condition at *t* = *t*_0_, due to the effective selection *s*′; however, this can be interrupted, albeit definitionally rarely, by any or all of the interactions described by the coupling action in Equation 15. The free propagator in Equation 14 will be used extensively in all subsequent analyses as an approximate functional form for the moment 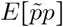.

#### Perturbation theory around the free action

The functional nature of the path integral accounts for, at least theoretically, contributions from the set of *all possible paths* the allele frequency can take through time, between (*p*_0_, *t*_0_) and (*p, t*) (i.e., the set of all possible stochastic changes in frequency due to selection, mutation, and drift that can occur at every moment between *t*_0_ and *t*). However, the contribution from each path is weighted by the probability that it occurs (i.e., is exponentially suppressed by the action *e^−S^*), which can be vanishingly small for all but the simplest paths. Perturbative field theory, detailed using a simplified example below, provides a means of identifying which paths are non-negligible, evaluating contributions from these paths approximately by using the free propagator to characterize the basic dynamics, and ignoring all other possible paths that leave the result largely unchanged. In principle, this allows for calculation to arbitrary accuracy by successively evaluating and adding contributions from the next-most important paths, but, the higher the desired accuracy, the more tedious the required calculations.

This section describes a brute force method for calculating moments from the partition function, which has, in practice, largely been replaced by more elegant methods (e.g., diagrammatic techniques, techniques related to various symmetries, etc.), depending on the context. However, most perturbative analyses perform essentially the same steps and other presentations tend to obscure the procedure to some extent. In this sense, the following prescription for directly applying functional derivatives is perhaps the clearest way of presenting perturbation theory and is the most directly analogous to more familiar moment generating function methods used in population genetics. Having said that, readers may choose to skip this section and return to it, if necessary, after reading about Feynman diagrams.

To evaluate a given moment approximately, one can Taylor expand the coupling action around the free action.

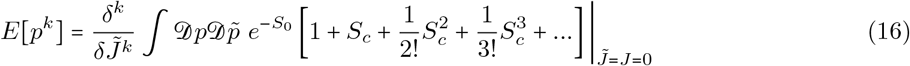

This moment can be approximated by truncating this series at an arbitrary order in *S_c_* for the desired accuracy and evaluating all remaining terms. Since *S_c_* is a sum of several terms with distinct *coupling constants* (i.e., parameters defining the strength of various interactions: *s, μ, p*_0_, and 1/4*N*) and frequency dependence, along with the source functions 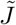 and *J*, there are a combinatoric number of distinct contributions that are generated from the expansion at a given order of *S_c_*. However, the majority of terms representing potential contributions to a given moment vanish for a number of reasons. The *k*^th^ moment *E*[*p^k^*] only has nonvanishing contributions from terms containing exactly *k* copies of 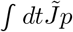 and zero copies of 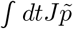; applying the functional derivatives and taking 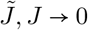, *J* → 0 removes all other terms. Thus, the lowest order contributions (i.e., the smallest power *n* for the expansion in 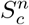 that does not completely vanish) to the expectation value *E*[*p^k^*] must be at least of order 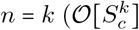, where terms that appear at order *n* in the *S_c_* series are denoted as 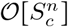 in the context of this subsection), but may be higher due to other considerations. Additionally, the first two sets of terms in the expansion trivially vanish for all moments (except in the zeroth moment *E*[*p*^0^], the normalization of the partition functional 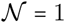): the constant term at 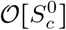 is removed by the functional derivative(s) and the only possible remaining term at 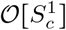 (originating from 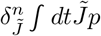) has no 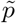 dependence and thus vanishes (because the free action *S*_0_ in Equation 10 is bilinear in 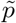 and *p*; see below). Thus, the simplest non-vanishing examples one can use are found in the 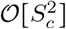 contributions to the mean frequency *E*[*p*], one of which is written explicitly below (the second is analogous, but describes the initial condition, rather than mutational influx).

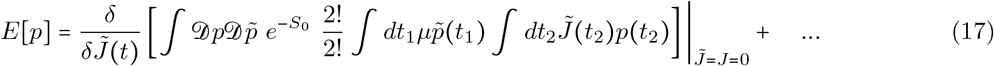

Here, the ellipsis represents all additional non-vanishing terms in the infinite series. The prefactor 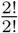 comes from the Taylor expansion of the exponential (i.e., 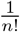 for the 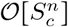 term in the series for *e^−S_c_^*; in the above 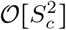 term, this is 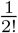) and the multinomial coefficient from expanding the sum of terms in *S_c_* to the *n*^th^ power (2! in this example). More generally, the multinomial coefficient at 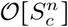 is 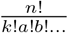; all terms at 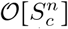 have a common overall factor of n!, contributions to the moment *E*[*p^k^*] have a common overall factor of 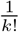 (from the *k* copies of the integral 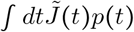 that will be pulled out by *k* functional derivatives), and the remaining factorials 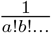 (= 1 in the above example) count multiple copies of any additional repeated integrals (e.g., a term with two copies of the integral 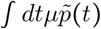 and three copies of 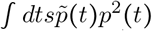 will have an overall factor of 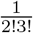; any non-repeated integrals have a factor of 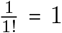). The product 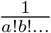 (equal to one in Equation 17 since the only remaining integral, 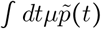, is not repeated), where the factor of 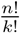 has been removed, is related to the *counting factor* of this contribution, which is discussed below in the context of Feynman diagrams.

To evaluate the path integral in Equation 16 for a moment *E*[*p^k^*] at 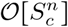 (in the current example, the first moment *E*[*p*] expanded to 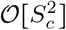), one can make use of the bilinear functional form of the free action *S*_0_.

Equation 17 can be evaluated by first applying the functional derivative and setting 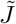 to zero.

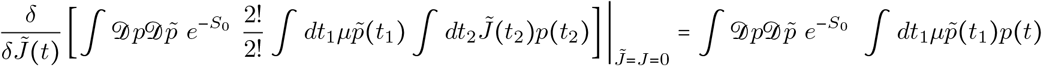

Note that the resulting path integral is bilinear in *p* and 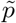, which implies it is non-vanishing when integrating using the (bilinear) free action. The path integration elements 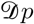 and 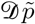 commute with *dt*_1_, allowing for a change in the order of integration. As the free action is independent of *t*_1_, this can be rewritten as follows.

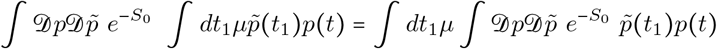

This is simply the two-point function defined as the expectation 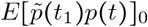 evaluated using the free action (i.e., path integrated over the density *e*^-*S*_0_^, as denoted by the subscript); this is equivalent to the propagator *G*(*t, t*_1_), which can be seen immediately from the moment generating function in Equation 12. Substituting in the free propagator and using the definition in Equation 14, the path integral can now be evaluated directly by integrating over the time *t*_1_ (i.e., the time when mutation occurs, changing the allele frequency).

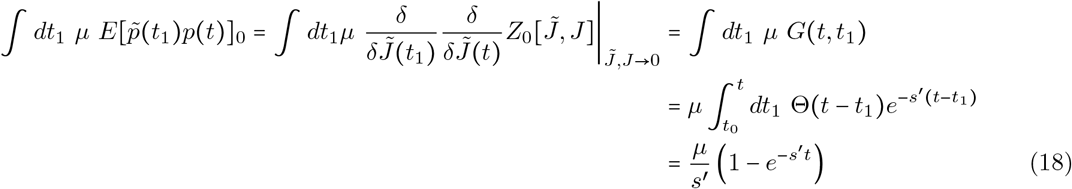

The bilinear nature of the free action *S*_0_, which is analogous to a Gaussian integral, makes taking moments of any power of 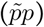 relatively simple; all higher moments vanish such that a path integral of the form 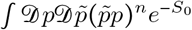 (where each 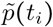 and *p*(*t_j_*) can occur the same or different times *t_i_* and *t_j_*) is equivalent to replacing all possible combinations of 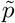 and *p* with *n* copies of the propagator *G*(*t_j_, t_i_*), taking care to maintain the proper time-ordering. Thus, for the example above (the simplest case), the final integral form can be obtained immediately by substituting the dependence on 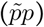 with the propagator, without the need for any intermediate steps. After substituting in the definition of the propagator from Equation 14, the time integration can be performed, and an approximate functional form for the moment of interest can be obtained by summing the results from various contributions.

This integral represents mutational influx occurring at an arbitrary time *t*_1_ (where *t*_1_ ≥ *t*_0_) prior to the time *t* (i.e., the evaluation time, or observation time, for the moment *E*[*p*(*t*)]). Here, *t*_1_ is the *interaction time*, the time when the allele frequency is altered by mutation; the mutation term in the coupling action 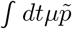 is only dependent on 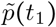 (i.e., is causal, or outgoing with respect to causality) such that a mutation at time *t*_1_ changes the allele frequency at later times (in this case, affecting the value of *E*[*p*(*t* > *t*_1_)]), rather than the reverse. The result in Equation 18 (i.e., the relationship between the mutational influx and the mean frequency *E*[*p*]) is a consequence of the time ordering imposed by free propagator, which applies to all results in perturbative PGFT; for example, this contribution vanishes if the interaction time is manually restricted to the future of the observation time, *t*_1_ ≥ *t*. Since the precise time of the new mutation is variable, the *t*_1_ integral sums contributions over all possible times *t*_0_ < *t*_1_ < *t* when the mutation could have occurred, resulting in changes to *E*[*p*(*t*)] at time *t*. Thus, this contribution represents not just a single path altered by a new mutation that increased the frequency at a specific time, but the sum of all paths with the same structure (i.e., those with a only single mutational event and without additional effects due to the initial condition, drift, or selection at high frequencies). This is clearly not an exact solution for allelic dynamics (drift, initial conditions, etc. are ignored), but instead is an approximation that assumes the allele propagates freely (according to the free action) after mutating once; the mutational influx is thus treated as a perturbation around the otherwise unaltered deterministic dynamics dictated by the functional form of the free propagator. Additionally, this is only one of a number of such contributions that are to be summed to approximate the expectation for the mean frequency at time *t* (for example, paths that take into account the initial condition enter at the same order).

Perturbative field theory makes use of the fact that there is an overall multiplicative constant for any given contribution that dictates its relative importance to the total quantitative value of a moment. If the population parameters *s, μ*, and 2*N* are constant in time, this overall constant is proportional to the product of the coupling constants for each involved interaction (this is equal to *μ* in the above example, as is evident from Equation 17 without computing any results). This is subsequently divided by some power of *s*′, the decay rate (or mass) of a freely propagating allele, where this power counts the number of explicit integrals computed over intermediate times *t* > *t_i_* > *t*_0_ (each *t_i_* integral pulls down a factor of 1/*s*′ from the propagator). For example, the constant of interest for the contribution in Equation 18 is *μ/s*′, where exactly one factor of 1/*s*′ appeared when the integral over the interaction time *t*_1_ was evaluated (this can again be seen without direct calculation); in contrast, the initial condition interaction *∫ dtp*_0_*δ*(*t – t*_0_) does not result in this dependence, as the interaction time is fixed at *t*_0_ (integration over a delta function does not result in a factor of 1/*s*′). This constant can be used to determine if this contribution is ‘technically relevant’ to the moment (i.e., whether or not it is of a comparable order of magnitude to the largest contribution and is thus an essential part of the roughest approximation). Since there are so many coupling constants in the action in Equation 15, it is worth mentioning that it is possible for the coupling constants to impact the dynamics asymmetrically, implying the expansion in one particular constant *c*_1_ must be computed to higher order than the others (e.g., when 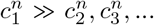 for all *n*). For simplicity, I will ignore this complication and (very roughly) assume that 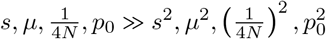, such that each coupling constant is roughly equally perturbatively small. In general, integral contributions to a given moment containing a large number of coupling constants (e.g., a prefactor of 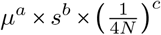 where *a* + *b* + *c* ≫ 1) usually represent subdominant terms that can be ignored. However, these statements are imprecise because the dependence on *s*′ plays an important role in rescaling the coupling constants, as *s*′ is pulled down a different number of times for each interaction (depending on the number of fields 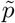 and *p* involved in each interaction); this can be seen in the calculations presented in Results.

Though I have made use of the simplest possible example, the same procedure can be applied to far more complicated contributions to arbitrary moments. The only complication comes in the form of the prefactor enumerating the symmetry of a given term, which is derived from evaluation of the number of possible Wick contractions: the number of ways to replace 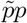 with the propagator while still describing the same time-ordered dynamics. A detailed discussion of Wick contractions (and a vast overview of the technical aspects of field theory) can be found in Peskin and Schroeder [60]. Additionally, Appendix A2 details the enumeration of Wick contractions for PGFT in the diagrammatic context.

Having throughly confused the reader at this point, it is now important to reiterate that this is the *hard way* of computing moments using perturbation theory; Feynman diagrams were invented to render this procedure largely moot by substantially simplifying the task at hand. While this is necessary background for understanding the construction of Feynman diagrams and their associated rules, direct evaluation using functional derivatives is usually too tedious for practical use, especially for particularly complicated coupling actions like Equation 15.

### Feynman diagrams for population genetics

Feynman diagrams, introduced by Richard Feynman in 1949 [70], a scant two years before Motoo Kimura’s first description of a diffusion model of population genetics [71], are graphical representations of perturbation theory used to compute quantities of interest from a path integral that cannot be integrated exactly (i.e., those describing all but the simplest theories). These can be viewed as cartoons of contributing paths, a mnemonic for the assembly of non-vanishing integral contributions, and at the same time a rigorous graph theoretic framing of a calculation when used in conjunction with a set of well-defined rules. For the present purposes, *directed* (i.e., time ordered) Feynman diagrams for the PGFT will be used as a means to quickly compute relevant contributions to the first few moments of the time-dependent allele frequency distribution, including in several specific non-equilibrium contexts. Each diagram is constructed from a set of *vertices* (representing interactions between the allele frequency and selection, mutation, drift, or the initial conditions) and edges (representing allele frequency propagating freely through time) connecting these vertices. The diagrammatic approach resembles the same conceptual steps as the manual perturbation theory described by Equations 16–18: edges represent the free propagation of allele frequency through time, and vertices represent perturbative interactions between, e.g., drift and allele frequency that interrupt this propagation.

#### Vertices representing selection, mutation, and drift

The zoology of interactions in PGFT can be drawn with the following vertices and their associated definitions. Each vertex represents a term in the coupling action shown in Equation 15 and describes an interaction between allele frequency and the initial conditions, mutation, selection, or drift at a given time *t_v_*. For each vertex there is an implicit time ordering represented by some number of outgoing (i.e., causal) allele frequencies 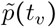 and incoming (i.e., affected) allele frequencies *p*(*t_v_*) that interact in some way at time *t_v_*. Now the meanings of ‘incoming’ and ‘outgoing’ allele frequencies become clearer: they represent the direction of the arrow of time with respect to the vertex time *t_v_*, the time at which the interaction represented by the vertex occurs (i.e., incoming frequencies propagate into the vertex from an earlier time, outgoing frequencies propagate away from the vertex towards later times). The higher order (in 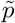 and *p*) a given term in the coupling action is, the more (incoming and outgoing) allele frequencies are drawn to represent connections to propagators from other vertices in the past (for *p*) or future (for 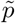). The relevant coupling vertices for a single allele, additive PGFT are enumerated as *V*1 – *V*5 below.

The first vertex represents the effect of the initial condition (i.e., *p*(*t*_0_) = *p*_0_) on the allele frequency. Unlike all other vertices in this theory, the initial frequency vertex occurs at a specific time (enforced by the delta function at *t* = *t*_0_).

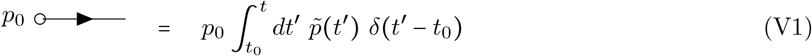

This vertex has a single connection to an outgoing allele frequency 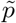, no dependence on *p*, and represents the above integral. Mutational influx, which is linear only in 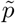 in the coupling action, is represented by a vertex of a similar form.

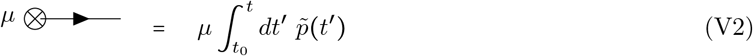

The vertices representing the initial frequency and mutational influx, denoted by an empty circle and crossed circle, respectively, can technically be combined due to the similar linear dependence on 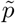. However, since these vertices represent distinct phenomena and will be used in various combinations to create distinct diagrams, I will maintain them as separate vertices for clarity throughout. I will refer to both as *terminal* vertices, as they are the only vertices that have only a single connection; both represent sources of allele frequency (and causality).

The dominant impact of purifying selection on the dynamics is due to the bilinear dependence on *p* and 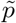 in the free action in Equation 10, which characterizes the exponential reduction of allele frequency over time. Thus, the linear effect of selection on allele frequency is contained in the propagator and will be addressed shortly. In contrast, the effect of selection near the fixation boundary (i.e., when allele frequency is high) is represented by the quadratic dependence on *p*, along with a linear dependence on 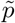, that appears in the coupling action. The latter effect is represented by the following vertex.

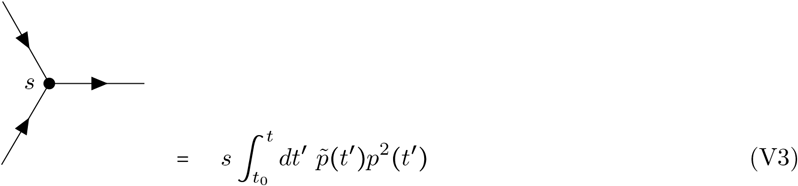

The black dot here represents the selection vertex with coupling constant *s*. The selection vertex represents an interaction between two incoming connections to *p* and one outgoing connection to 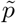. Following the time ordering from left to right, this represents a *narrowing* vertex [55] (the only one in PGFT) and has associated topological properties (i.e., the set of allowed diagrams is constrained by this vertex).

Genetic drift is incorporated as two separate vertices, one representing fluctuations due to low frequencies proximal to the extinction boundary (at zero frequency) and another representing fluctuations near the fixation boundary (at frequency one). The former is represented by the term proportional to 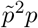 in the coupling action.

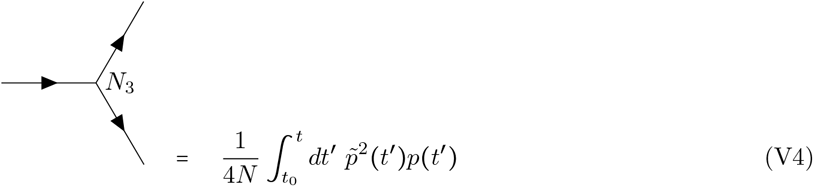

In contrast to the selection vertex V3, this is a *diverging* vertex [55] and has topological conditions associated with this fact. Given the three connections, I will refer to vertex V4 as the *N*_3_ vertex. The second drift vertex, which will be referred to as the *N*_4_ vertex (V5), takes the following form with four connections due to the dependence on 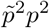.

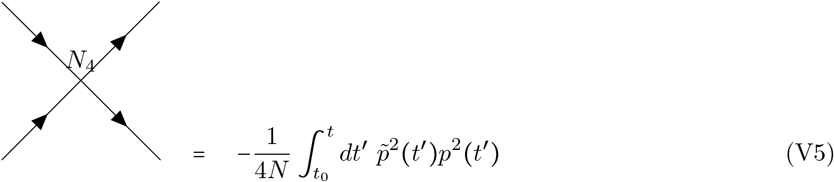

Note that this is the only vertex with a negative coupling constant – 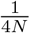, stemming from the quadratic part of the *p*(1 – *p*) dependence in Equation 5. The common feature of the two drift vertices is the quadratic dependence on 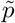, which can only stem from the stochastic contribution in the derivation of the action (see Appendix A1). Thus, any vertex with two outgoing connections inherently represents the contributions of stochastic fluctuations to the allele frequency dynamics.

Finally, there are *external* vertices that represent couplings to the arbitrary source function 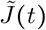 (couplings to *J*(*t*) will not be discussed here). Recall that moments of the allele frequency distribution *E*[*p^n^*(*t*)] are computed by applying functional derivatives 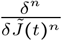 to the partition functional; these functional derivatives are defined at time *t* and resulting moments are evaluated (or observed) at this time after taking the limit 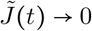 to remove the dependence on the arbitrary function. The role of 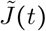 is to impose the time of evaluation *t* and to enumerate the appropriate number of simultaneously evaluated allele frequencies pulled out of the action by the functional derivatives (the power *n* in the moment *E*[*p^n^*(*t*)]). Analogously, external vertices represent properties of the field being measured at time *t* (with no remaining dependence on the coupling 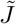). If the moment of interest is the mean frequency *E*[*p*(*t*)] (i.e., *n* = 1), each contributing diagram will have a single external vertex (i.e., diagrams for *n* = 1 all have one external vertex at time *t* connected to some internal vertex at time *t_v_* by a single edge that denotes propagation *t_v_* → *t*).

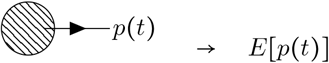

The shaded circle here represents some combination of internal vertices (connected by edges) for a given diagram. The edge propagating to the externally evaluated frequency *p*(*t*) represents the allele frequency distribution exiting the diagram; the termination point of this edge at time *t* is considered an external vertex. For the second moment *E*[*p*^2^ (*t*)], there are two external vertices.

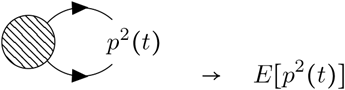

Similarly, all contributing diagrams representing the *n*^th^ moment *E*[*p^n^*(*t*)] have *n* external vertices, each connecting some internal vertex (at some time *t_v_* < *t*) to the external vertex at the evaluation time *t*. External vertices in the above diagram are shown as coincident (i.e., all pointing to *p*^2^(*t*)), which is an appropriate visualization given their meaning, but this will be implied for all external vertices appearing in diagrams below; external vertices are evaluated collectively at time *t*, regardless of their orientation.

Henceforth, the vertex annotations (e.g., *N*_3_, *s*, etc.) will be removed from the vertices in diagrams below, but the above section can be referenced to clarify the interactions represented by any given vertex. Each evolutionary force can be recognized by the corresponding style of circle on the vertex: an empty circle for the initial condition, a crossed circle for mutation, a black circle for selection, and the absence of a circle for drift.

#### Diagrammatic representation of the allele frequency propagator

As mentioned above, the propagator connects vertices in time. A complete diagram is formed when all vertices are connected to edges representing propagators. The way in which this is done can lead to a variety of diagram topologies, each of which represent a distinct contribution to the moment of interest. Each edge (i.e., propagator) between time points *t*_1_ and *t*_2_ will be represented as follows.

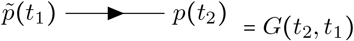

Here, *G*(*t*_2_, t_1_) is the free propagator defined in Equation 14, which strictly enforces the time ordering (i.e.,the causal relationship *t*_2_ > *t*_1_) via the Heaviside function such that the propagator vanishes for any value *t*_2_ ≤ *t*_1_. As mentioned above, *G*(*t_i_, t_j_*) = 0 corresponds to the Itô condition [55], which has implications for the diagrammatic rules presented below. As a result, the arrow appearing on each propagator should be interpreted quite literally as the flow of time from an earlier time point *t*_1_ to a later time point *t*_2_. Note, I have chosen to draw all vertices and the propagator from left to right with increasing time, but this is solely for visual clarity; the flow of time is defined by the arrows on the propagators, not the orientation of a given diagram.

#### Feynman rules for population genetics

To make use of this diagrammatic approach, one must establish rules for interpreting each diagram as the associated integral contribution to moments of interest. These rules allow one to replace the algebraic manipulation and evaluation of perturbative contributions shown in Equations 16–18 with a series of diagrams representing the same contributions. The Feynman rules for population genetic field theory diagrams are as follows (see [55] for a discussion of the rules for other example SFTs).

- <u>Rule 1:</u> For each vertex in a diagram multiply by the appropriate coupling constant. These are *p*_0_ for V1, *μ* for V2, *s* for V3, 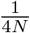 for V4, and 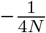 for V5.
- <u>Rule 2:</u> For *C* copies of each *internal* vertex in a given diagram, multiply by a factor of 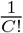 (this does not apply to the external vertices in this theory, and doing so will produce incorrect results for higher moments).
- <u>Rule 3:</u> Count the number of diagrams with identical topology that can be made by connecting edges (i.e., propagators) between the same vertices. This number is an overall *counting factor* of *T* for the number of diagrams with the same topology. This can also be computed by finding the number of unique ways to replace pairs of 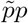 with a propagator when writing out the diagram in the form shown in Equation 17 (i.e., the number of unique *Wick contractions*). A description of the counting factor in terms of Wick contractions is given in Appendix A2, along with detailed examples. Note that this quantity is related to the ‘symmetry factor’ used to compensate for over-counting contributions in QFT [60, 61], but it is used here as an overall multiplicative factor (rather than dividing by T) for the same purpose, consistent with the formulation described in [55].
- <u>Rule 4:</u> Multiply by a factor of the propagator *G*(*t*_2_, *t*_1_), defined in Equation 14, for each edge connecting a vertex at *t*_1_ to a vertex at a later time *t*_2_. For propagators from the initial condition vertex V1, *t*_1_ is set to *t*_0_; for propagators exiting the diagram (i.e., connected to an external vertex), *t*_2_ is set to the evaluation time *t*.
- <u>Rule 5:</u> Any diagram containing a propagator that starts and ends at the same vertex (i.e., *G*(*t_v_, t_v_*)) has an overall factor of zero (i.e., does not contribute; *G*(*t_v_, t_v_*) ∞ Θ(0) = 0 disallows self-connections). Similarly, any diagram that contains a propagator going backwards in time has an overall factor of zero. In other words, all propagators in diagrams representing non-vanishing contributions must be *time-ordered* (i.e., all edges must flow from early to late time, independent of the vertices involved).
- <u>Rule 6:</u> Integrate over the time *t_v_* at which each vertex in the diagram could have occurred, between the initial time *t*_0_ and the final time *t*. Note that the Heaviside functions in the propagators will enforce the appropriate time ordering, so the integration can be performed from *t*_0_ to ∞, if desired (this produces equivalent results and may be easier in some cases).

### Heaviside integral evaluation in Mathematica

All integrals over Heaviside functions were either performed manually and confirmed using Mathematica [72] (version 12.3.1) or computed directly in Mathematica for accuracy. The relevant Heaviside integrals represented by each Feynman diagram are written out explicitly in Supplemental Material S3; Supplemental File S1 contains a Mathematica notebook evaluating each integral for the specific examples of constant sized, exponentially expanding, (square) bottlenecked, and cyclical populations described in this manuscript.

### Non-equilibrium simulations

All simulations were performed using `neqPopDynx`: Non-equilibrium Population Dynamics simulator (version 1.5), a custom-written Wright-Fisher simulator scripted in Python 3 for this manuscript (see Data Availability for download information). Allele frequencies evolve independently in the infinite recombination limit to assess properties of the allele frequency probability distribution and are subject to user-specified rates of mutation and back mutation rates (allowing for *μ, μ_b_* > 1/2*N*), selection and dominance coefficients, initial population size, and a choice of pre-specified demographic changes in the population size with specifiable parameters (e.g., growth rate of exponential expansion, bottleneck start time, duration, and diploid population size, etc.). Initial population sizes are specifiable as a number of diploid individuals *N*_0_, but alleles evolve in haploid form (dominance, if non-additive, is imposed via allele frequency under the assumption of Hardy-Weinberg equilibrium). Selection is imposed on the expected frequency (at the current generation *t* + 1) with the following definition.

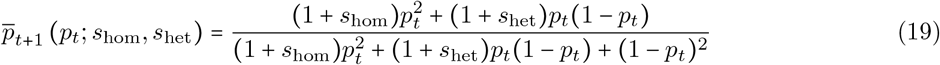

Here, the allele frequency *p_t_* is defined at the end of the previous generation *t* as *p_t_* = *n_t_*/2*N_t_*, where *n_t_* is the allele count and *N_t_* is the diploid population size for generation *t*. Drift is imposed by binomially sampling a number of counts from a haploid sample size of 2*N*_*t*+1_ (the population size of the current generation) around 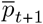.

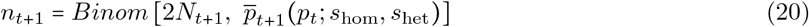

The mutational process is imposed by Poisson sampling *n_μ_* = *Pois*(*μ*(2*N – n_t_*)) mutations and *n_μ_b__* = *Pois*(*μ_b_n_t_*) back mutations. The total counts *n*_*t*+1_ = *n_t_ + n_μ_ – n_μ_b__* are then restricted to the range *n*_*t*+1_ ∈ [0, 2*N*_*t*+1_] such that mutations outside that range are assigned to extinction (*n*_*t*+1_ = 0) or fixation (*n*_*t*+1_ = 2*N*_*t*+1_), if necessary. This procedure allows for both recurrent mutation and ‘un-fixation’ (i.e., back mutation of a fixed allele). Finally, the population size is altered according to the user-specified choice from the following pre-coded demographies: `equilib, exponential, bottleneck, oscillating`, and several other options not relevant to this manuscript. The equilibrium demography maintains a constant population size at the initial diploid size *N*_0_. The exponential demography models exponential population growth with a specifiable growth rate Γ and initial population size *N*_0_ such that for generation *t* the population contains *N_t_* = *N*_0_*e*^Γ*t*^ (rounded to the nearest integer) diploid individuals. The bottleneck demography models a ‘square’ bottleneck that instantaneously reduces in population size and instantaneously returns to the initial population size at a later time. This demography has specifiable parameters for the initial diploid population size *N*_0_, the generation when the bottleneck begins *T_i_*) (not to be confused with the initial condition defined at *t*_0_ above), the duration of the bottleneck *T_b_*, and the number of diploid individuals during the bottleneck *N_b_*. The oscillating demography models a cyclical population with regular oscillations in population size using the parameterization defined in Equation 73 (rounded to the nearest integer), which is specific to this manuscript. This parameterization allows for specification of the maximum population size *N*_max_ (where the initial population size is fixed by *N*_0_ = *N*_max_), the minimum population size *N*_min_, and the period of oscillation *τ*. `neqPopDynx` allows for a burn-in period (using the `--burnin` option) of a specified number of generations prior to the beginning of the demography that can be used to equilibrate the population, but no burn-in was used for this manuscript due to a focus on the dynamics of the equilibration process itself. Finally, the initial frequency *p*(*t*_0_) can be specified using the `--initcount` command as a count between 0 and 2*N*_0_. This models a delta function initial condition and is applied to all sites to track the temporal evolution of the distribution of frequencies.

Each run simultaneously simulates *L* independently evolving alleles for a total of `gen` generations through the specified time-dependent demography. For the first ten generations and every `printgen` generations thereafter, the following moments are (optionally) computed by averaging over L allele frequencies: the first five non-central moments (with the command `--moments`), the first five central moments (`--central`), and the first five cumulants (`--cumulants`); all fifteen moments can be reported with the command `--recordall`. This generates a temporal output of the first several moments for comparison to the analytic results presented in this manuscript. To evaluate the analytic approximations generated in this manuscript, *L* = 10^5^ sites were simulated separately for each demography of interest (see parameterizations above), for each of two mutation rates, for each of nine selection coefficients: six selection coefficients were used for all demographies *s*_het_ = 10^-1/2^, 10^-1^, 10^-2^, 10^-3^, 10^-4^, 10^-5^ under the assumption of additive selection (i.e.,co-dominance with *h* = 1/2) such that *s*_hom_ = 2*s*_het_. Note that, since additive selection is assumed throughout this manuscript, I have used the convention *s* = *s*_het_ in all analytic formulae (in contrast to the simulator input which uses the diploid definition *s* = *s*_hom_ = 2*s*_het_, though *s*_het_ may be defined directly). The remaining three selection coefficients were chosen to interrogate the breakdown of the analytic approximations, which occurs in a demography-specific parameter regime: coefficients of 2*Ns*_het_ = 3, 5, 10 were used for populations with constant size; 2*N*_0_*s*_het_ = 3, 5, 10 were used for the exponentially growing populations, where *N*_0_ is the initial population size; 2*N_b_s*_het_ = 3, 5, 10 were used for bottleneck demographies, where *N_b_* is the population size during the bottleneck; and 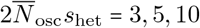 were used for cyclical populations, where 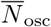 is the effective population size defined as the harmonic mean of the population size *N*(*t*) over one period of oscillation. Mutation and back mutation rates were set to *μ* = 10^-6^, *μ_b_* = 10^-8^ and separately *μ* = 10^-8^, *μ_b_* = 10^-10^ per individual per generation for all demographies (i.e., two mutation rates were simulated for each demography, where *μ_b_* = *μ*/100 was used in all cases to reduce the parameter space). The total number of generations used for each simulation was chosen from `gen` = 2 × 10^3^, 10^4^, 10^5^, 2 × 10^5^ generations to suit the relevant dynamics (e.g., very rapid exponential growth was only simulated for a modest 2000 generations to focus on semi-realistic scenarios). Moments of interest were recorded every generation for the first 10 generations and then every 10 subsequent generations to produce robust temporal output for each simulation.

Simulations were run across a broad range of selection, mutation, and demographic parameters to assess the analytic results presented in this manuscript; the majority of these simulations are presented in Supplemental Material S4 in Figures *S*1–*S*44, but some minor discretion was used to limit redundancy and the total number of plots by omitting plots produced with duplicate parameters, except with respect to mutation rate (in all cases the more relevant mutation rate was chosen to highlight any notable observations). For each parameter combination, dynamics of the allele frequency probability distribution were tracked by densely outputting the first five cumulants and first five non-central moments, estimated by averaging over *L* = 10^5^ independently evolving sites, and tracked over time for between 2 × 10^3^ and 2 × 10^5^ generations to estimate time-dependent trajectories for each quantity. A total of 540 distinct parameter combinations were simulated: each demographic parameter was varied over a range of 1–3 orders of magnitude (selection coefficients were varied over 5.5 and mutation rates over 2 orders of magnitude; see details above) to systematically explore the parameter space of demographic scenarios. Simulations for each parameter combination were initialized with a delta function at generation *t* = 0 by setting the initial frequency *p*(*t* = 0) = 1/2*N*_0_ (i.e., one derived allele in a single haploid individual in the initial population 2*N*_0_ at generation *t* = 0) and the subsequent non-equilibrium dynamics of the allele frequency probability distribution were output. Exploration of each parameter combination provided a means of assessing the regime of validity for the analytic approximations presented herein for a number of non-equilibrium scenarios of interest. Supplemental Material S4 presents plots spanning roughly 60% of the explored parameter space (see Figures S1–S44). The equilibrium demography was simulated with population sizes *N* = 10^2^, 10^3^, 10^4^; all parameter combinations (*N, μ, s*) are presented in Supplemental Material S4.1. The exponential demography was simulated over all combinations of three initial population sizes *N*_0_ = 10^2^, 10^3^, 10^4^ and growth rates Γ = 10^-2^, 10^-3^, 10^-4^ per generation; plots are presented in Supplemental Material S4.2 for *μ* = 10^-8^ with all combinations of (*N*_0_, Γ, *s*) and for *μ* = 10^-6^ with all combinations of (*N*_0_ = 10^4^, Γ, *s*). The bottleneck demography was run with the initial size *N*_0_ = 10^4^ and all combinations of the bottleneck size *N_b_* = 10^2^, 10^3^, bottleneck onset time *T_i_* = 10^3^, 10^4^, and bottleneck duration *T_b_* = 10^2^, 10^3^; Supplemental Material S4.3 presents results using *μ* = 10^-6^ to generate all figures due to the subtlety of the time-dependent change for many quantities of interest (*μ* = 10^-8^ results in inherently noisier plots, as discussed in Supplemental Material S4.1, effectively masking subtle changes in the simulated trajectories) and contains plots of all parameter combinations of (*N*_0_ = 10^4^, *N_b_, T_i_, T_b_, s*). Cyclical demographies were simulated using the maximum population size *N*_max_ = 10^4^ and all combinations of the minimum population sizes *N*_min_ = 10^3^, 5 × 10^3^ (i.e., 10× and 2× change in population size over each period, respectively), and oscillation periods of *τ* = 10^2^, 10^3^, 10^4^ generations; Supplemental Material S4.4 presents plots with *μ* = 10^-6^ for all combinations of (*N*_max_ = 10^4^, *N*_min_, *τ, s*) and with *μ* = 10^-8^ for all combinations of (*N*_max_ = 10^4^, *N*_min_ = 5 × 10^3^, *τ, s*) for all simulated periods.

## Results

There are two distinct classes of non-equilibrium behavior presented in this manuscript. First, the dynamics of equilibration from a known initial frequency is discussed, describing the approach to the standard equilibrium frequency distribution. Second, I will consider non-equilibrium demography and describe the associated dynamics of the allele frequency distribution for specific examples of particular interest in population genetics: bottlenecks, exponential growth, and population size oscillations (e.g., seasonal variation).

### Equilibration from a known initial frequency

While the equilibrium distribution of allele frequencies has a well known steady-state (pseudo-)equilibrium description [33], the approach to equilibrium from an arbitrary initial frequency is a topic of less discussion in the literature, outside of simulation studies; in the latter context, it is understood that a ‘burn-in’ period is needed to erase transient effects that remain from the arbitrary choice of initial condition. This alone should motivate careful study of the equilibration process, as it is of practical relevance to the optimization of simulation techniques, which may want to minimize the necessary burn-in period (e.g., to choose the pre-computation time as a function of the selection coefficient). However, equilibration is also of empirical interest when the frequency of an allele at a given time is known (or can be inferred). For example, with the advent of ancient DNA sequencing it has become possible to estimate the historical allele frequency in an ancient population using contemporaneous geographically proximate samples (resources like, e.g., the Ancient mtDNA database [73] contain a growing number of such ancient population samples, there for mitochondrial DNA). This can then be used as a starting point to model the expected distribution of frequencies today, varying any number of population parameters (e.g., for inference of selection or demographic properties). While this can be accomplished via simulation, working with an analytic model of equilibration (or non-equilibrium trajectories) allows for the construction of an explicit likelihood from first principles. A second case when the initial frequency is known is more ubiquitous: all new mutations enter the population at frequency 1/2*N*(*t_μ_*) (or slightly above if recurrent), which provides an initial frequency at some time *t_μ_* (i.e., the approximate date of origin) if the population size history *N*(*t*) and age of the allele can be jointly inferred. In some demographic histories, one can even assume that the population size is roughly constant on short time scales (which is more accurate for stronger selection) such that the dynamics immediately after the emergence of a novel allele temporarily equilibrate to the mutation-selection-drift balance described by [33], rather than taking a more general non-equilibrium path. Additionally, the assumption of constant population size provides an opportunity to apply the field theoretic machinery developed above to the simplest case and yields a null model for comparison to the dynamics in non-equilibrium demographies (discussed in detail below). In this section, I focus on this scenario, the transient equilibration of an allele in a finite population with a known initial state and constant diploid population size *N*. Here, the model consists of a single mutation originating at a frequency *p*_0_ at time *t*_0_ (which will be set to zero) and the equilibration period is studied in the form of the first few moments of the probability distribution as time progresses to some final time *t*. With a sufficient number of moments considered (the first four are explicitly computed herein), this provides a full description of the temporal evolution of the probability density of the allele frequency.

The dynamics of an allele in a constant sized population can be computed using the field theoretic formalism described above, but perturbative field theory makes an implicit assumption about the relative values of coupling constants, which restricts the regime of validity of all results. For the perturbative PGFT partition functional in Equation 16, selection must remain efficient with 2*Ns*′ > 1 (which reduces to 2*Ns* > 1 when *s* ≫ *μ, μ_b_*) because all terms in the coupling action must remain subdominant to the free action to truncate the Taylor expansion at a finite point (i.e., the series must converge). However, it is unclear at what value of *s*′ the perturbative theory breaks down due to the additional dependence on frequency (i.e., the relevant selection term 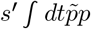 is linear in 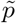, while the relevant drift term 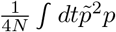 has a quadratic dependence on 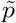). One might choose to proceed under the assumption of very strong selection 2*Ns*′ ≫ 1 and compute more limited results by ignoring higher order corrections entirely. A more precise cutoff for an approximation is partly dependent on how many terms in the Taylor series in *S_c_* are considered before truncating the rest. Computing corrections to the lowest order approximation (i.e., the first non-vanishing contributions to a given moment) by adding higher order contributions extends the regime of validity due to a more accurate dependence on the perturbative parameter 1/2*Ns*′. The same is true of the diagrammatic formalism: computing higher order diagrams provides a more accurate approximation and can thus account for comparatively weaker selection coefficients – up to a point. That said, the series in its current form will never converge for sub-drift selection coefficients 2*Ns*′ ≤ 1 and approaching the drift limit from above requires successively more contributions for the same accuracy. Treating sub-drift dynamics requires either a different version of perturbative field theory (e.g., a theory that results in an expansion in orders of 2*Ns* rather than 1/2*Ns*; see [51]), some means of evaluating an infinite number of terms (see discussion of the adiabatic expansion in [55]), or a non-perturbative technique to evaluate the full partition functional defined by the action in Equation 6. For the perturbative field theory presented here, I will primarily be concerned with the point at which the lowest order approximation breaks down, rather than attempting to obtain more accurate approximations, but will compute the first subdominant corrections (in the form of more complicated Feynman diagrams) for the lowest moments to clarify the procedure. Thus, all results presented below assume alleles evolve under efficient selection, but this statement will become more quantitative along the way and simulations will be used to confirm a reasonable cutoff for the regime of validity.

#### Temporal evolution of the mean frequency *E*[*p*(*t*)]

The temporal evolution of the mean frequency *E*[*p*(*t*)], henceforth 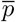 for notational convenience, is of particular interest as it describes the dynamics of the mutation burden and thus the mutation load under additive selection. PGFT can be used to find the time dependence of the mean frequency; the lowest order dynamics can be obtained relatively easily by using the formal perturbative analysis described above (i.e., by applying functional derivatives to the partition functional; Equation 18 shows one such contribution). I will instead focus on the diagrammatic series expansion as it is both visually clearer and easier to compute for the general moments presented below. However, understanding the mean frequency dynamics of equilibration under strong selection does not require a complicated framework. The first moment o the Langevin equation shown in Equation 5 reduces to a deterministic equation when selection is strong, as alleles are maintained at low frequency due to selection. In this regime, the equation can be approximately linearized in the same way as for Equation 3 (recall the linear operator 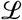), as corrections due to drift are largely negligible. Under strong selection, the mean frequency dynamics are well-approximated by the following equation.

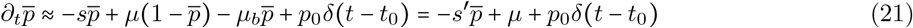

Note that this is identical to the linearized version of Equation 3 (with *p*_det_ replaced with 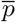) including the homogeneous linear operator 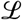. This approximation is valid if the initial frequency is well below fixation and deleterious selection is sufficiently strong to suppress the quadratic frequency dependence at all times. Since the equation is the same, the solution has the same functional form as Equation 4, which is written explicitly below for 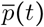. Henceforth, the initial time will be set to *t*_0_ = 0 for convenience.

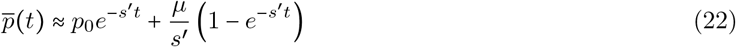

This is again expressed in terms of the effective selection coefficient *s*′, which drives the deterministic dynamics of exponential decay. The following additional variables defined in terms of *s*′ will become convenient for representing the perturbative parameters: the mutation-selection balance parameter *λ*′ and the population-scaled effective selection parameter *γ*′, where the prime denotes that they are defined in terms of *s*′ (rather than *s*).

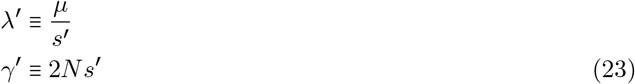

As with *s*′, these combined parameters approach their normal definitions when *s* ≫ *μ, μ_b_*, where *λ*′ → *λ* ≡ *μ/s* and *γ*′ → *γ* ≡ 2*Ns*. The equilibration of the mean frequency described in Equation 22 is easily interpreted: the mean exponentially decays away from the initial frequency 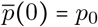 on a timescale of order ~ 1/*s*′ and approaches the mutation-selection balance frequency 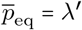 on the same timescale. At asymptotically late times (*t* → ∞), this equilibrium frequency dominates and persists.

In PGFT, the same result is obtained from the sum of the lowest order diagrams for the mean frequency. Both diagrams contain only a single internal vertex, either the terminal vertex representing the initial frequency V1 or the mutational influx terminal vertex V2, and a single propagator connecting the internal vertex to an external vertex. These diagrams are drawn as follows. To differentiate them from equations, diagrams will be numbered separately from equations using *DX* where *X* the appropriate number.

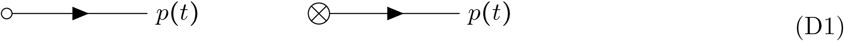

The two diagrams shown (left and right, above) represent separate contributions to the mean frequency 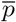 and can be calculated using the Feynman rules to find the appropriate time dependence. Here, the external vertices have been labeled with the field *p*(*t*) to indicate the quantity exiting the diagram, but this is implied for all external vertices by the lack of internal vertex decoration (black dot, crossed circle, etc.) and will be dropped henceforth. Contributions to each moment represented by a given diagram are labeled with subscripts denoting the diagram number DX and their placement (left, center, right in Diagram *DX*) for clarity (e.g., contributions to 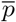 from the left diagram in D1 are labeled as 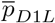, where *L, C*, and *R* denote left, center, and right diagrams, respectively).

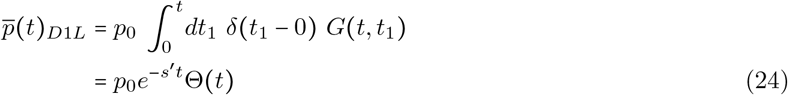

Since the Heaviside function Θ(*t*) = 1 for all positive values of time *t* > 0, the final dependence on Θ(*t*) can be dropped and will not be shown in subsequent results. The contribution from the right diagram in D1 is computed in the same way.

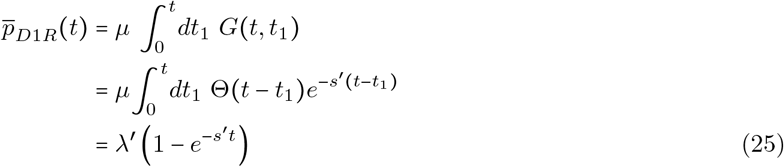

The sum of these two contributions provides the lowest order time dependence for the mean frequency. Higher order corrections are guaranteed to be 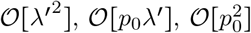 (i.e., involve at least two vertices V1 and/or V2), or are dependent on some other perturbative parameters (e.g., 1/2*Ns*′ in diagrams containing the drift vertices V4 and V5). Here, the notation 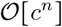 is used in the usual context to denote a sum of terms that are of order *n* or higher in the perturbative parameter *c* (e.g., *λ′, p*_0_, etc.) and suppressed by a series of powers of *c*. Under the assumption that each perturbative parameter *c* ≪ 1 converges at roughly the same rate (see Materials and Methods on perturbation theory), I will henceforth annotate the truncation point for a series of diagrams by the minimum number of internal vertices in all additional non-explicit diagrams: contributions from n-vertex and higher-vertex order are denoted 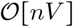, where *n* is the number of *internal* vertices (i.e., V1–V5) in the next diagram in the series, as each vertex contributes a factor of the appropriate coupling constant (proportional to a perturbative parameter) according to Feynman <u>Rule 1</u>. The one-vertex approximation for the mean frequency, which is given by the sum of contributions from all allowed one-vertex diagrams (i.e., the two shown in Diagram D1), is as follows.

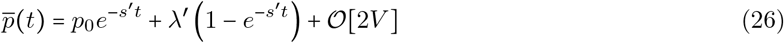

As discussed in Materials and Methods, the effective selection coefficient *s*′ is combined with the coupling constants to rescale the overall prefactor, changing the relevant parameter controlling the convergence of the series of diagrams from *μ* to *λ*′ = *μ/s*′ in the above expression. This factor appears only on the mutational-influx diagram (Diagram D1*R*) because the internal vertex time *t_v_* = *t*_1_ was integrated over (here, corresponding to contributions from mutations at all possible times *t*_0_ ≤ *t*_1_ ≤ *t*). This is in contrast to the initial condition diagram (Diagram D1*L*) because of the fixed vertex time *t*_v_ = *t*_0_ (i.e., the integral of an exponential over a delta function does not pull down a factor of 1/*s*′). This can also be seen from the result of the brute force calculation shown in Equation 18, which is now clearly equivalent to the contribution represented by the simplest mutational influx Diagram D1*R*. In the same way, each diagrams represents a term in the Taylor expansion of the coupling action and should be interpreted in this context, as well as quite literally as, e.g., mutational influx at some unspecified time *t*_1_ (to be integrated over) contributing to the mean frequency at a later time *t*. This literal interpretation of Feynman diagrams in PGFT is allowed, in part, due to the absence of divergences that often plague field theories (detailed explanations are provided in [60, 61]; Appendix A1.1 includes a brief discussion) and due to Feynman <u>Rule 5</u>, which ensures time-ordering (there is no quantum behavior or propagation backward in time to ponder in this theory).

With respect to the regime of validity, the dependence on perturbative parameters like *λ*′ (or 1/*γ*′), which naturally emerged from perturbation theory, indicates that there is a breakdown in the approximation roughly when any of these parameters is above order 1/10 (i.e., using the rescaled coupling constant *λ*′ as an example, this is the point when *λ*′^2^ is of the same order of magnitude as *λ*′ and represents an order 10% correction). This puts strict limits on the regimes that are well described by the present PGFT, most notably restricting selection to above the order of the drift constant (*γ*′ > 10 or *s*′ > 10/2*N*). I will choose to assume a slightly more generous cutoff of roughly *λ*′ ⪆ 5, representing a 20% correction to the 1/*γ*′ contribution from the order 1/*γ*′^2^ term in the series, a 1/25 = 4% correction from the order 1/*γ*′^3^ term, etc. Later, this proposed cutoff will be assessed via simulations to confirm the dynamics are still reasonably well described at *γ*′ = 5. The remainder of this manuscript will therefore assume strong selection using the rough definition *γ*′ ~ 2*Ns* ≳ 5.

Higher order corrections to the deterministic behavior described by Equation 26 (and Equation 22) can be obtained by enumerating more complex diagrams. First, note that there are no allowed two-vertex order diagrams that contribute to the mean frequency due to topological constraints imposed by the vertices in this theory and application of the Feynman rules; the only two-vertex diagrams with one external vertex that can be assembled from the available vertices include a self-connecting vertex and thus vanish because <u>Rule 5</u> enforces strict time-ordering. The next order contributions are instead represented by the five diagrams shown below, each containing three internal vertices. The first three non-vanishing three-vertex diagrams that have the same topology, but differ in their terminal vertices V1 and V2.

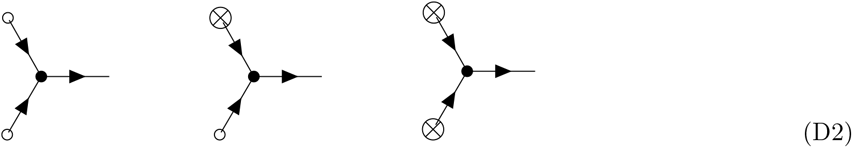

These diagrams are slightly more complex than those in Diagram D1 and not much more difficult to compute. However, they do require counting factors (i.e., Feynman <u>Rule 3</u>) that were unneeded in Diagram D1 (see Appendix A2). In each of these diagrams, there is an overall counting factor of 2 corresponding to the ability to swap the pair of propagators connecting the selection vertex V3 to the terminal vertices without altering the topology of the diagram. Additionally, Diagrams D2*L* and D2*R* each contain one repeated vertex (V1 and V2, respectively), resulting in an overall factor of 1/2! from Feynman <u>Rule 2</u>. The contributions to the mean frequency represented by these diagrams are as follows; for brevity, I will henceforth omit the step of substituting in the definition of the propagator in Equation 14 (but for completeness, the appropriate Heaviside integrals can be found in Supplemental Material S3).

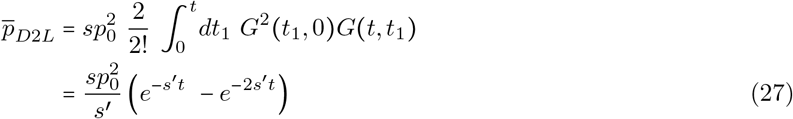

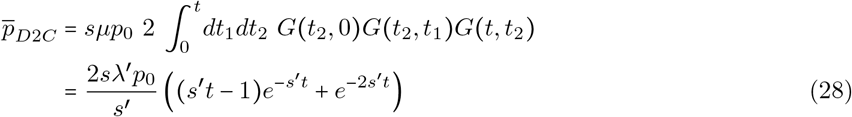

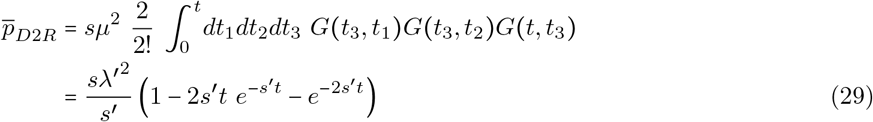

Note that any positive integer power of the Heaviside function is equal to the Heaviside function (i.e., Θ*^n^*(*t*) = Θ(*t*) for 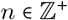); this fact is convenient for evaluating integrals over the propagator when multiple propagators take the same path, which, for example, leads to the *G*^2^ (*t*_1_, 0) dependence in Equation 27). Diagrams that have this simple branching topology are referred to as *tree diagrams* (see [61] for some general properties and applicable techniques). For the mean frequency, specifically, tree diagrams may only contain some number of selection vertices V3 and an appropriate number of initial frequency vertices V1 and/or mutation vertices V2 to terminate edges from each selection vertex. Notably, these are all *deterministic* vertices. In other words, tree diagrams for the mean frequency are independent of drift vertices and thus can only describe the deterministic approach to mutation-selection balance in constant-sized populations. Higher order tree diagrams (e.g., those shown in Diagram D2 and analogous diagrams with more branches) provide small corrections to the deterministic solution in Equation 22, approximating the contribution of the quadratic behavior suppressed to obtain this solution (i.e., the selection vertex describes the quadratic dependence of additive selection from –*sp*(1 – *p*) that appears in the coupling action in Equation 15). The sum of the infinite series of tree diagrams would thus reconstruct the non-linear dependence on selection at high frequencies in the form shown in Equation 3, with *p*_det_ replaced with 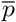, and reproduce the time-dependent solution to this equation (which can be obtained by other means). For some partition functionals, such an infinite series of tree diagrams can be computed in closed form, which forms the basis for a class of perturbative field theories in a distinct limit (see, e.g., [55] for an outline of the general procedure in a different context), but will not be presented in this manuscript.

In contrast to tree diagrams, additional allowed diagrams for the mean frequency that include a dependence on the drift vertices can be drawn, including two diagrams at three-vertex order. These diagrams are referred to as *loop diagrams* due to their topology (i.e., they contain a closed loop, but note that they remain time-ordered). Loop diagrams that contribute to the mean frequency at three-vertex order, shown below, depend only on the *N*_3_ drift vertex V4, which captures the low frequency effects of drift.

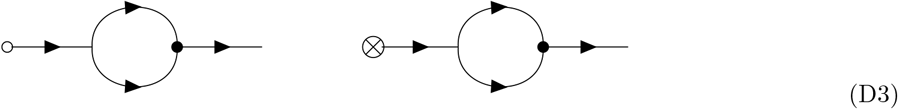

These are examples of one-loop diagrams; diagrams topologies with *n* distinct loops are referred to as *n*-loop diagrams. The particular one-loop diagrams in Diagram D3 provide a clear example of the effect of time ordering. If the selection and drift vertices were reversed (i.e., the *s* vertex connected by a propagator to the terminal vertex and the *N*_3_ vertex connected by a propagator to the external vertex), one propagator would be forced to flow backward in time and the contribution of the entire diagram would vanish due to <u>Rule 5</u>. This indicates that contributions to the mean frequency from substantial drift-driven fluctuations must subsequently be mitigated by the high frequency selection vertex at a later time. Phrased another way, the lowest order effects of drift on the mean frequency are necessarily mediated by natural selection, which is an indirect effect; this statement extends to all effects of drift on 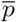 due to the topological constraints imposed by the unique narrowing property of the selection vertex V3 in this theory. The contributions from these diagrams can be computed in the same way as above, but note the counting factor of 2 from the equivalent topology that results from swapping the propagators inside the loop. Additionally, they each contain two propagators that take the same path (i.e., the two propagators that comprise the loop begin and end at the same vertices, which results in *G*^2^ between the drift and selection vertices; this was observed in Diagram D2*L* for a different reason).

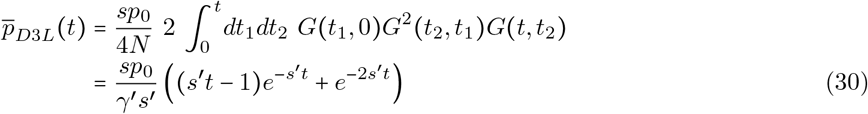

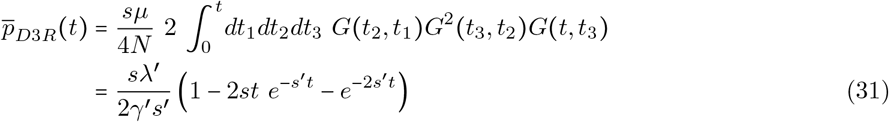

Note that these contributions depend on the previously unseen perturbative parameter 1/*γ*′, which defines the efficacy of selection in a constant population, characterizes the relative importance of drift to the dynamics, and is the primary topic of interest for the extension of PGFT to non-equilibrium demography introduced at a later point. Henceforth, for brevity, I will only show the resulting contribution for each diagram without explicitly writing the propagators. Instead, each relevant integral is presented in detail in Supplemental Material S3. Together with the tree diagrams, these are the only non-vanishing contributions at three-vertex order and can be summed to find the three-vertex approximation for the time dependence of the mean frequency, which provides the largest corrections to the deterministic behavior characterized by Equation 26.

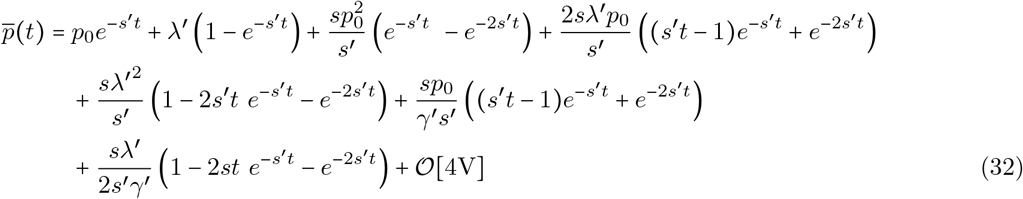

In practice, the values of the perturbative parameters here will generally be small enough to safely ignore even the three vertex contributions, unless the selection coefficient of interest is sufficiently small, the mutation rate is exceptionally high, or the population size is small. However, as *γ*′ → 1, three-vertex and higher order corrections become technically relevant, given that the difference between the leading order and at least one of the sub-leading order contributions is roughly a factor of 1/*γ*′ (assuming that *s/s*′ ≈ 1 to roughly compare the prefactors). One can now assess the accuracy of both Equation 26 and 32 by comparing the analytic approximations for the functional form to simulations (see Materials and Methods) for a range of parameter values: an example set of parameters is shown in Figure 1 (using the three-vertex approximation), along with other moments computed below,. The same moments, including those for the mean frequency, are shown for a large range of parameters (*N, μ, s*) in Supplemental Material S4.1 in Figures S1–S12. As expected, the deterministic approximation (i.e., Equation 26 without the three-vertex corrections) is an exceptionally good description of the dynamics of the mean frequency dynamics that persists until roughly *γ*′ ~ 5, after which it breaks down as expected. The three-vertex corrections add little to the approximation, even near the breakdown of the approximation, until the population size is very small, after which they provide a very modest correction in the correct direction. Thus, the one-vertex approximation in Equation 26 provides a robust approximation for the equilibration dynamics of the mean frequency in a constant sized population and can be used over a wide range of population sizes, provided selection remains efficient.

**Figure 1:**
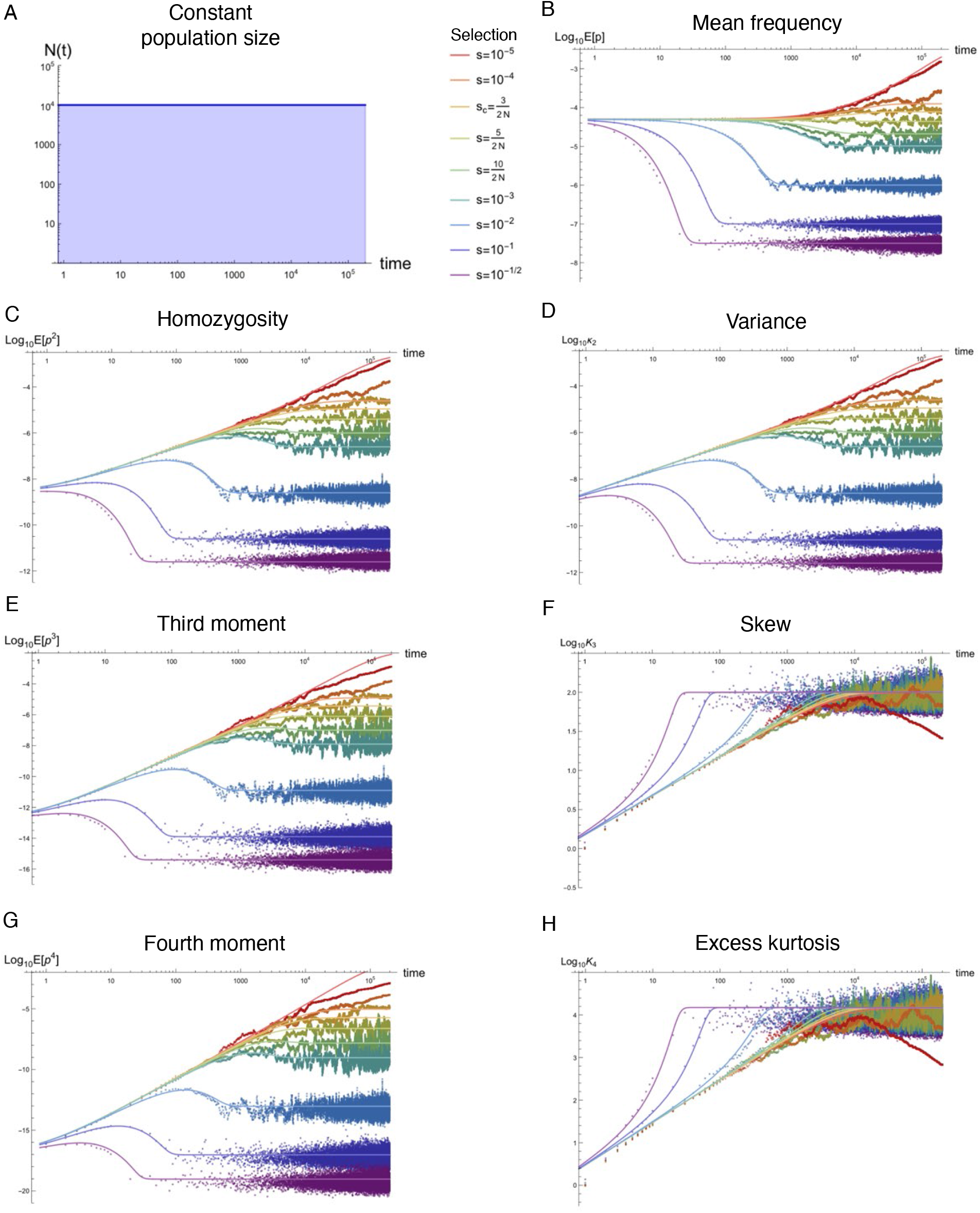
Equilibration of the first four non-central moments, variance, skew, and excess kurtosis in a population of constant size. From top to bottom, the left column of plots show the time dependence of the population size, homozygosity *E*[*p*^2^(*t*)], third moment *E*[*p*^3^(*t*)], and fourth moment *E*[*p*^4^(*t*)] of the allele frequency probability distribution for a population of constant (haploid) size 2*N*. The right column of plots shows the mean 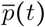, variance *κ*_2_(*t*), skew *K*_3_(*t*) = *κ*_3_/*κ*_2_^3/2^, and excess kurtosis *K*_4_(*t*) = *κ*_3_/*κ*_2_^2^ for the same parameter values. One example set of parameters with diploid population size *N* = 10^4^, mutation rate *μ* = 10^-8^, and back mutation rate *μ_b_* = 10^-10^ is used for all plots; a range of additional parameter combinations are presented in Supplemental Material S4.1 in Figures S1–S12. Solid lines in each plot represent analytic approximations from the text with selection strengths ranging from *s* = 10^-1/2^ (purple) to *s* = 10^-5^ (red), including those near where the approximation first breaks down at roughly *s*′ ~ 5/2*N*. Scattered points represent the results of estimated moments from temporal simulations using `neqPopDynx` with the same parameter values; darker shades of the each color correspond to the analytic curve with the same selection coefficient. *L* = 10^5^ sites were simulated to estimate each moment.

Looking at the functional form in Equation 32, the time dependence of the lowest order contributions in Equation 32 is dominated by exponential decay. However, many of the contributions from three-vertex diagrams involve *transient* terms (i.e., those that vanish at both early and asymptotically late times) with a time dependence of the form (*s′t*)*e^−s′t^*, which describes a maximum at *t* = 1/*s*′ and vanishes in either direction. Although these contributions are quantitatively very small and only affect the dynamics near the weak selection limit, they describe non-monotonic behavior en route to equilibrium, which may be surprising given the parameter description: constant population size and mutation rates with strong additive selection. Additionally, this non-monotonicity is not only due to stochastic fluctuations, as it is also present in the contributions to the mean frequency represented by the deterministic tree diagrams (D1–D2).

#### Time dependence of the variance of the frequency distribution and the homozygosity

The dynamics of equilibration for homozygosity are less obvious than for the mean frequency, given that there is no deterministic description. However, the field theoretic description of the partition functional allows for approximation of any moment of interest using the same techniques. All diagrams for the homozygosity have two external vertices. One can categorize diagrams with multiple external vertices into two classes: disconnected and fully connected diagrams. A diagram is considered disconnected if its topology can be reduced into distinct sub-diagrams, as depicted by the following example (left diagram below). The second diagram shown below (right) is similar, but with an additional drift vertex (V4) connecting the two subdiagrams of the disconnected example (left); this diagram is fully connected (i.e., cannot be separated into two sub-diagrams without cutting an edge).

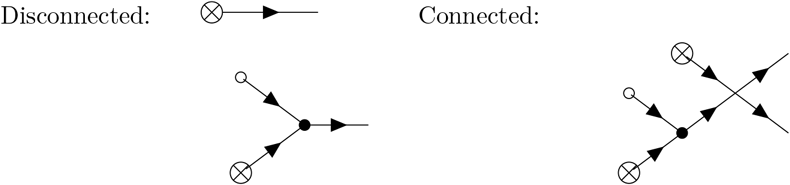

The disconnected diagram above represents a net contribution to the variance (and homozygosity), but each sub-diagram represents a contribution to the mean frequency corresponding to integrals already computed in the previous subsection in Equations 25 and 28. In fact, all disconnected diagrams represent a product of contributions to lower moments. As a corollary, if all lower moments are evaluated (or approximated), the only additional diagrams needed to compute a given non-central moment are connected diagrams; the sum of all connected diagrams with *n* external vertices represents the *n*^th^ cumulant *κ_n_* and, thus, all unique information for a given moment can be obtained from summing connected diagrams. The first three cumulants of the allele frequency are equal to the first three non-standardized central moments, the mean, variance, and the non-standardized skew (i.e., Skew × Var^3/2^) of the allele frequency probability distribution. The fourth cumulant *κ*_4_ is equal to the non-standardized excess kurtosis (i.e., Ex.Kurt × Var^2^), rather than the fourth central moment.

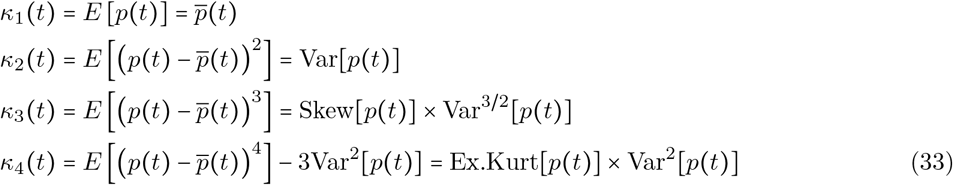

Like the fourth cumulant, all cumulants above the third (*κ_n_* for *n* > 3) differ from their respective central moments. Non-central moments *E*[*p^k^*] can be computed once all cumulants *κ_n_* with *n* ≤ *k* are calculated from the sum of connected diagrams. For example, the time dependence of the homozygosity, the second non-central moment, can be found by adding 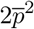 (using 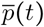) from Equation 32) to the sum of connected two-vertex diagrams (i.e., 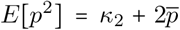, where *κ*_2_ = Var[*p*]). Like the approximations for the mean frequency, these time-dependent cumulants, as well as their respective moments, are not represented by a single specific path that an allele takes to travel through the population; they are properties of the time-dependent probability distribution of all possible paths.

The lowest order approximation to the variance (or second cumulant *κ*_2_) of the allele frequency is represented by the sum of contributions from the following two-vertex diagrams, both of which are related to genetic drift (i.e., contain the vertex V4).

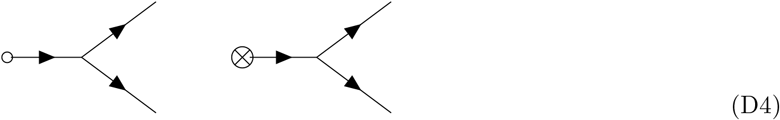

The fact that the largest contributions to the variance are dependent on drift becomes important in the context of non-equilibrium demography, as discussed below. The sum of these two-vertex diagrams yields the following approximation for the variance of the allele frequency.

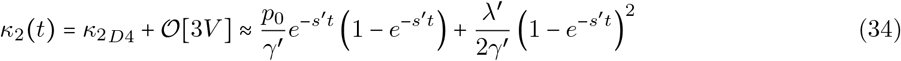

Note that all diagrams contributing to the variance have at least a counting factor of 2 from the choice of which external vertex connects to each propagator exiting the diagram (<u>Rule 3</u>), but can have additional counting factors associated with the specific diagram topology. These counting factors can again be determined by the number of Wick contractions allowed for a given vertex, conditional on the time-ordering requirement of <u>Rule 5</u> (see A2 describing the use of Wick contractions to assess counting factors for each diagram). Analogously, for all cumulants *κ_n_* above the mean (*n* > 1) there is at least an overall counting factor of *n*! from the number of ways to connect exiting propagators to the external vertices; however, the external vertices should not be counted as repeated vertices (i.e., they are not subject to the 1/*n*! factor from <u>Rule 3</u> since this is already accounted for in the stated Feynman rules). The first (and largest) corrections to this approximation are represented by several three-vertex diagrams: three tree diagrams and two loop diagrams. The three-vertex tree diagrams have the following topology, which, notably, contains an *N*_4_ drift vertex V5.

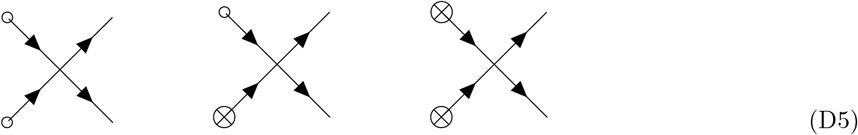

Unlike previous diagrams, contributions from the above diagrams are negative due to the *N*_4_ coupling constant (i.e., 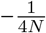 in V5). By comparing Diagram D4 to D5, one can see that the *N*_3_ vertex can always be replaced with an *N*_4_ vertex by connecting the additional edge that is introduced in the process to a terminal vertex (V1 or V2); this produces a new higher-vertex order diagram with the opposite overall sign, which is suppressed by a factor of the appropriate perturbative parameter from the added vertex (e.g., *p*_0_ or *λ*′). Heuristically, this accounts for the fact that fluctuations due to drift occur at both low and high frequencies (represented by V4 and V5, respectively), but an allele must be driven to high frequencies by some other force (here, e.g., mutation or the initial condition) before the high-frequency effects of drift are relevant. Evaluating and summing the (negative) contributions shown in Diagram D5 yields the following expression.

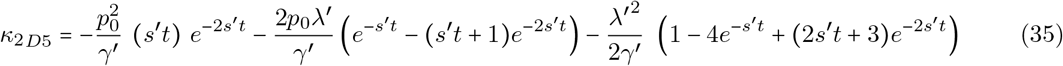

The following one-loop diagrams also represent three-vertex corrections to the variance.

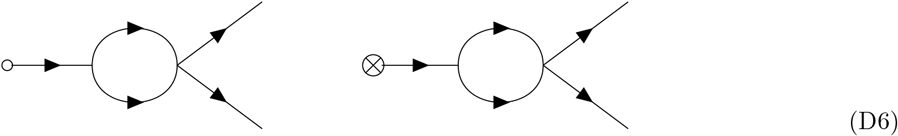

These diagrams depict the same scenario described above: some force drives alleles to high frequencies, which makes the *N*_4_ vertex relevant, but in this case that force is drift, itself. The resulting loop diagrams thus describe interactions between stochastic fluctuations in the allele frequency distribution, or, biologically speaking, self-interactions in genetic drift. Evaluating the contributions represented by these diagrams yields the impact of the lowest order drift-drift interactions on the allele frequency variance.

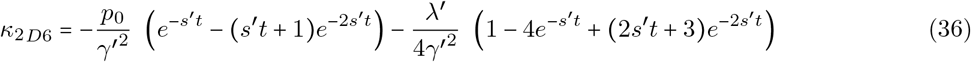

As one might expect, self-interactions are generally suppressed when selection is efficient, depending on 1/*γ*′^2^, as is required for the perturbative PGFT presented here. Comparing these terms to Equation 34 makes it clear that interactions between fluctuations (and the diagrams describing this phenomenon) become technically relevant to the variance as 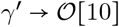, or, roughly when the lowest order approximation breaks down; assuming a range of validity of roughly *γ*′ > 5, self-interactions due to drift are relevant at the very edge of this regime and are largely to blame for the failure of the perturbative approximation. The late time asymptotic behavior *t* → ∞ also makes it clear that the three-vertex approximation, which is negative, dominates over the two-vertex approximation in Equation 34 when *γ* > 1/2, at which point the variance becomes negative (and thus ill defined); this represents a catastrophic breakdown of the three-vertex approximation (this is plotted explicitly in Supplemental Material S4.1 in Figure S6 for intuition). However, in this regime, the infinite number of higher order terms of 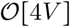 become technically relevant such that the weak selection behavior is very poorly approximated by the three-vertex approximation of the variance (i.e., the variance is still strictly positive, but the perturbative theory can no longer be truncated at a finite point). Comparison of the prefactors in Equation 36 to those in Equation D5 makes it clear that the three-vertex tree diagrams, in contrast to the loop diagrams, describe corrections due to very large mutation rates, rather than small population sizes controlled by *γ*′; the tree-diagrams compete with drift self-interactions roughly when mutations approach the recurrent regime 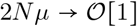 (recurrence is nicely depicted by the presence of two mutational influx events in Diagram D5*R*, leading to the *λ*′ dependence). Equation D5 also becomes technically relevant when the initial frequency is sufficiently high (namely, when *p*_0_ is of order 10%), but I have assumed throughout that this is not the case in order to focus on the equilibration of new mutations. The preceding analysis gives a sense of the power of this formalism: PGFT is capable of identifying complex self-interactions between drift-based fluctuations and uses straightforward algorithmic steps to produce closed-form analytic approximations that model their impact on the dynamics (without the need for numerical solutions); the theory itself details the mode of failure, which, in turn, defines the regime of validity for the perturbative approximations.

The full three-vertex approximation, defined below, is the sum of Diagrams D4, D5, and D6. The time-dependence described by this expression is used for comparison to simulations in the form of the three-vertex homozygosity (see Supplemental Figure S6), but I will focus subsequent discussion on the leading order (two-vertex) approximation presented in Equation 34.

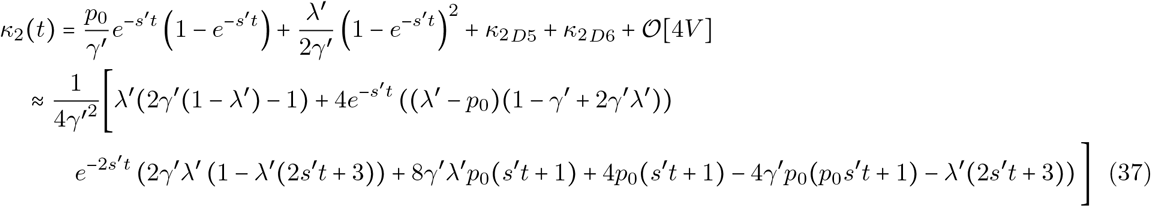

From the two-vertex part of this expression (i.e., the first two terms), it is clear that at early times *t* → 0 the variance (= *κ*_2_) approaches zero (i.e., the variance of the delta function initial condition), which is the case for the three-vertex expression, as well. At asymptotically late times, the leading contribution to the variance approaches *κ*_2_ ≈ *μ*/4*Ns*′^2^. This can be easily interpreted by writing the functional form for the homozygosity E [p^2^], which is a familiar quantity.

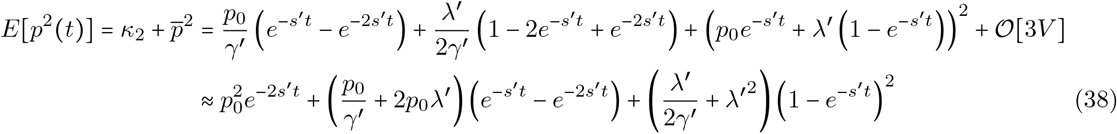

Considering the limit where *s* ≫ *μ* > *μ_b_* and in the absence of recurrent mutations 2*N_μ_* ≪ 1 (i.e., the infinite sites limit), the population parameters reduce to their normal definitions *γ*′ → *γ* and *λ*′ → *λ* and Equation 38 can be compared directly to the second non-central moment of the site frequency spectrum with target size *L* = 1. At the initial time *t* = 0, this expression approaches the homozygosity of the delta function imposed by the (non-equilibrium) initial condition.

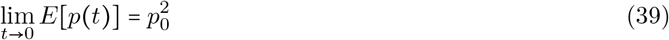

This is simply the homozygosity when the variance vanishes (i.e., 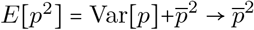). At asymptotically late times *t* → ∞, the homozygosity approaches that of a population in mutation-selection-drift balance.

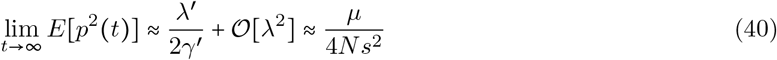

The same equilibrium value can be obtained by taking the strong selection limit of Kimura’s solution for the equilibrium frequency spectrum [33], which was given explicitly by Nei [34], and integrating over *p*^2^.

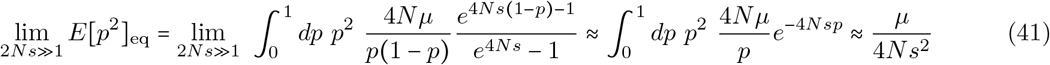

Thus, the late time behavior is simply the homozygosity in mutation-selection-balance, as expected for constant population size. The lowest order expressions line up well with simulations run under a variety of parameters, as shown in Figure 1 and Supplemental Material S4.1 (see Figures S1–S12 for all relevant moments).

#### Time dependence of the third cumulant, the third moment, and the skew

The third and fourth moments of the allele frequency probability distribution are the subject of significantly less study than the lower moments, suffering from more opaque biological interpretations. A brief calculation of each has been included here for two reasons. The first is precisely because they are understudied, despite containing unique information about the population parameters, history, and dynamics. The second is that they both share an interesting property, namely that they become independent of the strength of selection in some parameter regimes, which will be discussed in the comparison between non-equilibrium demographies. I have presented their respective computations briefly here to establish a null for later comparison.

For the third and higher moments, only the lowest order contributions will be computed for algebraic simplicity, but the associated approximations are sufficiently accurate when compared to simulations in the same parameter regime described above. The lowest order contributions to the third cumulant *κ*_3_, or the non-standardized skew, are represented by the following diagrams.

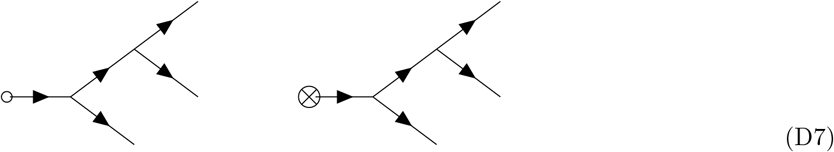

The leading dependence for the third cumulant is computed from the sum of the two three-vertex diagrams shown above.

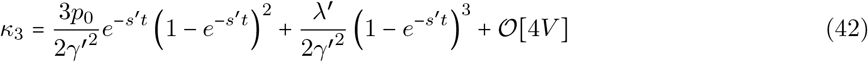

The third non-central moment *E*[*p*^3^(*t*)] can be approximated to three-vertex order using this expression and Equations 32 and 37 (or the expressions for the non-central moments) using the following relationship.

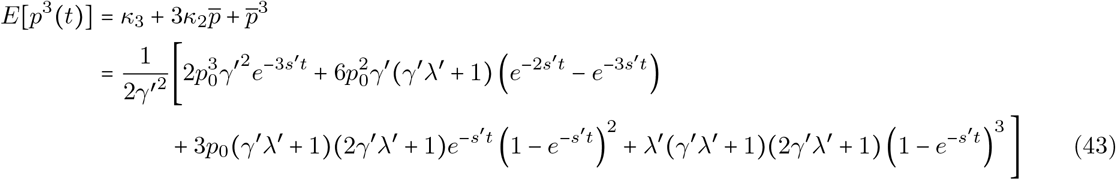

Here, the lowest-vertex approximations for the mean and variance were used for analytic simplicity (but Equations 32 and 37 can be used, if desired, along with higher-vertex corrections to Equation 42). The expression above is organized to make it clear that, at the initial time *t* → 0, all but the first term vanish and the third moment reduces to 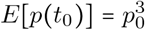 (i.e., the third non-central moment of the delta function initial condition). As with the variance, the non-standardized skew vanishes at early times. In the strong selection limit (*γ*′ → *γ* = 2*Ns* ≫ 1) of the solution to the equilibrium frequency spectrum [34] (without recurrent mutation 2*N_μ_* ≪ 1), the expression for *E*[*p*^3^(*t*)] has the following dependence at asymptotically late times *t* → ∞.

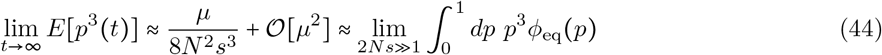

This is the same value obtained for the third non-central moment by integrating over the the equilibrium frequency distribution *ϕ*_eq_(*p*) (far right in the above equation) assuming strong selection, as was the case for the second moment in Equation 41. Surprisingly, the third *standardized* cumulant (the properly defined skew *K*_3_ ≡ *κ*_3_/*κ*_2_^3/2^) approaches a fixed value independent of the strength of selection, provided selection remains strong (i.e., *γ*′ > 5).

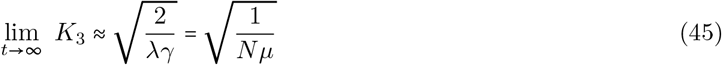

In a population with constant size and mutation rates, the equilibrium value of the skew is therefore approximately fixed over a range of selection coefficients, depending only on the population scaled mutation rate *θ* ≡ 4*N_μ_* (see Figure 1 and Supplemental Figure S9). This indicates that the relevant asymptotic dependence on selection in the non-standardized third cumulant *κ*_3_ is largely inherited from the dependence of the second cumulant, the variance *κ*_2_. Understandably, this is only the case after equilibration to mutationselection-drift balance (note the time dependence in Equation 42 and see simulation plots in Figure 1 and Supplemental Figure S9), which suggests that this quantity may be sensitive to the non-equilibrium dynamics during the equilibration process itself given the theoretical value above. Additionally, as described below, the dependence of the asymptotic value on the population size becomes relevant when the demography is time-dependent, or when the mutation rate is a function of time.

#### Time dependence of the fourth and higher cumulants and moments and the excess kurtosis

Here, the fourth cumulant *κ*_4_ and non-central moment *E*[*p*^4^] are presented. Although higher moments can be computed analogously, enumeration of the diagrams and evaluation of the resulting integrals can quickly become complicated. While the diagrammatic procedure itself is similar, calculation of higher cumulants does require identification of any missing diagrams, including those that do not have topological analogs to diagrams for the first four cumulants (e.g., there is one new type of diagram for the lowest order contributions to the fifth moment, etc.). To motivate the methodology and for future applications that may require higher moments, Supplemental Material S5 provides general formulae derived for the subset of contributions to general moments *κ_n_* that have directly analogous diagrams to those for the fourth and lower cumulants. However, generalized analytic approximations for arbitrary moments is beyond the scope of this manuscript. For the fourth cumulant, specifically, the primary topic of interest in this section, there are two classes of diagrams: those that have direct analogs to diagrams for the first three cumulants and those with without lower-moment analogs.

The fourth cumulant has four contributing diagrams at lowest order, each containing four vertices. The first two are analogous to the diagrams for *κ*_3_ in Diagram D7. These diagrams have a single terminal vertex (one with V1 and another with V2) and three copies of V4 in a *minimally connected* (or ‘maximally long’) chain of drift vertices.

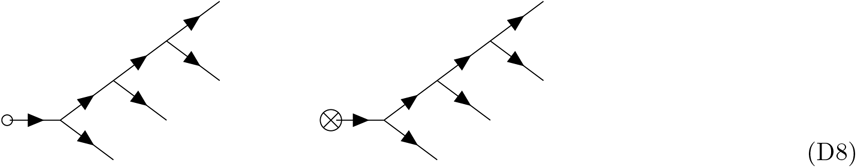

The associated contributions sum to the following expression.

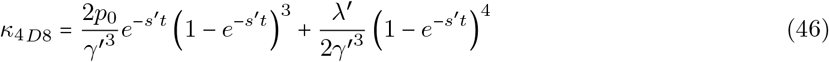

The second pair of four-vertex diagrams for *κ*_4_ have the following topologies.

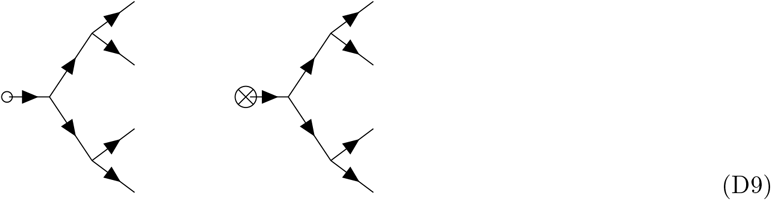

These diagrams are a series of *maximally connected N*_3_ drift vertices (i.e., these diagrams have the minimal average number of propagators between the terminal and external vertices in a connected diagram). Note that this concept is not well defined for arbitrary cumulants, but provides an adequate descriptor for distinguishing contributions to the the fourth cumulant. This is the first example thus far of multiple diagrams contributing to a single cumulant with identical vertices and distinct topologies (e.g., the two leftmost or two rightmost in Diagrams D8 and D9). After calculating the associated contributions for these diagrams, the resulting time-dependence is identical to the contributions represented by Diagram D8 up to an overall constant. These diagrams represent exactly half of the contributions of those in Diagram D8, but are of the same order and thus non-negligible when computing *κ*_4_. The additional factor of two found in Equation 46 can be attributed to a counting factor of 2^2^ = 4 associated with the choice of which 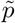 exiting each of the *N*_3_ vertices connects to a subsequent *N*_3_ vertex (as opposed to an external vertex) and an additional factor of 1/2 from integration due to the difference in the propagators. The counting factor for the Diagrams in D9 is one (i.e., once the first propagator has been used to connect two *N*_3_ vertices, the time dependences of all remaining propagators are fully determined; see Appendix A2).

Summing the above contributions, the time dependence of the fourth cumulant is well approximated (in the perturbative regime *γ*′ ≳ 5) by the following functional form.

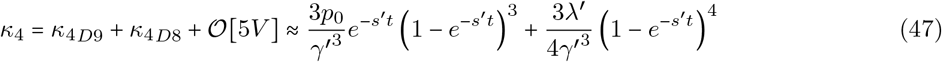

The fourth non-central moment can be computed using the results for the first four cumulants, noting that 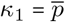 in the definition below, all of which are now computed.

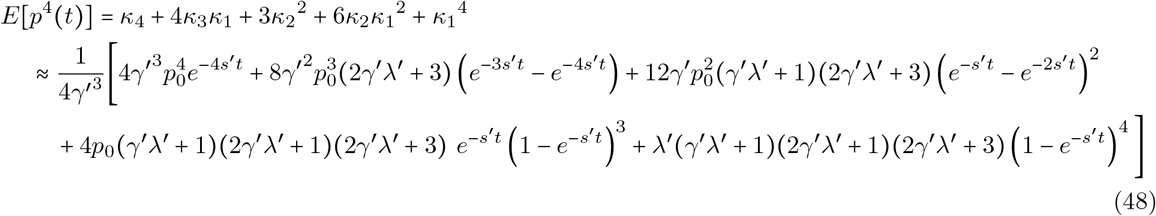

In order, other than the approximation of *κ*_4_ given immediately above, the expressions for the cumulants on the right hand side in the above expression use the lowest-vertex approximations in Equations 42, 26, and 34 (higher-vertex approximations can be used for all four cumulants, if desired). The trajectory described by Equation 48 aligns well with simulations (see Figure 1, for example) over a wide parameter range presented in Supplemental Material in Figure S11; Figure S10 shows the same agreement to simulations for the fourth cumulant expressed in Equation 47. One can see by inspection that all terms but the first vanish at early times as *t* → 0, which again results in the fourth moment of the delta function initial condition. As expected from the prior moments, at asymptotically late times and taking the weak mutation limits *γ*′ → *γ* and *λ*′ → *λ*, the fourth non-central moment equilibrates to the strong selection approximation of the equilibrium solution for the allele frequency distribution *ϕ*_eq_(*p*) derived in [34].

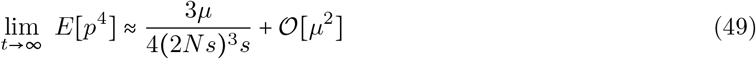

However, unlike the lower moments, this is the result of a sum over two diagrams containing the mutational influx vertex V2, which sum to the correct asymptotic value.

Notably, as with the standardized skew *K*_3_, the standardized excess kurtosis *K*_4_ ≡ *κ*_4_/*κ*_2_^2^ approaches a selection-independent value at asymptotically late times when the population size is constant.

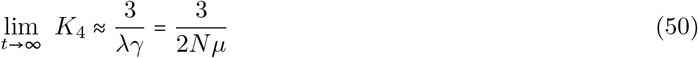

This again indicates that, at equilibrium (i.e., in mutation-selection-drift balance), the excess kurtosis of the allele frequency probability distribution is independent of the strength of selection, provided it remains efficient (*γ*′ > 5), which is clear from the plot in Figure 1. The dynamics of the excess kurtosis is presented over a wide parameter range in Supplemental Figure S12. This effect appears across the regime of validity for the approximations (i.e., for roughly *γ*′ ≳ 5). As with the standardized skew, the excess kurtosis is dependent only on the population scaled mutation rate *θ*, but the dependence on the population size is relevant when the demography is nontrivial or the mutation rate is time-dependent. Finally, note that this dependence is simple enough to identify a relationship between the asymptotic values for the skew and excess kurtosis, as discussed below.

### On the equilibrated allele frequency probability distribution

Under strong purifying selection, the equilibration process for a constant sized population approaches an equilibrium allele frequency probability distribution described by Nei (1968) [34]. This takes the form of a Gamma distribution Γ(*α, β*) [34], where *α* = *θ* ≡ 4*N_μ_* and *β* = *γ* ≡ 2*Ns*, and is the strong selection limit of the mutation-selection-drift balance distribution derived by Kimura [33]. While the equilibrium distribution itself is well known, one feature of this distribution is, to my knowledge, less widely appreciated. After standardizing the third and fourth cumulants (*κ*_3_ and *κ*_4_) by the appropriate powers of the variance (*κ*_2_), the skew *K*_3_ and excess kurtosis *K*_4_ are entirely independent of the selection coefficient *s* (discussed above in the context of asymptotic equilibration dynamics). The skew and excess kurtosis of the Gamma distribution are 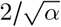 and 6/*α*, respectively, which are independent of the *β* parameter and, thus, independent of the strength of selection. While selection must be sufficiently strong that Nei’s limit of the distribution is valid (i.e., 2*Ns* ≫ 1), for all selection coefficients fitting this criteria, the skew (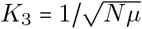; see Equation 45) and excess kurtosis (*K*_4_ = 3/2*N_μ_*; see Equation 50) are indeed only a function of the population scaled mutation rate *θ*. As a direct corollary, the ratio of the standardized excess kurtosis to the square of the standardized skew, which I will refer to as *ξ* (up to a constant factor 2/3), must have a constant value for the probability distribution of alleles under strong selection in equilibrium.

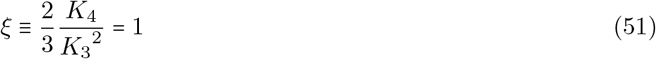

This constant *ξ*, which is a unique feature of the Gamma distribution, provides a measurable quantity that represents equilibrium for genetic variation under strong purifying selection. As such, if a set of alleles is known to be under strong selection (e.g., a set of highly conserved loss-of-function mutations), assessment of the average value of the per-allele *ξ* provides a quantitative test for equilibrium at any given time point: if *ξ* significantly deviates from 1, it is unlikely that the population is in equilibrium, while, if *ξ* is statistically indistinguishable from 1, the population is consistent with mutation-selection-drift balance equilibrium in a constant sized population. Stated another way, *ξ*(*t*|*γ* ≫ 1) = 1 appears to be a ‘non-equilibrium index’ for population genetics across a subset of the genome. The insensitivity to the selection coefficient *s* and, assuming *μ* is constant, mutation rate, implies that data can be pooled across a wide range of selection coefficients and mutational classes to generate power needed to estimate *ξ* from data (i.e., the precise selection coefficient need not be known for a pooled subset of highly conserved sites). In practice, the values of {*κ*_2_, *κ*_3_, *κ*_4_} or {*K*_3_, *K*_4_} must be estimated from a population sample and the associated effects of binomial downsampling must be corrected before comparing to *ξ* = 1. Additionally, there is no guarantee that this particular statistic, which is a ratio of potentially noisy observables, has sufficient power to detect any given non-equilibrium scenario. However, identification of a ‘non-equilibrium index’ of this kind (*ξ* or some analogous alternative) could prove useful for data quality assessment, classification, and to determine which population genetic statistics are applicable to a given sample.

### Equilibration times for moments of the allele frequency probability distribution

With a description of the equilibration process for the first several moments of the frequency distribution, one can assess the equilibration time for the allele frequency probability distribution, starting from a known initial frequency *p*_0_. In other words, it is of interest to characterize the time at which the effects of the initial condition can largely be ignored. As in the previous sections, I will assume *p*_0_ is on the order of 1/2*N* for a new mutation in a constant sized population and thus relatively small, but the perturbative limit extends to initial frequencies below roughly 10%. The equilibration time can be analyzed on a per-moment basis to assess whether the timescale differs between distinct features of the distribution (e.g., to determine whether the variance and homozygosity equilibrate on the same timescale as the mean frequency). Beginning with the mean, the previous theoretical and simulation-based analysis indicates that equilibration is well-described by the deterministic approximation in Equation 26 when selection is efficient (i.e., in the regime of validity *γ*′ ≳ 5). The equilibration time for the mean 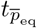 can be assessed by asking when the time dependence in Equation 26 is negligible relative to the asymptotic state of mutation-selection balance for the mean frequency, 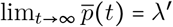. This can be phrased as the condition 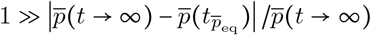, which requires that the magnitude of the difference between the mean at the time of equilibration and at the final time (i.e., the percent difference) is relatively small, where the absolute value asserts that the direction of deviation from the asymptotic value is unimportant. After inserting the functional form for 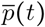 and the final value of *λ*′ = *μ/s*′, this condition becomes the following inequality.

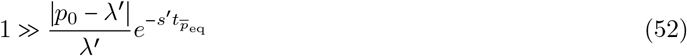

The equilibration time 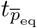 corresponds to when the right side of this inequality is roughly an order of magnitude less than the left side (i.e., replacing the ‘≫‘ symbol with a factor of ten). Re-expressing this condition in terms of 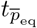 results in the following parameter dependence for the the equilibration timescale.

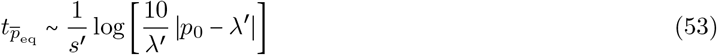

This expression is well defined, provided *p*_0_ ≠ *λ*′, and the dependence is smooth away from this pole; the logarithm will usually be dominated by either *p*_0_ or *λ*′, depending on the parameter regime. Using an example set of parameters, for a new mutation with initial frequency *p*_0_ = 1/2*N* in a population with constant size *N* = 10^4^ (i.e., *p*_0_ ~ 10^-4.5^), the logarithm is well approximated by the *p*_0_ dependence when *μ/s*′ ≪ 10^-5^. For a mutation rate of *μ* = 10^-8^ under the assumption of perturbative mutational effects *s* ≫ *μ* > *μ_b_* such that *s*′ → *s*, this corresponds to selection coefficients stronger than roughly *s* ~ 10^-3^ (i.e., the regime *γ* ≳ 20); in this strong selection regime, equilibration of the mean frequency occurs on a timescale of roughly 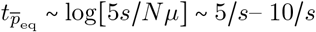. Figure 1 and Supplemental Figures S2 and S3, which plot the equilibration of the mean frequency over a range of parameters that includes this combination, confirm this to be the correct timescale. For smaller population sizes (i.e., larger *p*_0_ for new mutations) or weaker mutation rates, the equilibration time for strongly selected alleles generically exceeds 10/*s* generations (e.g., for strong selection with *N* = 10^3^ and the same mutation rate); this is also true if the allele frequency is initially higher than 1/2*N* (i.e., when the dynamics of interest are not related to new mutations). Intuitively, the equilibration of alleles under selection is usually considered to happen rapidly on the order of 1/*s* generations, trivially ascertained from the decay of the exponential, which is sometimes used as a rule of thumb to determine a sufficient burn-in time for simulations (i.e., the initial period used to erase transient effects of an arbitrary initial condition in population genetic simulators) when selection is much stronger than drift by elapsing 10/*s* generations. From the above equation, it is clear that the equilibration time depends logarithmically on both the initial conditions and the mutation-selection balance parameter *λ*′ (or *λ* = *μ/s* for small, oneway mutation rates), and can span an additional order of magnitude. This equilibration time for the mean frequency also has non-trivial variance around the expected time presented above, where the variance of the expected trajectory for the mean frequency is proportional to the variance of the probability distribution in Equation 34; accounting for the variability of the equilibration time inherently extends the minimum timescale needed to ensure the equilibration of all sites. Thus, to completely remove transient effects from the initial condition in a simulation with these parameters, the burn-in period used should be at least of order 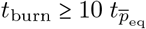 (accounting for estimation of the simulated mean frequency alone), which is substantially longer than 1/*s* and, in some situations, begins to compete with the coalescent time for neutral variation [74]. To simulate features of the distribution beyond the mean, this minimal time must be extended by an amount dependent on the feature of the distribution of interest, requiring an assessment of the equilibration of other moments (see below).

For higher cumulants (and moments), which have inherently more complicated time dependence due to drift, the equilibration time *t*_eq_ (temporarily dropping the moment subscript) can be assessed by analyzing the near-asymptotic dynamics in the large, but finite limit of *t*. The time-dependence for each quantity is expressed as a sum of powers of exponentials, which makes separation of this timescale straightforward: the square of an exponential (*e^−s′t^*)^2^ is simply an exponential with twice the decay rate, and is quickly suppressed relative to *e^−s′t^*. Thus, all exponential dependence other than on the slowest decay rates are negligible at late times, provided *e*^*-s′t*_eq_^ ≪ 1. Noting that the various exponentials appear in, e.g., Equation 34 after expanding the square, all time dependence proportional to *e*^-2*s′t*^ can be dropped for this reason when the evolution of the variance approaches the equilibration time *t* → *t*_*κ*_2_eq___ < ∞. The remaining time dependence is a simple exponential of the same form used to obtain Equation 52 for 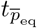, which can be used to find the equilibration time. The parametric dependence of the resulting equilibration time time can subsequently be used to confirm that this finite time limit was applicable by verifying that 1 ≫ *e*^*-s′t*_eq_^. Applying this to the variance, the equilibration time is readily obtained from Equation 34 by repeating the procedure above: subtract the asymptotic parameter dependence *μ*/2*γ*′^2^, normalize by the same quantity, multiply the exponentially dependent terms by a factor of roughly ten, set this to one, and solve for time.

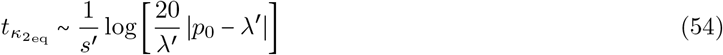

From this expression, the rate of exponential decay can be assessed to confirm that 1 ≫ *e*^-*s′t*_eq_^. This condition becomes 1 ≫ *λ*′/2|(*p*_0_ – *λ*′)| and is satisfied so long as *p*_0_ > *λ*′, which is the regime of primary interest for the present analysis (i.e., weak mutation and strong selection). In the strong mutation limit, the condition for the variance, specifically, can be determined from the full inequality by taking the positive real root of the quadratic expression in powers of *e*^-*s′tκ*_2_eq__^. Analogously, the equilibration time for the third and fourth cumulants, the non-normalized skew and excess kurtosis of the frequency distribution, can be computed in the same way, again dropping all powers of *e*^-*s′t*_eq_^ above one.

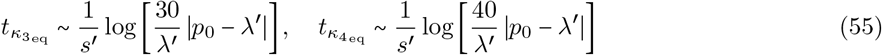

These equilibration times are again only valid for *p*_0_ > *λ*′. The factor of *n* inside the logarithm (e.g., 2 for the variance *κ*_2_, 3 for *κ*_3_, etc.) appears from the linear expansion of the binomial in (1 – *e^−s′t^*)*^n^* for the mutation term in Equations 34, 42, and 47. If this pattern is consistent at higher cumulants, which will not be verified here beyond *κ*_4_, the generalized equilibration time for the lowest order approximation to the *n*^th^ cumulant *κ_n_* is, by extension, 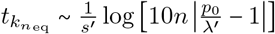, Note that the subset of lowest-order diagrams computed for general cumulants *κ_n_* in Supplemental Material S5 contains a dependence on (1 – *e^−s′t^*)*^n^* (i.e., the term responsible for the factor of *n*), which likely justifies this generalization; however, this is insufficient to prove the generalized behavior of the equilibration times *t*_*k*_*n*_eq___ without accounting for all diagrams for general moments. These equilibration times differ from 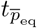 by roughly 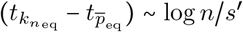 generations (at least for the first four moments), which is insignificant when *p*_0_/*λ*′ ≫ *n*, and is particularly impactful when *p*_0_/*λ*′ ≲ *n*. For the regime 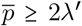, the equilibration time of the *n*^th^ cumulant is also bounded by *t*_*k*_*n*_eq___ ≥ log[10*n*]/*s*′. Thus, the equilibration time of the allele frequency probability distribution as a whole is governed by the largest observable moment, where ‘observable’ is determined by the inverse of the haploid population size 2*N* (or inverse sample size when working with a sample) used to estimate the distribution at time *t*, and is obviously bounded by the neutral equilibration time (*t*_eq_ = 8*N* from the coalescence time of 2*N* individuals [74]). Perhaps surprisingly, the naive equilibration time for alleles under strong additive selection is thus only realistic for the equilibration of the lowest moments, and only in a limited parameter range. All cumulants of the frequency distribution appear to equilibrate on a longer timescale of at least roughly *t*_eq_ ~ 10/*s*′, but potentially much longer for higher cumulants and moments *n* ≥ 10, provided *t*_*k*_*n*_eq___ ≤ 8*N*.

The above approximations are specific to cumulants, though the equilibration times for non-central moments are definitionally on the same order. However, the standardized skew and kurtosis of the allele frequency probability distribution have different equilibration times due to the normalization by powers of the variance. To approximate the equilibration time for these moments, one can Taylor expand the analytic expressions for the standardized skew (*K*_3_ ≡ *κ*_3_/*κ*_2_^3/2^) and for the standardized excess kurtosis (*K*_4_ ≡ *κ*_4_/*κ*_2_^2^) to first order in the small parameter *e^−s′t^* before performing an analogous analysis to that shown above. For the standardized skew of the frequency distribution, the equilibration time is approximately the following (again using a factor of 10 for the ≫ inequality).

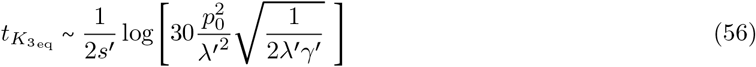

Note that the denominator has a factor of two relative to the equilibration times above. In the Taylor expansion, the first order dependence on *e^−s′t^* from *κ*_3_ is cancelled by that from *κ*_2_^3/2^ such that the quadratic dependence *e^−s′t^* is the leading non-asymptotic term. The equilibration time for the standardized excess kurtosis can be expressed in a similar form.

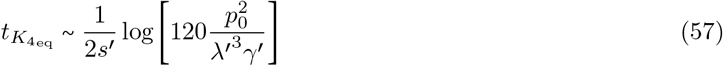

The dependence inside the log in the above expression for *t*_*K*_4_eq___ differs from *t*_*K*_3_eq___ by a factor of 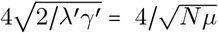, indicating that the excess kurtosis is slower to equilibrate by log[32/*λ′γ′*]/4*s*′ = log[16/*N_μ_*]/4*s*′ generations (which is of order 5/2*s*′ generations for *N* = 10^4^ and *μ* = 10^-8^, consistent with only a slight difference seen in the respective plots in Figure 1).

### Dynamics of the first four moments in non-equilibrium demographies

After describing the temporal process of equilibration of the allele frequency probability distribution for a population of constant size, I now extend this analysis to examples where the population size changes over time. Analytic approximations for the the temporal evolution of the first four moments of the allele frequency distribution are provided below for three cases of non-equilibrium demographic history: the exponential growth of the population, a population with a temporary ‘square bottleneck’, and a population with cyclical oscillations in size. The same diagrams shown above for populations of constant size provide a representation of the integrals contributing to various moments in the case of a population with time-dependent population size *N*(*t*); no additional diagrams need be considered. The difference is that the coupling constants for the two drift vertices V4 and V5 cannot be pulled out of the integration as constants, and instead are a time-dependent function of the vertex time. Diagrams that do not contain the drift vertex, namely only tree diagrams for the mean frequency shown in Diagrams D1–D2, are unchanged by the change in demography (but would be treated in a similar way if the mutation rate or selection coefficient were time-dependent). The remaining diagrams, all of which that contain drift vertices and are thus sensitive to demography, are appropriately altered, as shown below for the integral form of Diagram D4.

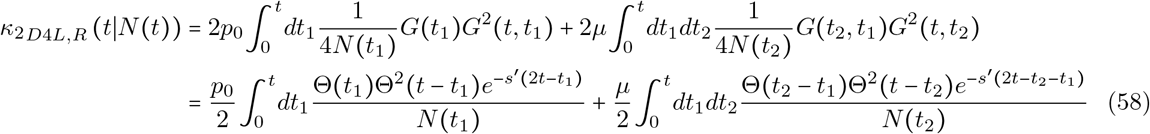

Integration in the above expressions cannot be performed until a functional form is specified for the demography *N*(*t*). Note that in the first integral (for Diagram D4*L*), the population size is a function of the vertex time *t*_1_ in contrast to the second integral (for Diagram D4*R*), which is a function of *N*(*t*_2_). Similarly, if multiple drift vertices appear in a diagram, they will each carry a different integration variable *t*_1_, as is the case, for example, for the integrals represented by Diagram D7. The right diagram in Diagram D7 represents the following integral contribution to the third cumulant *κ*_3_.

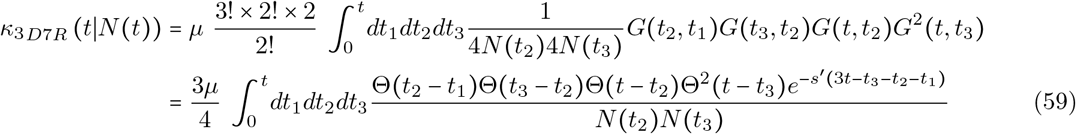

It is important to note that this integral form applies to *any* demography that can be written as a deterministic function of time. If the functional form is complicated or intractible, numerical integration can be performed to evaluate each integral for a set of parameters of interest. Additionally, as implied above, any of the coupling constants, selection, mutation, or the population size, can be treated as time-dependent in the same way, provided a deterministic functional form is used (it is possible to assume a stochastic contribution, as well, but that lies beyond the scope of this manuscript). The only caveat associated with the generalization to time-dependent selection is that the free propagator must be re-computed to solve the appropriate Green’s function equation and may take an integral form even after linearization; this can be avoided for time-dependent mutation rates simply by disallowing recurrence.

For the remainder of this manuscript I will focus on the leading order approximation to the mean, variance and homozygosity, skew and third non-central moment, and excess kurtosis and fourth non-central moment; higher vertex-order contributions shown above for the first two moments (which are computed explicitly for each demography in Supplemental Material S3 and Supplemental File S1) will be incorporated into plots comparing the appropriate approximations to simulation results, but will not be discussed further. Unlike the mutation-selection balance parameter *λ*′ = *μ/s*′, the combined variable *γ*′ = 2*Ns*′ is poorly defined for time-dependent demographies. I will instead express most results below as explicit functions of *s*′ and relevant parameterizations of the population size (e.g., *N*_0_ for the initial population size, *N_b_* for the population size of a bottleneck, *N*_max_ and *N*_min_ for an oscillatory demography). Combined variables analogous to *γ*′ are introduced in the context of each demography, all of which are defined in Supplemental Material S1 for reference.

#### Exponentially growing population size

Exponentially growing populations can be parameterized by an initial population size *N*_0_ at the initial time *t*_0_ and a finite exponential growth rate Γ per generation such that the time-dependent population size takes the following form.

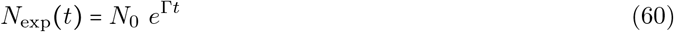

The coupling constant for the drift vertices V4 and V5 is then proportional to 1/*N*_exp_(*t_i_*) = *e*^-Γ*t_i_*/*N*_0_^ such that the population size enters each integrand in a similar form to the exponential introduced by the propagator *G*(*t_i_, t_j_*) ∞ *e*^*-s*′(*t_i_ – t_j_*)^. This makes exponential growth (or decay) particularly easy to treat analytically in PGFT and the resulting expressions remain relatively simple.

The leading order approximation for the mean frequency is identical to the deterministic expression in Equation 26, as there is no drift-dependence (i.e., no population size-dependence) at lowest order. The dependence of the mean frequency on the population size first occurs at three-vertex order, as the subdominant loop corrections scale as 1/2*N*_exp_(*t*)*s* due to their dependence on the drift vertex V4 and thus a coupling constant dependent on the population size. In contrast, the lowest order approximations for all higher cumulants are explicitly dependent on the population size, which no longer commutes with integration. When selection is efficient, the time dependence of the variance can be approximated by the following expression for the lowest-vertex contribution.

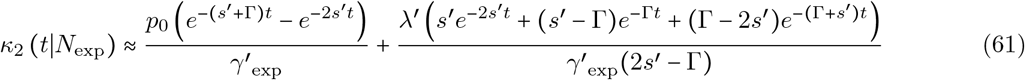

I have defined a population scaled effective selection coefficient *γ*′_exp_ as follows, which is an emergent parameter in the above expression and in all subsequent moments.

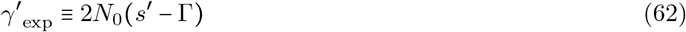

This parameter is discussed again below the calculation of the fourth cumulant *κ*_4_(*t*|*N*_exp_). Reversing the sign of Γ in Equation 61 gives the corresponding solution for exponential decay, but this approximation will break down when 2*N*_exp_(*t*)*s*′ < 5; thus, use of this approximation for exponential decay is restricted to selection coefficients that remain strong throughout the monotonic decrease in population size, with 2*N*_exp_(*t*)*s* ≳ 5 for all values of *t*. In the case of exponential growth (Γ > 0), this condition becomes *γ*′_0_ = 2*N*_0_*s*′ ≳ 5, since this will be satisfied at all later times due to the monotonic increase of *N*_exp_(*t*). The asymptotic limit of variance at late times vanishes as the population grows large enough to be well approximated by a deterministic solution, albeit with at an unrealistically large (i.e., infinite) population size. One additional concern is the appearance of poles at *s*′ = Γ and 2*s*′ = Γ, indicating that the above approximation fails for selection coefficients in the neighborhoods of both the growth rate and double the growth rate, but is well behaved both above and below each pole, provided selection remains strong relative to drift. The homozygosity can be approximated at the same order by using the deterministic expression for the mean and the variance in Equation 61 to find the following dependence.

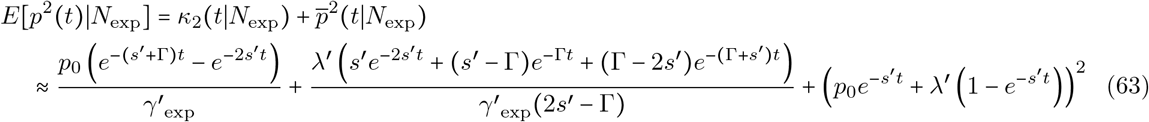

When the growth rate is large, both the variance and the homozygosity differ substantially from those in populations of constant size, given in Equations 37 and 38. Note that the homozygosity under exponential growth approaches the asymptotic value of lim_t→∞_ *E*[*p*^2^] = (*E*[*p*(*t* → ∞)])^2^ = *μ*^2^/*s*′^2^ at late times. This describes the approach to a delta function distribution at mutation-selection balance, which is expected for an infinitely large population size. All higher cumulants also vanish at asymptotically late times such that each non-central moment *E*[*p^n^*(*t* → ∞)] approaches *μ^n^*/*s′^n^*. As a result, the allele frequency probability distribution starts at a delta function *δ*(*p – p*_0_), diffuses to as the population grows, narrows as the population grows larger, and eventually approaches the final state, a delta function 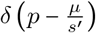 (see Figure 2 and Supplemental Figures S6 and S7). Interestingly, this introduces the possibility of a maximal variance at some finite intermediate time *t*_*κ*_2_max___, which is dependent on both *s*′ and Γ. Although it cannot be solved for exactly due to an associated transcendental equation, provided *s*′ ≠ Γ and the mutation rate is sufficiently small (*μ* ≪ *p*_0_), the value of *t*_*κ*_2_max___ can be found by taking the time derivative of 61 and taking *μ* → 0; this approximation is quite accurate because the variance and its derivative are dominated by the *p*_0_-dependent terms at early times when the maximum occurs, which can be confirmed via plot or a numerical root finder.

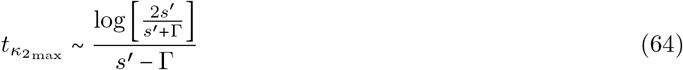

**Figure 2:**
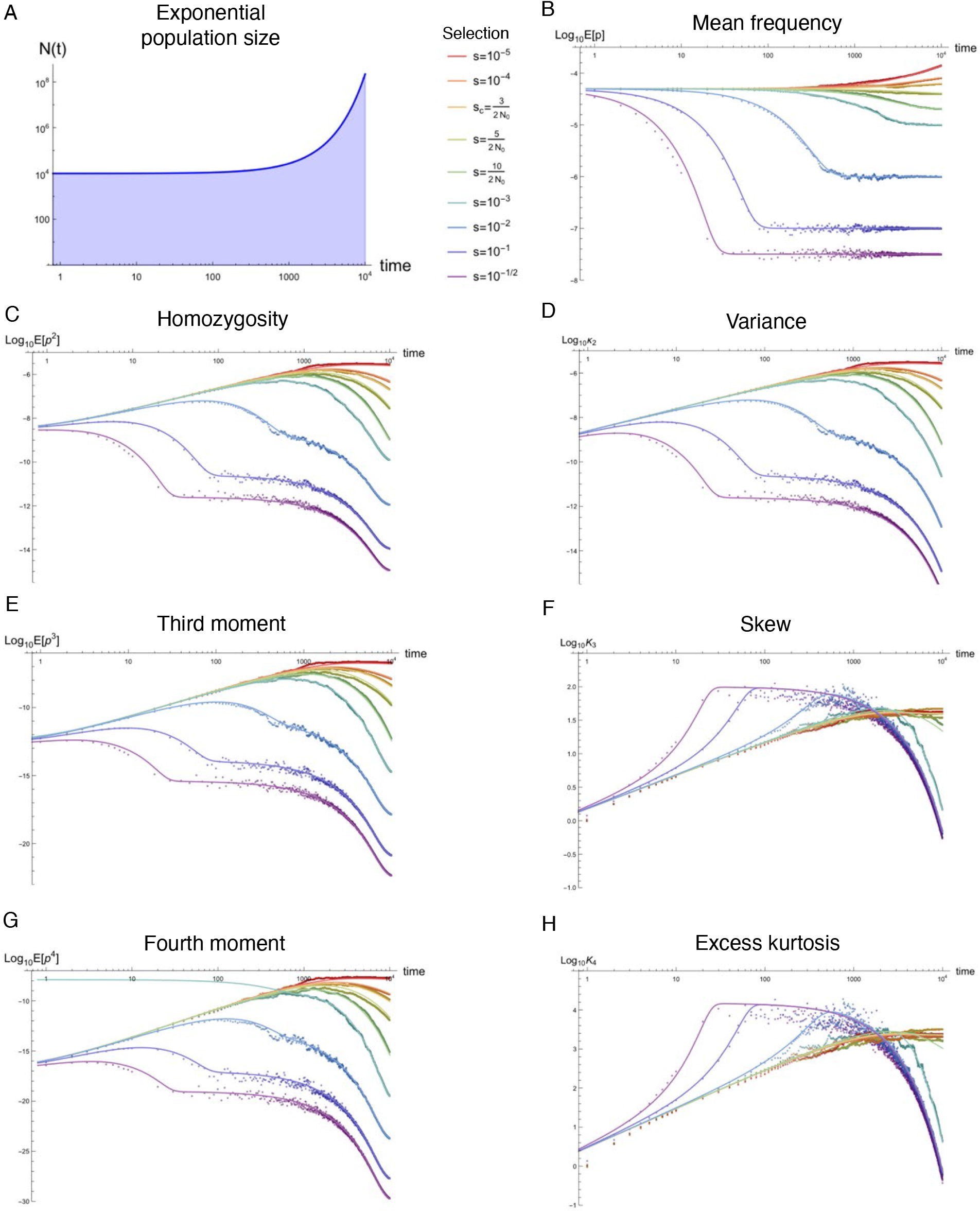
Equilibration of the first four moments, variance, skew, and excess kurtosis in an exponentially growing population. From top to bottom, the left column of plots show the time dependence of the population size, homozygosity *E*[*p*^2^(*t*)], third moment *E*[*p*^3^(*t*)], and fourth moment *E*[*p*^4^(*t*)] of the allele frequency probability distribution in an exponentially expanding population. The right column of plots shows the mean 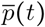, variance 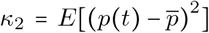, skew *K*_3_ = *κ*_3_(*t*)/*κ*_2_^3/2^, and excess kurtosis *K*_4_ = *κ*_3_/*κ*_2_^2^ for the same parameter values. One example set of parameters is plotted with an initial population size *N*_0_ = 10^4^, growth rate Γ = 10^-3^, mutation rate *μ* = 10^-8^, and back mutation rate *μ_b_* = 10^-10^; additional parameter combinations are presented in Supplemental Material S4.2 in Figures S13–S23. Solid lines in each plot represent analytic approximations from the text with selection strengths ranging from *s* = 10^-1/2^ (purple) to *s* = 10^-5^ (red), including those near where the approximation first breaks down at roughly *s*′ ~ 5/2*N*. Scattered points represent the results of estimated moments from temporal simulations using `neqPopDynx` with the same parameter values; darker shades of the each color correspond to the analytic curve with the same selection coefficient. *L* = 10^5^ sites were simulated to estimate each moment.

Note that, when *s*′ < Γ, both the logarithm and the denominator switch signs such that the above expression is strictly positive, provided *s*′ ≠ Γ. Importantly, the derivative and the above approximation are both well defined in the limit Γ → 0 describing the asymptotic approach to a demography of constant size. This indicates that the equilibration process in a constant-sized population experiences an increase in the variance of the frequency distribution, with a maximum at *t*_*κ*_2_max___ ~ log[2]/*s*′, prior to reaching mutation-selection-drift balance, which can be confirmed in the plots shown in Figure 1 and in Supplemental Figures S4 and S5 for a range of parameters (the maximum occurs in the first few time points for the strongest selection coefficient shown, *s* = 10^-1/2^); the same maximum can be seen in plots of the homozygosity, as the square of the first moment is negligible. In both exponentially growing and constant-sized populations, the variance is maximized at a much earlier time than the equilibration time (i.e., *t*_eq_ ≫ *t*_*κ*_2_max___) and therefore describes a transient effect that subsequently reduces before equilibrium is reached. However, one of the consequences of rapid exponential growth Γ ≫ *s*′ is to push this maximum to later times *t*_*κ*_2_max___ ~ log[2*s*′/Γ]/Γ than in a population of constant size *t*_*κ*_2_max___ ~ log[2]/*s*′.

The third cumulant, the skew, is given by the following expression, again only using the two lowest order contributions shown in Diagram D7.

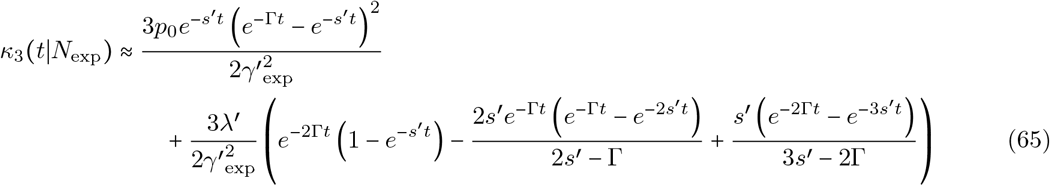

The corresponding non-central moment can be computed by substituting this into the expression below, with the second cumulant given in Equation 61 and the mean frequency, unchanged by the demography, given in Equation 26.

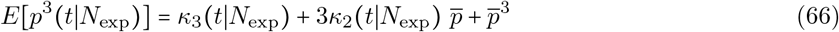

To avoid excessive algebra, I will leave this expression in this general form, but the third moment will be compared to simulations shortly. As mentioned above, this equilibrates to 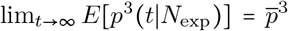 at late times as the population approaches infinite size. In an exponentially expanding population, the excess kurtosis of the allele frequency has the following form from the four lowest-vertex order diagrams.

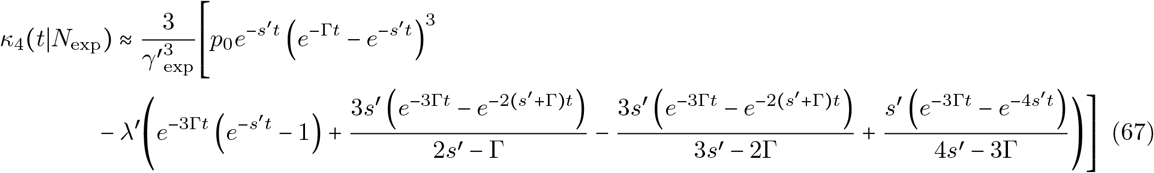

The fourth non-central moment is given by the following expression in terms of the above equation and the three lower cumulants.

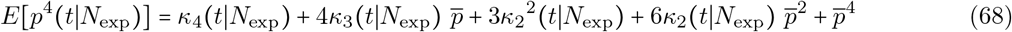

In each of the three non-trivial moments, the parameter *γ*′_exp_ defined in Equation 62 emerges, replacing the dependence on *γ*′ in the expressions for constant population size. This naively suggests that the growth rate Γ may act as a kind of additional selection pressure, but this analogy is less than straightforward due to the varied dependences on *s*′ – Γ, 2*s*′ – Γ, 3*s*′ – 2Γ, 4*s*′ – 3Γ (all four of which can be re-expressed as a sum of the form *ns*′ + *m*(*s*′ – Γ)), and, importantly, *s*′ alone. Additionally, the timescales expressed in the exponential dependence vary in a similar way (i.e., have the form *ns*′ – *m*Γ for various *n* and *m*), resulting in the non-monotonic behavior of non-central moments above the mean (see Figure 2); these curves dramatically differ from the strong selection curves appearing in Figure 1 for a population of constant size (though both complaints are unimportant when *s*′ ≫ Γ or Γ ≫ *s*′). Admittedly, these are only the lowest order expressions for each moment, making it possible that a full non-perturbative analysis contains some kind of effective *γ*′ parameter during exponential expansion (e.g., if the parameters found in the perturbative approximation are some expansion of (*s*′ – Γ)*^n^*); however, this cannot be seen directly in the perturbative regime, which does not *a priori* justify this interpretation for the growth rate.

Figure 2 shows a comparison between the approximations for the first four cumulants and moments and simulations for exponentially expanding population size; additional plots exploring a range of parameter combinations for (*N*, Γ, *s, μ*) are provided in Supplemental Material S4.2 in Figures S13–S23. From these plots, it is clear that the initial population size controls the breakdown of the strong selection approximation and that the late time behavior is that of deterministic mutation-selection balance, given sufficient time for the population to grow. The mean reaches a constant value of *λ*′, and the other moments tend towards zero in this limit. The non-monotonicity in the latter moments shows a competition between the equilibration process and the constantly shifting equilibrium point due to the rapid growth of the population (i.e., all moments above the mean have an equilibrium value dependent on the population size). While all cumulants asymptotically vanish, it is worth noting that the standardized skew *K*_3_ and excess kurtosis *K*_4_ both vanish under strong selection, as well, indicating that at asymptotically late times the corresponding cumulants, *κ*_3_ and *κ*_4_, vanish faster than the appropriate power of the variance. Additionally, for sufficiently strong selection, the variance, standardized skew, and standardized excess kurtosis approach an unstable equilibrium at relatively early times that takes on roughly the same values for these moments as for a population with constant size *N*_0_ (see Figure 2 and Supplemental Figures S6, S20, and S23). The variance reaches a transient value of *λ*′/4*N*_0_*s*′, the skew reaches 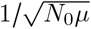, the excess kurtosis reaches 3/2*N*_0*μ*_, and temporarily approaches the equilibrium value of one (see Equation 51). This occurs due to the sufficiently rapid action of selection, which attempts to equilibrate to the initial population size when *s*′ ≫ Γ. This unstable state is eventually overtaken by the growth of the population at roughly the e-folding time of the population size *t* ~ 1/Γ (the change in population size is small compared to *N*_0_ before this time; 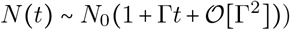, after which all three moments rapidly vanish as the allele frequency probability distribution approaches a delta function *δ*(*p – μ/s*). This behavior is of particular interest for alleles under strong selection that are often mutations associated with disease, which are usually at low frequency, but may transiently elevate in frequency in the tails of the distribution prior to the e-folding time 1/Γ. The duration of this effect and the range of selection coefficients affected by these dynamics are primarily dependent on rate of population growth.

Interestingly, an apparently stable equilibrium is reached for both the skew and the excess kurtosis for sufficiently weak selection. Under weak selection, the late-time decay of the third cumulant matches the latetime decay of the variance (to the 3/2 power), such that the skew equilibrates in the weak selection limit. Similarly, the excess kurtosis decays at the same rate as the square of the variance. This occurs for a range of selection coefficients *s*′ ≲ Γ, for which *γ*′_exp_ ≡ 2*N*_0_(*s*′ – Γ) ≈ −2*N*_0_Γ (the negative sign, as defined here, here acts like a beneficial selection coefficient, since *s* is defined as positive for deleterious selection), and is described by the analytic approximations above (unless the population size is very small) due to the exponentially decaying drift constant *e*^-Γ*t*^/2*N*_0_. In addition, a smaller range of selection coefficients in a ‘delayed maximum’ regime interpolate between the pseudo-equilibrated and asymptotically equilibrated regimes, reaching maximum values for the skew and excess kurtosis at later times when the pseudo-equilibration of selection coefficients with *s*′ ≫ Γ has already begun to exponentially decay (i.e., at times later than the e-folding time *t* ~ 1/Γ). For weak selection coefficients, the asymptotically stable skew and excess kurtosis remain at fixed values when the population size is large enough that additional sustained growth seems unrealistic in nature (e.g., for Γ = 10^-2^, these curves are flat at *N*(*t*) ~ 10^12^), after which the growth is unlikely to proceed indefinitely. This may result in an extended equilibrium as the growth rate saturates for variation under weak selection. The pseudo-equilibrated but asymptotically-time vanishing, delayed maximum but vanishing, and asymptotically equilibrated regimes of selection appear across a wide range of parameter combinations simulated for this analysis, which are shown in Supplemental Material S4.2 in Figures S13–S23.

#### A population bottleneck and subsequent re-expansion

The simplified bottleneck demography discussed here starts from a constant diploid initial size of *N*_0_ at the initial time *t*_0_ = 0, instantaneously reduces the population size to *N_b_* diploid individuals after *T_i_* generations, and subsequently re-expands to the initial population size after a duration of *T_b_* generations. In this demography, the transient response of the allele frequency probability distribution to both the reduced and increased population size are of primary interest, as well as how this response depends on the bottleneck duration, intensity, and time of onset (i.e., at what stage of equilibration from the initial condition the bottleneck occurs). The following expression for the time-dependent diploid population size *N*(*t*) can be used to parameterize a square bottleneck with the variables *N*_0_, *N_b_*, *T_i_*, and *T_b_*.

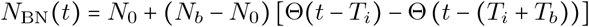

The Heaviside functions Θ(*t* – *t*′) in the above definition ensure that the population changes instantaneously from size *N*_0_ to *N_b_* and then back to *N*_0_. While this functional form is intuitive for a square bottleneck, the integrals for various cumulants depend on the in inverse of the population size 1/2*N*(*t*) (e.g., Equation 58); this would place Heaviside functions in the denominator and can thus introduce spurious divergences due to the inherent discontinuities of the Heaviside functions. To avoid this, it is more convenient to use the following definition, which is formally equivalent and preserves the interpretations of the bottleneck parameters introduced above.

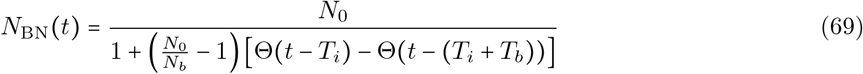

The Heaviside functions now appear in the denominator of the population size such that they are in the numerator of the dependence on 1/2*N*(*t*) in the cumulant integrals, making this parameterization substantially easier to work with analytically.

The allele frequency probability distribution in this demography has identical time dependence to that of a demography with constant population size, until the generation *T_i_* when the bottleneck begins. First, note that there are two distinct conditions on the efficacy of selection, 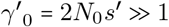, and 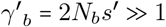. The perturbative analysis here technically breaks down during the bottleneck if 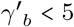, but via comparison to simulations I will show that the approximation for the first four moments detailed below is somewhat robust if the bottleneck is sufficiently short or if the population size remains within an order of magnitude of the initial size. There are additional considerations when analyzing the effects of a temporary bottleneck (i.e., one that re-expands at some point before all alleles are equilibrated to the bottleneck size). If the bottleneck duration *T_b_* is sufficiently long, the distribution equilibrates to the new population size 2*N_b_*, but the extent to which this is true also depends on the value of *s*′, which largely dictates the equilibration time [27]. For extremely high selection coefficients (i.e., nearly lethal mutations), moments of the probability distribution will equilibrate almost immediately, independent of the population size. Weaker purifying selection (while still maintaining large 2*Ns*′) implies a longer timescale before the moments settle within the bottleneck and a longer time to settle after re-expansion [27]. Additionally, if the bottleneck occurs prior to the equilibration time, this non-equilibrated state interacts with the excessive drift due to the smaller population size, which can cause additional non-monotonicity due to the increasing frequencies of polymorphic alleles during the bottleneck. This aspect is similar to the exponential population discussed above; the change in population size can disrupt the equilibration process because the equilibrium values of all moments higher than the mean are dependent on the drift coupling constant at lowest order and are thus sensitive to the time dependence of *N*_BN_(*t*).

As with all choices of demography, the lowest order behavior of the time-dependent mean frequency 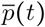 remains largely unaltered by a population bottleneck while selection remains efficient. However, the variance of the (sample) mean frequency increases dramatically during a period of reduced population size because it is proportional to the variance of the allele frequency probability distribution, as shown below/. This is visible as an increase in the sampling noise in simulations of the mean frequency presented in Figure 3 and persists across a wide range of parameters shown in Supplemental Figure S26.

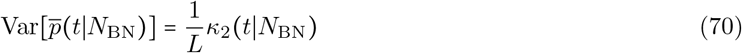

**Figure 3:**
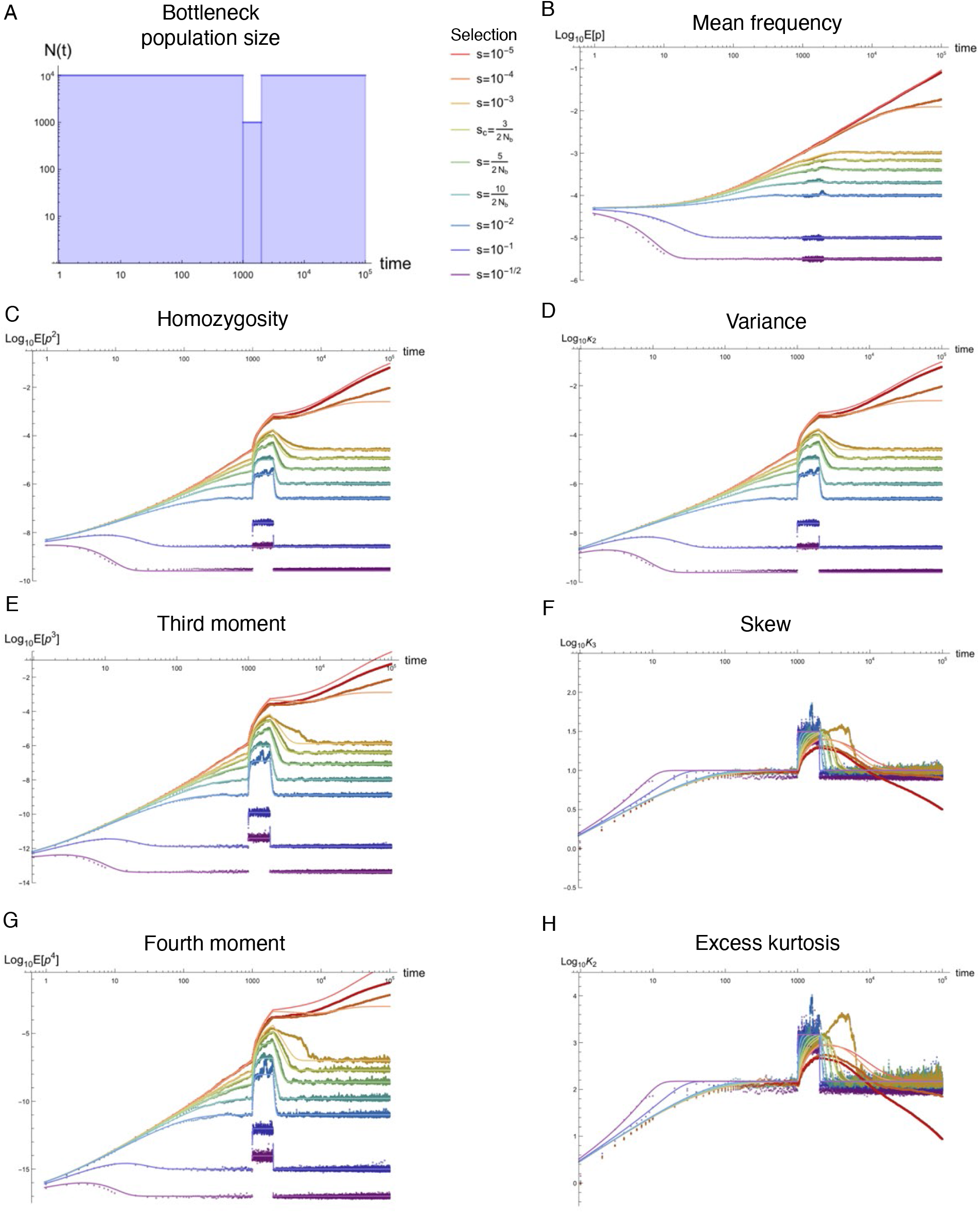
Equilibration of the first four moments, variance, skew, and excess kurtosis in population with an instantaneous bottleneck and subsequent re-expansion. From top to bottom, the left column of plots show the time dependence of the population size, homozygosity *E*[*p*^2^(*t*)], third moment *E*[*p*^3^(*t*)], and fourth moment *E*[*p*^4^(*t*)] of the allele frequency probability distribution in an exponentially expanding population. The right column of plots shows the mean 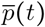, variance *κ*_2_(*t*), skew *K*_3_(*t*) = *κ*_3_/*κ*_2_^3/2^, and excess kurtosis *K*_4_(*t*) = *κ*_3_/*κ*_2_^2^ for the same parameter values. One example set of parameters is plotted with an initial population size *N*_0_ = 10^4^, *N_b_* = 10^3^, onset of the bottleneck at generation *T_i_* = 10^3^, a duration of *T_b_* = 10^3^ generations, mutation rate *μ* = 10^-6^, and back mutation rate *μ_b_* = 10^-8^; additional parameter combinations are presented in the Supplemental Material S4.3 in Figures S24–S34. Solid lines in each plot represent analytic approximations from the text with selection strengths ranging from *s* = 10^-1/2^ (purple) to *s* = 10^-5^ (red), including those near where the approximation first breaks down at roughly *s*′ ~ 5/2*N*. Scattered points represent the results of estimated moments from temporal simulations using `neqPopDynx` with the same parameter values; darker shades of the each color correspond to the analytic curve with the same selection coefficient. *L* = 10^5^ sites were simulated to estimate each moment. Note that an increased mutation rate in these plots, relative to the previous plots, allows for clearer separation between simulated selection values by reducing the variance of each moment.

Here, *L* is the number of loci, including monomorphic sites, used to compute the mean frequency, which is *L* = 10^5^ for all simulations presented in this manuscript. Since the variance increases during the bottleneck for any selection strength, the variance of the mean frequency estimated from a sample of *L* sites will increase, as well. That said, the estimated mean frequency, itself, remains well-described by the deterministic solution in Equation 26. Like exponential growth, the next three cumulants were computed from the same diagrams used for the constant population size analysis, above, but with time-dependent drift vertices (as shown in, e.g., Equations 58 and 59). The second cumulant *κ*_2_ has the following lowest-vertex dependence in a demography with a square bottleneck described by Equation 69.

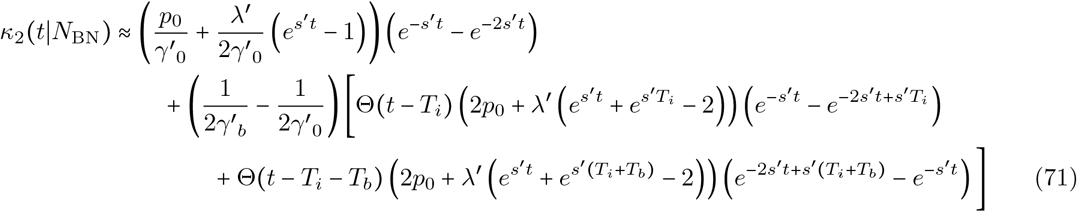

Here, Θ(*t*) is the Heaviside function and I have defined the initial population scaled selection coefficient and the bottleneck population scaled selection coefficient as 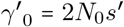 and 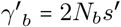, respectively. The solution naturally collects into three distinct temporal slices: prior to the bottleneck there are terms that persist throughout time, during the bottleneck there are terms proportional to Θ(*t* – *T_i_*), and after reexpansion there are terms proportional to Θ(*t* – *T_i_* – *T_b_*), where the sum (*T_i_* + *T_b_*) is the time at which the bottleneck ends (i.e., the bottleneck onset time plus the bottleneck duration). There is a distinct time dependence for the variance (and, thus, homozygosity) during each time slice. The only persistent term at late times comes from the first time slice (the *e*^*s*′*t*^ × *e*^-*s*′*t*^ term in Equation 71) and equilibrates to 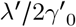, the same value for the variance of the allele frequency probability distribution in a population of constant size at asymptotically late times. In other words, the terms introduced at the beginning and end of the bottleneck are transient responses to the immediate increase and subsequent decrease in drift due to the smaller population size during the bottleneck. Figure 3 shows the dynamics of the variance (and homozygosity): beginning at the onset of the bottleneck, the variance rises exponentially towards a new equilibrium value, but this temporary increase in variance is halted when the bottleneck re-expands, after which the variance decays exponentially to return to the original equilibrium value for the larger population size (see Supplemental Figures S27 and S28 for the variance and homozygosity, respectively, plotted over a range of parameter values).

At each successive order, the resulting moments for this demography become increasingly long and tedious to write down, but contributions for each diagram can be found in Supplemental Material S3 and the full functional forms for the first four cumulants and non-central moments are directly accessible for plotting and analytic manipulation in Supplemental File S1 (for all four demographies described in this manuscript, including the bottleneck). The third cumulant, or the non-standardized skew, takes the following form after substantial simplification.

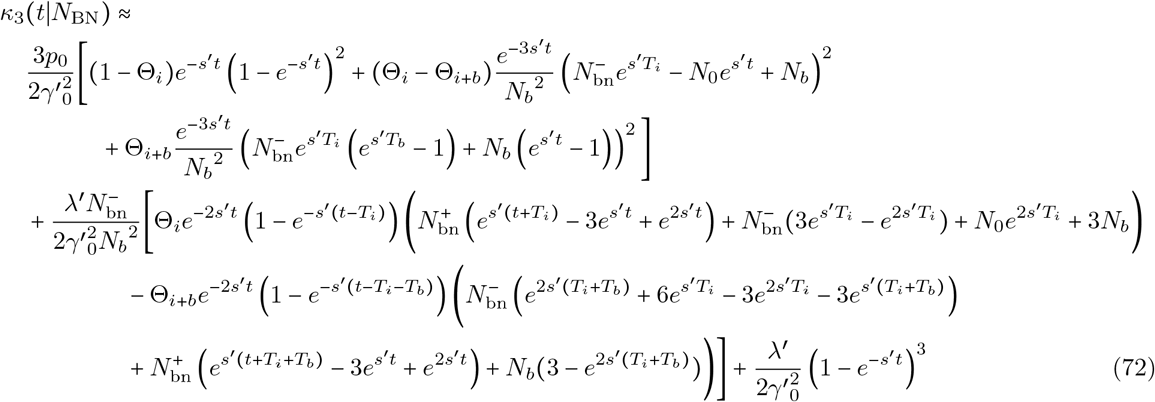

Here, I have defined 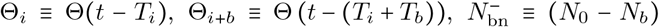, and 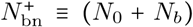 to slightly abbreviate the above expression. The fourth cumulant and moment are particularly long and space consuming to write out in full, but, for completeness, the leading order approximation for the fourth cumulant can be found in Appendix A3. All four cumulants and moments are plotted for an example parameter combination in Figure 3; plots spanning a range of bottleneck sizes, starting times, and durations provided in Supplemental Material S4.3 in Figures S24–S34 for the first four cumulants and moments, along with the standardized skew and excess kurtosis. Note that the onset of the bottleneck at time *T_i_* can occur prior to equilibration of each moment for a given selection coefficient (see, for example, *s* = 10/2*N* in Figure 3). The associated dynamics of this ‘interrupted equilibration’ are fully captured by the analytic expressions above, provided selection is within the regime of validity.

#### Cyclical demography with oscillatory population size

Cyclical demographies are comparatively less well studied in the literature relative to the demographic features described above, but regular, periodic changes in population size, amongst other quantities are known to exist in nature (e.g., [9–12]), albeit in more limited examples. Fluctuations in population size can hypothetically occur due to seasonal changes related to environmental factors like resource availability or harsher conditions (e.g., temperature or habitat availability) or due to natural organismal cycles on a multigenerational scale. Additionally, cyclical population size changes can be viewed as regular, serial bottlenecks, which have been discussed in the context of range expansion (e.g., [13–15]), for example. When viewed on the timescale of a single oscillation, the dynamics and subsequent response of the allele frequency spectrum indeed resemble that of a population bottleneck (i.e., with a temporary period of increased genetic drift) and, depending on how rapidly the population recovers, re-expansion to a maximal size can resemble brief periods of exponential expansion; the dynamics are thus linked to the two previous analyses, but there are additional observable properties unique to the periodicity, as discussed below. From a modeling standpoint, this class of demographies provides a convenient example for the application of PGFT to a relatively complex demography that can be defined using a relatively simple deterministic function for the population size. Cyclical demographies can be parameterized in a number of ways (see, e.g., [37] for a recent example), which do not all describe equivalent demographies, as the shape of the population size trajectory over the course of each oscillation may differ in different models. Here, I will analyze one class of cyclical demographies as an example, which are defined by the following three-parameter function representing the time-dependence of the diploid population size *N*(*t*).

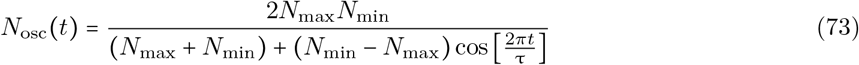

The above expression is parameterized in terms of the maximum and minimum diploid population size, *N*_max_ and *N*_min_, respectively, that each reoccur once every period of oscillation τ, the number of generations between adjacent peaks (or troughs) in the number of individuals. The maximum population size occurs at times defined by *N*_osc_(*t* = *n*τ), for all integer values of *n*, including the initial time at *n* = 0 (i.e., *N*_osc_(0) = *N*_max_). The minimum population size occurs at times *t* = *τ*/2 + *nτ* (every other half-period) such that troughs in population size *N*_osc_(*t* = *τ*/2 + *nτ*) = *N*_min_ are spaced halfway between each pair of peaks. To make integration easier, as with the bottleneck definition in Equation 69, I have chosen to place the oscillatory term in the denominator of Equation 73 (i.e., the drift coupling constants 1/4*N*(*t*) for vertex V4 are now linear in the cosine). The convenient properties of this parameterization allow for results to be computed analytically in PGFT, without resorting to numerical integration. However, this parameterization does not describe the same demography as, for example, a simple sinusoidal definition like 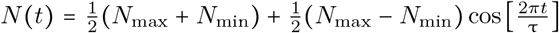 (which is not necessarily any better description of a natural population), but the defining feature of an oscillating population size remains (see Figure 4 and Supplemental Figure S35 for the general shape described by Equation 73, but note the log-log scale used to emphasize transient dynamics at early times). Relative to a function linear in the cosine, *N*_osc_(*t*) has sharper transitions to and from the peaks that more closely resemble exponential expansion (to an extent), more extended troughs that more closely resemble a prolonged bottleneck, and a different, more intuitive effective population size (see below), all of which are potentially attractive features for modeling populations. The definition of *N*_osc_(*t*) is constructed using three parameters, but as a result, the population is restricted to initially start at maximum size *N*_osc_(0) = *N*_max_; this has consequences for the transient behavior at early times, as discussed below. The initial size was an arbitrary choice (for example, the population starts at *N*_min_ if the cosine is replaced with a sine), but can be generalized to begin at any size between *N*_min_ and *N*_max_ by adding a phase *α* to the argument of the cosine (i.e, 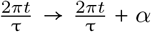; the effects of this four parameter model were not explored). Consequently, it is important to point out that some of the results discussed below are not necessarily generalizable to *all* cyclical demographies (e.g., the early-time transient dynamics, parameter regimes related to the initial population size, etc.), but various observations owing to the periodicity are likely generic. The lowest order PGFT approximations (see below) are valid for selection coefficients determined by the inverse of the effective population size 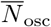 (I am using non-standard notation for *N_e_* here to avoid notational clutter), which is the harmonic mean of *N*_osc_(*t*) integrated over one period (i.e., 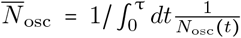). Due to the convenient properties of this particular parameterization, the effective population size is the following.

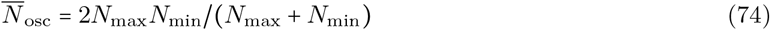

**Figure 4:**
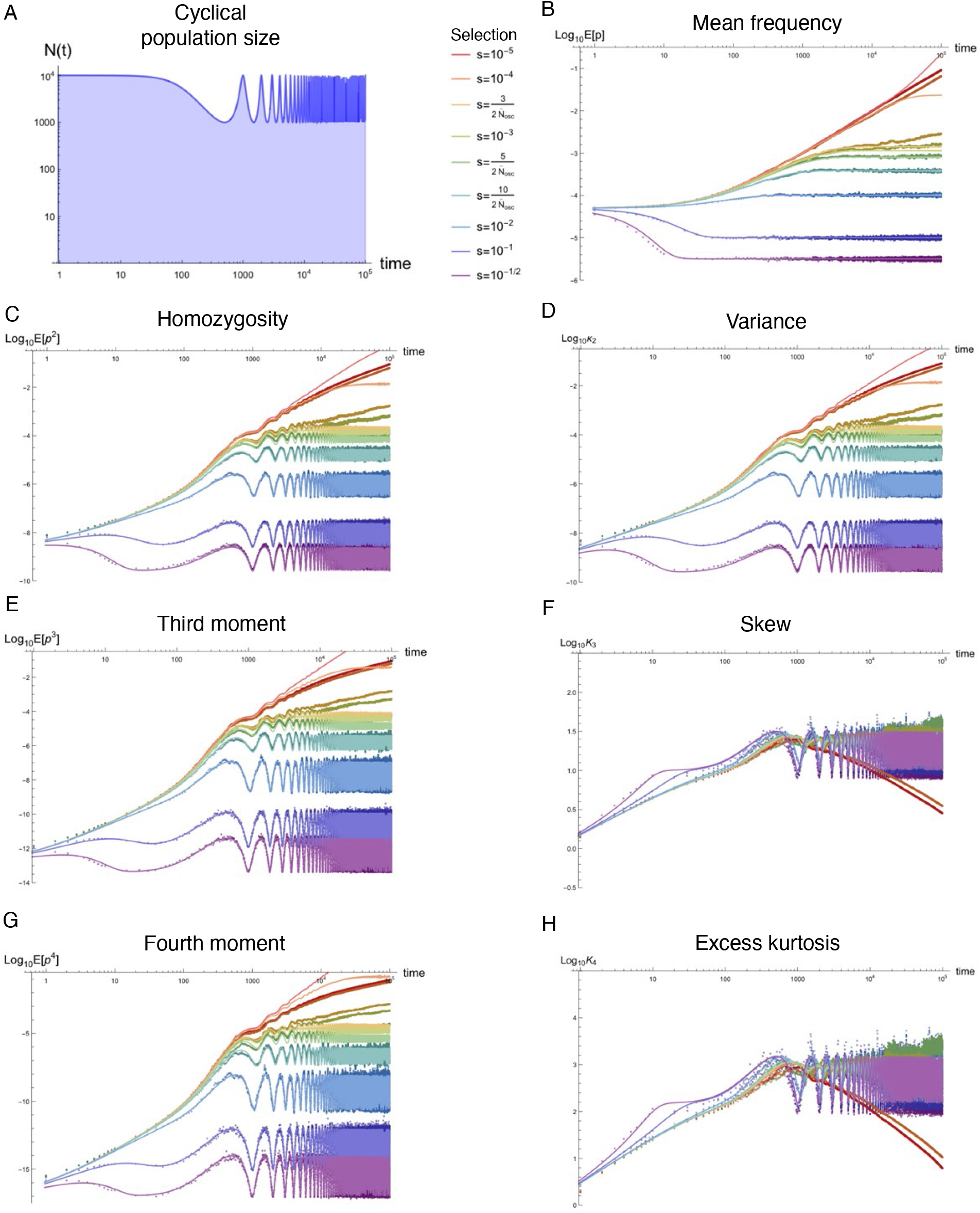
Equilibration of the first four moments, variance, skew, and excess kurtosis in cyclical population with oscillating size.. From top to bottom, the left column of plots show the time dependence of the population size, homozygosity *E*[*p*^2^(*t*)], third moment *E*[*p*^3^(*t*)], and fourth moment *E*[*p*^4^(*t*)] of the allele frequency probability distribution in a cyclical population with oscillating size. The right column of plots shows the mean 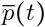, variance 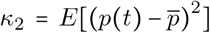, skew 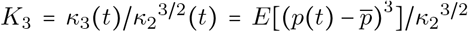, and excess kurtosis 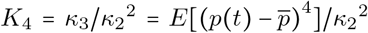 for the same parameter values. One example set of parameters is plotted with an initial population size *N*_max_ = 10^4^, *N*_min_ = 10^3^, a period of τ = 10^3^ generations, mutation rate *μ* = 10^-6^, and back mutation rate *μ_b_* = 10^-8^; additional parameter combinations are presented in Supplemental Material S4.4 in Figures S35–S44. Solid lines in each plot represent analytic approximations from the text with selection strengths ranging from *s* = 10^-1/2^ (purple) to *s* = 10^-5^ (red), including those near where the approximation first breaks down at roughly 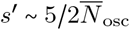 (where *s* ≈ *s*′ for the parameters plotted and 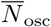 is the effective population size averaged over one oscillation). Scattered points represent the results of estimated moments from temporal simulations using `neqPopDynx` with the same parameter values; darker shades of the each color correspond to the analytic curve with the same selection coefficient. *L* = 10^5^ sites were simulated to estimate each moment. Note that the mutation rate is again increased, relative to previous plots, to reduce the variance of the moments estimated from simulation for clearer separation between selection coefficients..

In other words, 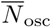 is simply the harmonic mean of *N*_min_ and *N*_min_. The regime of validity for perturbation theory is then roughly 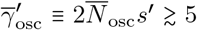, corresponding to that of a constant sized population with 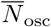 diploid individuals. To ensure that the approximations are valid at all times, one could instead require 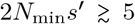, but 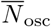 which is very close to the same boundary (due to the harmonic averaging, the effective size is dominated by the smallest population size), but 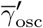 emerges from the equations below as an appropriate scale. This quantity is revisited in the context of transient dynamics and the asymptotic behavior.

Computing cumulants for this demography (i.e., using *N*_osc_(*t*)) is more laborious and time consuming than those for the previous demographies due to the trigonometric dependence. Applying the same generalized integral form for each PGFT diagram (i.e., the Heaviside integrals expressed using the free propagator), analytic results were computed using Mathematica [72] by plugging in the time-dependence defined in Equation 73 (see Supplemental File S1 for the appropriate script). The results are quite complicated and involve very long algebraic expressions, but the integral form and the associated result for each diagram can be found in Supplemental Material S3 in a form that was simplified as much as was possible. As all moments and the cumulants beyond the variance are extremely long to write out, the net time-dependence for all four cumulants and non-central moments for this demography are available in full form in Supplemental File S1, which can be accessed directly for analytic and plotting use. Despite the extended computational time to compute and simplify the appropriate Heaviside integrals, there was no need for numerical integration (i.e., the integrals were evaluated exactly), largely owing to the properties of the parameterization in Equation 73 discussed above. As a result, ignoring the unfriendly nature of these expressions, all results for this demography are variable with respect to population genetic and demographic parameters (e.g., values of *N*_max_, *N*_min_, and τ, but also *μ* and *s*), provided one remains in the regime of validity of the approximation 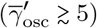. This is the primary benefit of using PGFT: the procedure only needs to be performed once to produce analytic expressions and can be used thereafter across all admissible parameter regimes (in contrast to numerical solvers for, e.g., the diffusion equation, which require computational time to solve for the dynamics for each parameter combination of interest).

As in the previous demographic models, the lowest order approximation to the mean frequency for the cyclical demography defined in Equation 73 is deterministic and thus characterized by Equation 26. The three-vertex corrections to the mean frequency can be computed by re-calculating contributions from Diagram D3 (see Supplemental Material S3.1.6 and S3.1.7) and adding the resulting expressions to to Equation 32. The full three-vertex approximation is compared to simulations in Figure 4 and Supplemental Figure S36. Unlike the deterministic approximation, and in contrast to the previous demographies, the three-vertex approximation for the mean frequency visibly oscillates in a narrow window of parameter space near the edge of the regime of validity. For example, this can be seen for a selection strength of 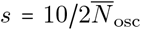 (of the same order as the edge of the regime of validity) in Supplemental Figure S36g which shows simulation results for an oscillatory population with parameters (*N*_max_ = 10^4^, *N*_min_ = 10^3^, *τ* = 10^4^). In that plot, the late-time trajectory begins to oscillate, which can be seen in both the analytic approximation and simulation output. This is the effect of the one-loop contributions to the mean frequency, which correctly characterize the emerging effects of drift as the deterministic behavior breaks down. However, relative to the magnitude of the mean, particularly in the presence of more substantial noise at lower mutation rate, the amplitude of oscillations is relatively small. Similar, drift-based corrections can be seen in the bottleneck demography, but the effect is more recognizable here due to the characteristic oscillations. In the same figure, stronger selection coefficients exhibit a distinct behavior: the variance of the mean oscillates in time. This is due to time-dependent oscillations of the allele frequency variance, which is linearly related to the variance of the mean (see Equation 70).

For this demography, all cumulants above the mean exhibit oscillatory behavior, as can be seen in Supplemental Figures S37–S43. The variance of the allele frequency probability distribution, starting from an initial frequency *p*_0_ in a demography parameterized by Equation 73 is given by the expression below. As before, this result was computed using the diagrams in Diagram D4 while integrating over the time-dependent drift coupling constant 1/4*N*_osc_(*t*) associated with the V4 vertex.

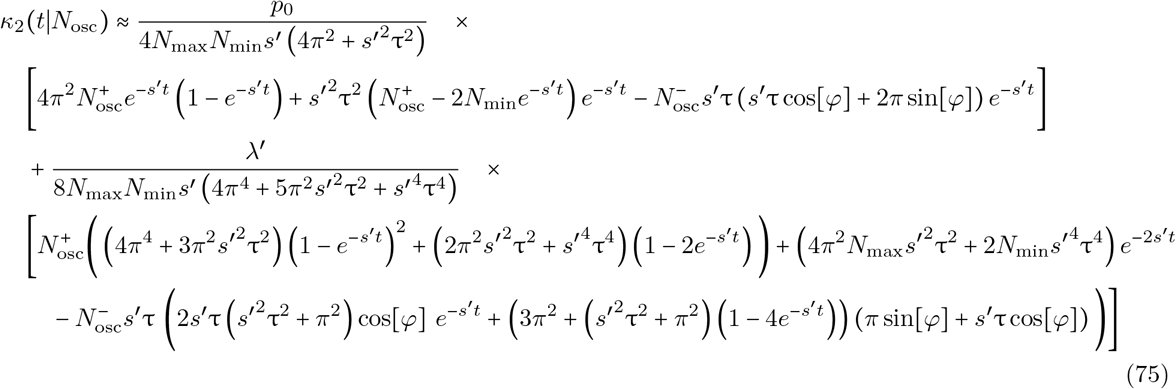

Note that I have abbreviated the above result using the definitions 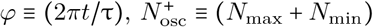, and 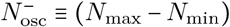. While this functional form is quite opaque, the plots of the variance in Supplemental Figure S37 (see Supplemental Figure S38 for the homozygosity) confirm that it has a few recognizable properties in asymptotic regimes. First, the variance begins at zero, as expected from the delta function initial condition. At early times, the behavior is highly variable; there are several different transient responses that occur during the first period. The effective population size is highly time-dependent until reaching the first trough, starting at an initial value of *N*_max_ and eventually reaching 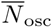 (i.e., 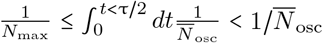; at later times, this integral resembles the lowest Bessel function, beginning at 1/*N*_max_, followed by damped oscillations that asymptote to 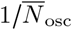 at *t* ≫ *τ*). During the first half period, the regime of validity of the approximation, and thus the minimum selection coefficient that behaves efficiently, extends below 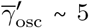 at early times. This is relevant to the transient state, as selection coefficients above this cutoff will pseudo-equilibrate to the variance corresponding very roughly to a constant population with initial population size *N*_max_ (or slightly smaller). However, assuming the initial dynamics of a constant population size of *N*_max_, the allele frequency variance due to selection equilibrates on a timescale of order 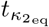 defined in Equation 54. Thus, if 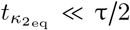, the variance will approach a fixed value before the population size reaches *N*_min_, after which drift becomes more significant and the variance increases in turn. A local variance minimization due to pseudo-equilibration at early times is seen most clearly for the *μ* = 10^-6^ plots in Supplemental Figure S37 (middle and bottom rows S37d–S37i) and occurs only for the strongest selection coefficients with *s* ≥ 10^-1^; note the flattening of the variance at early times for these selection coefficients. This effect fades entirely as the oscillation period is reduced from *τ* = 10^4^ to 10^2^ generations (moving left to right, e.g., from Supplemental Figures S37g to S37i); for τ = 10^2^, the first half period occurs on a timescale of 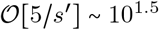. Using the parameter values for Supplemental Figures S37g, S37h, and S37i, selection coefficients of *s* = {10^-2^, 10^-1^, 10^-1/2^} have a variance that equilibrates on timescales of roughly 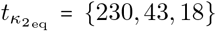 generations (see Equation 54), respectively, in a constant population of size *N*_max_ = 10^4^. When the period decreases to 100 generations, only the strongest selection coefficient has had time to equilibrate to the variance defined by *N*_osc_(0) = *N*_max_ before time *t* = τ/2. The remaining selection coefficients are overwhelmed by the decreased effective population size (i.e., increased drift coefficient) and do not pseudo-equilibrate to the initial state, but instead begin to increase in an approach a value closer to 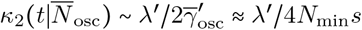. The strongest selection briefly pseudo-equilibrates until a little before the half period τ/2 = 50, but quickly equilibrates to the new value of 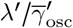 shortly thereafter; this results in an additional minimum for the strongest selection coefficient, which acts quickly enough to immediately compensate for the change in population size.

This behavior is counterintuitive: the strongest selection, near lethality, is the most sensitive to changes in population size and responds to changes in the drift coefficient; the weakest selection coefficients effectively decouple from the dynamics of the population size, instead responding to the harmonic average of these changes. However, selection coefficients between these limits appear to fall into one category or the other, depending on the period and the selection coefficient. This is a result of existence of three timescales relevant to the transient dynamics: the timescale of selection 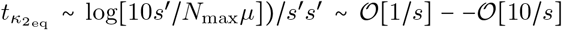 controlling the rate of exponential decay, the timescale of the effective drift coefficient 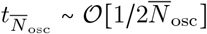 that determines the rough scale of the asymptotic time-averaged variance, and the timescale of the period τ that determines the rate of change in the drift coefficient. For the vast majority of selection coefficients, extremely rapid oscillations essentially remove the transient dynamics, as selection does not have sufficient time to act before the strength of drift shifts to a larger value of 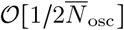. Supplemental Figures S39 and S42 confirm this is the case for higher moments, as well. In fact, the entirety of the probability distribution becomes effectively immune to τ for weaker selection, instead approaching an equilibrium distribution much like a population with constant size of 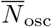. As a result, at asymptotically late times *t* → ∞, one can coarse-grain over the changes in population size to describe a steady-state equilibrium for the allele frequency probability distribution determined by mutation-selection-drift balance in a population with 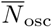 diploid individuals. This explains the lack of oscillations for weakest selection coefficients within the regime of validity 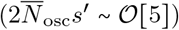 in Supplemental Figures S37f and S37i; if oscillations in the variance exist at all, their amplitude, which I have argued must be a function of *s*′*τ*, is effectively zero under weak selection. Note that this behavior is completely captured by the PGFT approximation, which matches simulations quite well (for both the variance and the amplitude of oscillations for the variance, when present). For lower values of *μ* (see Supplemental Figure S37c), the equilibration time of the variance is longer and more selection coefficients behave in this way, but this is masked by the sampling noise from fewer segregating sites. For very strong selection coefficients with *s*′*τ* > 1, the rapidity with which alleles respond to the change in effective population size allows them to constantly shift to approach new equilibria, which drags the variance into an oscillatory dance at asymptotic times *t* → ∞, constantly playing catch-up with the changing population size. Supplemental Figures S37, S39, and S42 suggest that this is true for the lowest order behavior of all cumulants, who’s dynamics are are increasingly inversely dependent on the drift vertex V4 with coupling constant 1/4*N*_osc_(*t*) (rescaled by *s*′ after integration over time), as the lowest order diagrams successively contain more drift vertices. This is well described by the time dependence in Equation 75, as can be seen from the functional form of the variance at asymptotically late times shown below.

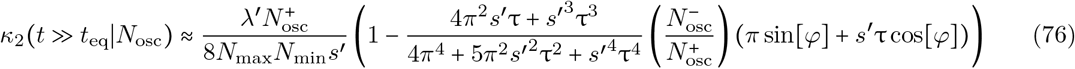

The trajectory for the variance at late times is now easily interpretable. First, the variance is oscillatory, with a period determined by the argument of the sine and cosine. The asymptotic variance oscillates around a mean value given by the constant (non-time-dependent) term in the above expression. The constant term, the average variance, is simply *λ*′(*N*_max_ + *N*_min_)/8*N*_max_*N*_min_*s*′, which, using the definition of the effective population size 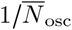 in Equation 74 (which naturally emerges in the asymptotic limit), corresponds to the variance of a population with constant size 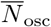 in mutation-selection-drift balance (i.e., 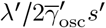). The rate of oscillations is determined by the argument of the sine and cosine functions, which is unchanged relative to the cosine in *N*_osc_(*t*) (i.e., 1/*τ*). However, instead of being proportional to a cosine alone, this is a sum of oscillatory terms, which describes a phase shift dependent on their relative weight *s*′*τ/π*. Thus, the asymptotic behavior describes a variance that lags after the change in population size, with a longer delay for weaker selection coefficients, and a maximum phase shift of *π* (definitional due to the periodicity of the sinusoidal functions). The oscillatory terms have an amplitude of oscillations determined by *s*′*τ* and 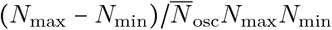 such that these oscillations vanish when selection is weak, which describes the approach to a coarse-grained distribution (determined by 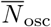), or when the population size does not fluctuate (i.e.” *N*_max_ = *N*_min_); in contrast, the oscillations are magnified when *N*_max_ ≫ *N*_min_. The eventual variance of allele frequency in a cyclical population is therefore oscillatory around the ‘effective variance’ 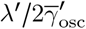. All of these properties of the oscillatory and coarse-grained behavior can be seen in plots of the analytic functions for the higher cumulants and non-central moments, as well, with primarily differ in their functional dependence on *s*′τ and 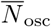 (e.g., the respective amplitudes for the oscillatory components).

Thus, after the transient phase, instead of approaching an equilibrium distribution described by a static envelope determined by 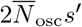 (i.e., the case for weaker selection coefficients), the allele frequency probability distribution remains in a fundamentally non-equilibrium state, constantly shifting with a time-dependent cycle akin to a perpetual waltz: the variance rises and falls at the same pace as the population size, always lagging behind by the same amount of time, keeping pace with the oscillatory dance of *N*_osc_(*t*). The third and fourth cumulants are written in full form in Appendix A3 in A3.2 and A3.3, respectively. Due to their extended functional forms, asymptotic dynamics are easiest to assess via the cumulant plots provided in Supplemental Material S4.4 in Figures S39 and S42; the third and fourth non-central moments are presented in Figures S40 and S43 and the skew and excess kurtosis are presented in Figures S41 and S44. The higher cumulants mimic the behavior of the variance, and describe a similar waltz with altered magnitudes and phases. Notably, the *standardized* moments oscillate under strong selection, as well. This indicates that the oscillatory behavior is not completely accounted for when normalizing by the appropriate power of the variance. This suggests that the Gamma function distribution for strong selection described in [34] is inapplicable in this regime, as the amplitudes and phases of the oscillations of the skew and excess kurtosis are determined by distinct dependences on *s*′τ and separately on 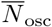, *N*_max_, and *N*_min_, Thus, in addition to the non-equilibrium oscillations of the variance, properties, like the asymmetry dictated by the skew and the relative weight of the tail determined by the excess kurtosis, waltz along in the same way, which provides a probabilistic opportunity for highly deleterious variants to be observed at uncharacteristically high frequencies.

### Discussion

This manuscript describes the construction and application of a population genetic field theory capable of generating straightforward descriptions of the temporal behavior of the allele frequency probability distribution for mutations under strong selection. For constant sized populations, this amounts to a relatively simple description of the equilibration of a newly spawned mutation to a stationary state of mutation-selection-drift balance. Three additional demographies, each specified by a deterministic time-dependent function relevant to natural populations, were chosen to elucidate the analogous processes under distinct non-equilibrium conditions; the resulting time dependences differ in end state: exponential population growth results in an approach to deterministic behavior, temporary population bottlenecks re-equilibrate to the re-expanded constant population size, while a cyclical population can result in either oscillatory or coarse-grained distributions that maintain parametric dependence on the effective population size. While the example parameterizations used here are quite specific, some of the observations are generic (e.g., transient inflation of the variance), and, more importantly, the methodology, itself, is fully generalizable. Results for any demography of interest can be computed using the same integral forms (using, for example, expressions along the lines of Equation 58, which generates the leading order contribution to the variance for a fully specifiable time-dependent population size *N*(*t*)). Thus, a wide range of non-equilibrium population genetic phenomena can be described using the population genetic field theory developed in this manuscript.

The formalism itself, i.e., field theory, is in no way novel, particularly in applications to SDEs [59]; it is simply under-explored in the discipline of theoretical population genetics (but see, e.g., [50, 51, 65]), in which many of the outstanding questions are related to non-equilibrium temporal dynamics. Furthermore, the description provided here is simply a first attempt at using this formalism to describe population genetics. A number of other phenomena of interest, including linkage disequilibrium and other multi-site dynamics, genetic dominance, the effects of inbreeding and non-random mating, clonal dynamics in asexual systems, and a number of drift-dominated dynamics, provide avenues for further applications. I have focused on the simplest possible case: a single allele under strong additive purifying selection in an obligate outcrossing species. Nonetheless, several interesting observations arise in even this simple example, including the extended equilibration time (i.e., 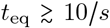, rather than the naive assumption of 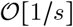 in populations of constant size, analytic approximations parameterizing the cumulants and moments of the frequency distribution, non-monotonicity of the variance (and homozygosity) during equilibration, and an approach to selection-independent values for the skew and excess kurtosis of the distribution under strong selection. Additionally, exponential growth and population bottlenecks are ubiquitous in nature [1–8], and examples of oscillatory population parameters–seasonal, in short lived organisms [9–12], or on a longer time-scale during demographic events like range expansion [13–15]–are known to occur in a range of organisms (interestingly, oscillations are more often discussed in the context of population dynamics rather than population genetics, e.g., [75]). A full understanding the associated dynamics in these demographic scenarios requires an appropriate mathematical representation of the time-dependence of the allele frequency distribution, along the lines of that presented here. Once this is accomplished, the pantheon of classical test statistics that assume equilibrium dynamics can be appropriately altered and new observables can be developed, both of which aid in our understanding of the genetics of samples from natural populations.

Towards this end, PGFT is not *a priori* necessary; there have been many attempts at describing non-equilibrium dynamics in a number of ways, ranging from a focus on fixation probability in a changing population size [38, 39] or changing environment [40] (or both simultaneously [37]), integral solutions to neutral non-equilibrium dynamics of the allele frequency spectrum [41], exact solutions for the transition density in the form of an infinite series (for constant population size [42] and variable population size [43]), a system of ODEs for the moments of the allele frequency distribution [44, 46], numerical solutions to the time-dependent diffusion equations [44–46], analysis of the stochastic dynamics of traveling waves of fitness [52], amongst many others. Several approaches involve the use of path integrals [47–51, 65], which is a natural way to characterize the trajectory that an allele takes through a population over time, also employed in the present manuscript. A particular approach of note, described in Schraiber (2014) [51], focuses on the construction of a path integral representation of the transition density for changes in the allele frequency in a finite population. As mentioned in the introduction, the formalism developed in [51] is in many ways complementary to the analysis provided here. The path integral is derived directly from the diffusion equation, with an appropriate diffusion kernel defining the transition density. In contrast, the partition functional constructed here, following the conventions in [55], uses a Langevin equation to construct an action from the stochastic differential equation, which is an equivalent, but distinct description of the diffusion limit of population genetics. The two path integrals look fundamentally different and cannot be naturally combined, but both are used in analogous ways to compute properties of the allele frequency probability distribution. Most prominently, the parameter regimes appropriate for each theory apply to distinct regimes of natural selection. The allelic dynamics of Schraiber’s path integral are assumed to follow a neutral, drift-driven path that is occasionally interrupted by the intermediary effect of natural selection, which can be expressed in a perturbative expansion in powers of the selection coefficient. This yields a controlled approximation under weak selection; in other words, the sub-drift regime emerges from the path integral as a natural perturbative regime. This complements the natural limit of the PGFT developed here, in which the strong selection regime analogously arises as a direct consequence of perturbation theory around the free action that appears in the partition functional. As a result, the dynamics between interaction times are largely deterministic, inherited from the free propagator, but allele propagation is occasionally interrupted by fluctuations from drift and mutational influx. As seen in the calculation of the all cumulants above the mean, which are at lowest order coupled to genetic drift, this formalism accounts for stochastic effects when they are relatively infrequent. The regime of validity can be extended somewhat, albeit laboriously, by computing several orders of corrections to the lowest-vertex behavior. That said, as it is currently phrased, the theory is restricted to the regime of efficient natural selection defined by *γ*′ ≳ 5. Schraiber’s perturbation series behaves similarly, as corrections can be computed by accounting for higher order terms in the 2*Ns* expansion, but the regime of validity has a hard boundary at 2*Ns* ≲ 1, beyond which perturbation theory fails to account for the full effects of selection; as it is currently phrased, the path integral also lacks a detailed incorporation of time-dependent parameters, including population size changes, which are inherently relevant to the dynamics of drift. In these ways, the two formalisms are complementary and collectively provide descriptions of the weak and strong selection limits of the diffusion approximation of population genetics. However, it is unclear exactly how the two formulations, which are very different in their partition functional (i.e., path integral), are related or if they can be combined into a stable and self-consistent field theory. To provide a unified description along these lines that describes time-dependent allele frequency dynamics across the entire range of selection strengths would be akin to writing down a systematic prescription for fully describing non-equilibrium population genetics, albeit for additive natural selection.

(e.g., to fully account for interactions between stochastic fluctuations)

The limitations to the PGFT description of population genetics presented in this manuscript are self-evident: there is no meaningful methodology developed for sub-drift natural selection (i.e., the regime where interactions between stochastic fluctuations dominate the dynamics) and many of the aforementioned complications that arise in population genetics are left out in favor of intuitive simplicity and analytic tractability. Many of these possible extensions beg to be pursued in this context. In particular, a full extension to diploid selection appears to be difficult to accomplish for technical reasons. Naively attempt to incorporate recessive natural selection using the same steps results in divergences akin to those seen in particle physics. Navigating the associated issues to provide a robust description of recessive (or general diploid) natural selection, particularly in a non-equilibrium demography, is obviously of interest, as most mutations that cause deadly phenotypes [54] or severe disease [53], arguably the mutations most likely to be under strong purifying selection, are known to be recessive. This is especially important in human genetics, where all population samples are from natural populations that underwent a recent exponential (or faster) expansion [1, 4, 6] and some of which underwent a recent or relatively recent population bottleneck. For the same reasons, understanding recessive dynamics is important to the conservation genetics of species with recent rapid decline in population size [76, 77]. Finally, it is worth noting that all of the expressions provided here are only approximate, although they appear to be largely accurate approximations over a sizeable range of parameters (provided 2*Ns*′ > 5).

While there are a few established ways to solve the time-dependent diffusion equation, use of numerical solutions, in particular-perhaps the most practically useful methods for empirical analysis and inference–provide a rather opaque understanding of the parameters controlling the dynamics, as well as for generalization across variations in these parameters. Infinite series solutions suffer in similar ways, unless the series can be truncated sufficiently quickly to provide a tractable approximate expression. In this sense, while less exact and in the strong selection limit, the closed form approximations provided here may serve to elucidate features of the dynamics, such as the equilibration time or the time of maximal variance and homozygosity. Hopefully, extensions of this analysis will provide a broader intuition for these quantities under weaker selection and when other complexities observed in nature are introduced.

## Data availability

Supplemental Material is available on FigShare. The following supplemental material is available for this manuscript: Supplemental Material S1 contains Supplemental Tables S1–S3 defining quantities and parameters referred to in this manuscript. Supplemental Material S2 contains a brief note on the evaluation of functional derivatives. Supplemental Material S3 contains detailed calculations for all Feynman diagrams that appear in this manuscript. Supplemental Material S4 contains Supplemental Figures S1-S44 that compare analytic results to population genetic simulations over a wide range of parameters for each demographic scenario discussed in this manuscript (Supplemental Material S4.1: constant population sizes; S4.2: exponential growth; S4.3: population bottlenecks; S4.4: cyclical demographies). Supplemental Material S5 contains a brief note on generalized results for arbitrary cumulants *κ_n_*. Supplemental File S1 (described in Materials and Methods on page 18) is available on FigShare and is a Mathematica notebook that contains all relevant Feynman diagram integrals and analytic expressions for the first four cumulants and non-central moments for each of the four demographies discussed in this manuscript All simulations (detailed in Materials and Methods on pages 18–20) were performed using `neqPopDynx`, a Wright-Fisher population genetic simulator custom written in Python 3 for this manuscript. Simulator details are detailed in Materials and Methods on pages 18–20. The current version (v1.5) of this script is freely available for download on GitHub at https://github.com/dbalick/neqPopDynx and at http://genetics.bwh.harvard.edu/wiki/sunyaevlab/dbalick; a final version will be made publicly available at the time of publication.

## Acknowledgements

The author would like to sincerely thank Joshua Schraiber and Shamil Sunyaev for many helpful discussions during the preparation of this manuscript. Additionally, the author would like to acknowledge the late Joseph Polchinski, who’s instruction many years ago was foundational to his (comparatively modest) understanding of non-equilibrium field theory, without which this manuscript could not have been written.

## Funding

This research was supported through R35GM127131 from the National Institute of General Medical Sciences of the National Institutes of Health under the supervision of Shamil Sunyaev.

## Conflicts of interest

The author declares no conflicting interests.

## Appendix

### A1 Derivation of the action and partition functional from the Langevin equation

Here I will derive the partition functional, the primary object used for computation in field theory, from the Langevin equation. The derivation below largely follows the derivation of Chow and Buice (2015) [55]. For a more detailed and accessible introduction to the construction of path integral formulations of stochastic differential equations, please see their review. An analogous derivation can be found in Wio (2013) [59], which constructs a different class of path integrals (that more closely resemble the formulation in [51]); this contains a more detailed discussion of the use of various discretization conventions, a historical presentation of the application of path integrals to stochastic processes, and covers a wider range of topics in depth.

Starting from the Langevin formalism, Equation 5 can be rewritten by putting all terms on the left hand side and separating the linear terms for later convenience. As in the manuscript text, 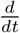 will be written as *∂_t_* for notational compactness.

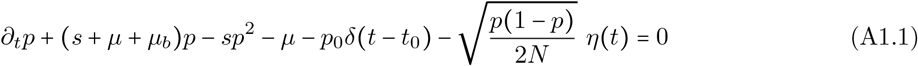

Note that the initial condition *p*_0_*δ*(*t* – *t*_0_) only affects the time point *t* = *t*_0_ at which *p*(*t* = *t*_0_) = *p*_0_. For notational simplicity, I have again defined the parameter *s*′ ≡ (*s* + *μ* + *μ_b_*), which plays an important role as the effective selection coefficient, summarizing a reduction in frequency due to natural selection Δ*p* ~ −*sp*, saturation of the number of mutated individuals Δ*p* ~ −*μp*, and the loss of frequency due to back mutations from the derived allele *p* to the wild-type allele (1 – *p*) at a rate Δ*p* ~ −*μ_b_p*. At this point it is convenient to separate and collect the terms that include *η*(*t*) and those that do not.

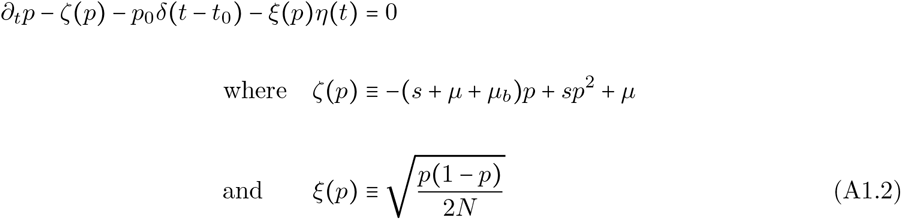

This can be rewritten in differential form, noting that the Gaussian white noise term *η*(*t*) defines a Wiener process *W* such that the time differential is *dtη* → *dW* (very loosely speaking, like Brownian motion, *η*(*t*) scales with 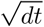). In this infinitesimal limit, the Dirac delta function becomes a Kronecker delta symbol *δ*_*t*,*t*_0__, which only evaluates to one when *t* = *t*_0_ (formally this occurs after discretization, but I have separated each step for clarity).

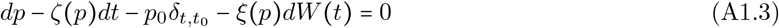

The infinitesimal time dependence *dt* of the function *ζ*(*p*(*t*)) has been extracted in this equation, such that *ζ*(*p*) now simply represents the magnitude of *ζ* at time *t*. This equation can be discretized as follows, but importantly I will use the Itô (or *prepoint*) discretization convention to avoid tedium later. In this convention, the discretization step size *dt* → *ϵ*, and the symbols *ζ_k_* and *ξ_k_* represent the evaluation of the functions *ζ*(*p*(*t*)) and *ξ*(*p*(*t*)), respectively, at time step *k*. The finite time interval (*t* – *t*_0_) → *T* is discretized into *K* steps of size *ξ* such that *T* = *Kξ* = *const* for any choice of *ξ*. The stochastic increment 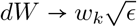 when discretized, displaying the square root dependence on the step size.

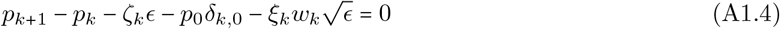

Here, the initial time *t*_0_ is defined as *k* = 0 for convenience. The discrete random variables *w_k_* for each *k* now take the place of *η*(*t*) (up to 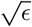) and are normally distributed with mean *E*[*w_k_*] = 0 and *E*[*w_i_w_j_*] = *δ_i,j_*. Discretization using the Stratonovich (or *midpoint*) convention would have resulted in 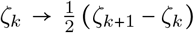, amongst several other differences at a later point, which will not be elaborated on henceforth (more details can be found in [59]).

At this point, the above describes a vector of time points with frequencies 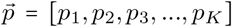 and corresponding stochastic random variables 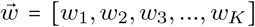 for each time step that are related by Equation A1.4. A probability density function, which should be interpreted as related to the transition density from (*p*_0_, *t*_0_) to (*p, t*) (conditional on a sequence of stochastic variables {*w_k_*}), can be constructed from this equation by introducing a Dirac delta function with an argument corresponding to the solution to Equation A1.4.

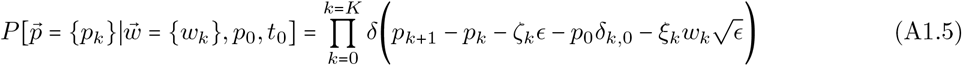

For every discrete time point *k*, the values of {*p_k_*} are restricted to those which satisfy Equation A1.4. Additionally, the values of *p_k_* are not independent (due to the dependence on *p*_*k*+1_) and are dependent on the discrete stochastic variable *w_k_*. In other words, the delta function constrains the values of the discrete frequencies *p_k_* to those that solve the discrete dynamical equation derived form the Langevin equation. The delta functions can be expressed via Fourier transform as follows.

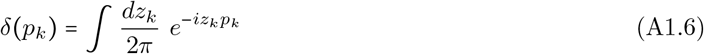

Fourier transforming for each time point, the following probability density function emerges.

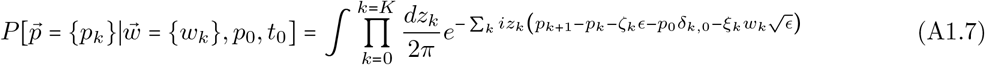

Here, the sum over the interval *k* ∈ [0, *K*] in the exponential comes from adding the exponentials for each of the *K* Fourier transforms. Since the *w_k_* happen to be Gaussian distributed for each value of *k*, one can integrate over the probability density of the noise terms to find a probability density for 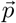 that is independent of 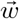, effectively by averaging over the noise at each discrete time *k* (referred to as *integrating out* the variables *w_k_* in field theory).

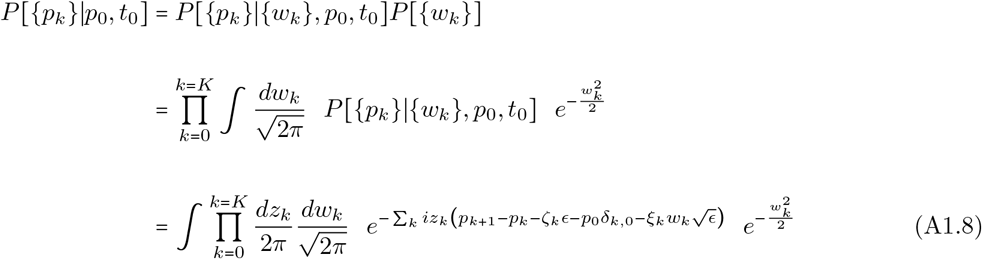

At this point, note that the linear dependence on *w_k_* in the original Fourier transformed delta function can be combined with the Gaussian integral by completing the square. Taking one *w_k_* at a time, the following integral can be performed.

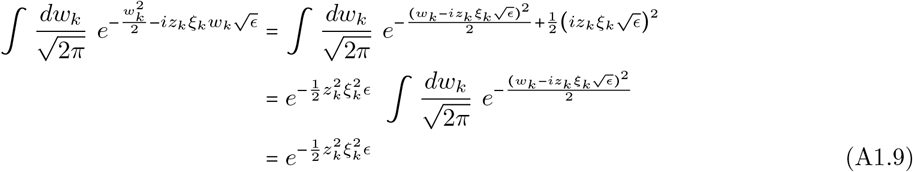

Thus, each *w_k_* can be integrated out completely and replaced with the remaining exponential from completing the square. Noting that 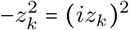, Equation A1.8, can be re-expressed as follows.

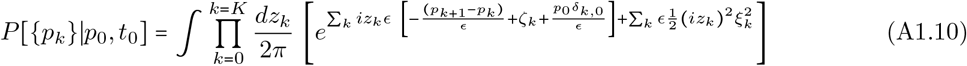

This expression is now ready to be taken to the continuum limit by taking *ξ* → 0 while *K* → ∞ such that *T* remains finite and constant. As it represents the dual variable to *p*, the variable *iz_k_* can be redefined as 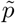, the significance of which is interpreted in the manuscript text. The continuum expression after this change of variables and replacing *ζ_k_* and *ξ_k_* with the continuum *ζ*(*p*) = *s*′*p* – *sp*^2^ – *μ* and 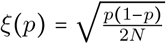 is the following.

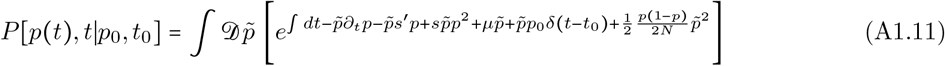

Finally, the exponential expression can be manipulated to collect the linear terms, which are of particular interest, and expand the terms related to drift.

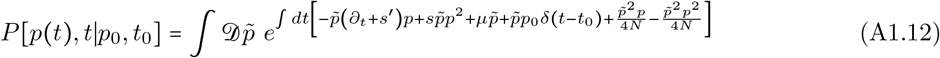

This is now a path integral describing the transition density *P*[*p, t*|*p*_0_, *t*_0_]. To formulate a partition functional, usually referred to as the moment generating functional in the stochastic context, one can take the Laplace transform (of the path integrand of Equation A1.12) with respect to source functions 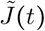 and *J*(*t*), transforming *p*(*t*) and 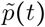, respectively. The partition functional can be expressed as follows.

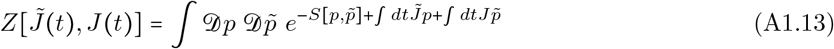

Here, the *action* 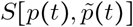 has been defined, which both describes the model of interest (e.g., the diffusion or Langevin equation can be directly written down from the action) and provides the weighting scheme for all possible paths, in this case, paths of the allele frequency. The relevant action for the population genetic field theory (PGFT) can be read from the exponentiated part of Equation A1.12.

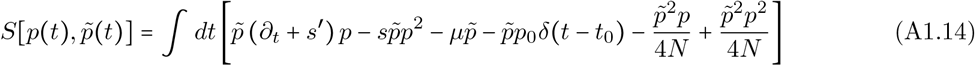

Note the sign change relative to Equation A1.12 due to the convention of defining the action as the negative of the exponentiated functional in Equation A1.13. Again, *s*′ is the effective selection due to the combination of selection and mutational effects removing derived alleles.

#### A1.1 Normalization of the path integral, the Itô discretization scheme, and loop diagrams

In quantum field theories there is usually a normalization factor for the partition functional that can be particularly useful for cancelling divergences that arise from computing various loop diagrams (due to integrating over all possible values of the momenta of internal propagators) [60, 61]. The normalization factor, often denoted as 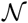, can be computed from a series of diagrams describing the vacuum state of the theory, each of which takes the form of a diagrams with no external vertices; no 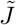 or *J* functional derivatives are applied for a diagram with no legs, so the vacuum energy is the evaluation of the path integral over the action (i.e., 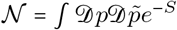, which can be computed perturbatively) [60, 61]. The divergent contributions that pop up in loop diagrams happen to exactly cancel the divergences of the vacua, since an observable of interest now contains an overall factor of 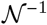.

The population genetic field theory of interest to this manuscript does not contain such divergences to begin with. The normalization turns out to be determined only by the discretization scheme chosen: pre-point (the Itô discretization convention used here), midpoint (Stratonovich convention), or endpoint discretization schemes [59]. Use of a different discretization scheme makes the normalization of the partition functional depend on a discretization parameter (1/2 for the Stratonovich convention), but this should be cancelled by additional factors in the action and thus in the results of computing a given moment [59]. Instead, the Itô scheme yields the simplest normalization to work with.

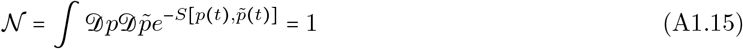

Here, the action 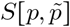 is the action in Equation 6. This discretization scheme also partially determines the free action (specifically, its lack of dependence on the discretization parameter, amongst other complications) and thus the propagator. Solving the Green’s function for the propagator yields Equation 14, which is proportional to Θ(*t*_2_ – *t*_1_); importantly, Θ(0) = 0 such that *G*(0) = 0. This fact ensures that all propagators must be time-ordered and therefore both that they cannot begin and end at the same vertex and that they cannot point backwards in time [55]. As a result, diagrams for PGFT are subject to <u>Rule 5</u> (i.e., diagrams that are not time-ordered evaluate to zero), which severely limits the allowed structures for loop diagrams and is a consequence of the Itô condition.

### A2 Calculating counting factors using Wick contractions

Much of the following discussion can be found in Peskin and Schroeder [60], along with a more extended discussion on the interpretation and application of Wick’s Theorem. As discussed above in <u>Rule 3</u>, the number of diagrams that can be constructed with identical topology provides an overall counting factor multiplying the contribution represented by each diagram. Wick contractions, in this context, amount to the number of possible ways to replace pairs of *p*(*t*_2_) and 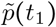 with a propagator *G*(*t*_2_, *t*_1_), where *t*_2_ > *t*_1_, indicating propagation of the free action between these time points. This can be visualized by writing out the integral form of each vertex for a given diagram and finding unique ways to pair all connections *p* and 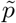 contained within.

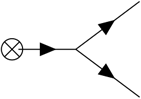

For example, for the right diagram in Diagram D4, repeated above for clarity, one can write down all vertices used to construct the diagram as follows. I will separate each vertex with whitespace denoting integrals over the same time within each block and over different times between blocks (e..g, *t*_a_, *t_b_*).

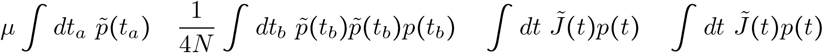

The integrals and coupling constants can be dropped for the present purposes and each group of connections *p* and 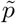 separated by a vertical line are implied to have the same time coordinate.

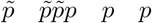

There are two distinct sets of contractions for this diagram, each of which is denoted by a specific set of paired braces, as shown below.

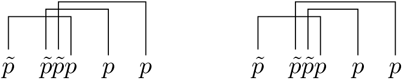

Here, each bracket corresponds to a propagator, and each grouping of connections corresponds to a vertex. The distinction between these sets of contractions is the choice of which copy of 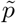 in the drift vertex (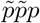 in the notation used above) to pair to the first, as opposed to the second, external vertex (*p* in this notation) on the right side. Note that the following contraction is disallowed.

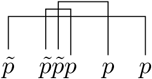

There are two reasons for why this contraction does not contribute to the counting factor for this diagram. First, the topology represented by this contraction differs from the initial two such that it corresponds to a completely different (and disconnected) diagram, shown below.

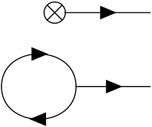

The second issue is that there is a contraction between connections in a single vertex, implying a closed loop, as seen in the above diagram. This is disallowed by <u>Rule 5</u>, which dictates that any diagram (or, in this case, Wick contraction represented by a diagram) with a propagator that starts and ends at a single vertex evaluates to zero. This is clearly not the case for Diagram D4, as can be seen in Equation 34. Thus, the overall counting factor for this diagram is two, corresponding to two distinct Wick contractions.

This method is particularly useful for determining the counting factor for more complex diagrams, which tend to be less obvious by inspection. As an example, the left side of Diagram D7 is repeated below.

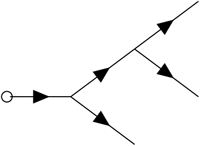

The counting factor for this diagram is 2 × 2! × 3!, which may be difficult to identify without showing the Wick contractions.

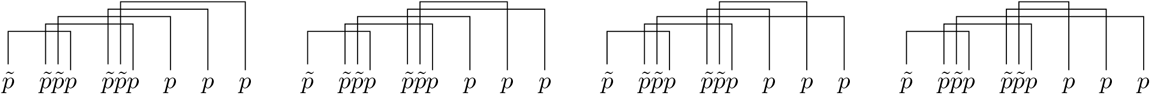

The first two sets of contractions are the same other than swapping the connections between the rightmost drift vertex and the first two external vertices. In the third set, the three connections to the external vertices are reordered (the third is connected to the first drift vertex, rather than the first). The fourth set is the same as the third, but with the two connections from the final drift vertex to the first two external vertices reversed. The remaining reordering of the external vertices is captured in the following two contractions, making six re-orderings in all.

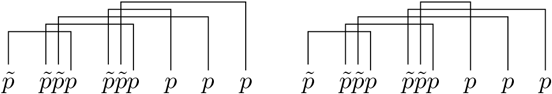

There are six additional sets of contractions that are identical to these other than changing which copy of 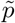 in the first (leftmost) drift vertex is connected to the external vertex. For example, the first set of contractions above (left) becomes the following set.

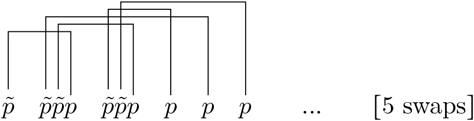

Here, ‘5 swaps’ represents the five additional sets of contractions with this reversal, for a total of 12 sets of Wick contractions thus far. Finally, all twelve of the above can be rewritten by swapping which of the two drift vertices is connected to the terminal vertex (a *p*_0_ vertex in this case) and which is connected to the external vertices. For example, the very first contraction is repeated below with swapped drift vertices.

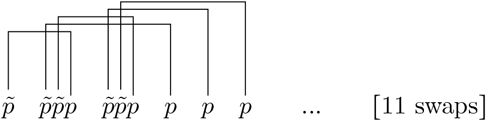

Again, ‘11 swaps’ represents the additional eleven contractions that result in a total of 24 unique sets of Wick contractions that produce diagrams of the same topology. Thus, Diagram D7 has an overall counting factor of 2 × 2! × 3!: the 3! comes from re-arranging the order of propagators exiting the internal vertices and connecting to the external vertices; the factor of 2 comes from swapping the connections of the first drift vertex; the factor of 2! comes from swapping the drift vertices entirely, of which there are two in this example. This type of counting, by identifying the *symmetries* of the Wick contractions, substantially reduces the time required to compute counting factors. Note that in the example of Diagram D7, there are two (repeated) drift vertices, which results in a factor of 1/2! from <u>Rule 2</u>. This cancels with the factor of 2! from swapping the drift vertices; all diagrams with repeated vertices have an analogous cancellation.

### A3 Higher cumulants for bottleneck and cyclical demographies

Given the extended functional form of the fourth cumulant in the bottleneck demography and both the third and fourth cumulants in the cyclical demography, the final expressions do not appear in the main text. These results are printed in full form here, for completeness, and are based on summing the appropriate diagram integrals, separately computed for each demography, each of which is written explicitly in Supplemental Material S3 (in integral form and calculated result). However, due to the length of these expressions, Supplemental File S1 should be used to directly access all of the following cumulants and non-central moments in a form that can be further manipulated and plotted in Mathematica [72]. The third cumulant in the cyclical demography is given by summing the contributions represented by Diagram D7 (see Supplemental Material S3.3.1 and S3.3.2). For both demographies, the fourth cumulant is given by the following sum of contributions from Diagrams D8 and D9.

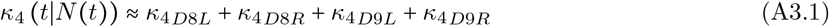

The expressions appearing in this sum are given by the results printed in Supplemental Material S3.4.1, S3.4.2, S3.4.3, and S3.4.4. Note that the following expressions have been simplified as much as was possible (which is not much) using the `FullSimplify` function in Mathematica followed by manual manipulation and abbreviation (via definitions provided below and defined in Supplemental Table S3) to make them more compact. To access these expressions prior to simplification, please see Supplemental File S1.

#### A3.1 Fourth cumulant in the presence of a population bottleneck

For convenience, the following expressions are again abbreviated using the definitions 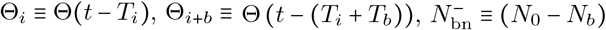, and 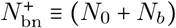.

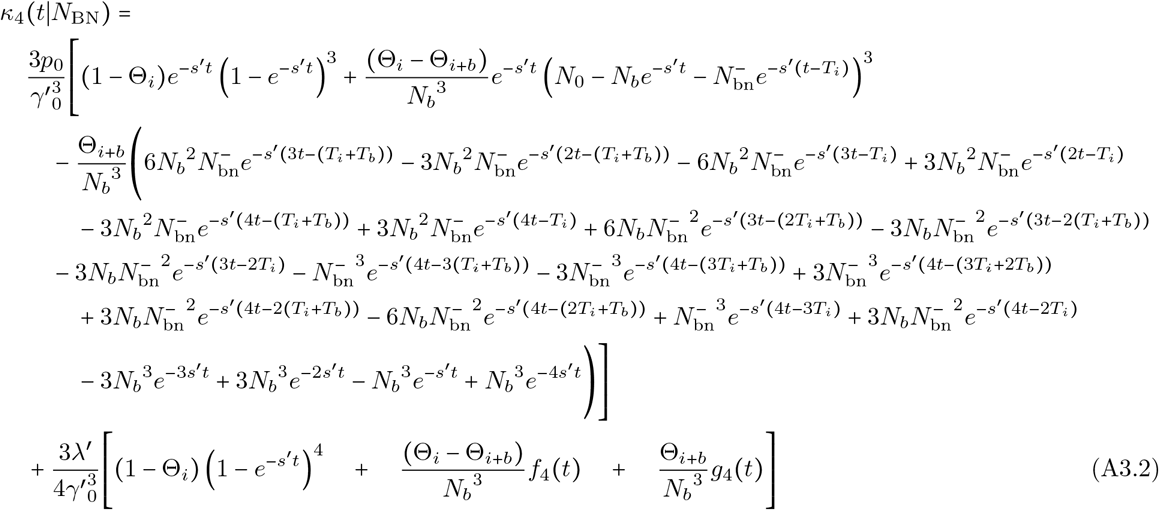

Here, the functions *f*_4_ and *g*_4_ are defined as follows.

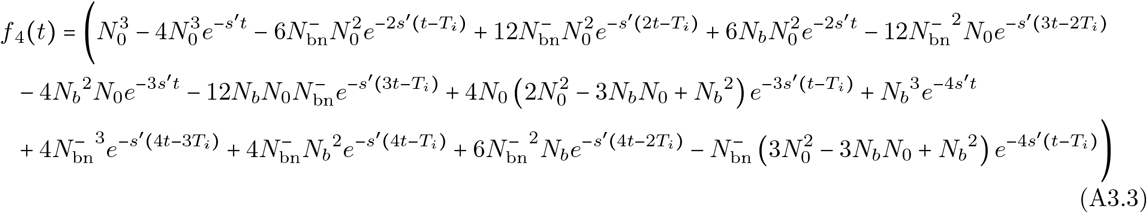

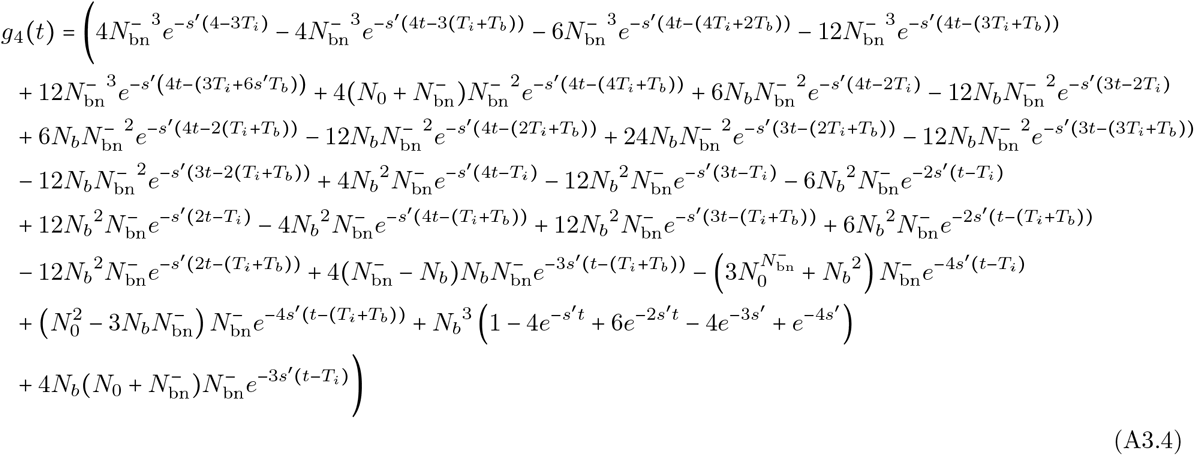

Note the overall prefactor of 3/2 the expressions in Equation A3.2 relative to the diagram results for D8 in Supplemental Material S3.4.1 and S3.4.2. This is due to the sum over both Diagrams D8 and D9, the results of which have the same functional dependence up to an overall constant. The fourth non-central moment can be expressed in terms of the above result, the third cumulant in Equation 72, the variance in Equation 71, and the mean frequency in Equation 26 using the relationship 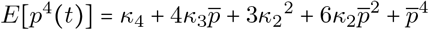 to define the fourth moment in terms of all lower cumulants. Here, 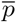, the mean frequency, is also the first cumulant of the allele frequency probability distribution.

#### A3.2 Third cumulant in a cyclical population

Due to the extended forms of all cumulants in a cyclical demography defined by Equation 73, the following results have been abbreviated using the definitions 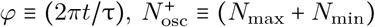, and 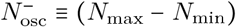. The time dependence of the third cumulant is given by the following expression, broken into slightly more palatable components.

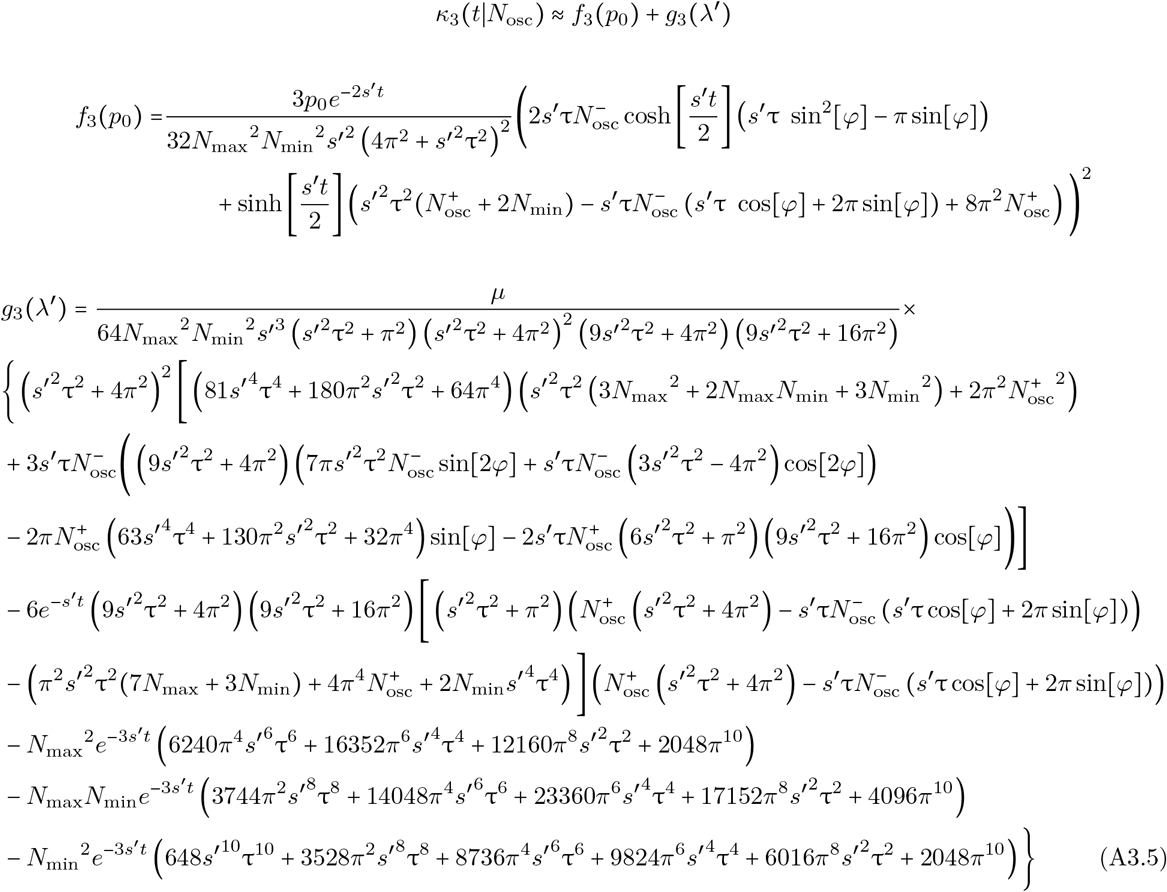

The third non-central moment can be expressed in terms of the above result, the variance in Equation 75, and the mean frequency in Equation 26 using the relationship 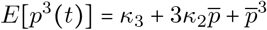 to define the third moment in terms of all lower cumulants. Here, 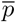, the mean frequency, is also the first cumulant of the allele frequency probability distribution.

#### A3.3 Fourth cumulant in a cyclical population

As for the third cumulant, the following expressions are abbreviated using the definitions 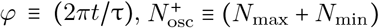, and 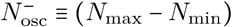.

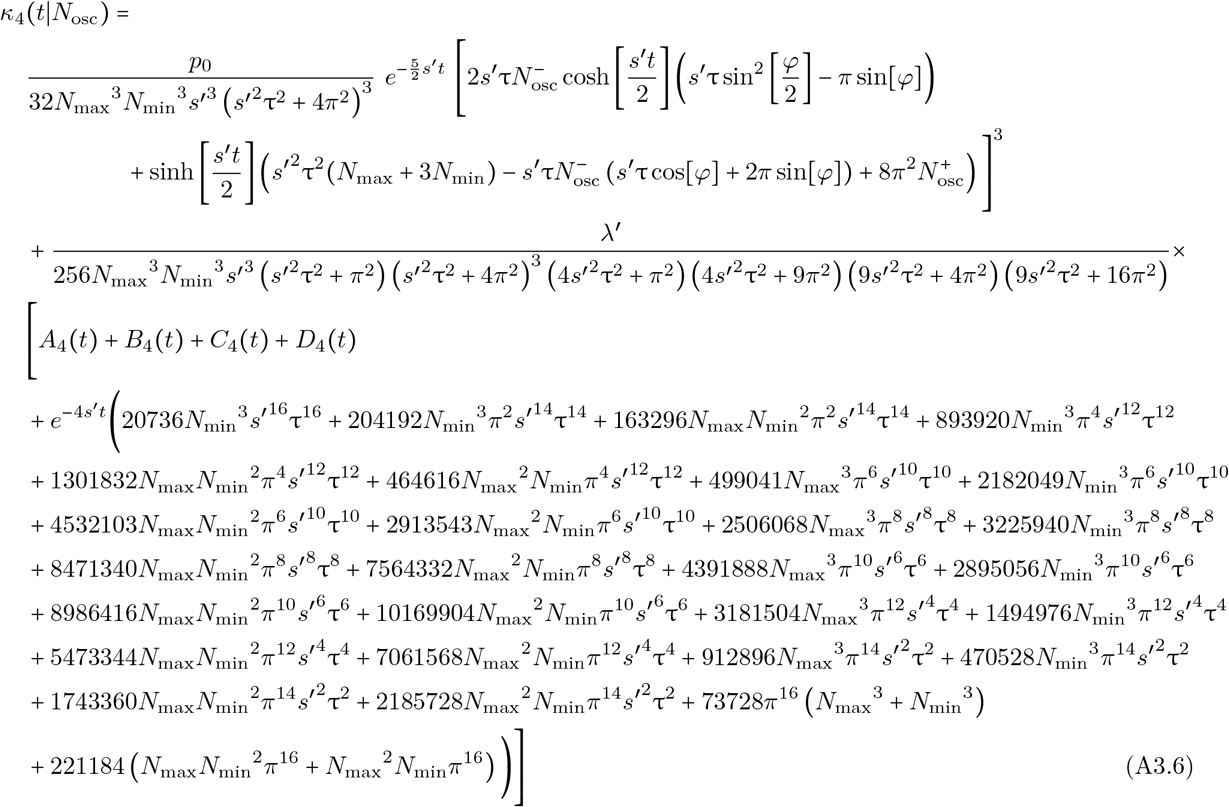

The functions *A*_4_, *B*_4_, *C*_4_, and *D*_4_ are defined as follows.

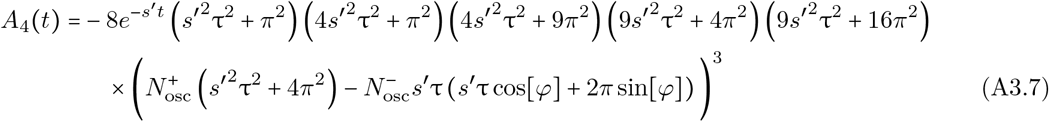

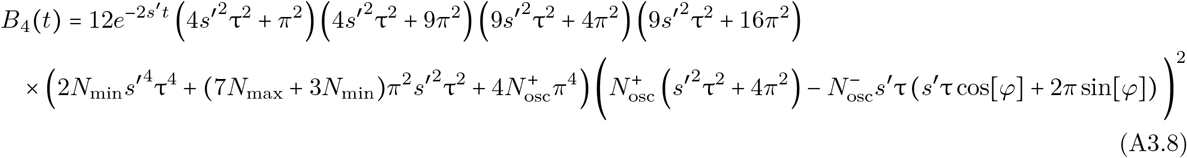

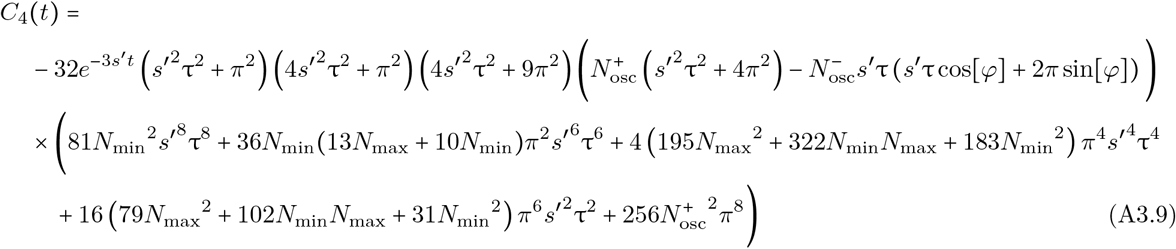

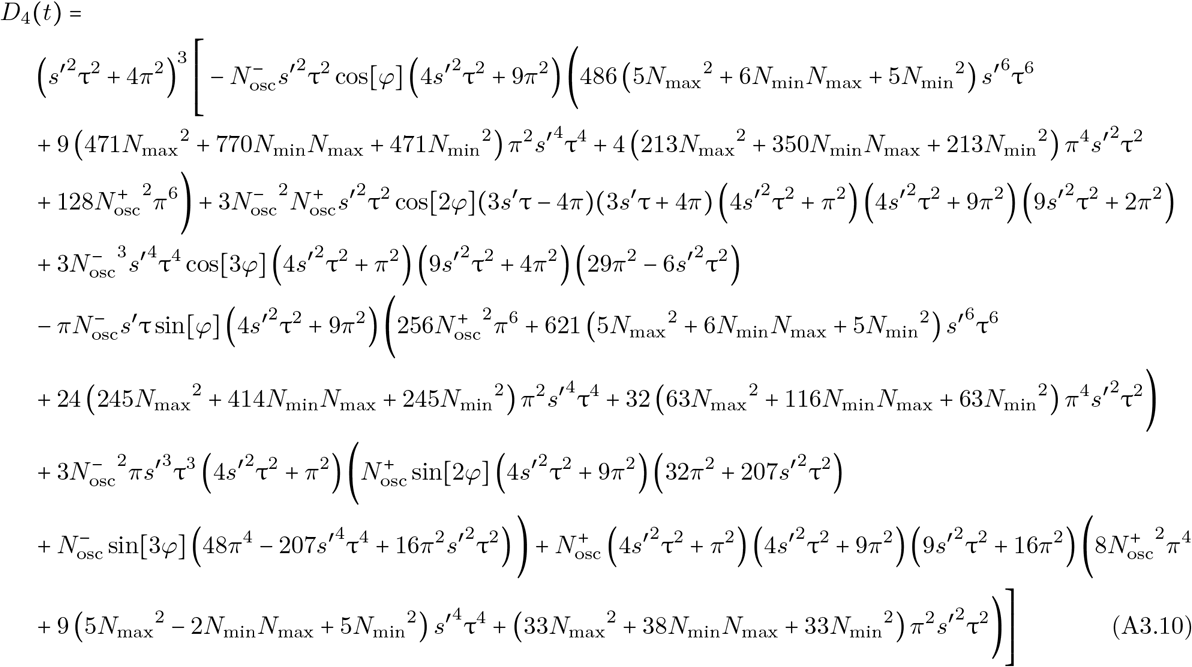

An overall factor of 3/2 again appears from the sum of Diagrams D8 and D9, which represent contributions with the same functional dependence up to an overall constant. The fourth non-central moment can be expressed in terms of the above result, the result in Appendix A3.2, the variance in Equation 75, and the mean frequency in Equation 26 using the relationship 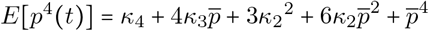 to define the fourth moment in terms of all lower cumulants. Here, 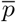, the mean frequency, is also the first cumulant of the allele frequency probability distribution.

## Notes

### Competing Interest Statement

The authors have declared no competing interest.

