## Supplemental Material for "A field theoretic approach to non-equilibrium population genetics in the strong selection regime"

for

Daniel J. Balick

#### Table of Contents

S1 Supplemental tables: quantity and parameter definitions, *p.S2*
Table S1: Function, functional, and symbol definitions, *p.S2*
Table S2: Population parameters definitions, *p.S4*
Table S3: Demography-specific parameter definitions, *p.S5*
S2 A brief note on evaluating functional derivatives, *p.S6*
S3 Feynman diagrams and corresponding integrals, *p.S7*
S3.1 Diagrams for the mean frequency $\overline{p}(t)$, *p.S8*
S3.2 Diagrams for the second cumulant $\kappa_2(t)$, *p.S11*
S3.3 Diagrams for the third cumulant $\kappa_3(t)$, *p.S20*
S3.4 Diagrams for the fourth cumulant $\kappa_4(t)$, *p.S23*
S4 Comparison of analytic approximations to simulations over a range of parameters, *p.S31*
S4.1 Constant population size, *p.S31*
S4.1.1 Constant demography: time dependence of the population size, *p.S32*
S4.1.2 Constant demography: mean frequency $\overline{p}(t)$ to one-vertex order, *p.S33*
...
S4.1.12 Constant demography: excess kurtosis $K_4(t)$ to four-vertex order, *p.S43*
S4.2 Exponential population growth, *p.S44*
S4.2.1 Exponential demography: time dependence of the population size, *p.S45*
S4.2.2 Exponential demography: mean frequency $\overline{p}(t)$ to three-vertex order, *p.S46*
...
S4.2.10 Exponential demography: excess kurtosis $K_4(t)$ to four-vertex order, *p.S54*
S4.3 A population bottleneck and re-expansion, *p.S55*
S4.3.1 Bottleneck demography: time dependence of the population size, *p.S57*
S4.3.2 Bottleneck demography: mean frequency $\overline{p}(t)$ to three-vertex order, *p.S58*
...
S4.3.10 Bottleneck demography: excess kurtosis $K_4(t)$ to four-vertex order, *p.S66*
S4.4 Oscillating population size, *p.S67*
S4.4.1 Cyclical demography: time dependence of the population size, *p.S69*
S4.4.2 Cyclical demography: mean frequency $\overline{p}(t)$ to three-vertex order, *p.S70*
...
S4.4.10 Cyclical demography: excess kurtosis $K_4(t)$ to four-vertex order, *p.S78*
S5 On the computation of the fourth and higher cumulants of the allele frequency distribution, *p.S79*
S5.3Literature cited in Supplemental Material, *p.S82*

### S1 Supplemental tables

Table S1: *Function, functional, and symbol definitions*

| | |
|---|---|
| $\delta(t-t')$ | Dirac delta function centered at time $(t-t')$. |
| $\Theta(t-t')$ | Heaviside function; defined in this manuscript using $\Theta(0)=0$. |
| $\frac{\delta^k}{\delta f(t)^k}$ | The $k^{\text{th}}$ functional derivative of a function $f(t)$; acts on a functional, analogous to a partial derivative. See Section S2. |
| $p(t)$ | Allele frequency field; time-dependent function treated as a field, a scalar valued function at all time points $t$, where $t \geq t_0(=0)$ is the generation time in the continuum limit. |
| $p_0$ | Initial allele frequency $p_0 \equiv p(t_0)$ (setting $t_0 = 0$ for convenience); treated as a tunable parameter, but various approximations and simulation comparisons assume the initial frequency $p_0 = 1/2N$ (i.e., a single allele in one haploid individual) for convenience and biological relevance to new mutations. |
| $\tilde{p}(t)$ | Frequency dual field; time-dependent function treated as a field. Defined by taking the continuum time limit of $\tilde{p}(t_i) = iz(t_i)$; here, $t_i$ represents discrete generation time and $z(t_i)$ is the Fourier transform of allele frequency $p(t_i)$ at each time point $t_i$. This is the dual field to frequency $p(t)$ such that an allele frequency path between times $t_1$ and $t_2$ originates at $\tilde{p}(t_1)$ and ends at $p(t_2)$; analogous to the relationship between $x$ and $p$ in the forward diffusion equation for $\phi(p, t_2 \mid x, t_1)$, where $p(t_1) = x$. |
| $\eta(t)$ | Gaussian white noise parameter; stochastic contribution to the allele frequency due to genetic drift, when the dynamics are represented in the form of a stochastic differential equation (i.e., the Langevin equation); defined to have mean $E[\eta(t)] = 0$, variance $\text{Var}[\eta(t)] = E[\eta^2(t)] = 1$, and no temporal correlations (i.e., satisfies a Markov condition $E[\eta(t)\eta(t')] = \delta(t-t')$). |
| $\mathscr{D}p(t)$ | Differential element of the path integral; the product of an infinite number of differential elements $dp$ for the allele frequency $p$ at each time point $t_k$: $\prod_{i=0}^{i=f} dp(t_k)$, where $t_f = t$ is the time of observation. $\mathscr{D}\tilde{p}(t)$ is defined in the same way. See Appendix A1. |
| $\tilde{J}(t)$ | Sink function for $p(t)$; arbitrary time-dependent function defining an absorbing 'sink' (or 'annihilation' operator) for allele frequency; used in the partition functional to generate moments of the distribution for $p(t)$ via functional derivatives (analogous to the the Laplace transform variable for $p(t)$ in a moment generating function). |
| $J(t)$ | Source function for $\tilde{p}(t)$; arbitrary time-dependent function defining an emitting 'source' (or 'creation' operator) for allele frequency; used in the partition functional to generate moments of the distribution for $\tilde{p}(t)$ via functional derivatives. |
| $S[\tilde{p}(t'), p(t'')]$ | The action; exponentiated in the partition functional; defines the probabilistic weights for allele trajectories (paths); dependent on the functional forms of the functions $\tilde{p}(t)$ and $p(t)$. A field theory is fully defined by its action. |

Table S1(continued): *Function, functional, and symbol definitions*

| | |
|---|---|
| $S_0[\tilde{p}(t'), p(t'')]$ | The free action; bilinear (in $\tilde{p}$ and $p$) part of the action used to determine the propagator(s) for a field theory. The free action is usually defined to be fully soluble and is subsequently used extensively in perturbation theory (to approximate propagation between interaction events). Note that the free action for PGFT has a single time derivative (usually indicates diffusion-like or advection-like propagation, as opposed to wave-like propagation). |
| $S_c[\tilde{p}(t'), p(t'')]$ | The coupling action; summarizes all possible interactions in the theory; generally defined as all non-bilinear parts of the action. In PGFT, the coupling action contains terms describing drift and the quadratic effect of natural selection when frequency is high (i.e., the $p^2$ dependence in $-sp(1-p)$ in the diffusion or Langevin equations). The initial condition and mutational influx (derived allele mutation into the frequency bin $1/2N$) have been included in the coupling action for intuitive clarity, but can be incorporated into the free action by redefining the allele frequency, if desired. |
| $G(t, t')$ | Propagator of the 'free' allele frequency (i.e., from the free action) between an earlier time $t'$ and a later time $t$; found by solving the Green's function equation for the operator defined in the free action (i.e., the propagator is the inverse of the linear operator acting on $p(t)$ in the free action $S_0$). PGFT uses the following free propagator: $G(t, t') \equiv \Theta(t - t')e^{-s'(t-t')}$. Note that the decay rate of the exponential in this propagator defines $s'$, the rate of frequency loss due to selection, back mutation, and saturation of the derived allele target size (see Population genetic parameters table). |
| $Z[J(t), \tilde{J}(t)]$ | Partition functional; path integral analog of the moment generating function. Time-dependent moments of the form $E[p^n(t)\tilde{p}^m(t)]$ can be computed by applying functional derivatives to $Z$ and taking $J(t), \tilde{J}(t) \to 0$ at all time points: $E[\tilde{p}^n(t)p^m(t)] = \frac{\delta^n}{\delta J(t)^n}\frac{\delta^m}{\delta \tilde{J}(t)^m} Z[J, \tilde{J}]\|_{J,\tilde{J}\to 0}$. Setting the arguments to $J(t')$ and $\tilde{J}(t)$ yields properties of the relationship between time points (e.g., the correlation function $E[\tilde{p}(t')p(t)]$ between the distributions at times $t'$ and $t$). |
| $E[p^n(t)]$<br><br>$\overline{p}(t) \equiv E[p(t)]$ | Time-dependent non-central moments of the allele frequency probability distribution. The first moment ($n = 1$) defines the time-dependence of the mean frequency and is denoted as $\overline{p}(t)$ for convenience; $k = 2$ defines the time-dependent homozygosity; higher moments describe the asymmetry and tails. |
| $\kappa_n(t)$ | Time-dependent cumulants of the allele frequency probability distribution; first cumulant is equivalent to $\overline{p}(t)$; second, third, and fourth cumulants represent variance, non-standardized skew, and non-standardized excess kurtosis. Cumulants are the primary object of interest in the theory, as they can be computed as the sum of connected Feynman diagrams (disconnected diagrams are related to lower cumulants). |
| $K_n(t)$ | Standardized time-dependent cumulants of the allele frequency probability distribution; the skew $K_3 = \frac{\kappa_3}{\kappa_2^3/2}$ and excess kurtosis $K_4 = \frac{\kappa_4}{\kappa_2^2}$ are computed explicitly and compared to simulations. |
| $\xi \equiv \frac{2K_4}{3K_3^2}$ | Equilibrium parameter approximately equal to one when selection is sufficiently strong ($2Ns' \gtrsim 5$); equivalently expressed as $\xi = \frac{2\kappa_4\kappa_2}{3\kappa_3^2}$. Under mutation-selection-drift balance, this ratio of cumulants becomes roughly independent of the parameters $s$, $\mu$, and $N = const.$; this is a property of the Gamma distribution for allele frequencies under strong selection described by Nei (1968) [1]. After correcting for sampling, estimation of this parameter may provide a test for consistency with mutation-selection-drift balance in a population of constant size. |

Table S2: *Population parameters definitions*

| | |
|---|---|
| $N(t)$ | Population size in number of diploid individuals ($2N(t)$ haploids); treated throughout as a deterministic function with a parametric definition specified for each demography; see Supplemental table ST3. For constant population size, $N \equiv N(t) = const.$ is used for notational convenience. |
| $\mu$ | Mutation rate from ancestral to derived allele. |
| $\mu_b$ | Back mutation rate from derived to ancestral allele. |
| $L$ | Target size for derived mutations; when not explicitly shown, assumed to represent a single locus (i.e., taken to be $L = 1$). |
| $s$ | Selection coefficient; defined as a positive quantity ($s \geq 0$) such that the negative sign is made explicit when selection is deleterious (i.e., purifying selection is represented as $-s$ and beneficial selection is represented as $s$ assuming $s > 0$). |
| $s_{\text{hom}}$ $s_{\text{het}}$ | Homozygous and heterozygous selection coefficients, respectively. Additive natural selection ($s_{\text{hom}} = 2s_{\text{het}}$) is assumed throughout the manuscript, but input for `neqPopDynx` allows for specification of a general diploid selection coefficients; see Materials and Methods for description of neqPopDynx and Data availability for a link to download and input parameter instructions. |
| $s' \equiv (s + \mu + \mu_b)$ | Effective selection coefficient defined by the decay rate of the allele frequency of the propagator $G(t_2, t_1) \propto e^{-s(t_2 - t_1)}$ for the free action. Note, also referred to as the 'mass' (e.g., in particle physics). Accounts for all effects that deterministically reduce allele frequency: purifying selection, back mutation, and saturation of the derived mutational target size. |
| $\theta \equiv 4N(t)\mu$ | Population scaled mutation rate. Not to be confused with the Heaviside function $\Theta(t)$. |
| $\gamma \equiv 2N(t)s$ | Population scaled selection coefficient (dependent on $s$). Low mutation rate limit of the population scaled effective selection coefficient $\gamma'$. Not to be confused with the growth rate $\Gamma$. |
| $\gamma' \equiv 2N(t)s'$ | Population scaled effective selection coefficient (dependent on $s'$). Generalization of the population scaled selection coefficient $\gamma$ such that mutation and back mutation rates may be appreciable. Not to be confused with the growth rate $\Gamma$. |
| $\lambda \equiv \frac{\mu}{s}$ | Mutation-selection balance parameter with negligible back mutation rate ($\mu_b \ll \mu$) and without recurrent mutations ($\theta \ll 1$). Low mutation rate limit of the effective mutation-selection balance parameter $\lambda'$ when $\mu \ll s$. |
| $\lambda' \equiv \frac{\mu}{s'}$ | Effective mutation-selection balance parameter for general mutation and back mutation rates. Generalization of the mutation-selection balance parameter $\lambda$ such that mutation and/or back mutation rates may be appreciable. Note that this parameter is bounded by $\lambda' \leq 1$. |

Table S3: *Demography-specific parameter definitions*

| | |
|---|---|
| $N_0 \equiv N(t_0)$ | Initial population size in number of diploid individuals; used in parameterizations of exponential and bottleneck demographies. |
| $N_{\text{exp}}(t) \equiv N_0 e^{\Gamma t}$ | Time-dependent diploid population size for exponential demography; treated as a deterministic function (i.e., independent of the state of other population parameters. |
| $\Gamma$ | Population growth rate for exponential demography. |
| ${\gamma'}_{\text{exp}} \equiv 2N_0(s' - \Gamma)$ | Modified population-scaled selection coefficient; emergent parameter appearing in moments in the exponential demography. |
| $N_{\text{BN}}(t)$ | Time-dependent diploid population size for bottleneck demography; treated as a deterministic function. See Equation S5 for definition used for computation; equivalent to simpler definition provided in the main text. |
| $N_b$ | Number of diploid individuals during bottleneck. |
| $T_i$ | Time when population is reduced from initial population size $N_0$ to bottleneck population size $N_b$. |
| $T_b$ | Duration of bottleneck; re-expansion from bottleneck to original population size occurs at time $t = T_i + T_b$ |
| $N^-_{\text{bn}} \equiv (N_0 - N_b)$<br>$N^+_{\text{bn}} \equiv (N_0 + N_b)$ | Linear combinations of $N_0$ and $N_b$ defined to reduce the algebraic complexity of bottleneck results. |
| $\Theta_i \equiv \Theta(t - T_i)$<br>$\Theta_{i+b} \equiv \Theta\left(t - (T_i + T_b)\right)$ | Heaviside functions centered after the bottleneck *begins* and after the bottleneck *ends*, respectively; terms proportional to $(\Theta_i - \Theta_{i+b})$ are only nonzero *during* the bottleneck; defined to reduce algebraic complexity of bottleneck results. |
| ${\gamma'}_0 \equiv 2N_0 s'$ | Modified population-scaled effective selection coefficient for the original population size $N_0$. |
| ${\gamma'}_b \equiv 2N_b s'$ | Modified population-scaled effective selection coefficient for the bottleneck population size $N_b$. |
| $N_{\text{osc}}(t)$ | Time-dependent diploid population size for cyclical/oscillatory demography; treated as a deterministic function. See definition in Equation S6. |
| $\tau$ | Period of oscillation period (i.e., time to complete one cycle). |
| $\varphi \equiv 2\pi t/\tau$ | Angular parameterization of oscillation; this enforces correct behavior of cosine parameterization such that the population size is the same when $t = n\tau$ for any integer multiple $n$; defined to reduce algebraic complexity of oscillatory results. |
| $N_{\text{max}}$<br>$N_{\text{min}}$ | Maximum and minimum diploid population size, respectively. Note that the definition for $N_{\text{osc}}(t)$ implies $N_{\text{osc}}(t_0) = N_{\text{max}}$ after setting $t_0 = 0$ for convenience (due to the cosine, rather than sine); this is simply an initialization condition and is not meant to hold biological significance. |
| $N^-_{\text{bn}} \equiv (N_{\text{max}} - N_{\text{min}})$<br>$N^+_{\text{bn}} \equiv (N_{\text{max}} + N_{\text{min}})$ | Linear combinations of $N_{\text{max}}$ and $N_{\text{min}}$ defined to reduce algebraic complexity of oscillatory results. |

#### S2 A brief note on evaluating functional derivatives

Readers unfamiliar with functional calculus may want to consult Srednicki (2007) [2] for a more comprehensive discussion and context for the application of functional calculus to field theory. Though functional calculus may be less familiar, the rules are straightforward and analogous to multivariable calculus. In the present context, the application is the same as in [2]: functional derivatives are applied to the partition functional, a path integral defining the probability distribution of all possible allele trajectories, to evaluate moments of the allele frequency probability distribution . The partition functional is analogous to a moment generating function, but instead of applying partial derivatives to the density to evaluate scalar valued moments, functional derivatives are used to evaluate moments in the form of time-dependent functions (e.g., to find the time dependence of the mean frequency). In the manuscript text, it is noted that such moments can be computed from the partition functional $Z[\tilde{J}(t_a), J(t_b)]$ as follows.

$$E[\tilde{p}^n(t)p^m(t)] = \left[\frac{\delta^n}{\delta J^n(t)}\frac{\delta^m}{\delta J^m(t)} Z[\tilde{J}(t_a), J(t_b)]\right]\bigg|_{\tilde{J},J\to 0} \tag{S1}$$

Here, the functional derivative $\frac{\delta}{\delta\tilde{J}(t)}$ is defined by the following application to $\tilde{J}(t)$.

$$\frac{\delta}{\delta\tilde{J}(t)}\tilde{J}(t_a) = \delta(t - t_a) \tag{S2}$$

Applying the functional derivative to an arbitrary test function $f(\tilde{J}(t_a), t_a)$ uses the functional equivalent of the chain rule.

$$\begin{aligned}\frac{\delta}{\delta\tilde{J}(t)} f(\tilde{J}(t_a), t_a) &= \left(\frac{\partial}{\partial\tilde{J}} f(\tilde{J}, t)\right)\left(\frac{\delta}{\delta\tilde{J}(t)}\tilde{J}(t_a)\right)\\ &= f'(\tilde{J}(t), t)\delta(t - t_a)\end{aligned} \tag{S3}$$

In the second line, the partial derivative treats the function $\tilde{J}$ as if it is a variable rather than a function such that $\frac{\partial}{\partial_{\tilde{J}}} f(\tilde{J}) = f'(\tilde{J})$, which is followed by applying Equation S2. Functional derivatives of $\frac{\delta}{\delta J(t)}$ are defined in the same way. Additionally, functional derivatives commute with both integrals and path integrals. Since the partition functional is a functional integral, the functional derivatives are applied to the exponentiated action directly (i.e., the test function $f$ is the function $e^{-S[\tilde{p},p]}$).

After computing the derivatives, a limit must be taken to remove the source functions: $\tilde{J}(t) \to 0$ and $J(t) \to 0$ Formally, this limit is taken by defining the source functions $\tilde{J}(t) = \varepsilon\tilde{j}(t)$ and $J(t) = \varepsilon j(t)$, where $\varepsilon$ is a scalar valued real number, and taking the limit $\varepsilon \to 0$.

#### S3 Feynman diagrams and corresponding integrals

Here, integrals for all diagrams presented in the accompanying manuscript (repeated here for clarity) are computed using the appropriate Feynman rules. For an accessible overview relevant to the present context, please consult Chow and Buice (2015) [3]; further discussion of the mathematical underpinnings of Feynman diagrams can be found in [2] and [4]. For each integral, the relevant allele frequency propagators $G(t,t') = \Theta(t-t')e^{-s'(t-t')}$ are shown and expressed in terms of the Heaviside function $\Theta(x)$. Results are presented for all relevant expressions that appear in the manuscript text for the four demographies discussed: constant population size, exponential population growth, a population that experiences an instantaneous bottleneck and subsequent re-expansion, and a cyclical population with oscillating size. Note that all expressions below are also presented in Supplemental File SF1 for easier use in any future applications. For simplicity, the integrals and corresponding results below are written using several definitions to shorten notation. First, all results are expressed in terms of the following, using the same notation as in the manuscript text.

$$s' \equiv (s+\mu+\mu_b)$$
$$\lambda' \equiv \frac{\mu}{s'}$$

For the population of constant size, specifically, the following notation is used.

$$\gamma' \equiv 2Ns'$$

In expressions for the exponential demography, the following definitions are used when appropriate.

$$N_{\text{exp}}(t) = N_0 e^{\Gamma t} \tag{S4}$$

$$\gamma'_{\text{exp}} \equiv 2N_0(s'-\Gamma)$$

For the bottleneck demography, the integrals are expressed using the following notation.

$$N_{\text{BN}}(t) = \frac{N_0}{1+\left(\frac{N_0}{N_b}-1\right)\left[\Theta(t-T_i)-\Theta(t-(T_i+T_b))\right]} \tag{S5}$$

$$\gamma'_0 \equiv 2N_0 s'$$
$$\gamma'_b \equiv 2N_b s'$$
$$\Theta_{i+b} \equiv \Theta(t-T_i)$$
$$\Theta_{i+b} \equiv \Theta\left(t-(T_i+T_b)\right)$$
$$N^-_{\text{bn}} \equiv (N_0-N_b)$$
$$N^+_{\text{bn}} \equiv (N_0+N_b)$$

Integrals for the cyclical demography with oscillating size are expressed in terms of the following variables.

$$N_{\text{osc}}(t) = \frac{2N_{\max}N_{\min}}{N^+_{\text{osc}}-N^-_{\text{osc}}\cos[\varphi]} \tag{S6}$$

$$\varphi \equiv \frac{2\pi t}{\tau}$$
$$N^-_{\text{osc}} \equiv (N_{\max}-N_{\min})$$
$$N^+_{\text{osc}} \equiv (N_{\max}+N_{\min})$$

Additionally, the initial time will be set to $t_0 = 0$ in all cases such that the generation time $t$ is a strictly positive quantity. Thus, all results should be considered as being implicitly multiplied by an overall factor of $\Theta(t)$.

#### S3.1 Diagrams for the mean frequency $\overline{p}(t)$

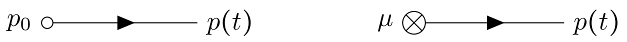


(D1)

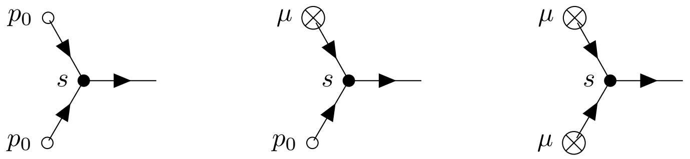


(D2)

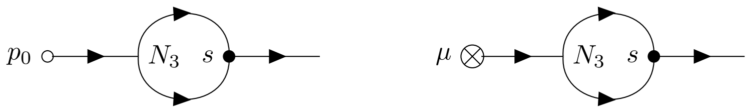


(D3)

Diagrams D2 and D3 are higher order corrections to the mean frequency $\overline{p}(t)$, as discussed in the manuscript. Diagram D3 represents the first (i.e., lowest-vertex order) contributions to the mean frequency that are dependent on drift (i.e., on the $N_3$ vertex). Thus, these diagrams become demography-dependent, and must be evaluated separately for the exponential, bottleneck, and cyclical populations. However, the expressions below make clear that the corresponding contributions to the mean frequency are small; these expressions are suppressed by $1/\gamma'$, or the analogous variable defined for each demography (e.g., ${\gamma'}_{\text{exp}}$, ${\gamma'}_0$, ${\gamma'}_{\text{b}}$). Note that the results for nontrivial demography are not presented in the manuscript text, but appear here and in Supplemental File SF1.

##### S3.1.1 Diagram 1 (left)

**Any demography:**

$$\begin{aligned}\overline{p}(t)_{D1L} &= p_0 \int_0^t dt_1\ \delta(t_1 - 0)\ G(t, t_1) \\ &= p_0 e^{-s't}\Theta(t) \\ &= p_0 e^{-s't}\end{aligned} \tag{S7}$$

##### S3.1.2 Diagram 1 (right)

**Any demography:**

$$\begin{aligned}\overline{p}_{D1R}(t) &= \mu \int_0^t dt_1\ G(t, t_1) \\ &= \mu \int_0^t dt_1\ \Theta(t - t_1) e^{-s'(t-t_1} \\ &= \frac{\mu}{s'}\left(1 - e^{-s't}\right)\end{aligned} \tag{S8}$$

###### S3.1.3 Diagram 2 (left)

**Any demography:**

$$\begin{aligned}\overline{p}_{D2L} &= sp_0^2\ \frac{2}{2!}\ \int_0^t dt_1\ G^2(t_1,0)G(t,t_1)\\ &= sp_0^2\ \int_0^t \Theta^2(t_1)\Theta(t-t_1)\ e^{-s'(t+t_1)}\\ &= \frac{sp_0^2}{s'}\left(e^{-s't}\ - e^{-2s't}\right)\end{aligned} \tag{S9}$$

###### S3.1.4 Diagram 2 (center)

**Any demography:**

$$\begin{aligned}\overline{p}_{D2C} &= s\mu p_0\ 2\ \int_0^t dt_1 dt_2\ G(t_2,0)G(t_2,t_1)G(t,t_2)\\ &= 2s\mu p_0\ \int_0^t dt_1 dt_2\ \Theta(t_2)\Theta(t_2-t_1)\Theta(t-t_2)\ e^{-s'(t+t_2-t_1)}\\ &= \frac{2s\lambda' p_0}{s'}\left((s't-1)e^{-s't}+e^{-2s't}\right)\end{aligned} \tag{S10}$$

###### S3.1.5 Diagram 2 (right)

**Any demography:**

$$\begin{aligned}\overline{p}_{D2R} &= s\mu^2\ \frac{2}{2!}\ \int_0^t dt_1 dt_2 dt_3\ G(t_3,t_1)G(t_3,t_2)G(t,t_3)\\ &= s\mu^2\ \frac{2}{2!}\ \int_0^t dt_1 dt_2 dt_3\ \Theta(t_3-t_1)\Theta(t_3-t_2)\Theta(t-t_3)\ e^{-s'(t+t_3-t_2-t_1)}\\ &= \frac{s\lambda'^2}{s'}\left(1-2s't\ e^{-s't}-e^{-2s't}\right)\end{aligned} \tag{S11}$$

###### S3.1.6 Diagram 3 (left)

**Constant size:**

$$\begin{aligned}\overline{p}_{D3L}(t) &= \frac{sp_0}{4N}\ 2\ \int_0^t dt_1 dt_2\ G(t_1,0)G^2(t_2,t_1)G(t,t_2)\\ &= \frac{sp_0}{2N}\ \int_0^t dt_1 dt_2 \Theta(t_1)\Theta^2(t2-t1)\Theta(t-t_2)\ e^{-s'(t+t_2-t_1)}\\ &= \frac{sp_0}{\gamma' s'}\left((s't-1)e^{-s't}+e^{-2s't}\right)\end{aligned} \tag{S12}$$

**Exponential growth:**

$$\begin{aligned}\overline{p}_{D3L}(t) &= sp_0\ 2\ \int_0^t dt_1 dt_2\ \frac{1}{4N_{\exp}(t_1)} G(t_1,0)G^2(t_2,t_1)G(t,t_2)\\ &= sp_0\ \int_0^t dt_1 dt_2 \Theta(t_1)\Theta^2(t2-t1)\Theta(t-t_2)\ \frac{e^{-s'(t+t_2-t_1)}}{2N_{\exp}(t_1)}\\ &= \frac{p_0 s}{\gamma'_{\exp}}\left(\frac{(s'-\Gamma)e^{-s't}}{\Gamma s'}-\frac{e^{-(s'+\Gamma)t}}{\Gamma}+\frac{e^{-2s't}}{s'}\right)\end{aligned} \tag{S13}$$

**Bottleneck:**

$$
\begin{aligned}
\overline{p}_{D3L}(t) &= sp_0\ 2\int_0^t dt_1 dt_2\ \frac{1}{4N_{\mathrm{BN}}(t_1)} G(t_1,0)G^2(t_2,t_1)G(t,t_2) \\
&= sp_0 \int_0^t dt_1 dt_2 \Theta(t_1)\Theta^2(t2-t1)\Theta(t-t_2)\ \frac{e^{-s'(t+t_2-t_1)}}{2N_{\mathrm{BN}}(t_1)} \\
&= \frac{p_0 s}{2N_0 N_b s'^2}\Bigg[N_b e^{-s't}\left((s't-1)+e^{-s't}\right) \\
&\quad + N_{\mathrm{bn}}^{-}\Bigg(\Theta_i e^{-s't}\left((s't-s'T_i-1)+e^{-s'(t-T_i)}\right) - \Theta_{i+b}e^{-s't}\left((s'(t-(T_i+T_b))-1)+e^{-s'(t-(T_i+T_b))}\right)\Bigg)\Bigg] \qquad \text{(S14)}
\end{aligned}
$$

**Oscillating size:**

$$
\begin{aligned}
\overline{p}_{D3L}(t) &= sp_0\ 2\int_0^t dt_1 dt_2\ \frac{1}{4N_{\mathrm{osc}}(t_1)} G(t_1,0)G^2(t_2,t_1)G(t,t_2) \\
&= sp_0 \int_0^t dt_1 dt_2 \Theta(t_1)\Theta^2(t2-t1)\Theta(t-t_2)\ \frac{e^{-s'(t+t_2-t_1)}}{2N_{\mathrm{osc}}(t_1)} \\
&= \frac{p_0 s}{8\pi N_{\max} N_{\min} s'^2\left(s'^2\tau^2+4\pi^2\right)} \quad \times \\
&\Bigg[e^{-s't}\Bigg(2\pi N_{\mathrm{osc}}^{+}(s't-1)\left(s'^2\tau^2+4\pi^2\right)+s'^2\tau^2 N_{\mathrm{osc}}^{-}\left(2\pi\cos[\varphi]-s'\tau\sin[\varphi]\right)\Bigg)+4\pi e^{-2s't}\left(2\pi^2 N_{\mathrm{osc}}^{+}+N_{\min}s'^2\tau^2\right)\Bigg] \qquad \text{(S15)}
\end{aligned}
$$

Note that the above expression is proportional to $1/4N_{\max}N_{\min}s'$, which is part of the inverse of the population-scaled effective selection coefficient (i.e., the effective drift coefficient $1/2\overline{N}_{\mathrm{osc}}$ multiplied by $1/s'$) averaged over one cycle, $\overline{\gamma}'_{\mathrm{osc}} = 2\overline{N}_{\mathrm{osc}}s' = 4N_{\max}N_{\min}s'/N_{\mathrm{osc}}^{+}$. I have chosen not to express this result in terms of $\overline{\gamma}'_{\mathrm{osc}}$, as this can also be expressed in terms of the inverse of the scaled selection coefficient for the initial (or maximum) size $\gamma'_{\max} = 2N_{\max}s'$, if preferred. Similarly, all results for a cyclical population in this section are left in terms of $1/4N_{\max}N_{\min}s'$ for generality.

###### S3.1.7 Diagram 3 (right)

**Constant size:**

$$
\begin{aligned}
\overline{p}_{D3R}(t) &= \frac{s\mu}{4N}\ 2\int_0^t dt_1 dt_2 dt_3\ G(t_2,t_1)G^2(t_3,t_2)G(t,t_3) \\
&= \frac{s\mu}{2N}\int_0^t dt_1 dt_2 dt_3\ \Theta(t_2-t_1)\Theta^2(t_3-t_2)\Theta(t-t_3)\ e^{-s'(t+t_3-t_2-t_1)} \\
&= \frac{s\lambda'}{2\gamma' s'}\left(1-2st\ e^{-s't}-e^{-2s't}\right) \qquad \text{(S16)}
\end{aligned}
$$

**Exponential growth:**

$$\begin{aligned}\overline{p}_{D3R}(t) &= s\mu\ 2\int_0^t dt_1 dt_2 dt_3\ \frac{1}{4N_{\rm exp}(t_2)} G(t_2,t_1)G^2(t_3,t_2)G(t,t_3)\\ &= s\mu\int_0^t dt_1 dt_2 dt_3\ \Theta(t_2-t_1)\Theta^2(t_3-t_2)\Theta(t-t_3)\frac{e^{-s'(t+t_3-t_2-t_1)}}{2N_{\rm exp}(t_2)}\\ &= \frac{\lambda' s}{\gamma'_{\rm exp}}\left(\frac{\left(e^{-\Gamma t}-e^{-2s't}\right)}{2s'-\Gamma}-\frac{e^{-s't}\left(1-e^{-\Gamma t}\right)}{\Gamma}\right)\end{aligned} \tag{S17}$$

**Bottleneck:**

$$\begin{aligned}&\overline{p}_{D3R}(t) = s\mu\ 2\int_0^t dt_1 dt_2 dt_3\ \frac{1}{4N_{\rm BN}(t_2)} G(t_2,t_1)G^2(t_3,t_2)G(t,t_3)\\ &= s\mu\int_0^t dt_1 dt_2 dt_3\ \Theta(t_2-t_1)\Theta^2(t_3-t_2)\Theta(t-t_3)\frac{e^{-s'(t+t_3-t_2-t_1)}}{2N_{\rm BN}(t_2)}\\ &= \frac{\mu s}{4N_0 N_b s'^3}\Bigg[N_b\left(1-2s'te^{-s't}-e^{-2s't}\right)+N_{\rm bn}^-\Theta_i\Bigg(1+e^{-s't}\Big[2\left(1-s'(t-T_i)\right)-e^{s'T_i}\left(2+2e^{-s't}-e^{-s'(t-T_i)}\right)\Big]\Bigg)\\ &\qquad -N_{\rm bn}^-\Theta_{i+b}\Bigg(1+e^{-s't}\Big[2\left(1-s'\left[t-(T_i+T_b)\right]\right)-e^{s'(T_i+T_b)}\left(2+2e^{-s't}-e^{-s'(t-(T_i+T_b))}\right)\Big]\Bigg)\Bigg]\end{aligned} \tag{S18}$$

**Oscillating size:**

$$\begin{aligned}&\overline{p}_{D3R}(t) = s\mu\ 2\int_0^t dt_1 dt_2 dt_3\ \frac{1}{4N_{\rm osc}(t_2)} G(t_2,t_1)G^2(t_3,t_2)G(t,t_3)\\ &\quad = s\mu\int_0^t dt_1 dt_2 dt_3\ \Theta(t_2-t_1)\Theta^2(t_3-t_2)\Theta(t-t_3)\frac{e^{-s'(t+t_3-t_2-t_1)}}{2N_{\rm osc}(t_2)}\\ &\quad = \frac{\lambda' s}{8N_{\rm max}N_{\rm min}s'^2\left(\pi s'^4\tau^4+5\pi^3 s'^2\tau^2+4\pi^5\right)} \quad \times\\ &\Bigg[s'^2\tau^2 N_{\rm osc}^-\Bigg(\left[s'^2\tau^2 e^{-s't}-\pi^2\left(3-e^{-s't}\right)\right]\left(s'\tau\sin[\varphi]\right)+\left[2\pi^2\left(1-e^{-s't}\right)-s'^2\tau^2\left(1+2e^{-s't}\right)\right]\left(\pi\cos[\varphi]\right)\Bigg)\\ &\qquad +\pi N_{\rm osc}^+\left(s'^4\tau^4+5\pi^2 s'^2\tau^2+4\pi^4\right)-2\pi e^{-s't}\left(s'^2\tau^2+\pi^2\right)\left(s'^2\tau^2(N_{\rm osc}^+ s't-N_{\rm osc}^-)+4\pi^2 s'tN_{\rm osc}^+\right)\\ &\qquad -\pi e^{-2s't}\left(4\pi^4 N_{\rm osc}^+ + 2N_{\rm min}s'^4\tau^4+\pi^2 s'^2\tau^2(4N_{\rm max}+3N_{\rm osc}^+)\right)\Bigg]\end{aligned} \tag{S19}$$

#### S3.2 Diagrams for the second cumulant $\kappa_2(t)$

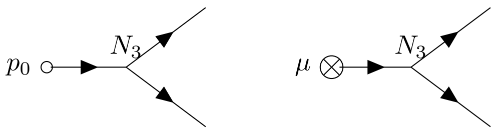


(D4)

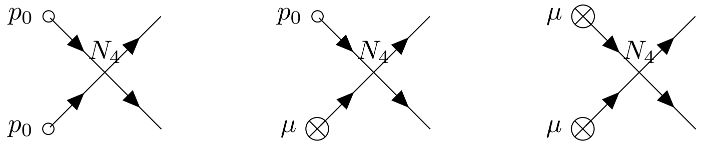


(D5)

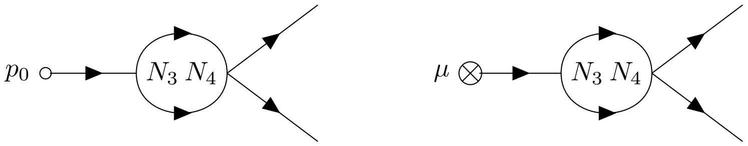


(D6)

As is the case for the first (lowest-vertex order) corrections (D2 and D3) to the one-vertex approximation for the mean frequency (given by the deterministic equation), Diagrams D4 and D5 represent the first corrections to the variance of the allele frequency probability distribution. These diagrams are only computed for a population of constant size, but are computed for the three specified demographies here and in Supplemental File SF1. Unlike D2 for the mean frequency, all of the following diagrams represent drift-dependent contributions, as drift is inextricably incorporated into the variance of the frequency distribution, and thus the homozygosity.

##### S3.2.1 Diagram 4 (left)

**Constant size:**

$$\begin{aligned}\kappa_{2D4L} &= \frac{p_0}{4N}\ 2\int_0^t dt_1\ G(t_1,0)G^2(t,t_1)\\ &= \frac{p_0}{2N}\int_0^t dt_1\ \Theta(t_1)\Theta^2(t-t_1)\ e^{-s'(2t-t_1)}\\ &= \frac{p_0}{\gamma'}e^{-s't}\left(1-e^{-s't}\right)\end{aligned}\tag{S20}$$

**Exponential growth:**

$$\begin{aligned}\kappa_{2D4L} &= p_0\ 2\int_0^t dt_1\ \frac{1}{4N_{\text{exp}}(t_1)}G(t_1,0)G^2(t,t_1)\\ &= p_0\int_0^t dt_1\ \Theta(t_1)\Theta^2(t-t_1)\frac{e^{-s'(2t-t_1)}}{2N_{\text{exp}}(t_1)}\\ &= \frac{p_0}{\gamma'_{\text{exp}}}e^{-s't}\left(e^{-\Gamma t}-e^{-s't}\right)\end{aligned}\tag{S21}$$

**Bottleneck:**

$$\begin{aligned}\kappa_{2D4L} &= p_0\ 2\int_0^t dt_1\ \frac{1}{4N_{\text{BN}}(t_1)}G(t_1,0)G^2(t,t_1)\\ &= p_0\int_0^t dt_1\ \Theta(t_1)\Theta^2(t-t_1)\frac{e^{-s'(2t-t_1)}}{2N_{\text{BN}}(t_1)}\\ &= \frac{p_0}{\gamma'_0}e^{-s't}\left(1-e^{-s't}\right)+\left(\frac{p_0}{\gamma'_b}-\frac{p_0}{\gamma'_0}\right)e^{-s't}\left[\Theta_i\left(1-e^{-s't+s'T_i}\right)-\Theta_{i+b}\left(1-e^{-s't+s'(T_i+T_b)}\right)\right]\end{aligned}\tag{S22}$$

**Oscillating size:**

$$
\begin{aligned}
\kappa_{2D4L} &= p_0 \; 2 \int_0^t dt_1 \; \frac{1}{4N_{\mathrm{BN}}(t_1)} G(t_1,0) G^2(t,t_1) \\
&= p_0 \int_0^t dt_1 \; \Theta(t_1)\Theta^2(t-t_1) \frac{e^{-s'(2t-t_1)}}{2N_{\mathrm{BN}}(t_1)} \\
&= \frac{p_0 \left(4\pi^2 N_{\mathrm{osc}}^+ e^{-s't}\left(1-e^{-s't}\right) + s'^2\tau^2\left(N_{\mathrm{osc}}^+ - 2N_{\mathrm{min}}e^{-s't}\right)e^{-s't} - N_{\mathrm{osc}}^- s'\tau\left(s'\tau\cos[\varphi] + 2\pi\sin[\varphi]\right)e^{-s't}\right)}{4N_{\mathrm{max}}N_{\mathrm{min}}s'\left(4\pi^2 + s'^2\tau^2\right)}
\end{aligned}
\tag{S23}
$$

##### S3.2.2 Diagram 4 (right)

**Constant size:**

$$
\begin{aligned}
\kappa_{2D4R} &= \frac{\mu}{4N} \; 2 \int_0^t dt_1 dt_2 \; G(t_2,t_1) G^2(t,t_2) \\
&= \frac{\mu}{4N} \int_0^t dt_1 dt_2 \; \Theta(t_2-t_1)\Theta^2(t-t_2) \; e^{-s'(2t-t_2-t_1)} \\
&= \frac{\lambda'}{2\gamma'}\left(1-e^{-s't}\right)^2
\end{aligned}
\tag{S24}
$$

**Exponential growth:**

$$
\begin{aligned}
\kappa_{2D4R} &= \mu \; 2 \int_0^t dt_1 dt_2 \; \frac{1}{4N_{\mathrm{exp}}(t_2)} G(t_2,t_1) G^2(t,t_2) \\
&= \mu \int_0^t dt_1 dt_2 \; \Theta(t_2-t_1)\Theta^2(t-t_2) \frac{e^{-s'(2t-t_2-t_1)}}{2N_{\mathrm{exp}}(t_2)} \\
&= \frac{\lambda'}{\gamma'_{\mathrm{exp}}(2s'-\Gamma)}\left(s'e^{-2s't} + (s'-\Gamma)e^{-\Gamma t} - (2s'-\Gamma)e^{-(\Gamma+s')t}\right)
\end{aligned}
\tag{S25}
$$

**Bottleneck:**

$$
\begin{aligned}
\kappa_{2D4R} &= \mu \; 2 \int_0^t dt_1 dt_2 \; \frac{1}{4N_{\mathrm{BN}}(t_2)} G(t_2,t_1) G^2(t,t_2) \\
&= \mu \int_0^t dt_1 dt_2 \; \Theta(t_2-t_1)\Theta^2(t-t_2) \frac{e^{-s'(2t-t_2-t_1)}}{2N_{\mathrm{BN}}(t_2)} \\
&= \frac{\lambda'}{2\gamma'_0}\left(1-e^{-s't}\right)^2 + \left(\frac{\lambda'}{2\gamma'_b} - \frac{\lambda'}{2\gamma'_0}\right)\Big[\Theta_i\left(1+e^{-s'(t-T_i)} - 2e^{-s't}\right)\left(1-e^{-s'(t-T_i)}\right) \\
&\qquad - \Theta_{i+b}\left(1+e^{-s'(t-T_i-T_b)} - 2e^{-s't}\right)\left(1-e^{-s'(t-T_i-T_b)}\right)\Big]
\end{aligned}
\tag{S26}
$$

**Oscillating size:**

$$\kappa_{2D4R} = \mu \; 2 \int_0^t dt_1 dt_2 \; \frac{1}{4N_{\text{osc}}(t_2)} G(t_2,t_1) G^2(t,t_2)$$
$$= \mu \int_0^t dt_1 dt_2 \; \Theta(t_2 - t_1)\Theta^2(t-t_2) \frac{e^{-s'(2t-t_2-t_1)}}{2N_{\text{osc}}(t_2)}$$
$$= \frac{\lambda'}{8N_{\max}N_{\min}s'\left(4\pi^4 + 5\pi^2 s'^2 \tau^2 + s'^4\tau^4\right)} \quad \times$$
$$\Bigg[N^+_{\text{osc}}\Bigg(\left(4\pi^4 + 3\pi^2 s'^2\tau^2\right)\left(1-e^{-s't}\right)^2 + \left(2\pi^2 s'^2 \tau^2 + s'^4\tau^4\right)\left(1-2e^{-s't}\right)\Bigg) + \left(4\pi^2 N_{\max} s'^2\tau^2 + 2N_{\min}s'^4\tau^4\right)e^{-2s't}$$
$$- N^-_{\text{osc}} s'\tau\left(2s'\tau\left(s'^2\tau^2+\pi^2\right)\cos[\varphi]\; e^{-s't} + \left(3\pi^2 + \left(s'^2\tau^2+\pi^2\right)\left(1-4e^{-s't}\right)\right)\left(\pi\sin[\varphi] + s'\tau\cos[\varphi]\right)\right)\Bigg] \tag{S27}$$

##### S3.2.3 Diagram 5 (left)

**Constant size:**

$$\kappa_{2D5L} = -\frac{p_0^2}{4N}\;\frac{2\times 2}{2!}\int_0^t dt_1\; G^2(t_1,0)G^2(t,t_1)$$
$$= -\frac{p_0^2}{2N}\int_0^t dt_1\; \Theta^2(t_1)\Theta^2(t-t_1)\; e^{-2s't}$$
$$= -\frac{p_0^2}{\gamma'}\;(s't)\; e^{-2s't} \tag{S28}$$

**Exponential growth:**

$$\kappa_{2D5L} = -p_0^2\;\frac{2\times 2}{2!}\int_0^t dt_1\;\frac{1}{4N_{\exp}(t_1)}G^2(t_1,0)G^2(t,t_1)$$
$$= -p_0^2\int_0^t dt_1\;\Theta^2(t_1)\Theta^2(t-t_1)\;\frac{e^{-2s't}}{2N_{\exp}(t_1)}$$
$$= -\frac{p_0^2}{\gamma'_{\exp}}\left(\frac{s'-\Gamma}{\Gamma}\right)e^{-2s't}\left(1-e^{-\Gamma t}\right) \tag{S29}$$

**Bottleneck:**

$$\kappa_{2D5L} = -p_0^2\;\frac{2\times 2}{2!}\int_0^t dt_1\;\frac{1}{4N_{\text{BN}}(t_1)}G^2(t_1,0)G^2(t,t_1)$$
$$= -p_0^2\int_0^t dt_1\;\Theta^2(t_1)\Theta^2(t-t_1)\;\frac{e^{-2s't}}{2N_{\text{BN}}(t_1)}$$
$$= -\frac{p_0^2}{\gamma'_0}\;e^{-2s't}\left((s't) + \left(\frac{N_0}{N_b}-1\right)\left[\Theta_i s'(t-T_i) - \Theta_{i+b}s'\left(t-(T_i+T_b)\right)\right]\right) \tag{S30}$$

**Oscillating size:**

$$
\begin{aligned}
\kappa_{2D5L} &= -p_0^2 \; \frac{2\times 2}{2!} \int_0^t dt_1 \; \frac{1}{4N_{\mathrm{osc}}(t_1)} G^2(t_1,0)G^2(t,t_1) \\
&= -p_0^2 \int_0^t dt_1 \; \Theta^2(t_1)\Theta^2(t-t_1) \; \frac{e^{-2s't}}{2N_{\mathrm{osc}}(t_1)} \\
&= -\frac{p_0^2}{4N_{\mathrm{max}}N_{\mathrm{min}}s'} \; e^{-2s't}\left(N_{\mathrm{osc}}^{+}(s't) - N_{\mathrm{osc}}^{-}\frac{s'\tau}{2\pi}\sin[\varphi]\right)
\end{aligned}
\tag{S31}
$$

##### S3.2.4 Diagram 5 (center)

**Constant size:**

$$
\begin{aligned}
\kappa_{2D5C} &= -\frac{p_0\mu}{4N} \; 2\times 2 \int_0^t dt_1 \; G(t_2,0)G(t_2,t_1)G^2(t,t_2) \\
&= -\frac{2p_0\mu}{2N} \int_0^t dt_1 \; \Theta(t_2)\Theta(t_2-t_1)\Theta^2(t-t_2) \; e^{-s'(2t-t1)} \\
&= -\frac{2p_0\lambda'}{\gamma'}\left(e^{-s't} - (s't+1)e^{-2s't}\right)
\end{aligned}
\tag{S32}
$$

**Exponential growth:**

$$
\begin{aligned}
\kappa_{2D5C} &= -p_0\mu \; 2\times 2 \int_0^t dt_1 \; \frac{1}{4N_{\mathrm{exp}}(t_2)} G(t_2,0)G(t_2,t_1)G^2(t,t_2) \\
&= -2p_0\mu \int_0^t dt_1 \; \Theta(t_2)\Theta(t_2-t_1)\Theta^2(t-t_2)\frac{e^{-s'(2t-t1)}}{2N_{\mathrm{exp}}(t_2)} \\
&= -\frac{2\lambda' p_0}{\gamma'_{\mathrm{exp}}}\left(\frac{s'}{\Gamma}\left(e^{-(\Gamma+2s')t} - e^{-2s't}\right) + \left(e^{-(s'+\Gamma)t} - e^{-(2s'+\Gamma)t}\right)\right)
\end{aligned}
\tag{S33}
$$

**Bottleneck:**

$$
\begin{aligned}
\kappa_{2D5C} &= -p_0\mu \; 2\times 2 \int_0^t dt_1 \; \frac{1}{4N_{\mathrm{BN}}(t_2)} G(t_2,0)G(t_2,t_1)G^2(t,t_2) \\
&= -2p_0\mu \int_0^t dt_1 \; \Theta(t_2)\Theta(t_2-t_1)\Theta^2(t-t_2)\frac{e^{-s'(2t-t1)}}{2N_{\mathrm{BN}}(t_2)} \\
&= -\frac{2\lambda' p_0}{{\gamma'}_0 N_b}\Bigg[N_b\left(e^{-s't} - (s't+1)e^{-2s't}\right) + N_{\mathrm{bn}}^{-}\Theta_i\left(e^{-s't} - e^{-s'(2t-T_i)} + s'(T_i - t)e^{-2s't}\right) \\
&\qquad - N_{\mathrm{bn}}^{-}\Theta_{i+b}\left(e^{-s't} + e^{-s'(2t-(T_i+T_b))} - s'\left(t-(T_i+T_b)\right)e^{-2s't}\right)\Bigg]
\end{aligned}
\tag{S34}
$$

**Oscillating size:**

$$
\begin{aligned}
\kappa_{2D5C} &= -p_0\mu\ 2\times 2\ \int_0^t dt_1\ \frac{1}{4N_{\text{osc}}(t_2)} G(t_2,0)G(t_2,t_1)G^2(t,t_2) \\
&= -2p_0\mu\ \int_0^t dt_1\ \Theta(t_2)\Theta(t_2-t_1)\Theta^2(t-t_2)\frac{e^{-s'(2t-t1)}}{2N_{\text{osc}}(t_2)} \\
&= -\frac{\lambda' p_0}{4\pi N_{\max}N_{\min}s'\left(s'^2\tau^2+4\pi^2\right)} \quad \times \\
&\Bigg[ -s'\tau N_{\text{osc}}^{-}e^{-s't}\Bigg(\left(4\pi^2-\left(4\pi^2+s'^2\tau^2\right)e^{-s't}\right)\sin[\varphi]+2\pi s'\tau\cos[\varphi]\Bigg) \\
&+2\pi N_{\text{osc}}^{+}e^{-s't}\Bigg(\left(s'^2\tau^2+4\pi^2\right)-\left(4\pi^2\left(s't+1\right)+s'^2\tau^2(s't)\right)e^{-s't}\Bigg)-4\pi N_{\min}s'^2\tau^2e^{-2s't}\Bigg] \qquad \text{(S35)}
\end{aligned}
$$

##### S3.2.5 Diagram 5 (right)

**Constant size:**

$$
\begin{aligned}
\kappa_{2D5R} &= -\frac{\mu^2}{4N}\ \frac{2\times 2}{2!}\ \int_0^t dt_1dt_2dt_3\ G(t_3,t_1)G(t_3,t_2)G^2(t,t_3) \\
&= -\frac{\mu^2}{2N}\ \int_0^t dt_1dt_2dt_3\ \Theta(t_3-t_1)\Theta(t_3-t_2)\Theta^2(t-t_3)\ e^{-s'(2t-t_2-t_1)} \\
&= -\frac{\lambda'^2}{2\gamma'}\ \left(1-4e^{-s't}+(2s't+3)e^{-2s't}\right) \qquad \text{(S36)}
\end{aligned}
$$

**Exponential growth:**

$$
\begin{aligned}
\kappa_{2D5R} &= -\mu^2\ \frac{2\times 2}{2!}\ \int_0^t dt_1dt_2dt_3\ \frac{1}{4N_{\exp}(t_3)}G(t_3,t_1)G(t_3,t_2)G^2(t,t_3) \\
&= -\mu^2\ \int_0^t dt_1dt_2dt_3\ \Theta(t_3-t_1)\Theta(t_3-t_2)\Theta^2(t-t_3)\frac{e^{-s'(2t-t_2-t_1)}}{2N_{\exp}(t_3)} \\
&= -\frac{\lambda'^2}{\gamma'_{\exp}}\left(\frac{2s'^2\left(e^{-2s't}-e^{-(\Gamma+2s')t}\right)}{\Gamma(2s'-\Gamma)}-\frac{\Gamma e^{-\Gamma t}\left(1-e^{-s't}\right)^2}{(2s'-\Gamma)}+\frac{s'\left(e^{-\Gamma t}-4e^{-(\Gamma+s')t}+3e^{-(\Gamma+2s')t}\right)}{(2s'-\Gamma)}\right) \qquad \text{(S37)}
\end{aligned}
$$

**Bottleneck:**

$$
\begin{aligned}
\kappa_{2D5R} &= -\mu^2\ \frac{2\times 2}{2!}\ \int_0^t dt_1dt_2dt_3\ \frac{1}{4N_{\text{BN}}(t_3)}G(t_3,t_1)G(t_3,t_2)G^2(t,t_3) \\
&= -\mu^2\ \int_0^t dt_1dt_2dt_3\ \Theta(t_3-t_1)\Theta(t_3-t_2)\Theta^2(t-t_3)\frac{e^{-s'(2t-t_2-t_1)}}{2N_{\text{BN}}(t_3)} \\
&= -\frac{\lambda'^2}{2\gamma'_0N_b}\Bigg[N_b\left(1-4e^{-s't}+(2s't+3)e^{-2s't}\right)+N_{\text{bn}}^{-}\Theta_i\left(1-4e^{-s't}+2s'(t-T_i)e^{-2s't}-e^{-2s'(t-T_i)}+4e^{-s'(2t-T_i)}\right) \\
&\quad -N_{\text{bn}}^{-}\Theta_{i+b}\left(1-4e^{-s't}+2s'(t-T_i-T_b)e^{-2s't}-e^{-2s'(t-(T_i+T_b))}+4e^{-s'(2t-(T_i+T_b))}\right)\Bigg] \qquad \text{(S38)}
\end{aligned}
$$

**Oscillating size:**

$$
\begin{aligned}
\kappa_{2D5R} &= -\mu^2 \ \frac{2\times 2}{2!} \ \int_0^t dt_1 dt_2 dt_3 \ \frac{1}{4N_{\text{osc}}(t_3)} G(t_3,t_1)G(t_3,t_2)G^2(t,t_3) \\
&= -\mu^2 \ \int_0^t dt_1 dt_2 dt_3 \ \Theta(t_3-t_1)\Theta(t_3-t_2)\Theta^2(t-t_3)\frac{e^{-s'(2t-t_2-t_1)}}{2N_{\text{osc}}(t_3)} \\
&= -\frac{\lambda'^2}{8\pi N_{\max}N_{\min}s'\left(s'^2\tau^2+\pi^2\right)\left(s'^2\tau^2+4\pi^2\right)}\Bigg[-\left(s'^2\tau^2\left(1-4e^{-s't}\right)+4\pi^2\left(1-e^{-s't}\right)\right)\pi N_{\text{osc}}^{-}s'^2\tau^2\cos[\varphi] \\
&\quad -\left(s'^2\tau^2\left(1-8e^{-s't}+5e^{-2s't}+s'^4\tau^4e^{-2s't}\right)+4\pi^2\left(1-e^{-s't}\right)^2\right)\pi^2N_{\text{osc}}^{-}s'\tau\sin[\varphi]+6\pi N_{\min}s'^4\tau^4e^{-2s't} \\
&\quad +\pi N_{\text{osc}}^{+}\left(s'^4\tau^4\left(1-4e^{-s't}+2s'te^{-2s't}\right)+\left(5\pi^2s'^2\tau^2+4\pi^4\right)\left(1-4e^{-s't}+(2s't+3)e^{-2s't}\right)\right)\Bigg]
\end{aligned}
\tag{S39}
$$

##### S3.2.6 Diagram 6 (left)

**Constant size:**

$$
\begin{aligned}
\kappa_{2D6L} &= -\frac{p_0}{(4N)^2} \ 2\times 2 \ \int_0^t dt_1 dt_2 \ G(t_1,0)G^2(t_2,t_1)G^2(t,t_2) \\
&= -\frac{p_0}{(2N)^2} \ \int_0^t dt_1 dt_2 \ \Theta(t_1)\Theta^2(t_2-t_1)\Theta^2(t-t_2) \ e^{-s'(2t-t_1)} \\
&= -\frac{p_0}{\gamma'^2} \ \left(e^{-s't}-(s't+1)e^{-2s't}\right)
\end{aligned}
\tag{S40}
$$

**Exponential growth:**

$$
\begin{aligned}
\kappa_{2D6L} &= -p_0 \ 2\times 2 \ \int_0^t dt_1 dt_2 \ \frac{1}{4N_{\exp}(t_1)\times 4N_{\exp}(t_2)} G(t_1,0)G^2(t_2,t_1)G^2(t,t_2) \\
&= -p_0 \ \int_0^t dt_1 dt_2 \ \Theta(t_1)\Theta^2(t_2-t_1)\Theta^2(t-t_2)\frac{e^{-s'(2t-t_1)}}{2N_{\exp}(t_1)\times 2N_{\exp}(t_2)} \\
&= -\frac{p_0}{\gamma'^2_{\exp}}\left(\frac{s'-\Gamma}{\Gamma}\right)\left(\frac{s'-\Gamma}{s'-2\Gamma}e^{-s't}\left(e^{-2\Gamma t}-e^{-s't}\right)-e^{-(s'+\Gamma)t}\left(e^{-\Gamma t}-e^{-s't}\right)\right)
\end{aligned}
\tag{S41}
$$

**Bottleneck:**

$$
\begin{aligned}
\kappa_{2D6L} &= -p_0 \; 2\times 2 \int_0^t dt_1 dt_2 \; \frac{1}{4N_{\mathrm{BN}}(t_1)\times 4N_{\mathrm{BN}}(t_2)} G(t_1,0)G^2(t_2,t_1)G^2(t,t_2) \\
&= -p_0 \int_0^t dt_1 dt_2 \; \Theta(t_1)\Theta^2(t_2-t_1)\Theta^2(t-t_2)\frac{e^{-s'(2t-t_1)}}{2N_{\mathrm{BN}}(t_1)\times 2N_{\mathrm{BN}}(t_2)} \\
&= -\frac{p_0}{{\gamma'}_0^2}\Bigg[ (1-\Theta_i)\, e^{-s't}\left(1-(s't+1)e^{-s't}\right) \\
&\quad + \frac{1}{{N_b}^2}(\Theta_i-\Theta_{i+b})\left(N_0^2 e^{-s't} - N_{\mathrm{bn}}^- e^{-s'(2t-T_i)}(N_0 s'(t-T_i)+N_{\mathrm{bn}}^+) - N_b e^{-2s't}\left(N_0 s'(t-T_i)+N_b(s'T_i+1)\right)\right) \\
&\quad + \frac{1}{{N_b}^2}\Theta_{i+b}\Bigg(N_{\mathrm{bn}}^- e^{-s'(2t-T_i+T_b)}(N_{\mathrm{bn}}^+ + N_b s'(t-T_i-T_b)) - N_{\mathrm{bn}}^- e^{-s'(2t-s'T_i)}(N_{\mathrm{bn}}^+ + N_0 s'T_b + N_b s'(t-T_i-T_b)) \\
&\quad - N_b e^{-2s't}\left(N_0 s'T_b + N_b\left(s'(t-T_b)+1\right)\right) + {N_b}^2 e^{-s't}\Bigg)\Bigg]
\end{aligned}
\tag{S42}
$$

**Oscillating size:**

$$
\begin{aligned}
\kappa_{2D6L} &= -p_0 \; 2\times 2 \int_0^t dt_1 dt_2 \; \frac{1}{4N_{\mathrm{osc}}(t_1)\times 4N_{\mathrm{osc}}(t_2)} G(t_1,0)G^2(t_2,t_1)G^2(t,t_2) \\
&= -p_0 \int_0^t dt_1 dt_2 \; \Theta(t_1)\Theta^2(t_2-t_1)\Theta^2(t-t_2)\frac{e^{-s'(2t-t_1)}}{2N_{\mathrm{osc}}(t_1)\times 2N_{\mathrm{osc}}(t_2)} \\
&= -\frac{p_0}{32{N_{\max}}^2{N_{\min}}^2}\left[ -\frac{4{s'}^2\tau^4 N_{\mathrm{osc}}^- N_{\mathrm{osc}}^+}{\left({s'}^2\tau^2+4\pi^2\right)^2} e^{-s't}\cos[\varphi] \quad + \quad \frac{1}{\left(\pi {s'}^2\tau^2+16\pi^3\right)\left({s'}^3\tau^2+4\pi^2 s'\right)^2}\times f^{\mathrm{osc}}_{D6L}(t)\right]
\end{aligned}
\tag{S43}
$$

The function $f^{\mathrm{osc}}_{D6L}$ in the latter part of the above expression, used only to simplify the result, has the following dependence on time, $s'$, and the parameters that define $N_{\mathrm{osc}}$.

$$
\begin{aligned}
f^{\mathrm{osc}}_{D6L}(t) &= 2s'\tau N_{\mathrm{osc}}^-\left({s'}^2\tau^2+16\pi^2\right)\Bigg(2\pi^2{s'}^2\tau^2(N_{\max}+3N_{\min})e^{-2s't} - 2\pi^2 N_{\mathrm{osc}}^+ e^{-s't}\left(3{s'}^2\tau^2+4\pi^2\right) \\
&+ \left(8\pi^4 N_{\mathrm{osc}}^+ + N_{\min}{s'}^4\tau^4\right)e^{-2s't}\Bigg)\sin[\varphi] - \pi{s'}^2\tau^2 {N_{\mathrm{osc}}^-}^2 e^{-s't}\left(32\pi^4+4\pi^2{s'}^2\tau^2-{s'}^4\tau^4\right)\cos[2\varphi] \\
&+ 6\pi^2{s'}^3\tau^3{N_{\mathrm{osc}}^-}^2 e^{-s't}\left({s'}^2\tau^2+4\pi^2\right)\sin[2\varphi] + \pi{N_{\min}}^2\Bigg(-32\pi^4{s'}^2\tau^2 e^{-s't}\left(11e^{s't}-(13s't+10)e^{-s't}\right) \\
&+ 512\pi^6 e^{-s't}\left(1-(s't+1)e^{-s't}\right) + {s'}^6\tau^6 e^{-s't}\left(3-4(s't+2)e^{-s't}\right) + 4\pi^2{s'}^4\tau^4 e^{-s't}\left(17-(22s't+32)e^{-s't}\right)\Bigg) \\
&+ \pi{N_{\max}}^2\Bigg(512\pi^6 e^{-s't}\left(1-(s't+1)e^{-s't}\right) + 3{s'}^6\tau^6 e^{-s't} + 4\pi^2{s'}^4\tau^4 e^{-s't}\left(17-2s'te^{-s't}\right) \\
&+ 32\pi^4{s'}^2\tau^2 e^{-s't}\left(11-5(s't+2)e^{-s't}\right)\Bigg) + 2\pi N_{\max}N_{\min}\Bigg(e^{-s't}\left({s'}^6\tau^6+28\pi^2{s'}^4\tau^4+224\pi^4{s'}^2\tau^2+512\pi^6\right) \\
&- 2e^{-2s't}\left({s'}^2\tau^2+4\pi^2\right)^2\left({s'}^2\tau^2(s't)+16\pi^2(s't+1)\right)\Bigg)
\end{aligned}
\tag{S44}
$$

###### S3.2.7 Diagram 6 (right)

**Constant size:**

$$\begin{aligned}
\kappa_{2D6R} &= -\frac{\mu}{(4N)^2}\ 2\times 2\int_0^t dt_1 dt_2 dt_3\ G(t_2,t_1)G^2(t_3,t_2)G^2(t,t_3)\\
&= -\frac{\mu}{(2N)^2}\int_0^t dt_1 dt_2 dt_3\ \Theta(t_2-t_1)\Theta^2(t_3-t_2)\Theta^2(t-t_3)\ e^{-s'(2t-t_2-t_1)}\\
&= -\frac{\lambda'}{4\gamma'^2}\left(1-4e^{-s't}+(2s't+3)e^{-2s't}\right) \qquad \text{(S45)}
\end{aligned}$$

**Exponential growth:**

$$\begin{aligned}
\kappa_{2D6R} &= -\mu\ 2\times 2\int_0^t dt_1 dt_2 dt_3\ \frac{1}{4N_{\text{exp}}(t_2)\times 4N_{\text{exp}}(t_3)}G(t_2,t_1)G^2(t_3,t_2)G^2(t,t_3)\\
&= -\mu\int_0^t dt_1 dt_2 dt_3\ \Theta(t_2-t_1)\Theta^2(t_3-t_2)\Theta^2(t-t_3)\frac{e^{-s'(2t-t_2-t_1)}}{2N_{\text{exp}}(t_2)\times 2N_{\text{exp}}(t_3)}\\
&= -\frac{\lambda'}{2\gamma'^2_{\text{exp}}}\left(\frac{s'-\Gamma}{\Gamma}\right)\left(\frac{s'}{(s'-2\Gamma)}e^{-2s't}+\Gamma e^{-2\Gamma t}\left(\frac{1}{(2s'-\Gamma)}-\frac{2}{(s'-2\Gamma)}e^{-s't}\right)-\frac{2s'}{(2s'-\Gamma)}e^{-(2s'+\Gamma)t}\right) \qquad \text{(S46)}
\end{aligned}$$

**Bottleneck:**

$$\begin{aligned}
\kappa_{2D6R} &= -\mu\ 2\times 2\int_0^t dt_1 dt_2 dt_3\ \frac{1}{4N_{\text{BN}}(t_2)\times 4N_{\text{BN}}(t_3)}G(t_2,t_1)G^2(t_3,t_2)G^2(t,t_3)\\
&= -\mu\int_0^t dt_1 dt_2 dt_3\ \Theta(t_2-t_1)\Theta^2(t_3-t_2)\Theta^2(t-t_3)\frac{e^{-s'(2t-t_2-t_1)}}{2N_{\text{BN}}(t_2)\times 2N_{\text{BN}}(t_3)}\\
&= -\frac{\lambda'}{4\gamma'^2_0 N_b}\Bigg[N_b\left(1-4e^{-s't}+(2s't+3)e^{-2s't}\right)+N^-_{\text{bn}}\Theta_i\left(\left(1-e^{-s'(t-T_i)}\right)\left(1-4e^{-s't}+e^{-s'(t-T_i)}\right)+2s'(t-T_i)e^{-2s't}\right)\\
&\qquad -N^-_{\text{bn}}\Theta_{i+b}\left(1-4e^{-s't}+2s'\left(t-(T_i+T_b)\right)e^{-2s't}+4e^{-s'(2t-(T_i+T_b))}-e^{-2s'(t-(T_i+T_b))}\right)\Bigg] \qquad \text{(S47)}
\end{aligned}$$

**Oscillating size:**

$$
\begin{aligned}
&\kappa_{2D6R} = -\mu\ 2\times 2 \int_0^t dt_1 dt_2 dt_3\ \frac{1}{4N_{\mathrm{osc}}(t_2)\times 4N_{\mathrm{osc}}(t_3)} G(t_2,t_1)G^2(t_3,t_2)G^2(t,t_3)\\
&= -\mu \int_0^t dt_1 dt_2 dt_3\ \Theta(t_2-t_1)\Theta^2(t_3-t_2)\Theta^2(t-t_3)\frac{e^{-s'(2t-t_2-t_1)}}{2N_{\mathrm{osc}}(t_2)\times 2N_{\mathrm{osc}}(t_3)}\\
&= \frac{\lambda'}{128{N_{\mathrm{max}}}^2{N_{\mathrm{min}}}^2 s'^2\left(\pi s'^2\tau^2+16\pi^3\right)\left(s'^4\tau^4+5\pi^2 s'^2\tau^2+4\pi^4\right)^2} \quad\times\\
&\Bigg[\pi\Bigg(e^{-2s't}\Big[-8{N_{\mathrm{min}}}^2 s'^{10}\tau^{10}(s't+3)-12{N_{\mathrm{min}}}^2\pi^2 s'^8\tau^8(15s't+32)-4{N_{\mathrm{min}}}^2\pi^4 s'^6\tau^6(235s't+291)\\
&-8{N_{\mathrm{min}}}^2\pi^6 s'^4\tau^4(232s't+333)\Big]+4N^-_{\mathrm{osc}}N^+_{\mathrm{osc}}s'^4\tau^4\left(s'^2\tau^2+16\pi^2\right)\left(\left(s'^2\tau^2+4\pi^2\right)^2-4e^{-s't}\left(s'^2\tau^2+\pi^2\right)^2\right)\cos[\varphi]\\
&-96{N_{\mathrm{min}}}^2\pi^8 s'^2\tau^2(22s't+41)e^{-2s't}+N^-_{\mathrm{osc}}s'\tau\Big[-16N^+_{\mathrm{osc}}\pi\left(3s'^4\tau^4+52\pi^2 s'^2\tau^2+64\pi^4\right)\left(s'^2\tau^2+\pi^2\right)^2 e^{-s't}\sin[\varphi]\\
&-N^-_{\mathrm{osc}}s'\tau\left(s'^4\tau^4+5\pi^2 s'^2\tau^2+4\pi^4\right)\left(\left(-4+e^{s't}\right)s'^4\tau^4 e^{-s't}+14\left(1+2e^{-s't}\right)\pi^2 s'^2\tau^2-32\left(1-e^{-s't}\right)\pi^4\right)\cos[\varphi]\Big]\\
&-512{N_{\mathrm{min}}}^2\pi^{10}(2s't+3)e^{-2s't}\\
&-\left(s'^2\tau^2+\pi^2\right)\left(s'^2\tau^2+4\pi^2\right)^2\left(s'^2\tau^2+16\pi^2\right)\left(2\pi^2{N^+_{\mathrm{osc}}}^2+\left(3{N_{\mathrm{max}}}^2+2N_{\mathrm{min}}N_{\mathrm{max}}+3{N_{\mathrm{min}}}^2\right)s'^2\tau^2\right)\\
&+4e^{-s't}\left(s'^2\tau^2+\pi^2\right)^2\left(8\pi^2{N^+_{\mathrm{osc}}}^2+\left(3{N_{\mathrm{max}}}^2+2N_{\mathrm{min}}N_{\mathrm{max}}+3{N_{\mathrm{min}}}^2\right)s'^2\tau^2\right)\left(s'^4\tau^4+20\pi^2 s'^2\tau^2+64\pi^4\right)\\
&-8N_{\mathrm{max}}N_{\mathrm{min}}e^{-2s't}\left(s'^2\tau^2+\pi^2\right)^2\left(s'^2\tau^2+4\pi^2\right)\left(s'^4\tau^4(s't)+10\pi^2(2s't+3)s'^2\tau^2+32\pi^4(2s't+3)\right)\\
&+4{N_{\mathrm{max}}}^2\pi^2 e^{-2s't}\Big[-7s'^8\tau^8(s't)-\pi^2(151s't+315)s'^6\tau^6-42\pi^4(16s't+25)s'^4\tau^4-8\pi^6(98s't+123)s'^2\tau^2\\
&-128\pi^8(2s't+3)\Big]\Bigg)+N^-_{\mathrm{osc}}s'\tau\Bigg(4\left(s'^4\tau^4+20\pi^2 s'^2\tau^2+64\pi^4\right)e^{-s't}\Big[\Big(N_{\mathrm{min}}s'^6\tau^6+(5N_{\mathrm{max}}+4N_{\mathrm{min}})\pi^2 s'^4\tau^4\\
&+2(6N_{\mathrm{max}}+5N_{\mathrm{min}})\pi^4 s'^2\tau^2+4N^+_{\mathrm{osc}}\pi^6\Big)\cosh[s't]\\
&-s'^2\tau^2\left(N_{\mathrm{min}}s'^4\tau^4+(2N_{\mathrm{max}}+N_{\mathrm{min}})\pi^2 s'^2\tau^2-(N_{\mathrm{max}}+3N_{\mathrm{min}})\pi^4\right)\sinh[s't]\Big]\sin[\varphi]\\
&-3N^-_{\mathrm{osc}}\pi^2 s'^2\tau^2\left(\left(1-8e^{-s't}\right)s'^2\tau^2+8\left(2-e^{-s't}\right)\pi^2\right)\left(s'^4\tau^4+5\pi^2 s'^2\tau^2+4\pi^4\right)\sin[2\varphi]\Bigg)\Bigg] \qquad \text{(S48)}
\end{aligned}
$$

Here, sinh and cosh are the hyperbolic sine and hyperbolic cosine functions, respectively.

#### S3.3 Diagrams for the third cumulant $\kappa_3(t)$

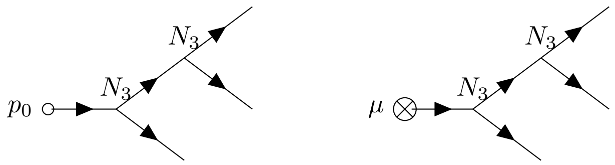


(D7)

The counting factors for these diagrams are less apparent than for the above diagrams. Here, the total multiplicative factor is $\frac{2!\times 3!\times 2}{2!} = 2\times 3!$. The factor of 2! is from the choice of the order of $N_3$ vertices to connect to the terminal ($p_0$ or $\mu$) vertex, which is cancelled by a repeated vertex factor of 2! in the denominator.

The factor of 3! denotes the number of ways to connect edges exiting the internal vertices to the three external vertices. The additional factor of two comes from the choice of which $\tilde{p}$ exiting the first $N_3$ vertex is connected to the internal ($N_3$) vs. external vertex. This final factor is generally the hardest to identify, as it doubles the number of allowed topologies without looking particularly visually symmetric. In contrast, note that both edges exiting the second $N_3$ vertex are connected to external vertices, so the symmetry of this feature of the topology is already accounted for in the 3! associated with the choice of three external vertices. This example is also detailed in Appendix A2, which enumerates the allowed Wick contractions for Diagram D7. The counting factor is the same for both the left and right diagrams in D7, as well as in all demographies detailed below.

##### S3.3.1 Diagram 7 (left)

**Constant size**

$$\begin{aligned}
\kappa_{3D7L} &= \frac{p_0}{(4N)^2}\,\frac{2!\times 3!\times 2}{2!}\int_0^t dt_1 dt_2\; G(t_1,0)G(t_2,t_1)G(t,t_1)G^2(t,t_2) \\
&= \frac{3p_0}{(2N)^2}\int_0^t dt_1 dt_2\;\Theta(t_1)\Theta(t_2-t_1)\Theta(t-t_1)\Theta^2(t-t_2)\; e{-}s'(3t-t_2-t_1) \\
&= \frac{3p_0}{2\gamma'^2}e^{-s't}\left(1-e^{-s't}\right)^2
\end{aligned} \tag{S49}$$

**Exponential growth**

$$\begin{aligned}
\kappa_{3D7L} &= p_0\,\frac{2!\times 3!\times 2}{2!}\int_0^t dt_1 dt_2\;\frac{1}{4N_{\mathrm{exp}}(t_1)\times 4N_{\mathrm{exp}}(t_2)}G(t_1,0)G(t_2,t_1)G(t,t_1)G^2(t,t_2) \\
&= 3p_0\int_0^t dt_1 dt_2\;\Theta(t_1)\Theta(t_2-t_1)\Theta(t-t_1)\Theta^2(t-t_2)\frac{e{-}s'(3t-t_2-t_1)}{2N_{\mathrm{exp}}(t_1)\times 2N_{\mathrm{exp}}(t_2)} \\
&= \frac{3p_0}{2\gamma'^2_{\mathrm{exp}}}\;e^{-s't}\;\left(e^{-\Gamma t}-e^{-s't}\right)^2
\end{aligned} \tag{S50}$$

**Bottleneck**

$$\begin{aligned}
\kappa_{3D7L} &= p_0\,\frac{2!\times 3!\times 2}{2!}\int_0^t dt_1 dt_2\;\frac{1}{4N_{\mathrm{BN}}(t_1)\times 4N_{\mathrm{BN}}(t_2)}G(t_1,0)G(t_2,t_1)G(t,t_1)G^2(t,t_2) \\
&= 3p_0\int_0^t dt_1 dt_2\;\Theta(t_1)\Theta(t_2-t_1)\Theta(t-t_1)\Theta^2(t-t_2)\frac{e{-}s'(3t-t_2-t_1)}{2N_{\mathrm{BN}}(t_1)\times 2N_{\mathrm{BN}}(t_2)} \\
&= \frac{3p_0}{2\gamma'^2_0}\Bigg[(1-\Theta_i)e^{-s't}\left(1-e^{-s't}\right)^2+(\Theta_i-\Theta_{i+b})\frac{e^{-3s't}}{{N_b}^2}\left(N_{\mathrm{bn}}^{-}e^{s'T_i}-N_0e^{s't}+N_b\right)^2 \\
&\qquad +\Theta_{i+b}\frac{e^{-3s't}}{{N_b}^2}\left(N_{\mathrm{bn}}^{-}e^{s'T_i}\left(e^{s'T_b}-1\right)+N_b\left(e^{s't}-1\right)\right)^2\Bigg]
\end{aligned} \tag{S51}$$

**Oscillating size**

$$
\begin{aligned}
\kappa_{3D7L} &= p_0 \, \frac{2! \times 3! \times 2}{2!} \int_0^t dt_1 dt_2 \, \frac{1}{4N_{\text{osc}}(t_1) \times 4N_{\text{osc}}(t_2)} G(t_1, 0) G(t_2, t_1) G(t, t_1) G^2(t, t_2) \\
&= 3p_0 \int_0^t dt_1 dt_2 \, \Theta(t_1)\Theta(t_2 - t_1)\Theta(t - t_1)\Theta^2(t - t_2) \frac{e{-}s'(3t - t_2 - t_1)}{2N_{\text{osc}}(t_1) \times 2N_{\text{osc}}(t_2)} \\
&= \frac{3p_0}{32{N_{\max}}^2 {N_{\min}}^2 s'^2 \left(4\pi^2 + s'^2\tau^2\right)^2} e^{-2s't} \Bigg( 2s'\tau N_{\text{osc}}^- \cosh\left[\frac{s't}{2}\right] \left(s'\tau \, \sin^2[\varphi] - \pi \sin[\varphi]\right) \\
&\quad + \sinh\left[\frac{s't}{2}\right] \left( {s'}^2 \tau^2 (N_{\text{osc}}^+ + 2N_{\min}) - s'\tau N_{\text{osc}}^- \left(s'\tau \, \cos[\varphi] + 2\pi \sin[\varphi]\right) + 8\pi^2 N_{\text{osc}}^+ \right) \Bigg)^2 \qquad \text{(S52)}
\end{aligned}
$$

##### S3.3.2 Diagram 7 (right)

**Constant size**

$$
\begin{aligned}
\kappa_{3D7R} &= \frac{\mu}{(4N)^2} \, \frac{2! \times 3! \times 2}{2!} \int_0^t dt_1 dt_2 dt_3 \, G(t_2, t_1) G(t_3, t_2) G(t, t_2) G^2(t, t_3) \\
&= \frac{3\mu}{(2N)^2} \int_0^t dt_1 dt_2 dt_3 \, \Theta(t_2 - t_1)\Theta(t_3 - t_2)\Theta(t - t_2)\Theta^2(t - t_3) \; e^{-s'(3t - t_3 - t_2 - t_1)} \\
&= \frac{\lambda'}{2\gamma'^2} \left(1 - e^{-s't}\right)^3 \qquad \text{(S53)}
\end{aligned}
$$

**Exponential growth**

$$
\begin{aligned}
\kappa_{3D7R} &= \mu \, \frac{2! \times 3! \times 2}{2!} \int_0^t dt_1 dt_2 dt_3 \, \frac{1}{4N_{\exp}(t_2) \times 4N_{\exp}(t_3)} G(t_2, t_1) G(t_3, t_2) G(t, t_2) G^2(t, t_3) \\
&= 3\mu \int_0^t dt_1 dt_2 dt_3 \, \Theta(t_2 - t_1)\Theta(t_3 - t_2)\Theta(t - t_2)\Theta^2(t - t_3) \frac{e^{-s'(3t - t_3 - t_2 - t_1)}}{2N_{\exp}(t_2) \times 2N_{\exp}(t_3)} \\
&= \frac{3\lambda'}{2{\gamma'}_{\exp}^2} \left( \left(e^{-2\Gamma t} - e^{-(2\Gamma + s')t}\right) - \frac{2s'\left(e^{-2\Gamma t} - e^{-(\Gamma + 2s')t}\right)}{2s' - \Gamma} + \frac{s'\left(e^{-2\Gamma t} - e^{-3s't}\right)}{3s' - 2\Gamma} \right) \qquad \text{(S54)}
\end{aligned}
$$

**Bottleneck**

$$
\begin{aligned}
\kappa_{3D7R} &= \mu \, \frac{2! \times 3! \times 2}{2!} \int_0^t dt_1 dt_2 dt_3 \, \frac{1}{4N_{\text{BN}}(t_2) \times 4N_{\text{BN}}(t_3)} G(t_2, t_1) G(t_3, t_2) G(t, t_2) G^2(t, t_3) \\
&= 3\mu \int_0^t dt_1 dt_2 dt_3 \, \Theta(t_2 - t_1)\Theta(t_3 - t_2)\Theta(t - t_2)\Theta^2(t - t_3) \frac{e^{-s'(3t - t_3 - t_2 - t_1)}}{2N_{\text{BN}}(t_2) \times 2N_{\text{BN}}(t_3)} \\
&= \frac{\lambda' N_{\text{bn}}^-}{2{\gamma'}_0^2 {N_b}^2} \Bigg[ \Theta_i e^{-2s't} \left(1 - e^{-s'(t - T_i)}\right) \left( N_{\text{bn}}^+ \left(e^{s'(t + T_i)} - 3e^{s't} + e^{2s't}\right) + N_{\text{bn}}^- (3e^{s'T_i} - e^{2s'T_i}) + N_0 e^{2s'T_i} + 3N_b \right) \\
&\quad - \Theta_{i+b} e^{-2s't} \left(1 - e^{-s'(t - T_i - T_b)}\right) \Bigg( N_{\text{bn}}^- \left(e^{2s'(T_i + T_b)} + 6e^{s'T_i} - 3e^{2s'T_i} - 3e^{s'(T_i + T_b)}\right) \\
&\quad + N_{\text{bn}}^+ \left(e^{s'(t + T_i + T_b)} - 3e^{s't} + e^{2s't}\right) + N_b(3 - e^{2s'(T_i + T_b)}) \Bigg) \Bigg] + \frac{\lambda'}{2{\gamma'}_0^2} \left(1 - e^{-s't}\right)^3 \qquad \text{(S55)}
\end{aligned}
$$

**Oscillating size**

$$
\begin{aligned}
\kappa_{3D7R} &= \mu\ \frac{2!\times 3!\times 2}{2!}\ \int_0^t dt_1 dt_2 dt_3\ \frac{1}{4N_{\mathrm{osc}}(t_2)\times 4N_{\mathrm{osc}}(t_3)} G(t_2,t_1)G(t_3,t_2)G(t,t_2)G^2(t,t_3)\\
&= 3\mu \int_0^t dt_1 dt_2 dt_3\ \Theta(t_2-t_1)\Theta(t_3-t_2)\Theta(t-t_2)\Theta^2(t-t_3)\frac{e^{-s'(3t-t_3-t_2-t_1)}}{2N_{\mathrm{osc}}(t_2)\times 2N_{\mathrm{osc}}(t_3)}\\
&= \frac{\lambda'}{64{N_{\mathrm{max}}}^2{N_{\mathrm{min}}}^2 s'^2\left(s'^2\tau^2+\pi^2\right)\left(s'^2\tau^2+4\pi^2\right)^2\left(9s'^2\tau^2+4\pi^2\right)\left(9s'^2\tau^2+16\pi^2\right)}\times\\
&\Bigg[\left(s'^2\tau^2+4\pi^2\right)^2\Bigg[\left(81s'^4\tau^4+180\pi^2s'^2\tau^2+64\pi^4\right)\left(s'^2\tau^2\left(3{N_{\mathrm{max}}}^2+2N_{\mathrm{max}}N_{\mathrm{min}}+3{N_{\mathrm{min}}}^2\right)+2\pi^2{N^+_{\mathrm{osc}}}^2\right)\\
&+3s'\tau N^-_{\mathrm{osc}}\Bigg(\left(9s'^2\tau^2+4\pi^2\right)\left(7\pi s'^2\tau^2N^-_{\mathrm{osc}}\sin[2\varphi]+s'\tau N^-_{\mathrm{osc}}\left(3s'^2\tau^2-4\pi^2\right)\cos[2\varphi]\right)\\
&-2\pi N^+_{\mathrm{osc}}\left(63s'^4\tau^4+130\pi^2s'^2\tau^2+32\pi^4\right)\sin[\varphi]-2s'\tau N^+_{\mathrm{osc}}\left(6s'^2\tau^2+\pi^2\right)\left(9s'^2\tau^2+16\pi^2\right)\cos[\varphi]\Bigg)\Bigg]\\
&-6e^{-s't}\left(9s'^2\tau^2+4\pi^2\right)\left(9s'^2\tau^2+16\pi^2\right)\Bigg[\left(s'^2\tau^2+\pi^2\right)\left(N^+_{\mathrm{osc}}\left(s'^2\tau^2+4\pi^2\right)-s'\tau N^-_{\mathrm{osc}}\left(s'\tau\cos[\varphi]+2\pi\sin[\varphi]\right)\right)\\
&-\left(\pi^2s'^2\tau^2(7N_{\mathrm{max}}+3N_{\mathrm{min}})+4\pi^4N^+_{\mathrm{osc}}+2N_{\mathrm{min}}s'^4\tau^4\right)\Bigg]\left(N^+_{\mathrm{osc}}\left(s'^2\tau^2+4\pi^2\right)-s'\tau N^-_{\mathrm{osc}}\left(s'\tau\cos[\varphi]+2\pi\sin[\varphi]\right)\right)\\
&-{N_{\mathrm{max}}}^2e^{-3s't}\left(6240\pi^4s'^6\tau^6+16352\pi^6s'^4\tau^4+12160\pi^8s'^2\tau^2+2048\pi^{10}\right)\\
&-N_{\mathrm{max}}N_{\mathrm{min}}e^{-3s't}\left(3744\pi^2s'^8\tau^8+14048\pi^4s'^6\tau^6+23360\pi^6s'^4\tau^4+17152\pi^8s'^2\tau^2+4096\pi^{10}\right)\\
&-{N_{\mathrm{min}}}^2e^{-3s't}\left(648s'^{10}\tau^{10}+3528\pi^2s'^8\tau^8+8736\pi^4s'^6\tau^6+9824\pi^6s'^4\tau^4+6016\pi^8s'^2\tau^2+2048\pi^{10}\right)\Bigg]
\end{aligned}
\tag{S56}
$$

#### S3.4 Diagrams for the fourth cumulant $\kappa_4(t)$

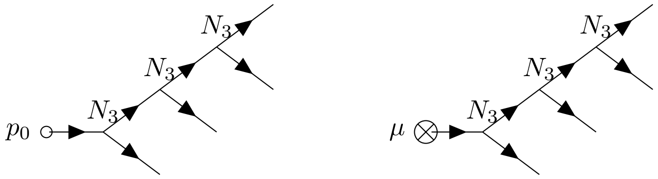


(D8)

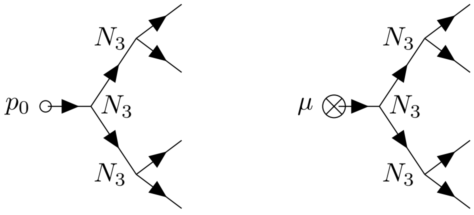


(D9)

As with the third cumulant $\kappa_3(t)$, the counting factors for the above diagrams are less obvious than those for the lower moments. The same principle discussed for Diagram D7 applies to Diagram D8: there is an overall multiplicative factor of $\frac{3!\times 4!\times 4}{3!}$, with the counting factor of 3! coming from the number of ways to order the $N_3$ vertices, along with the cancellation by the repeated vertex factor 1/3!, the factor of 4! comes from the number of ways to connect the internal to the four external vertices, and the factor of 4 comes from the two possible arrangements of the connections exiting first and second $N_3$ vertices, one of which connects to the subsequent $N_3$ vertex and the other to one of the external vertices. In contrast, the overall multiplicative factor for Diagram D9 is simpler: $\frac{3!\times 4!}{3!}$. In this case, the counting factors only represent the number of ways to order the $N_3$ vertices and connect to the external vertices. The symmetry associated with the maximally connected diagram structure does not allow for additional counting factors from the choice of connections between internal and external vertices; the factor of 3! already accounts for the choices associated with internal vertices and all external vertices are connected to the later-time $N_3$ vertices, with all possible choices of connections to the external vertices covered by the counting factor of 4! (similar to the final $N_3$ vertex in D7 or $D$8). Finally, note that the time dependence of the three coupling constants $1/4N(t_i)$ (for $i \in \{1,2,3,4\}$) associated with the time of the three $N_3$ vertices (to be integrated over) in D8 and D9 substantially complicates the integrals for the bottleneck and cyclical demographies.

##### S3.4.1 Diagram 8 (left)

**Constant size**

$$\begin{aligned}\kappa_{4D8L} &= \frac{p_0}{(4N)^3}\,\frac{4\times 3!\times 4!}{3!}\int_0^t dt_1dt_2dt_3\; G(t_1,0)G(t_2,t_1)G(t,t_1)G(t_3,t_2)G(t,t_2)G^2(t,t_3)\\ &= \frac{12p_0}{(4N)^3}\int_0^t dt_1dt_2dt_3\;\Theta(t_1)\Theta(t_2-t_1)\Theta(t-t_1)\Theta(t_3-t_2)\Theta(t-t_2)\Theta^2(t-t_3)\; e^{-s'(4t-t_3-t_2-t_1)}\\ &= \frac{2p_0}{\gamma'^3}e^{-s't}\left(1-e^{-s't}\right)^3 \end{aligned} \tag{S57}$$

**Exponential growth**

$$\begin{aligned}\kappa_{4D8L} &= p_0\;\frac{4\times 3!\times 4!}{3!}\int_0^t dt_1dt_2dt_3\;\frac{G(t_1,0)G(t_2,t_1)G(t,t_1)G(t_3,t_2)G(t,t_2)G^2(t,t_3)}{4N_{\text{exp}}(t_1)\times 4N_{\text{exp}}(t_2)\times 4N_{\text{exp}}(t_3)}\\ &= 12p_0\int_0^t dt_1dt_2dt_3\;\frac{\Theta(t_1)\Theta(t_2-t_1)\Theta(t-t_1)\Theta(t_3-t_2)\Theta(t-t_2)\Theta^2(t-t_3)\; e^{-s'(4t-t_3-t_2-t_1)}}{2N_{\text{exp}}(t_1)\times 2N_{\text{exp}}(t_2)\times 2N_{\text{exp}}(t_3)}\\ &= \frac{2p_0}{\gamma'^3_{\text{exp}}}e^{-s't}\left(e^{-\Gamma t}-e^{-s't}\right)^3 \end{aligned} \tag{S58}$$

**Bottleneck**

$$
\begin{aligned}
\kappa_{4D8L} &= p_0 \ \frac{4\times 3!\times 4!}{3!} \int_0^t dt_1 dt_2 dt_3 \ \frac{G(t_1,0)G(t_2,t_1)G(t,t_1)G(t_3,t_2)G(t,t_2)G^2(t,t_3)}{4N_{\mathrm{BN}}(t_1)\times 4N_{\mathrm{BN}}(t_2)\times 4N_{\mathrm{BN}}(t_3)} \\
&= 12p_0 \int_0^t dt_1 dt_2 dt_3 \ \frac{\Theta(t_1)\Theta(t_2-t_1)\Theta(t-t_1)\Theta(t_3-t_2)\Theta(t-t_2)\Theta^2(t-t_3) \ e^{-s'(4t-t_3-t_2-t_1)}}{2N_{\mathrm{BN}}(t_1)\times 2N_{\mathrm{BN}}(t_2)\times 2N_{\mathrm{BN}}(t_3)} \\
&= \frac{2p_0}{\gamma_0'^3}\Bigg[(1-\Theta_i)e^{-s't}\left(1-e^{-s't}\right)^3 + \frac{(\Theta_i-\Theta_{i+b})}{{N_b}^3}e^{-s't}\left(N_0 - N_b e^{-s't} - N_{\mathrm{bn}}^- e^{-s'(t-T_i)}\right)^3 \\
&\quad - \frac{\Theta_{i+b}}{{N_b}^3}\Bigg(6{N_b}^2 N_{\mathrm{bn}}^- e^{-s'(3t-(T_i+T_b))} - 3{N_b}^2 N_{\mathrm{bn}}^- e^{-s'(2t-(T_i+T_b))} - 6{N_b}^2 N_{\mathrm{bn}}^- e^{-s'(3t-T_i)} + 3{N_b}^2 N_{\mathrm{bn}}^- e^{-s'(2t-T_i)} \\
&\qquad - 3{N_b}^2 N_{\mathrm{bn}}^- e^{-s'(4t-(T_i+T_b))} + 3{N_b}^2 N_{\mathrm{bn}}^- e^{-s'(4t-T_i)} + 6N_b {N_{\mathrm{bn}}^-}^2 e^{-s'(3t-(2T_i+T_b))} - 3N_b {N_{\mathrm{bn}}^-}^2 e^{-s'(3t-2(T_i+T_b))} \\
&\quad - 3N_b {N_{\mathrm{bn}}^-}^2 e^{-s'(3t-2T_i)} - {N_{\mathrm{bn}}^-}^3 e^{-s'(4t-3(T_i+T_b))} - 3{N_{\mathrm{bn}}^-}^3 e^{-s'(4t-(3T_i+T_b))} + 3{N_{\mathrm{bn}}^-}^3 e^{-s'(4t-(3T_i+2T_b))} \\
&\qquad + 3N_b {N_{\mathrm{bn}}^-}^2 e^{-s'(4t-2(T_i+T_b))} - 6N_b {N_{\mathrm{bn}}^-}^2 e^{-s'(4t-(2T_i+T_b))} + {N_{\mathrm{bn}}^-}^3 e^{-s'(4t-3T_i)} + 3N_b {N_{\mathrm{bn}}^-}^2 e^{-s'(4t-2T_i)} \\
&\qquad - 3{N_b}^3 e^{-3s't} + 3{N_b}^3 e^{-2s't} - {N_b}^3 e^{-s't} + {N_b}^3 e^{-4s't}\Bigg)\Bigg] \qquad \text{(S59)}
\end{aligned}
$$

**Oscillating size**

$$
\begin{aligned}
\kappa_{4D8L} &= p_0 \ \frac{4\times 3!\times 4!}{3!} \int_0^t dt_1 dt_2 dt_3 \ \frac{G(t_1,0)G(t_2,t_1)G(t,t_1)G(t_3,t_2)G(t,t_2)G^2(t,t_3)}{4N_{\mathrm{osc}}(t_1)\times 4N_{\mathrm{osc}}(t_2)\times 4N_{\mathrm{osc}}(t_3)} \\
&= 12p_0 \int_0^t dt_1 dt_2 dt_3 \ \frac{\Theta(t_1)\Theta(t_2-t_1)\Theta(t-t_1)\Theta(t_3-t_2)\Theta(t-t_2)\Theta^2(t-t_3) \ e^{-s'(4t-t_3-t_2-t_1)}}{2N_{\mathrm{osc}}(t_1)\times 2N_{\mathrm{osc}}(t_2)\times 2N_{\mathrm{osc}}(t_3)} \\
&= \frac{p_0}{32{N_{\max}}^3 {N_{\min}}^3 s'^3\left(s'^2\tau^2+4\pi^2\right)^3} \ e^{-\frac{5}{2}s't} \ \Bigg(2s'\tau N_{\mathrm{osc}}^- \cosh\left[\frac{s't}{2}\right]\left(s'\tau\sin^2\left[\frac{\varphi}{2}\right] - \pi\sin[\varphi]\right) \\
&\qquad + \sinh\left[\frac{s't}{2}\right]\left(s'^2\tau^2(N_{\max}+3N_{\min}) - s'\tau N_{\mathrm{osc}}^-\left(s'\tau\cos[\varphi]+2\pi\sin[\varphi]\right) + 8\pi^2 N_{\mathrm{osc}}^+\right)\Bigg)^3 \qquad \text{(S60)}
\end{aligned}
$$

##### S3.4.2 Diagram 8 (right)

**Constant size**

$$
\begin{aligned}
\kappa_{4D8R} &= \frac{\mu}{(4N)^3} \ \frac{4\times 3!\times 4!}{3!} \int_0^t dt_1 dt_2 dt_3 dt_4 \ G(t_2,t_1)G(t_3,t_2)G(t,t_2)G(t_4,t_3)G(t,t_3)G^2(t,t_4) \\
&= \frac{12\mu}{(4N)^3} \int_0^t dt_1 dt_2 dt_3 dt_4 \ \Theta(t_2-t_1)\Theta(t_3-t_2)\Theta(t-t_2)\Theta(t_4-t_3)\Theta(t-t_3)\Theta^2(t-t_4) \ e^{-s'(4t-t_4-t_3-t_2-t_1)} \\
&= \frac{\lambda'}{2\gamma'^3}\left(1-e^{-s't}\right)^4 \qquad \text{(S61)}
\end{aligned}
$$

**Exponential growth**

$$
\begin{aligned}
\kappa_{4D8R} &= \mu \ \frac{4\times 3!\times 4!}{3!} \int_0^t dt_1 dt_2 dt_3 dt_4 \ \frac{G(t_2,t_1)G(t_3,t_2)G(t,t_2)G(t_4,t_3)G(t,t_3)G^2(t,t_4)}{4N_{\text{exp}}(t_2)\times 4N_{\text{exp}}(t_3)\times 4N_{\text{exp}}(t_4)} \\
&= 12\mu \int_0^t dt_1 dt_2 dt_3 dt_4 \ \frac{\Theta(t_2-t_1)\Theta(t_3-t_2)\Theta(t-t_2)\Theta(t_4-t_3)\Theta(t-t_3)\Theta^2(t-t_4) \ e^{-s'(4t-t_4-t_3-t_2-t_1)}}{2N_{\text{exp}}(t_2)\times 2N_{\text{exp}}(t_3)\times 2N_{\text{exp}}(t_4)} \\
&= -\frac{2\lambda'}{\gamma'^3_{\text{exp}}}\left(e^{-3\Gamma t}\left(e^{-s't}-1\right)+\frac{3s'\left(e^{-3\Gamma t}-e^{-2(s'+\Gamma)t}\right)}{2s'-\Gamma}-\frac{3s'\left(e^{-3\Gamma t}-e^{-2s't-2\Gamma t}\right)}{3s'-2\Gamma}+\frac{s'\left(e^{-3\Gamma t}-e^{-4s't}\right)}{4s'-3\Gamma}\right)
\end{aligned} \tag{S62}
$$

**Bottleneck**

$$
\begin{aligned}
\kappa_{4D8R} &= \mu \ \frac{4\times 3!\times 4!}{3!} \int_0^t dt_1 dt_2 dt_3 dt_4 \ \frac{G(t_2,t_1)G(t_3,t_2)G(t,t_2)G(t_4,t_3)G(t,t_3)G^2(t,t_4)}{4N_{\text{BN}}(t_2)\times 4N_{\text{BN}}(t_3)\times 4N_{\text{BN}}(t_4)} \\
&= 12\mu \int_0^t dt_1 dt_2 dt_3 dt_4 \ \frac{\Theta(t_2-t_1)\Theta(t_3-t_2)\Theta(t-t_2)\Theta(t_4-t_3)\Theta(t-t_3)\Theta^2(t-t_4) \ e^{-s'(4t-t_4-t_3-t_2-t_1)}}{2N_{\text{BN}}(t_2)\times 2N_{\text{BN}}(t_3)\times 2N_{\text{BN}}(t_4)} \\
&= \frac{\lambda'}{2\gamma'^3_0}\left((1-\Theta_i)\left(1-e^{-s't}\right)^4 \quad + \quad \frac{(\Theta_i-\Theta_{i+b})}{{N_b}^3} f^{\text{BN}}_{D8R}(t) \quad + \quad \frac{\Theta_{i+b}}{{N_b}^3} g^{\text{BN}}_{D8R}(t)\right)
\end{aligned} \tag{S63}
$$

The functions $f^{\text{BN}}_{D8R}$ and $g^{\text{BN}}_{D8R}$, used to simplify the the above expression, are defined below.

$$
\begin{aligned}
f^{\text{BN}}_{D8R}(t) = \Big(& N_0^3 - 4N_0^3 e^{-s't} - 6N^-_{\text{bn}}N_0^2 e^{-2s'(t-T_i)} + 12N^-_{\text{bn}}N_0^2 e^{-s'(2t-T_i)} + 6N_b N_0^2 e^{-2s't} - 12{N^-_{\text{bn}}}^2 N_0 e^{-s'(3t-2T_i)} \\
& - 4{N_b}^2 N_0 e^{-3s't} - 12N_b N_0 N^-_{\text{bn}} e^{-s'(3t-T_i)} + 4N_0\left(2N_0^2 - 3N_b N_0 + {N_b}^2\right) e^{-3s'(t-T_i)} + {N_b}^3 e^{-4s't} \\
& + 4{N^-_{\text{bn}}}^3 e^{-s'(4t-3T_i)} + 4N^-_{\text{bn}}{N_b}^2 e^{-s'(4t-T_i)} + 6{N^-_{\text{bn}}}^2 N_b e^{-s'(4t-2T_i)} - N^-_{\text{bn}}\left(3N_0^2 - 3N_b N_0 + {N_b}^2\right) e^{-4s'(t-T_i)}\Big)
\end{aligned} \tag{S64}
$$

$$
\begin{aligned}
g^{\text{BN}}_{D8R}(t) = e^{-4s't}\Big(& 4{N^-_{\text{bn}}}^3 e^{-s'(4-3T_i)} - 4{N^-_{\text{bn}}}^3 e^{-s'(4t-3(T_i+T_b))} - 6{N^-_{\text{bn}}}^3 e^{-s'(4t-(4T_i+2T_b))} - 12{N^-_{\text{bn}}}^3 e^{-s'(4t-(3T_i+T_b))} \\
& + 12{N^-_{\text{bn}}}^3 e^{-s'\left(4t-(3T_i+6s'T_b)\right)} + 4(N_0+N^-_{\text{bn}}){N^-_{\text{bn}}}^2 e^{-s'(4t-(4T_i+T_b))} + 6N_b{N^-_{\text{bn}}}^2 e^{-s'(4t-2T_i)} - 12N_b{N^-_{\text{bn}}}^2 e^{-s'(3t-2T_i)} \\
& + 6N_b{N^-_{\text{bn}}}^2 e^{-s'(4t-2(T_i+T_b))} - 12N_b{N^-_{\text{bn}}}^2 e^{-s'(4t-(2T_i+T_b))} + 24N_b{N^-_{\text{bn}}}^2 e^{-s'(3t-(2T_i+T_b))} - 12N_b{N^-_{\text{bn}}}^2 e^{-s'(3t-(3T_i+T_b))} \\
& - 12N_b{N^-_{\text{bn}}}^2 e^{-s'(3t-2(T_i+T_b))} + 4{N_b}^2 N^-_{\text{bn}} e^{-s'(4t-T_i)} - 12{N_b}^2 N^-_{\text{bn}} e^{-s'(3t-T_i)} - 6{N_b}^2 N^-_{\text{bn}} e^{-2s'(t-T_i)} \\
& + 12{N_b}^2 N^-_{\text{bn}} e^{-s'(2t-T_i)} - 4{N_b}^2 N^-_{\text{bn}} e^{-s'(4t-(T_i+T_b))} + 12{N_b}^2 N^-_{\text{bn}} e^{-s'(3t-(T_i+T_b))} + 6{N_b}^2 N^-_{\text{bn}} e^{-2s'(t-(T_i+T_b))} \\
& - 12{N_b}^2 N^-_{\text{bn}} e^{-s'(2t-(T_i+T_b))} + 4(N^-_{\text{bn}}-N_b)N_b N^-_{\text{bn}} e^{-3s'(t-(T_i+T_b))} - \left(3N_0^{N^-_{\text{bn}}} + {N_b}^2\right) N^-_{\text{bn}} e^{-4s'(t-T_i)} \\
& + \left(N_0^2 - 3N_b N^-_{\text{bn}}\right) N^-_{\text{bn}} e^{-4s'(t-(T_i+T_b))} + {N_b}^3\left(1 - 4e^{-s't} + 6e^{-2s't} - 4e^{-3s'} + e^{-4s'}\right) \\
& + 4N_b(N_0+N^-_{\text{bn}})N^-_{\text{bn}} e^{-3s'(t-T_i)}\Big)
\end{aligned} \tag{S65}
$$

**Oscillating size**

Note that the following expression is extremely tedious, but is shown here for completeness. Any application of this expression should use the corresponding result in Supplemental File SF1.

$$\kappa_{4D8R} = \mu \ \frac{4\times 3!\times 4!}{3!} \ \int_0^t dt_1 dt_2 dt_3 dt_4 \ \frac{G(t_2,t_1)G(t_3,t_2)G(t,t_2)G(t_4,t_3)G(t,t_3)G^2(t,t_4)}{4N_{\rm osc}(t_2)\times 4N_{\rm osc}(t_3)\times 4N_{\rm osc}(t_4)}$$
$$= 12\mu \ \int_0^t dt_1 dt_2 dt_3 dt_4 \ \frac{\Theta(t_2-t_1)\Theta(t_3-t_2)\Theta(t-t_2)\Theta(t_4-t_3)\Theta(t-t_3)\Theta^2(t-t_4) \ e^{-s'(4t-t_4-t_3-t_2-t_1)}}{2N_{\rm osc}(t_2)\times 2N_{\rm osc}(t_3)\times 2N_{\rm osc}(t_4)}$$
$$= \frac{\lambda'}{256{N_{\rm max}}^3{N_{\rm min}}^3 s'^3\left(s'^2\tau^2+\pi^2\right)\left(s'^2\tau^2+4\pi^2\right)^3\left(4s'^2\tau^2+\pi^2\right)\left(4s'^2\tau^2+9\pi^2\right)\left(9s'^2\tau^2+4\pi^2\right)\left(9s'^2\tau^2+16\pi^2\right)}\times$$
$$\Bigg[ A^{\rm osc}_{D8R}(t)+B^{\rm osc}_{D8R}(t)+C^{\rm osc}_{D8R}(t)+D^{\rm osc}_{D8R}(t)$$
$$+ e^{-4s't}\Bigg( 20736{N_{\rm min}}^3 s'^{16}\tau^{16} + 204192{N_{\rm min}}^3\pi^2 s'^{14}\tau^{14} + 163296N_{\rm max}{N_{\rm min}}^2\pi^2 s'^{14}\tau^{14} + 893920{N_{\rm min}}^3\pi^4 s'^{12}\tau^{12}$$
$$+ 1301832N_{\rm max}{N_{\rm min}}^2\pi^4 s'^{12}\tau^{12} + 464616{N_{\rm max}}^2N_{\rm min}\pi^4 s'^{12}\tau^{12} + 499041{N_{\rm max}}^3\pi^6 s'^{10}\tau^{10} + 2182049{N_{\rm min}}^3\pi^6 s'^{10}\tau^{10}$$
$$+ 4532103N_{\rm max}{N_{\rm min}}^2\pi^6 s'^{10}\tau^{10} + 2913543{N_{\rm max}}^2N_{\rm min}\pi^6 s'^{10}\tau^{10} + 2506068{N_{\rm max}}^3\pi^8 s'^8\tau^8 + 3225940{N_{\rm min}}^3\pi^8 s'^8\tau^8$$
$$+ 8471340N_{\rm max}{N_{\rm min}}^2\pi^8 s'^8\tau^8 + 7564332{N_{\rm max}}^2N_{\rm min}\pi^8 s'^8\tau^8 + 4391888{N_{\rm max}}^3\pi^{10} s'^6\tau^6 + 2895056{N_{\rm min}}^3\pi^{10} s'^6\tau^6$$
$$+ 8986416N_{\rm max}{N_{\rm min}}^2\pi^{10} s'^6\tau^6 + 10169904{N_{\rm max}}^2N_{\rm min}\pi^{10} s'^6\tau^6 + 3181504{N_{\rm max}}^3\pi^{12} s'^4\tau^4 + 1494976{N_{\rm min}}^3\pi^{12} s'^4\tau^4$$
$$+ 5473344N_{\rm max}{N_{\rm min}}^2\pi^{12} s'^4\tau^4 + 7061568{N_{\rm max}}^2N_{\rm min}\pi^{12} s'^4\tau^4 + 912896{N_{\rm max}}^3\pi^{14} s'^2\tau^2 + 470528{N_{\rm min}}^3\pi^{14} s'^2\tau^2$$
$$+ 1743360N_{\rm max}{N_{\rm min}}^2\pi^{14} s'^2\tau^2 + 2185728{N_{\rm max}}^2N_{\rm min}\pi^{14} s'^2\tau^2 + 73728\pi^{16}\left({N_{\rm max}}^3+{N_{\rm min}}^3\right)$$
$$+ 221184\left(N_{\rm max}{N_{\rm min}}^2\pi^{16}+{N_{\rm max}}^2N_{\rm min}\pi^{16}\right)\Bigg)\Bigg] \quad \text{(S66)}$$

The functions $A^{\rm osc}_{D8R}$, $B^{\rm osc}_{D8R}$, $C^{\rm osc}_{D8R}$, and $D^{\rm osc}_{D8R}$, defined below, are the oscillatory components of the above expression for $\kappa_{4D8R}$ in a cyclical population.

$$A^{\rm osc}_{D8R}(t) = -8e^{-s't}\left(s'^2\tau^2+\pi^2\right)\left(4s'^2\tau^2+\pi^2\right)\left(4s'^2\tau^2+9\pi^2\right)\left(9s'^2\tau^2+4\pi^2\right)\left(9s'^2\tau^2+16\pi^2\right)$$
$$\times\left(N^+_{\rm osc}\left(s'^2\tau^2+4\pi^2\right)-N^-_{\rm osc}s'\tau\left(s'\tau\cos[\varphi]+2\pi\sin[\varphi]\right)\right)^3 \quad \text{(S67)}$$

$$B^{\rm osc}_{D8R}(t) = 12e^{-2s't}\left(4s'^2\tau^2+\pi^2\right)\left(4s'^2\tau^2+9\pi^2\right)\left(9s'^2\tau^2+4\pi^2\right)\left(9s'^2\tau^2+16\pi^2\right)$$
$$\times\left(2N_{\rm min}s'^4\tau^4+(7N_{\rm max}+3N_{\rm min})\pi^2 s'^2\tau^2+4N^+_{\rm osc}\pi^4\right)\left(N^+_{\rm osc}\left(s'^2\tau^2+4\pi^2\right)-N^-_{\rm osc}s'\tau\left(s'\tau\cos[\varphi]+2\pi\sin[\varphi]\right)\right)^2$$
$$\text{(S68)}$$

$$C^{\rm osc}_{D8R}(t) =$$
$$-32e^{-3s't}\left(s'^2\tau^2+\pi^2\right)\left(4s'^2\tau^2+\pi^2\right)\left(4s'^2\tau^2+9\pi^2\right)\left(N^+_{\rm osc}\left(s'^2\tau^2+4\pi^2\right)-N^-_{\rm osc}s'\tau\left(s'\tau\cos[\varphi]+2\pi\sin[\varphi]\right)\right)$$
$$\times\Bigg(81{N_{\rm min}}^2 s'^8\tau^8+36N_{\rm min}(13N_{\rm max}+10N_{\rm min})\pi^2 s'^6\tau^6+4\left(195{N_{\rm max}}^2+322N_{\rm min}N_{\rm max}+183{N_{\rm min}}^2\right)\pi^4 s'^4\tau^4$$
$$+16\left(79{N_{\rm max}}^2+102N_{\rm min}N_{\rm max}+31{N_{\rm min}}^2\right)\pi^6 s'^2\tau^2+256{N^+_{\rm osc}}^2\pi^8\Bigg) \quad \text{(S69)}$$

$$D_{D8R}^{\rm osc}(t) = \left(s'^2\tau^2 + 4\pi^2\right)^3 \Bigg[ - N_{\rm osc}^- s'^2\tau^2 \cos[\varphi] \left(4s'^2\tau^2 + 9\pi^2\right) \Bigg(486 \left(5N_{\rm max}{}^2 + 6N_{\rm min}N_{\rm max} + 5N_{\rm min}{}^2\right) s'^6\tau^6$$
$$+ 9 \left(471N_{\rm max}{}^2 + 770N_{\rm min}N_{\rm max} + 471N_{\rm min}{}^2\right) \pi^2 s'^4\tau^4 + 4 \left(213N_{\rm max}{}^2 + 350N_{\rm min}N_{\rm max} + 213N_{\rm min}{}^2\right) \pi^4 s'^2\tau^2$$
$$+ 128N_{\rm osc}^+{}^2\pi^6\Bigg) + 3N_{\rm osc}^-{}^2 N_{\rm osc}^+ s'^2\tau^2 \cos[2\varphi](3s'\tau - 4\pi)(3s'\tau + 4\pi) \left(4s'^2\tau^2 + \pi^2\right) \left(4s'^2\tau^2 + 9\pi^2\right) \left(9s'^2\tau^2 + 2\pi^2\right)$$
$$+ 3N_{\rm osc}^-{}^3 s'^4\tau^4 \cos[3\varphi] \left(4s'^2\tau^2 + \pi^2\right) \left(9s'^2\tau^2 + 4\pi^2\right) \left(29\pi^2 - 6s'^2\tau^2\right)$$
$$- \pi N_{\rm osc}^- s'\tau \sin[\varphi] \left(4s'^2\tau^2 + 9\pi^2\right) \Bigg(256N_{\rm osc}^+{}^2\pi^6 + 621 \left(5N_{\rm max}{}^2 + 6N_{\rm min}N_{\rm max} + 5N_{\rm min}{}^2\right) s'^6\tau^6$$
$$+ 24 \left(245N_{\rm max}{}^2 + 414N_{\rm min}N_{\rm max} + 245N_{\rm min}{}^2\right) \pi^2 s'^4\tau^4 + 32 \left(63N_{\rm max}{}^2 + 116N_{\rm min}N_{\rm max} + 63N_{\rm min}{}^2\right) \pi^4 s'^2\tau^2\Bigg)$$
$$+ 3N_{\rm osc}^-{}^2 \pi s'^3\tau^3 \left(4s'^2\tau^2 + \pi^2\right) \Bigg(N_{\rm osc}^+ \sin[2\varphi] \left(4s'^2\tau^2 + 9\pi^2\right) \left(32\pi^2 + 207s'^2\tau^2\right)$$
$$+ N_{\rm osc}^- \sin[3\varphi] \left(48\pi^4 - 207s'^4\tau^4 + 16\pi^2 s'^2\tau^2\right)\Bigg) + N_{\rm osc}^+ \left(4s'^2\tau^2 + \pi^2\right) \left(4s'^2\tau^2 + 9\pi^2\right) \left(9s'^2\tau^2 + 16\pi^2\right) \Bigg(8N_{\rm osc}^+{}^2\pi^4$$
$$+ 9 \left(5N_{\rm max}{}^2 - 2N_{\rm min}N_{\rm max} + 5N_{\rm min}{}^2\right) s'^4\tau^4 + \left(33N_{\rm max}{}^2 + 38N_{\rm min}N_{\rm max} + 33N_{\rm min}{}^2\right) \pi^2 s'^2\tau^2\Bigg)\Bigg] \qquad \text{(S70)}$$

Note that, despite the difference in Heaviside functions, due to the different topologies of Diagrams D8 and D9, and an overall multiplicative factor, the following integrals have the same exponentiated time dependence and the same temporal coordinates in the time-dependent population size. The two left (D8L and D9L) and two right (D8R and D9R) diagrams also have identical vertices, but distinct topologies, as discussed in the main text. As a result, after integration, Diagrams D8 and D9 have identical contributions to the fourth cumulant, up to differing overall constants.

##### S3.4.3 Diagram 9 (left)

**Constant size**

$$\begin{aligned}
\kappa_{4D9L} &= \frac{p_0}{(4N)^3} \; \frac{3! \times 4!}{3!} \int_0^t dt_1 dt_2 dt_3 \; G(t_1,0)G(t_2,t_1)G(t_3,t_1)G^2(t,t_2)G^2(t,t_3) \\
&= \frac{3p_0}{(2N)^3} \int_0^t dt_1 dt_2 dt_3 \; \Theta(t_1)\Theta(t_2-t_1)\Theta(t_3-t_1)\Theta^2(t-t_2)\Theta^2(t-t_3) \; e^{-s'(4t-t_3-t_2-t_1)} \\
&= \frac{p_0 e^{-s't}}{\gamma'^3} \left(1 - e^{-s't}\right)^3 \qquad \text{(S71)}
\end{aligned}$$

**Exponential growth**

$$\begin{aligned}
\kappa_{4D9L} &= p_0 \; \frac{3! \times 4!}{3!} \int_0^t dt_1 dt_2 dt_3 \; \frac{G(t_1,0)G(t_2,t_1)G(t_3,t_1)G^2(t,t_2)G^2(t,t_3)}{4N_{\rm exp}(t_1) \times 4N_{\rm exp}(t_2) \times 4N_{\rm exp}(t_3)} \\
&= 3p_0 \int_0^t dt_1 dt_2 dt_3 \; \frac{\Theta(t_1)\Theta(t_2-t_1)\Theta(t_3-t_1)\Theta^2(t-t_2)\Theta^2(t-t_3) \; e^{-s'(4t-t_3-t_2-t_1)}}{2N_{\rm exp}(t_1) \times 2N_{\rm exp}(t_2) \times 2N_{\rm exp}(t_3)} \\
&= \frac{p_0}{\gamma'^3_{\rm exp}} e^{-s't} \left(e^{-\Gamma t} - e^{-s't}\right)^3
\end{aligned}$$

(S72)

**Bottleneck**

$$
\begin{aligned}
\kappa_{4D9L} &= p_0 \ \frac{3! \times 4!}{3!} \int_0^t dt_1 dt_2 dt_3 \ \frac{G(t_1,0)G(t_2,t_1)G(t_3,t_1)G^2(t,t_2)G^2(t,t_3)}{4N_{\mathrm{BN}}(t_1) \times 4N_{\mathrm{BN}}(t_2) \times 4N_{\mathrm{BN}}(t_3)} \\
&= 3p_0 \int_0^t dt_1 dt_2 dt_3 \ \frac{\Theta(t_1)\Theta(t_2-t_1)\Theta(t_3-t_1)\Theta^2(t-t_2)\Theta^2(t-t_3) \ e^{-s'(4t-t_3-t_2-t_1)}}{2N_{\mathrm{BN}}(t_1) \times 2N_{\mathrm{BN}}(t_2) \times 2N_{\mathrm{BN}}(t_3)} \\
&= \ \frac{1}{2} \times \ \text{Result of Diagram D8}L
\end{aligned} \tag{S73}
$$

**Oscillating size**

$$
\begin{aligned}
\kappa_{4D9L} &= p_0 \ \frac{3! \times 4!}{3!} \int_0^t dt_1 dt_2 dt_3 \ \frac{G(t_1,0)G(t_2,t_1)G(t_3,t_1)G^2(t,t_2)G^2(t,t_3)}{4N_{\mathrm{osc}}(t_1) \times 4N_{\mathrm{osc}}(t_2) \times 4N_{\mathrm{osc}}(t_3)} \\
&= 3p_0 \int_0^t dt_1 dt_2 dt_3 \ \frac{\Theta(t_1)\Theta(t_2-t_1)\Theta(t_3-t_1)\Theta^2(t-t_2)\Theta^2(t-t_3) \ e^{-s'(4t-t_3-t_2-t_1)}}{2N_{\mathrm{osc}}(t_1) \times 2N_{\mathrm{osc}}(t_2) \times 2N_{\mathrm{osc}}(t_3)} \\
&= \ \frac{1}{2} \times \ \text{Result of Diagram D8}L
\end{aligned} \tag{S74}
$$

##### S3.4.4 Diagram 9 (right)

**Constant size**

$$
\begin{aligned}
\kappa_{4D9R} &= \frac{\mu}{(4N)^3} \ \frac{3! \times 4!}{3!} \int_0^t dt_1 dt_2 dt_3 dt_4 \ G(t_2,t_1)G(t_3,t_2)G(t_4,t_2)G^2(t,t_3)G^2(t,t_4) \\
&= \frac{3p_0}{(2N)^3} \int_0^t dt_1 dt_2 dt_3 dt_4 \ \Theta(t_2-t_1)\Theta(t_3-t_2)\Theta(t_4-t_2)\Theta^2(t-t_3)\Theta^2(t-t_4) \ e^{-s'(4t-t_4-t_3-t_2-t_1)} \\
&= \frac{\lambda'}{4\gamma'^3}\left(1-e^{-s't}\right)^4
\end{aligned} \tag{S75}
$$

**Exponential growth**

$$
\begin{aligned}
\kappa_{4D9R} &= \mu \ \frac{3! \times 4!}{3!} \int_0^t dt_1 dt_2 dt_3 dt_4 \ \frac{G(t_2,t_1)G(t_3,t_2)G(t_4,t_2)G^2(t,t_3)G^2(t,t_4)}{4N_{\mathrm{exp}}(t_2) \times 4N_{\mathrm{exp}}(t_3) \times 4N_{\mathrm{exp}}(t_4)} \\
&= 3\mu \int_0^t dt_1 dt_2 dt_3 dt_4 \ \frac{\Theta(t_2-t_1)\Theta(t_3-t_2)\Theta(t_4-t_2)\Theta^2(t-t_3)\Theta^2(t-t_4) \ e^{-s'(4t-t_4-t_3-t_2-t_1)}}{2N_{\mathrm{exp}}(t_2) \times 2N_{\mathrm{exp}}(t_3) \times 2N_{\mathrm{exp}}(t_4)} \\
&= \frac{\lambda'}{\gamma'^3_{\mathrm{exp}}}\left(e^{-3\Gamma t}\left(1-e^{-s't}\right) - \frac{3s'\left(e^{-3\Gamma t}-e^{-2(s'+\Gamma)t}\right)}{2s'-\Gamma} + \frac{3s'\left(e^{-3\Gamma t}-e^{-2(s'+\Gamma)t}\right)}{3s'-2\Gamma} - \frac{s'\left(e^{-3\Gamma t}-e^{-4s't}\right)}{4s'-3\Gamma}\right)
\end{aligned} \tag{S76}
$$

**Bottleneck**

$$
\begin{aligned}
\kappa_{4D9R} &= \mu \ \frac{3! \times 4!}{3!} \int_0^t dt_1 dt_2 dt_3 dt_4 \ \frac{G(t_2,t_1)G(t_3,t_2)G(t_4,t_2)G^2(t,t_3)G^2(t,t_4)}{4N_{\mathrm{BN}}(t_2) \times 4N_{\mathrm{BN}}(t_3) \times 4N_{\mathrm{BN}}(t_4)} \\
&= 3\mu \int_0^t dt_1 dt_2 dt_3 dt_4 \ \frac{\Theta(t_2-t_1)\Theta(t_3-t_2)\Theta(t_4-t_2)\Theta^2(t-t_3)\Theta^2(t-t_4) \ e^{-s'(4t-t_4-t_3-t_2-t_1)}}{2N_{\mathrm{BN}}(t_2) \times 2N_{\mathrm{BN}}(t_3) \times 2N_{\mathrm{BN}}(t_4)} \\
&= \ \frac{1}{2} \times \ \text{Result of Diagram D8}R
\end{aligned} \tag{S77}
$$

**Oscillating size**

$$\begin{aligned}\kappa_{4D9R} &= \mu\ \frac{3!\times 4!}{3!}\ \int_0^t dt_1 dt_2 dt_3 dt_4\ \frac{G(t_2,t_1)G(t_3,t_2)G(t_4,t_2)G^2(t,t_3)G^2(t,t_4)}{4N_{\rm osc}(t_2)\times 4N_{\rm osc}(t_3)\times 4N_{\rm osc}(t_4)}\\ &= 3\mu\ \int_0^t dt_1 dt_2 dt_3 dt_4\ \frac{\Theta(t_2-t_1)\Theta(t_3-t_2)\Theta(t_4-t_2)\Theta^2(t-t_3)\Theta^2(t-t_4)\ e^{-s'(4t-t_4-t_3-t_2-t_1)}}{2N_{\rm osc}(t_2)\times 2N_{\rm osc}(t_3)\times 2N_{\rm osc}(t_4)}\\ &= \ \frac{1}{2}\times\ \text{Result of Diagram D8}R \end{aligned} \tag{S78}$$

### S4 Comparison of analytic approximations to simulations over a range of parameters

Throughout this section, which compares the PGFT-based approximations discussed in the main text to `neqPopDynx` simulations, solid curves represent analytic approximations for the quantity of interest (e.g., third moment, second cumulant) and simulations of the same parameters are shown as a collection of time-dependent points in a darker shade of the same color used for the analytic curve. Curves with the strongest selection ($s = 10^{-1/2}$) are shown in red and those for the weakest selection coefficient ($s = 10^{-5}$) are shown in purple. Additionally, note that, in all cases, the strongest selection curves deviate slightly from the simulated results because $s = 10^{-1/2} \approx 0.316$ is large enough that the linear approximation of fitness in the selection coefficient becomes inaccurate (i.e., $1 - e^{-s} = 0.271 < 0.316 = s$). In comparison, even at $s = 0.1$, the linearized approximation is quite accurate: $1 - e^{-s} \approx 0.095$. I have left the corresponding curves in their linearized form with the intention of plotting the functional form of the relevant PGFT approximations as expressed in the manuscript. If, instead, the dependence on $s$ is replaced with $1 - e^{-s}$, that analytic approximations in the extremely strong selection regime ($s = 10^{-1/2}$) are closer to the simulated results in all parameter regimes explored in this analysis (i.e., for any moment in any demography with any combination of demographic parameters).

For each demography, all plots are presented on a log-log scale (in base ten) to allow for easier visualization of dynamics that span many orders of magnitude; the simulations are roughly evenly spaced in generation time such that the density of points is greater at later values due to the logarithmic generation time axis. In presenting plots on a logarithmic scale, the accuracy or inaccuracy of an analytic expression can be judged with respect to the order (or half order) of magnitude due to the logarithmic spacing, rather than with respect to small deviations, which would be difficult to observe due to the stochastic noise in simulation results. Each simulated point plotted is the result of computing expectation values over $L = 10^5$ independent sites to characterize cumulants and moments of the allele frequency probability distribution (e.g., $E[p] = \frac{1}{L}\sum_{i=1}^{L} p_i$).

#### S4.1 Constant population size

For a population with a constant size of $2N$ haploid individuals, two parameters are varied in this section: the population size $N$, and the mutation rate $\mu$ (while maintaining $\mu_b = \mu/10^2$ except where otherwise specified). For each parameter combination of ($N$, $\mu$), the same range of selection coefficients will be presented, $s \in [10^{-5}, 10^{-1/2}]$, but coefficients near the value where the strong selection approximation breaks down (at roughly $2Ns' \sim 5$) will be chosen appropriately for the corresponding population size such that $2Ns \in \{3, 5, 10\}$. Finally, the initial frequency $p_0$ will be varied while fixing fixing all other parameters ($N = 10^4$, $\mu = 10^{-8}$, $\mu_b = 10^{-10}$ for a range of selection coefficients) to investigate the regime of validity of the approximations detailed in the main text; a breakdown is expected at some $p_0$, but the numerical value is unclear from the perturbation series of the partition functional (but see discussion of the equilibration time for constant $N$, where $p_0 > \lambda'$ for moments above the mean frequency). All moments will be shown for each parameter choice for comparison. Simulations with lower mutation rates ($\mu = 10^{-8}$) appear to be more noisy at asymptotically late times (i.e., after equilibration); both the variance and mean of each moment are linear in the mutation rate such that the ratio of the standard deviation to the mean is proportional to $\mu^{-1/2}$.

##### S4.1.1 Constant demography: time dependence of the population size

For a population of constant size, the time dependence is trivial. Analytic and simulation results in the following figures are shown in the same order as the population size plots below, from top to bottom: $N = 10^4$, $10^3$, and $10^2$. Note that the left and right columns in Figure S1 are identical; each row shows results for the same demography but with distinct mutation rates $\mu = 10^{-6}$ (left column) and $\mu = 10^{-8}$ (right column). Back mutation rates are set to a fixed fraction of the mutation rate $\mu_b = \mu/100$ in all cases.

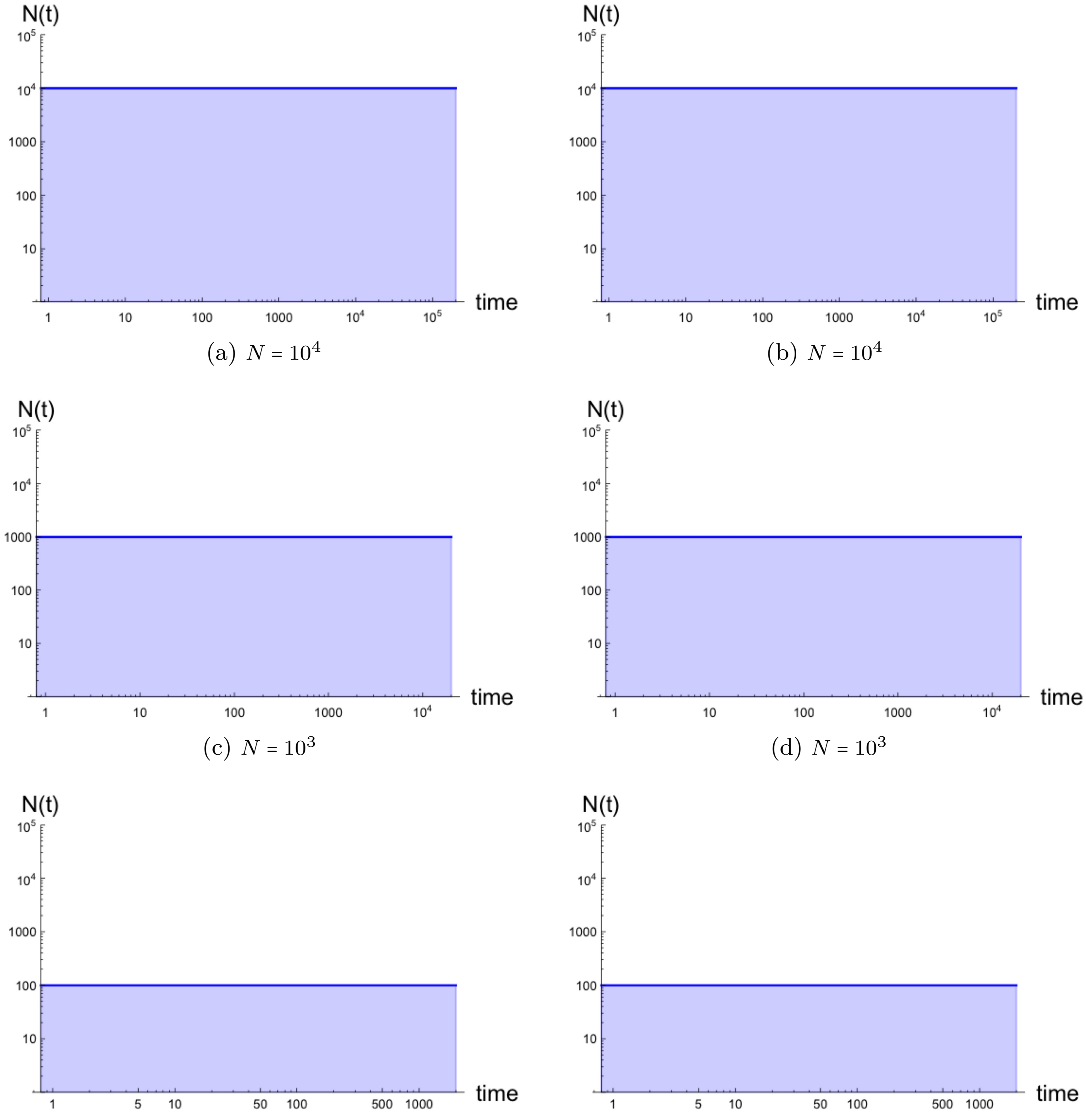


(a) $N = 10^4$ (b) $N = 10^4$

(c) $N = 10^3$ (d) $N = 10^3$

(e) $N = 10^2$ (f) $N = 10^2$

Figure S1: *Time dependence of the population size for simulated populations with constant size.* Demographies are presented with $N = 10^4$ (a, b), $10^3$ (c,d), $10^2$ (e,f). Left and right plots are identical here, but will correspond to distinct mutation rates in subsequent figures for constant population size.

##### S4.1.2 Constant demography: mean frequency $\overline{p}(t)$ to one-vertex order

The lowest order approximations to the mean frequency $\overline{p}(t) \equiv E[p(t)]$, the first cumulant and first moment of the allele frequency probability distribution, are plotted below for simulations with various population size and mutation rate combinations. These solutions are equivalent to the solution to the deterministic equation for allele frequency as a function of the initial frequency $p_0$, the mutation rates $\mu$ and $\mu_b$, and the (purifying) selection coefficient $s$. Each plot shows the dynamics of a range of selection coefficients, with a higher density of selection coefficients near the breakdown at roughly $s$ $5/2N$. The strong selection approximation breaks down at larger selection coefficients for smaller population size. Reduction of the mutation rates (right column) results in more variance of the mean, but the analytic approximation presented remains equally valid. For smaller population size, simulations become discretized under very strong selection when approaching mutation-selection balance at late times.

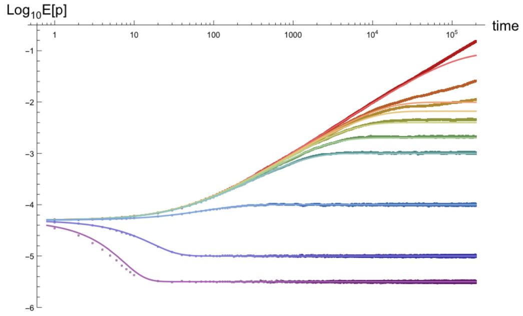


(a) $N = 10^4, \mu = 10^{-6}, \mu_b = 10^{-8}$

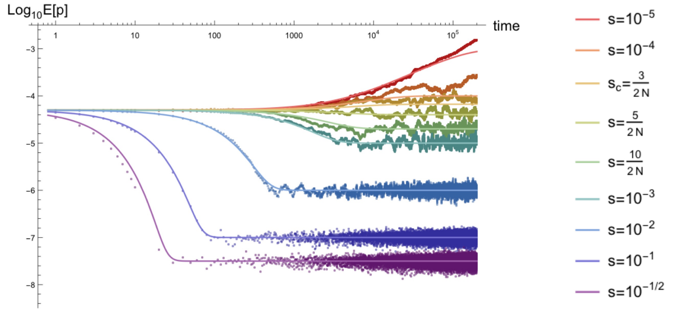


(b) $N = 10^4, \mu = 10^{-8}, \mu_b = 10^{-10}$

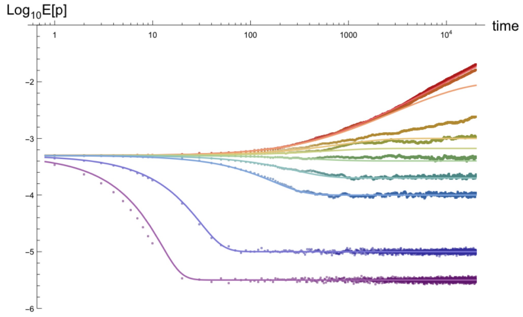


(c) $N = 10^3, \mu = 10^{-6}, \mu_b = 10^{-8}$

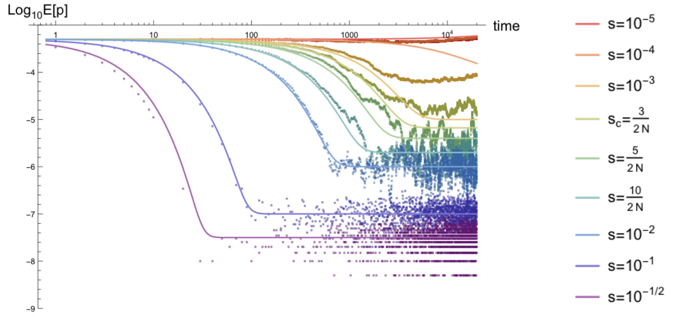


(d) $N = 10^3, \mu = 10^{-8}, \mu_b = 10^{-10}$

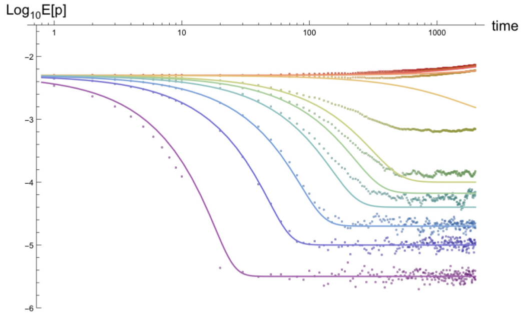


(e) $N = 10^2, \mu = 10^{-6}, \mu_b = 10^{-8}$

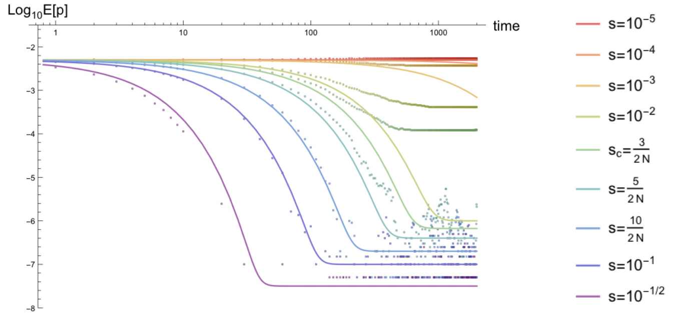


(f) $N = 10^2, \mu = 10^{-8}, \mu_b = 10^{-10}$

Figure S2: *Lowest order (one-vertex) approximation for the mean frequency $\overline{p}(t)$ dynamics for constant population size.* Plots are shown for mutation rates $\mu = 10^{-6}$ (left), $10^{-8}$ (right) and $\mu_b = \mu/100$, population size $N = 10^4$ (top), $10^3$ (middle), $10^2$ (bottom), and selection coefficients $s \in [10^{-5}, 10^{-1/2}]$.

##### S4.1.3 Constant demography: mean frequency $\overline{p}(t)$ to three-vertex order

The lowest-vertex order (3V) corrections to the one-vertex approximation of the mean frequency $\overline{p}(t)$ are shown below. Note that the corrections are very small in comparison to the leading order expression due to suppression by an additional factor of $p_0^2$, $\lambda'^2 = \mu^2/s'^2$, $1/\gamma' = 1/(2Ns')^2$, or some combination of these perturbative parameters. The effect of these corrections are clearest in the plots for the smallest population $N = 10^2$, where there is less deviation from simulated values in the curves with $2Ns = 3, 5, 10$ than seen in the analogous one-vertex mean plots above.

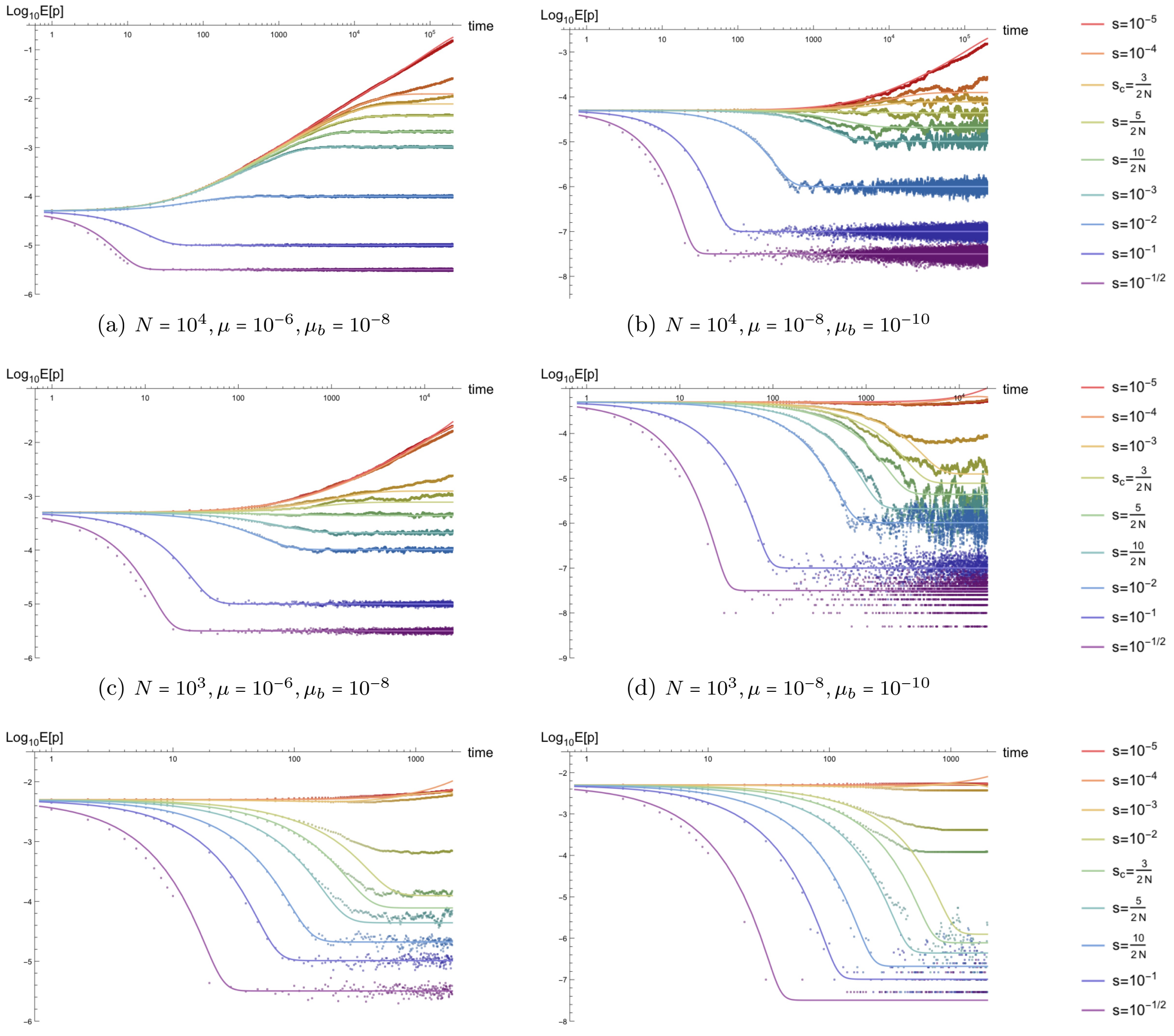


(a) $N = 10^4, \mu = 10^{-6}, \mu_b = 10^{-8}$ (b) $N = 10^4, \mu = 10^{-8}, \mu_b = 10^{-10}$

(c) $N = 10^3, \mu = 10^{-6}, \mu_b = 10^{-8}$ (d) $N = 10^3, \mu = 10^{-8}, \mu_b = 10^{-10}$

(e) $N = 10^2, \mu = 10^{-6}, \mu_b = 10^{-8}$ (f) $N = 10^2, \mu = 10^{-8}, \mu_b = 10^{-10}$

Figure S3: *Time dependence of the three-vertex approximations for the mean frequency $\overline{p}(t)$ for constant population size.* These plots incorporate the lowest-order non-vanishing corrections to the one-vertex approximations, which are first order in $p_0, \lambda', 1/\gamma'$. Parameter combinations correspond to those shown in Figure S2.

##### S4.1.4 Constant demography: variance $\kappa_2(t)$ to two-vertex order

Plots of the lowest order (two-vertex) approximation to the second cumulant, the variance $\kappa_2(t)$ of the allele frequency probability distribution, are presented here for the same range of parameters as above. The variance (computed from simulations as $E[(p-\overline{p})^2]$) is very close to the homozygosity $E[p^2]$ (shown below), since the square of the mean frequency only meaningfully alters the time-dependence when the variance is very small. The variance is initially zero at the initial delta function (unable to be seen on log-log plots) and equilibrates to values $\lambda'/2\gamma'$.

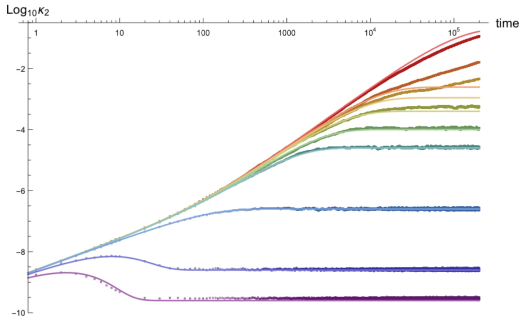


(a) $N = 10^4, \mu = 10^{-6}, \mu_b = 10^{-8}$

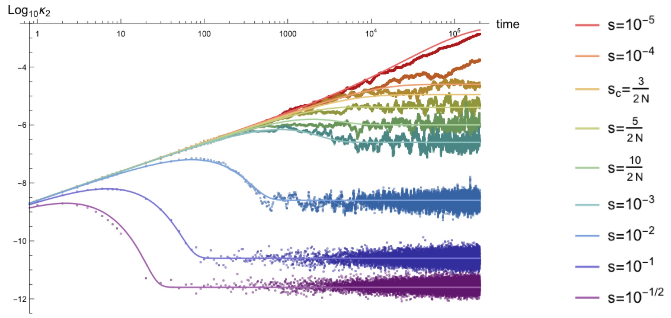


(b) $N = 10^4, \mu = 10^{-8}, \mu_b = 10^{-10}$

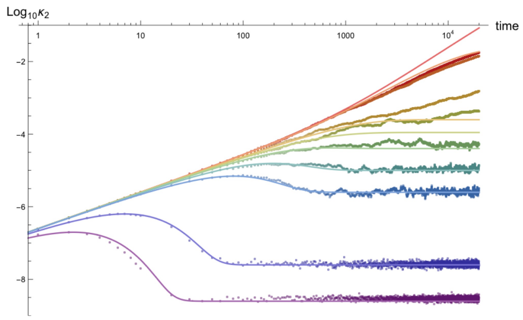


(c) $N = 10^3, \mu = 10^{-6}, \mu_b = 10^{-8}$

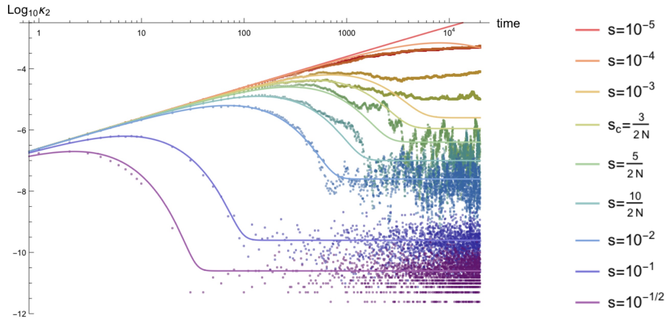


(d) $N = 10^3, \mu = 10^{-8}, \mu_b = 10^{-10}$

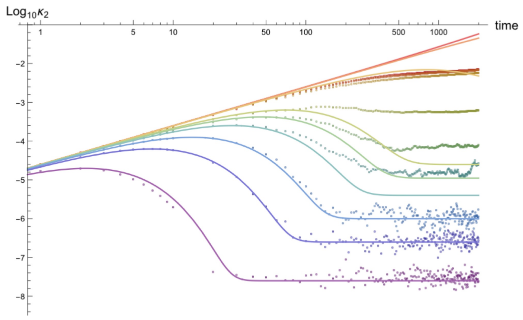


(e) $N = 10^2, \mu = 10^{-6}, \mu_b = 10^{-8}$

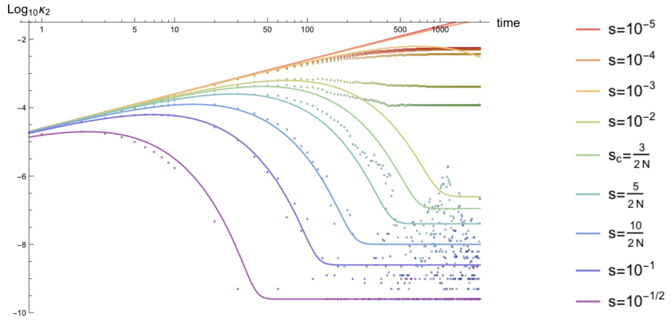


(f) $N = 10^2, \mu = 10^{-8}, \mu_b = 10^{-10}$

Figure S4: *Time dependence of the lowest-vertex order approximations for the variance of the allele frequency* $\kappa_2(t)$ *for constant population size.*

##### S4.1.5 Constant demography: homozygosity $E[p^2(t)]$ to two-vertex order

The lowest-vertex order (2V) approximation to the homozygosity is shown below using the same parameters as above. The homozygosity, computed as $\kappa_2(t) - (E[p(t)])^2$, is initially $p_0^2$ due to the delta function initial condition and equilibrates to roughly the same asymptotic value as the variance for the parameter combinations presented below.

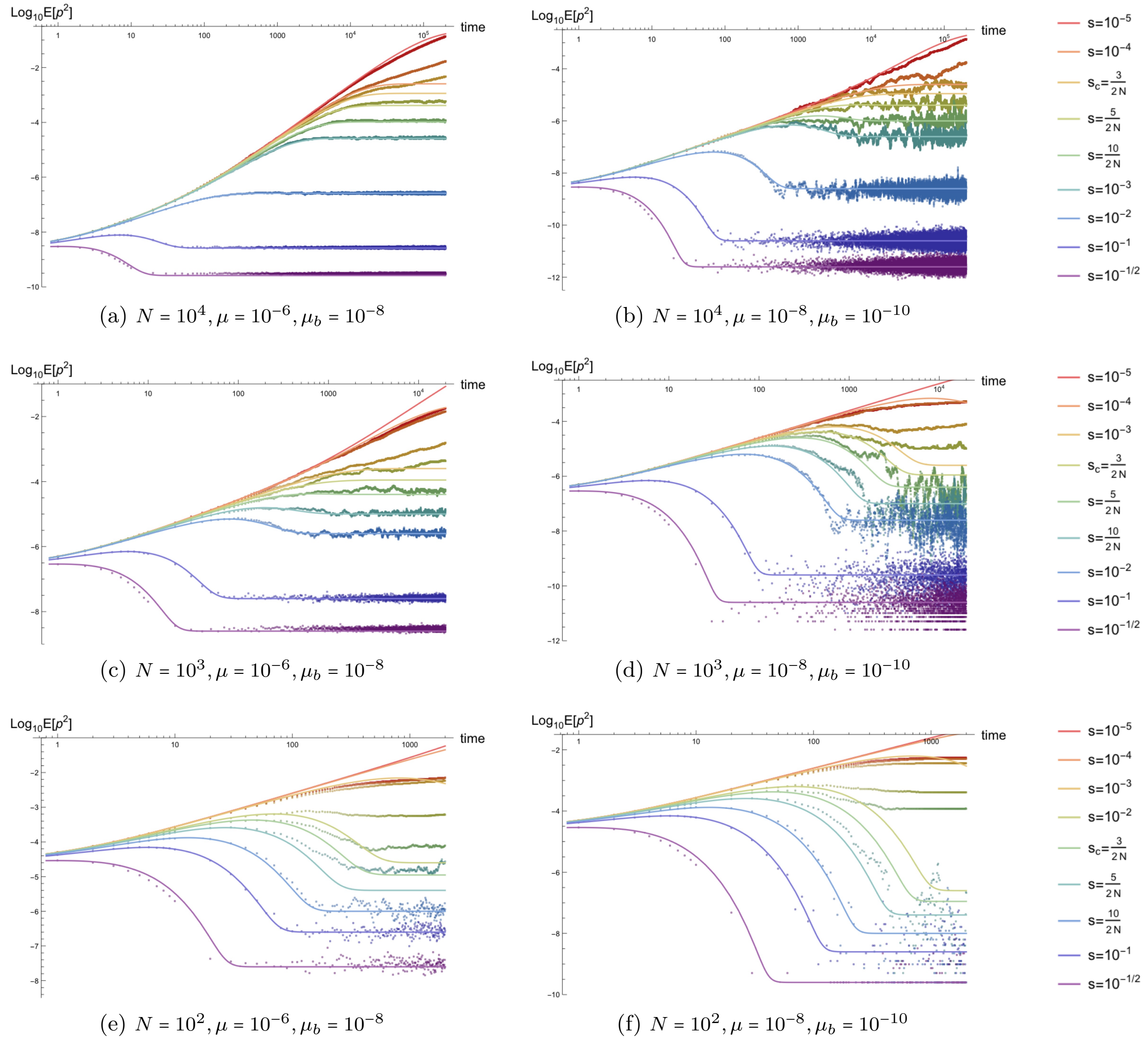


(a) $N = 10^4, \mu = 10^{-6}, \mu_b = 10^{-8}$ (b) $N = 10^4, \mu = 10^{-8}, \mu_b = 10^{-10}$

(c) $N = 10^3, \mu = 10^{-6}, \mu_b = 10^{-8}$ (d) $N = 10^3, \mu = 10^{-8}, \mu_b = 10^{-10}$

(e) $N = 10^2, \mu = 10^{-6}, \mu_b = 10^{-8}$ (f) $N = 10^2, \mu = 10^{-8}, \mu_b = 10^{-10}$

Figure S5: *Time dependence of the lowest-vertex order approximations for the homozygosity $E[p^2(t)]$ for constant population size.* Parameter combinations correspond to those shown in Figure S2.

##### S4.1.6 Constant demography: homozygosity $E[p^2(t)]$ to three-vertex order

The lowest-vertex order (3V) corrections to the two-vertex approximation to the homozygosity is shown below using the same parameters as above. The corrections are again small in comparison to the leading order terms, but become meaningfully closer to the simulation results when the population size is small. Importantly, these corrections fail catastrophically when the perturbative parameters (most relevantly, $\lambda'$ and $1/\gamma'$) become greater than one (see curves for the weakest selection coefficients below). This occurs because the three-vertex terms, which are strictly negative, grow larger than the two-vertex terms; this can be controlled by adding higher-vertex corrections, which overwhelm the three-vertex terms, but the approximation remains invalid in this regime unless an infinite series of corrections are computed.

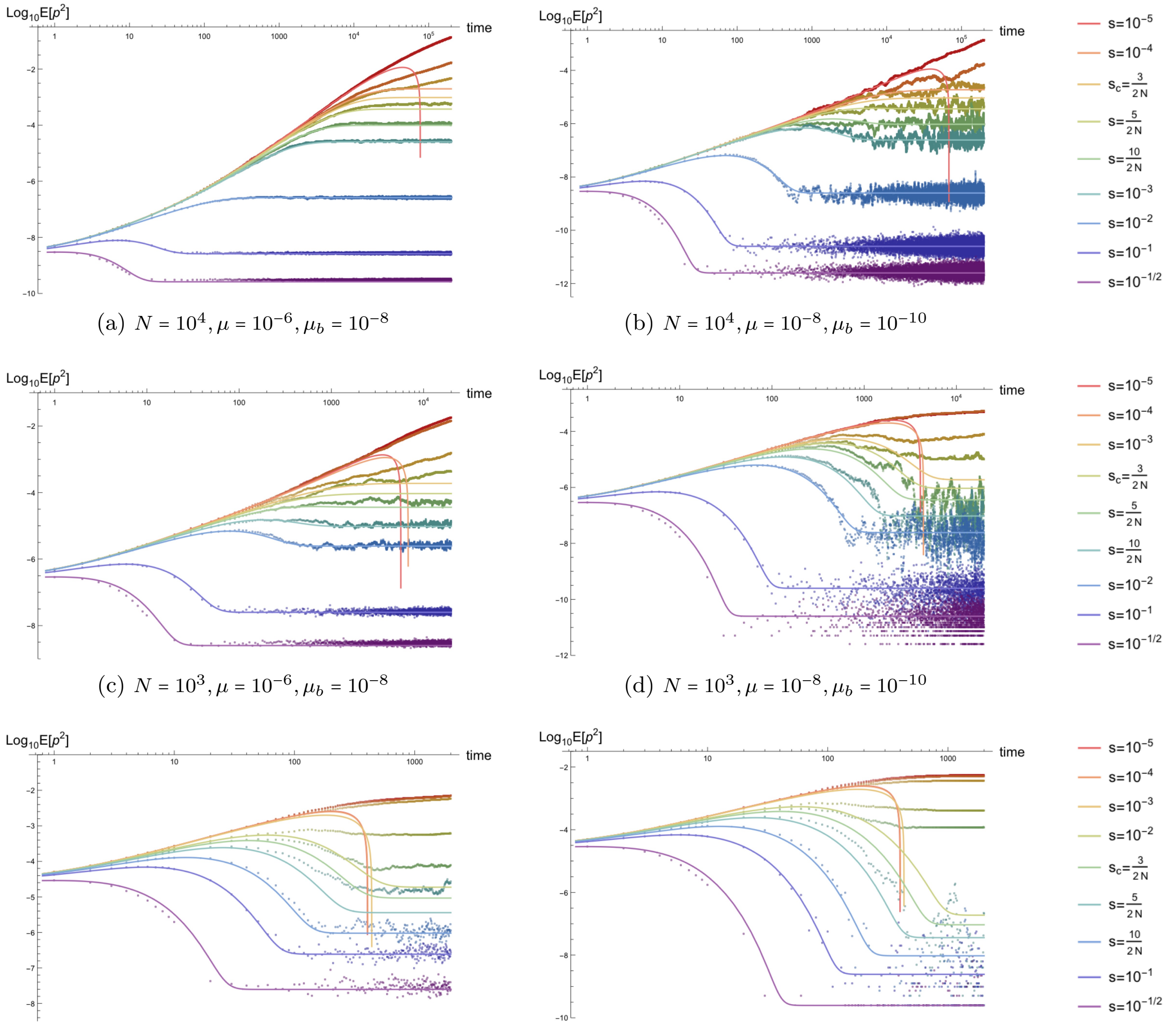


(a) $N = 10^4, \mu = 10^{-6}, \mu_b = 10^{-8}$

(b) $N = 10^4, \mu = 10^{-8}, \mu_b = 10^{-10}$

(c) $N = 10^3, \mu = 10^{-6}, \mu_b = 10^{-8}$

(d) $N = 10^3, \mu = 10^{-8}, \mu_b = 10^{-10}$

(e) $N = 10^2, \mu = 10^{-6}, \mu_b = 10^{-8}$

(f) $N = 10^2, \mu = 10^{-8}, \mu_b = 10^{-10}$

Figure S6: *Time dependence of the three-vertex approximations for the homozygosity $E[p^2(t)]$ for constant population size.* Parameter combinations correspond to those shown in Figure S2.

##### S4.1.7 Constant demography: third cumulant $\kappa_3(t)$ to three-vertex order

Below are plots of the lowest-vertex (3V) approximation to the time-dependence of the third cumulant of the allele frequency probability distribution $\kappa_3(t)$, the non-standardized skew, for the same parameters.

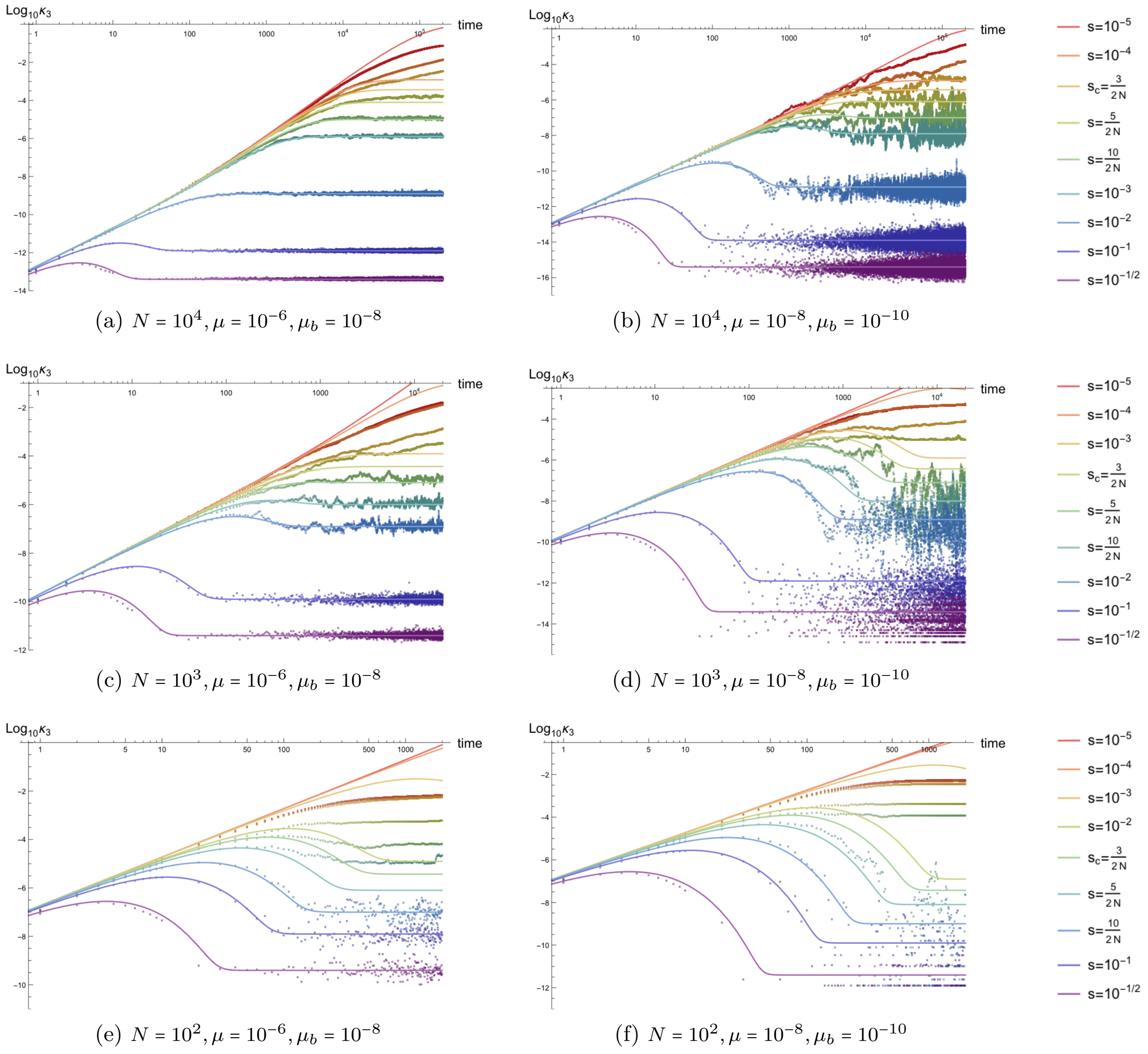


(a) $N = 10^4, \mu = 10^{-6}, \mu_b = 10^{-8}$

(b) $N = 10^4, \mu = 10^{-8}, \mu_b = 10^{-10}$

(c) $N = 10^3, \mu = 10^{-6}, \mu_b = 10^{-8}$

(d) $N = 10^3, \mu = 10^{-8}, \mu_b = 10^{-10}$

(e) $N = 10^2, \mu = 10^{-6}, \mu_b = 10^{-8}$

(f) $N = 10^2, \mu = 10^{-8}, \mu_b = 10^{-10}$

Figure S7: *Time dependence of the three-vertex approximations for the third cumulant $\kappa_3(t)$ for constant population size.* Parameter combinations correspond to those shown in Figure S2.

##### S4.1.8 Constant demography: third moment $E[p^3(t)]$ to three-vertex order

Below are plots of the lowest-vertex (3V) approximation to the time-dependence of the third non-central moment $E[p^3(t)]$ for the same parameters. Like the homozygosity, the third moment is very similar to the third cumulant other than at the initial time when it has a value of $p_0^3$ from the delta function initial condition.

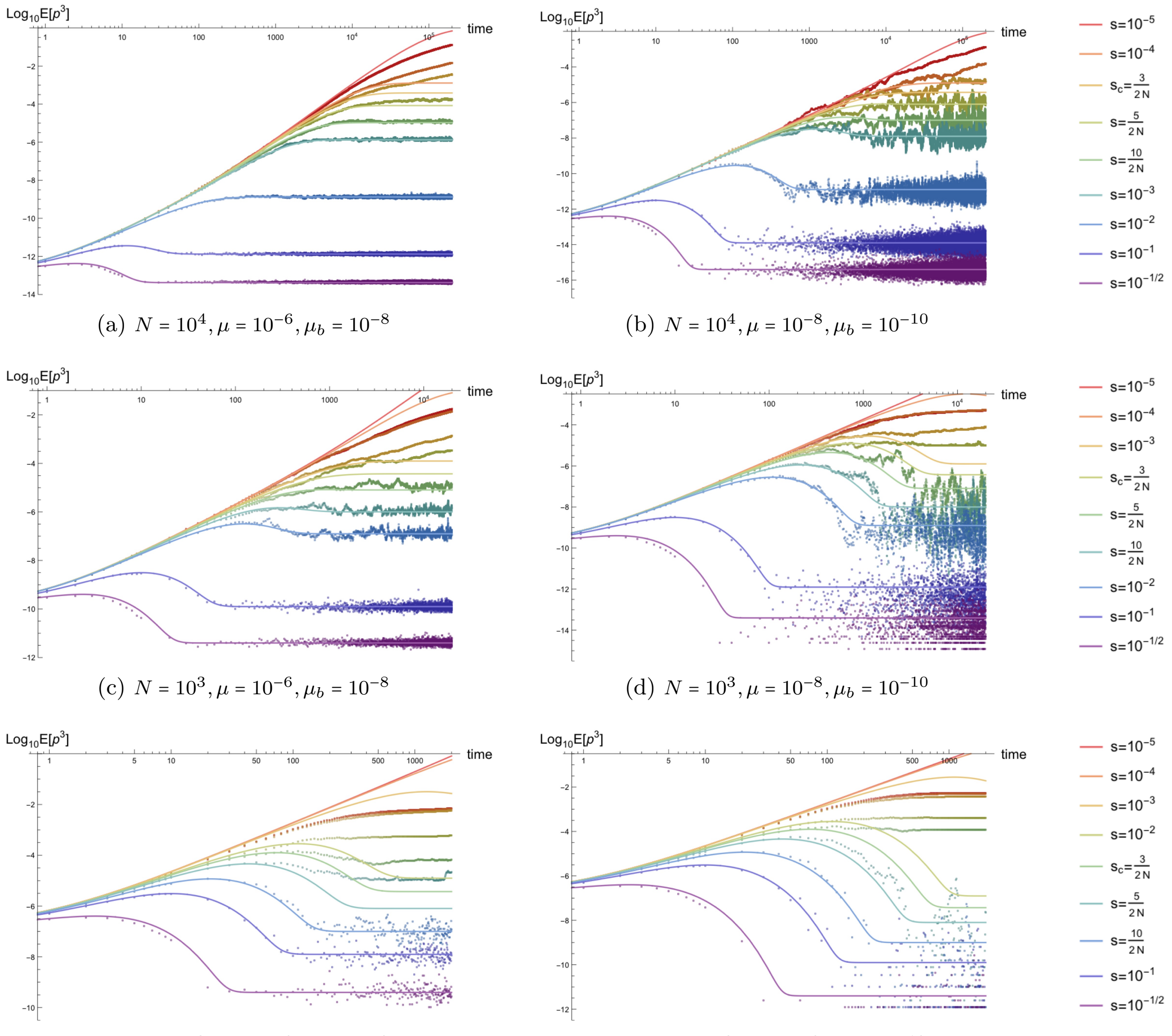


(a) $N = 10^4, \mu = 10^{-6}, \mu_b = 10^{-8}$

(b) $N = 10^4, \mu = 10^{-8}, \mu_b = 10^{-10}$

(c) $N = 10^3, \mu = 10^{-6}, \mu_b = 10^{-8}$

(d) $N = 10^3, \mu = 10^{-8}, \mu_b = 10^{-10}$

(e) $N = 10^2, \mu = 10^{-6}, \mu_b = 10^{-8}$

(f) $N = 10^2, \mu = 10^{-8}, \mu_b = 10^{-10}$

Figure S8: *Time dependence of the three-vertex approximations for the third non-central moment $E[p^3(t)]$ for constant population size.* Parameter combinations correspond to those shown in Figure S2.

##### S4.1.9 Constant demography: skew $K_3(t)$ to three-vertex order

Here, the time dependence of the lowest-vertex (3V) order approximation to the skew of the allele frequency probability distribution $K_3(t)$ is plotted. Note that, when the approximation describes the dynamics (for $2Ns > 5$), the skew is independent of the strength of selection at asymptotically late times. The equilibrium value $1/\sqrt{\lambda'\gamma'} = 1/\sqrt{\mu N}$ is dependent only on the population scaled mutation rate $\theta = 4N\mu$ (compare the left to right columns for each row below). Note that, for low mutation rate and very small population size, the skew fails to match analytic expectations due to discretization of the third cumulant, which can be seen in the Figures 9D and 9F below.

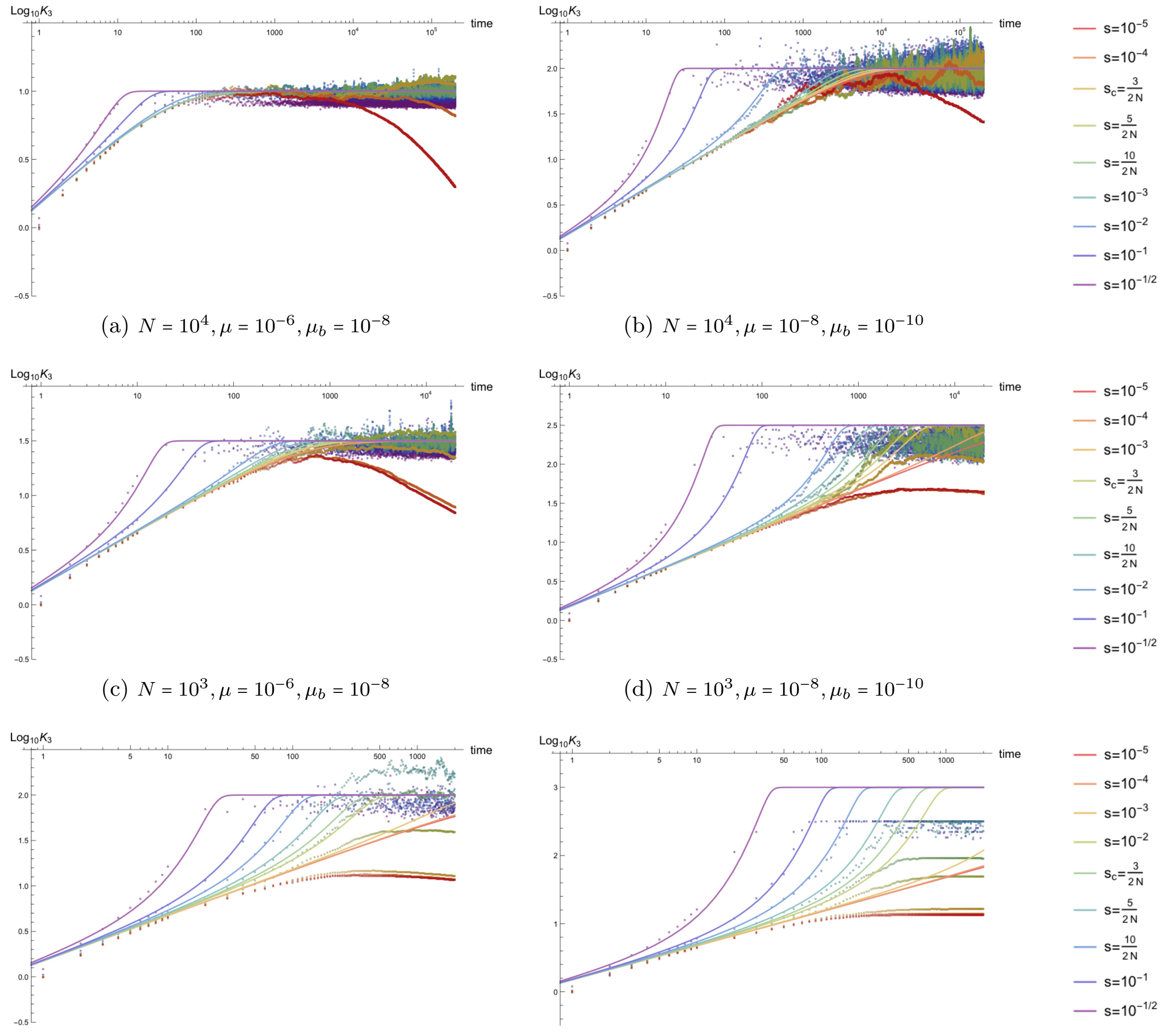


(a) $N = 10^4, \mu = 10^{-6}, \mu_b = 10^{-8}$

(b) $N = 10^4, \mu = 10^{-8}, \mu_b = 10^{-10}$

(c) $N = 10^3, \mu = 10^{-6}, \mu_b = 10^{-8}$

(d) $N = 10^3, \mu = 10^{-8}, \mu_b = 10^{-10}$

(e) $N = 10^2, \mu = 10^{-6}, \mu_b = 10^{-8}$

(f) $N = 10^2, \mu = 10^{-8}, \mu_b = 10^{-10}$

Figure S9: *Time dependence of the three-vertex approximations for the skew of the allele frequency probability distribution $K_3(t)$ in populations with constant size.* Parameter combinations correspond to those shown in Figure S2.

##### S4.1.10 Constant demography: fourth cumulant $\kappa_4(t)$ to four-vertex order

Below are plots of the lowest-vertex (4V) approximation to the time-dependence of the fourth cumulant of the allele frequency probability distribution $\kappa_4(t)$, the non-standardized excess kurtosis, for the same parameters. This is the first cumulant that differs from the central moment of the same order (i.e., the non-standardized kurtosis); all higher cumulants differ, as well.

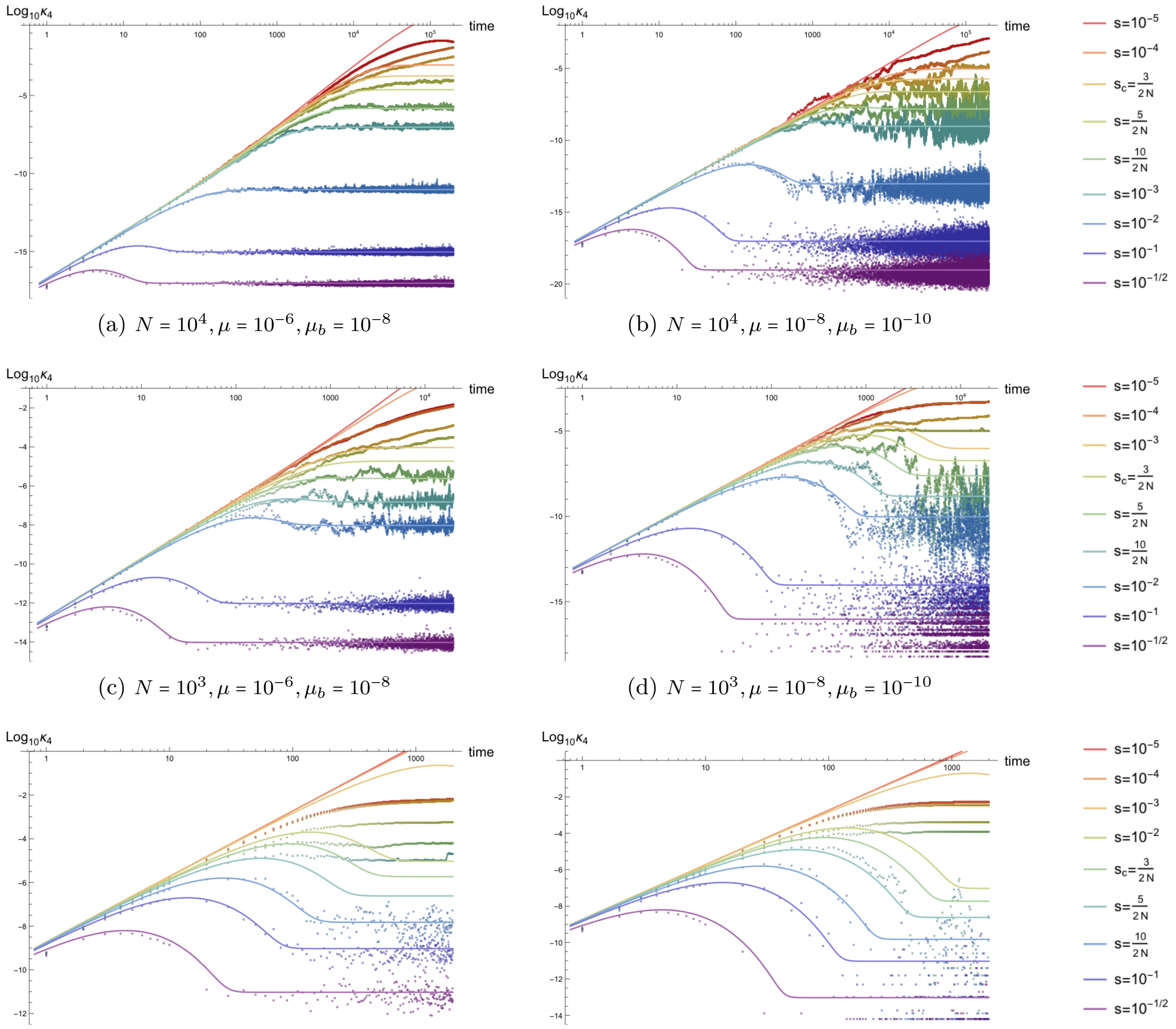


(a) $N = 10^4, \mu = 10^{-6}, \mu_b = 10^{-8}$
(b) $N = 10^4, \mu = 10^{-8}, \mu_b = 10^{-10}$
(c) $N = 10^3, \mu = 10^{-6}, \mu_b = 10^{-8}$
(d) $N = 10^3, \mu = 10^{-8}, \mu_b = 10^{-10}$
(e) $N = 10^2, \mu = 10^{-6}, \mu_b = 10^{-8}$
(f) $N = 10^2, \mu = 10^{-8}, \mu_b = 10^{-10}$

Figure S10: *Time dependence of the four-vertex approximations for the fourth cumulant $\kappa_4(t)$ for constant population size.* Parameter combinations correspond to those shown in Figure S2.

##### S4.1.11 Constant demography: fourth moment $E[p^4(t)]$ to four-vertex order

The lowest-vertex order approximation to the time dependence of the fourth non-central moment $E[p^4(t)]$ is plotted below for the same parameter combinations. Again, the fourth moment is initially $p_0^4$ at the delta function initial condition, but equilibrates to roughly the same value as the fourth cumulant.

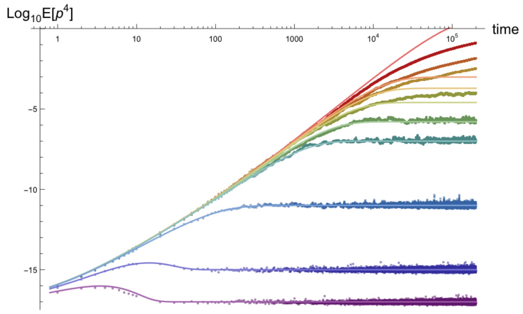


(a) $N = 10^4, \mu = 10^{-6}, \mu_b = 10^{-8}$

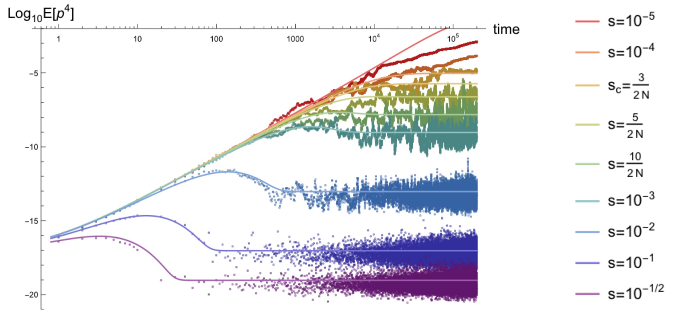


(b) $N = 10^4, \mu = 10^{-8}, \mu_b = 10^{-10}$

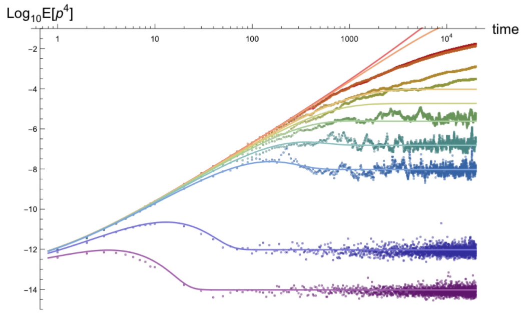


(c) $N = 10^3, \mu = 10^{-6}, \mu_b = 10^{-8}$

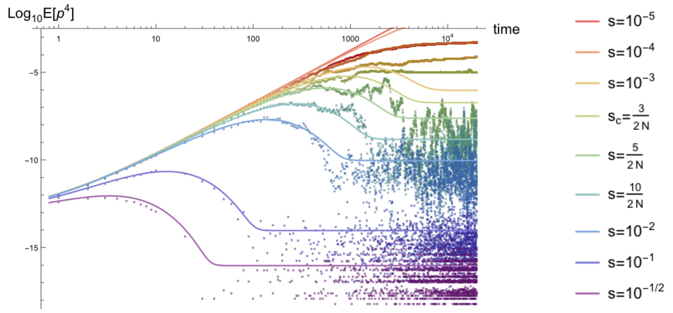


(d) $N = 10^3, \mu = 10^{-8}, \mu_b = 10^{-10}$

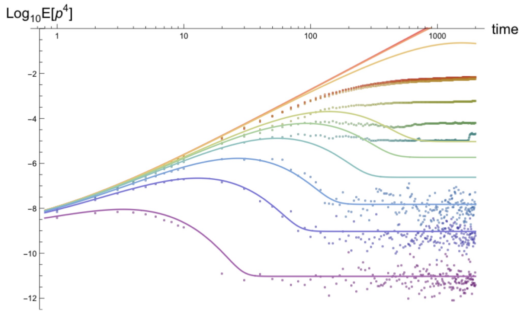


(e) $N = 10^2, \mu = 10^{-6}, \mu_b = 10^{-8}$

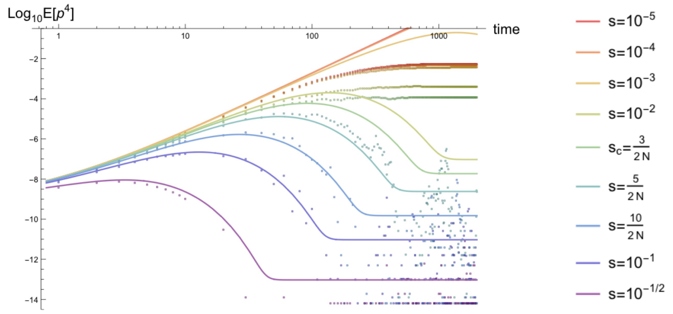


(f) $N = 10^2, \mu = 10^{-8}, \mu_b = 10^{-10}$

Figure S11: *Time dependence of the four-vertex approximations for the fourth non-central moment* $E[p^4(t)]$ *for constant population size.* Parameter combinations correspond to those shown in Figure S2.

##### S4.1.12 Constant demography: excess kurtosis $K_4(t)$ to four-vertex order

Here, the time dependence of the lowest-vertex (4V) order approximation to the excess kurtosis of the allele frequency probability distribution $K_4(t)$ is plotted. Like the skew, when the approximation describes the dynamics ($2Ns > 5$), the excess kurtosis is independent of the strength of selection at asymptotically late times. The equilibrium value $3/\lambda'\gamma' = 3/2N\mu$ is dependent only on the population scaled mutation rate $\theta = 4N\mu$ (compare the left to right columns for each row below). As with the skew, for low mutation rate and very small population size, the excess kurtosis fails to match analytic expectations due to discretization of the fourth cumulant, which can be seen in the Figures 12D and 12F below.

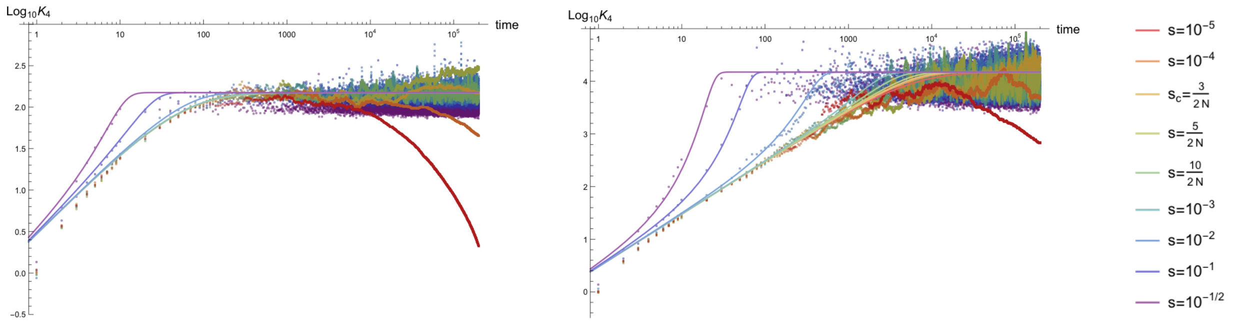


(a) $N = 10^4, \mu = 10^{-6}, \mu_b = 10^{-8}$ (b) $N = 10^4, \mu = 10^{-8}, \mu_b = 10^{-10}$

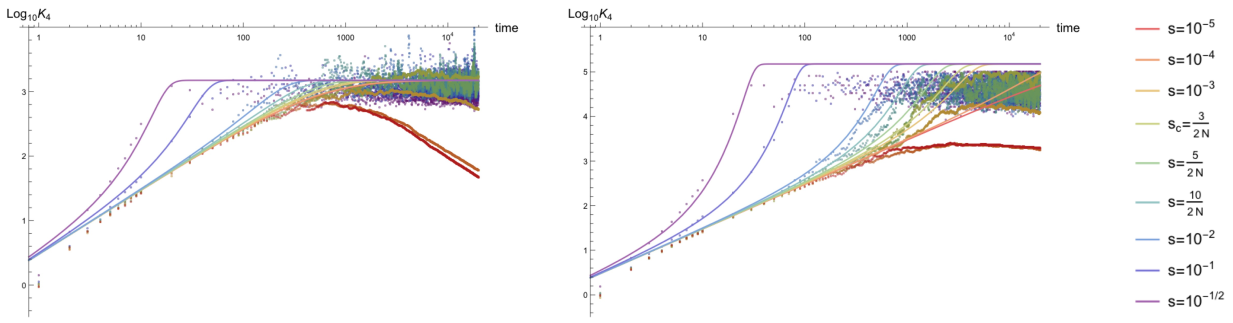


(c) $N = 10^3, \mu = 10^{-6}, \mu_b = 10^{-8}$ (d) $N = 10^3, \mu = 10^{-8}, \mu_b = 10^{-10}$

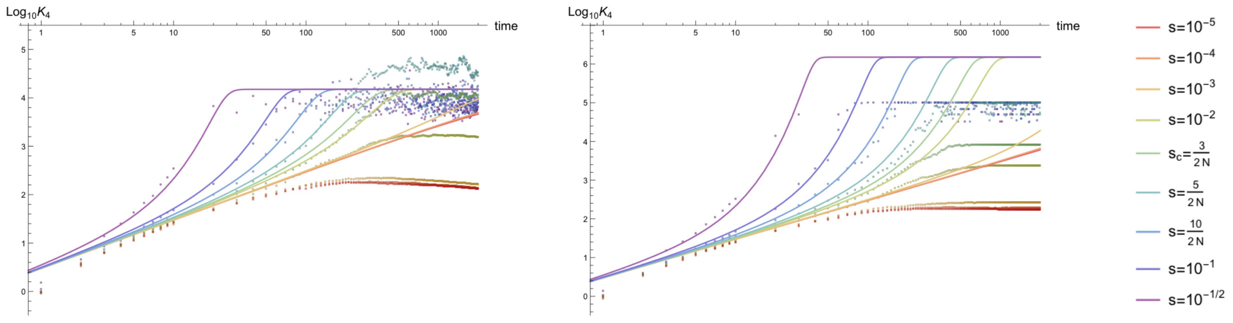


(e) $N = 10^2, \mu = 10^{-6}, \mu_b = 10^{-8}$ (f) $N = 10^2, \mu = 10^{-8}, \mu_b = 10^{-10}$

Figure S12: *Time dependence of the four-vertex approximations for the excess kurtosis of the allele frequency probability distribution $K_4(t)$ for constant population size.* Parameter combinations correspond to those shown in Figure S2.

#### S4.2 Exponential population growth

For a population undergoing exponential growth as parameterized in Equation S4, the plots in this section are shown for varying values of the initial population size $N_0$ and exponential growth rate $\Gamma$. As seen in the plots for constant sized populations in the above section, the primary effect of reducing the mutation rate $\mu$ (again keeping the back mutation rate $\mu_b$ fixed) is an increase in the relative amount of noise for each moment, since the equilibrated cumulants and moments are linear in the mutation rate at late times and thus equilibrate to lower quantitative values (the variance of each cumulant is also proportional to $\mu$ at late times such that the ratio of the standard deviation to the mean of each cumulant is proportional to $\sqrt{\mu}$). This can be seen, for example, in the comparison between the fourth cumulants in Figure S13 below, where the mutation rate in Figure S13a, but all other parameters are held fixed.

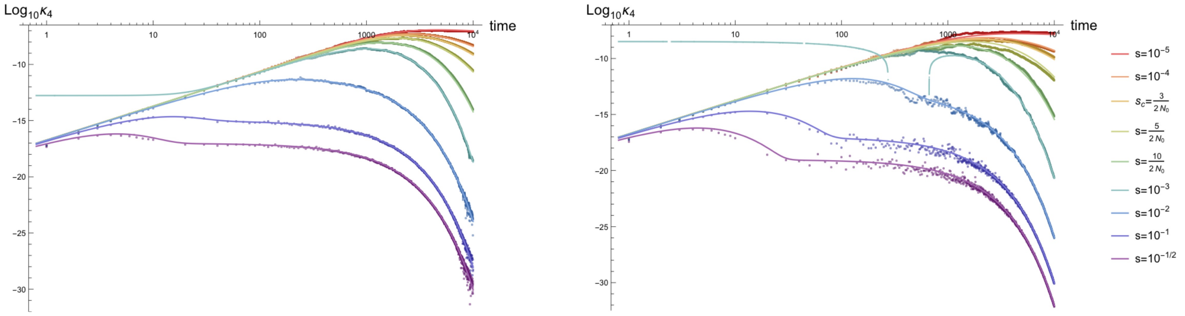


(a) Example $\kappa_4$ with $\mu = 10^{-6}, \mu_b = 10^{-8}$

(b) Example $\kappa_4$ with $\mu = 10^{-8}, \mu_b = 10^{-10}$

Figure S13: The effect of reducing the mutation rate is an increase the amount of noise relative to an exponentially growing population with the same parameters but increased mutation rate $\mu$ (with back mutation rate $\mu_b = \mu/10^2$). Here, the fourth cumulant is compared for different mutation rates in a population with exponential growth. The first, second, and third cumulants and moments and the fourth moment are affected in the same way.

In this section, figures are presented for a variety of parameter combinations of ($N_0$, $\Gamma$) and the same range of selection coefficients used for populations with constant size; selection coefficients between $10^{-5}$ and $10^{-1/2}$, along with selection coefficients chosen for each demography with $2Ns = 3, 5,$ and $10$. To reduce the number of figures, plots will only be shown comparing simulations to analytic curves with mutation rate $\mu = 10^{-8}$ for all demographies and $\mu = 10^{-6}$ for one value of the initial population size, $N_0 = 10^4$, for visual comparison; moments computed from all simulations with $\mu = 10^{-6}$ showed roughly the same shape with reduced noise and a simulated time-series nearly indistinguishable from the analytic curve in the regime of validity for the analytic approximations ($2N_0 s' > 5$). The one exception to this is for selection coefficients very close to the poles at $s' = \Gamma$ (or $\gamma'_{\text{exp}} = 0$), $2s' = \Gamma$, etc., which show the clearest deviation in the fourth cumulant and non-central moment. The example parameter combinations shown in Figure S13 were chosen to show the effect of selection coefficients in the neighborhood of the poles in the analytic expression for $\kappa_4(t|N_{\text{exp}})$. In both plots, the curves for selection coefficient $s = 10^{-3}$, which is very close in value to $s' = (s + \mu + \mu_b) = 10^{-3}$ (i.e., the value when $\Gamma = s'$ in the above plots), deviate wildly from the controlled analytic behavior of other values of strong selection. This behavior is due to the nearby pole where the approximation is no longer well-described. The third cumulant shows similarly pathological behavior in the neighborhood of one of the poles. Although a pole is present in the second cumulant under exponential population growth, the approximation is well controlled even for values close to $s = (\Gamma - \mu - \mu_b)$ (see Figure S16). However, the third and fourth cumulants (along with the corresponding moments, the skew, and the excess kurtosis) appear to be more sensitive to this issue at selection coefficients close to the pole, likely due to the quadratic and cubic dependence on $1/\gamma'_{\text{exp}}$ in the third and fourth cumulants, respectively, rather than the linear dependence appearing in $\kappa_2$ (i.e., $\kappa_3$ and $\kappa_4$ have higher order divergences than $\kappa_2$). For the following simulations, all moments and cumulants are computed as an average of $L = 10^5$ independent sites evolving through the same demography.

##### S4.2.1 Exponential demography: time dependence of the population size

The following log-log plots show the time dependence of the population size under exponential growth for all combinations of the initial population size $N_0 = 10^4$ (top two rows), $10^3$ (third row), $10^2$ (fourth row) and growth rate $\Gamma = 10^{-4}, 10^{-3}, 10^{-2}$ (from left to right). Analytic and simulation results will be presented in the same order as the time-dependent population sizes. The top two rows are identical in the figure below; the first row of all subsequent plots of cumulants and moments will be shown with mutation rates $\mu = 10^{-6}, \mu_b = 10^{-8}$ in the same demography as the second row with $\mu = 10^{-8}, \mu_b = 10^{-10}$. The second, third, and fourth rows of plots will show results for various demographies with mutation rates $\mu = 10^{-8}, \mu_b = 10^{-10}$.

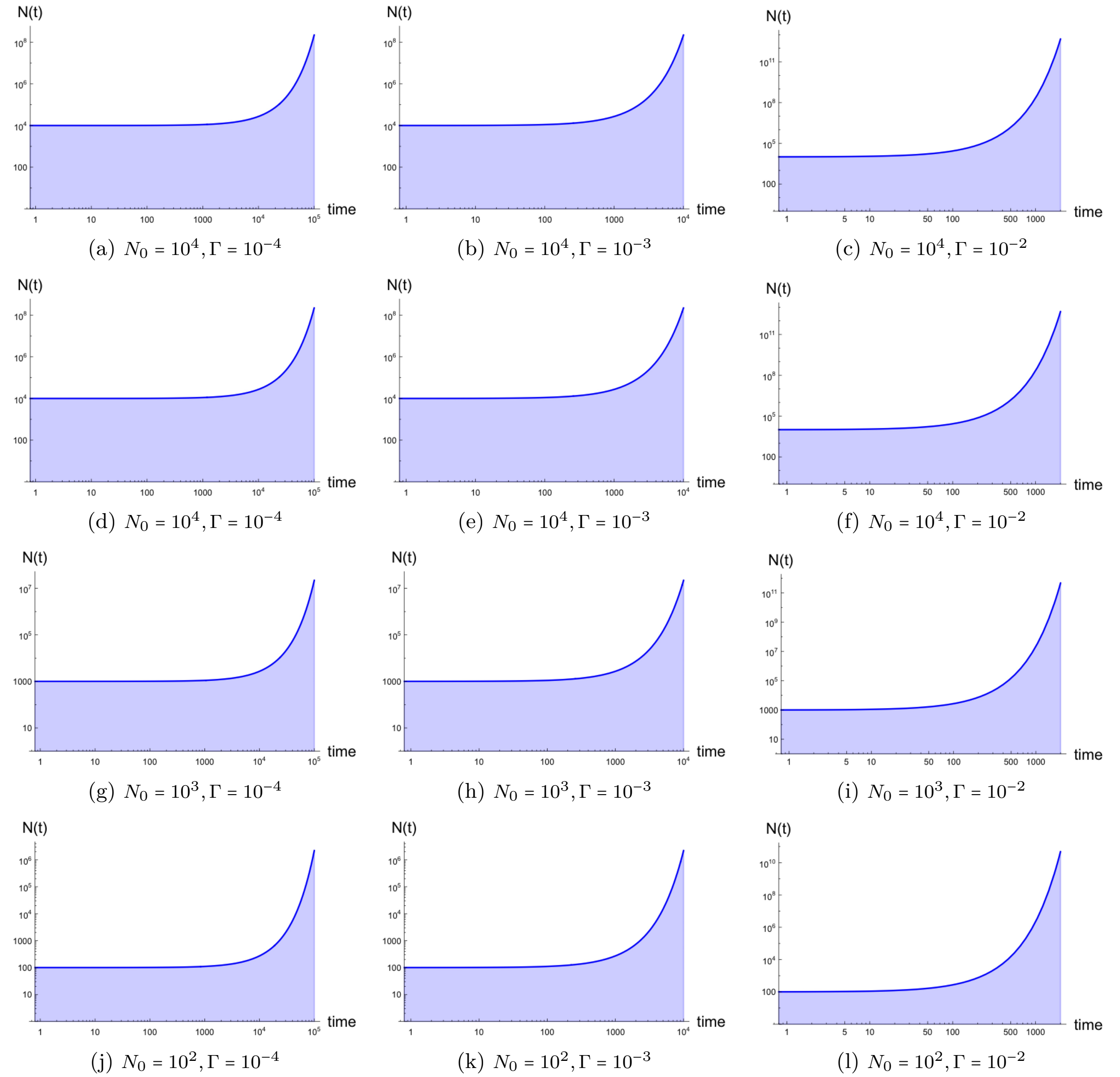


(a) $N_0 = 10^4, \Gamma = 10^{-4}$ (b) $N_0 = 10^4, \Gamma = 10^{-3}$ (c) $N_0 = 10^4, \Gamma = 10^{-2}$

(d) $N_0 = 10^4, \Gamma = 10^{-4}$ (e) $N_0 = 10^4, \Gamma = 10^{-3}$ (f) $N_0 = 10^4, \Gamma = 10^{-2}$

(g) $N_0 = 10^3, \Gamma = 10^{-4}$ (h) $N_0 = 10^3, \Gamma = 10^{-3}$ (i) $N_0 = 10^3, \Gamma = 10^{-2}$

(j) $N_0 = 10^2, \Gamma = 10^{-4}$ (k) $N_0 = 10^2, \Gamma = 10^{-3}$ (l) $N_0 = 10^2, \Gamma = 10^{-2}$

Figure S14: *Time dependence of the population size for simulated populations under exponential growth.* Demographies are presented with initial size $N_0 = 10^4$ (a–f), $10^3$ (g–i), $10^2$ (j–l) and growth rate $\Gamma = 10^{-4}$ (left), $10^{-3}$(center), $10^{-2}$ (right).

##### S4.2.2 Exponential demography: mean frequency $\overline{p}(t)$ to three-vertex order

Since the lowest order approximation of the mean frequency $\overline{p}(t)$ under strong selection is independent of the population size, and thus the demography, plots for an exponentially growing population are identical to those in Figure S2 up to the number of simulated generations. The figure below shows the three-vertex approximation of the mean frequency, with three-vertex corrections to the one-vertex deterministic solution. The effect of the demography on strong selection is infinitesimal. Simulations for weak selection show a marked difference from the lowest order approximation, but are not well-described by the analytic approximations presented in this manuscript. For all following subfigures in this demography, the mutation rate will be annotated, but the back mutation rate $\mu_b$ (where $\mu_b = \mu/10^2$ in all cases) will remain implicit.

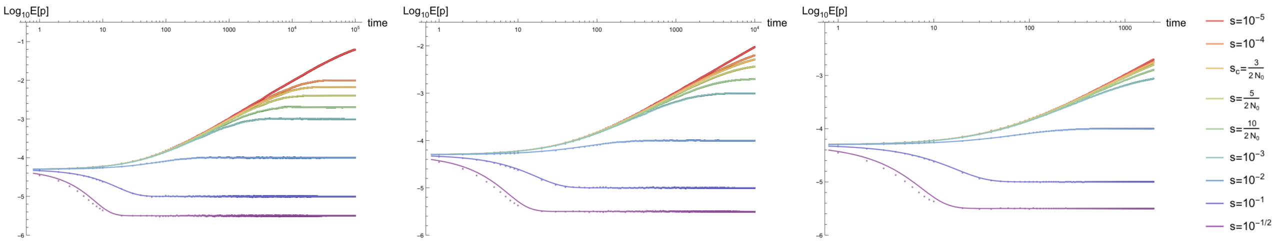


(a) $N_0 = 10^4, \Gamma = 10^{-4}, \mu = 10^{-6}$ (b) $N_0 = 10^4, \Gamma = 10^{-3}, \mu = 10^{-6}$ (c) $N_0 = 10^4, \Gamma = 10^{-2}, \mu = 10^{-6}$

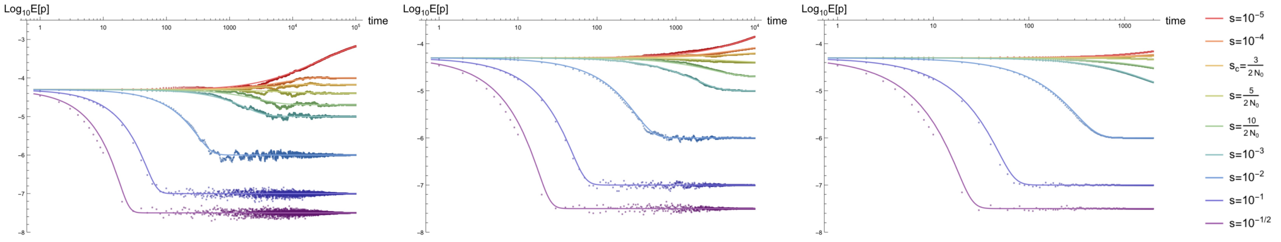

(d) $N_0 = 10^4, \Gamma = 10^{-4}, \mu = 10^{-8}$ (e) $N_0 = 10^4, \Gamma = 10^{-3}, \mu = 10^{-8}$ (f) $N_0 = 10^4, \Gamma = 10^{-2}, \mu = 10^{-8}$

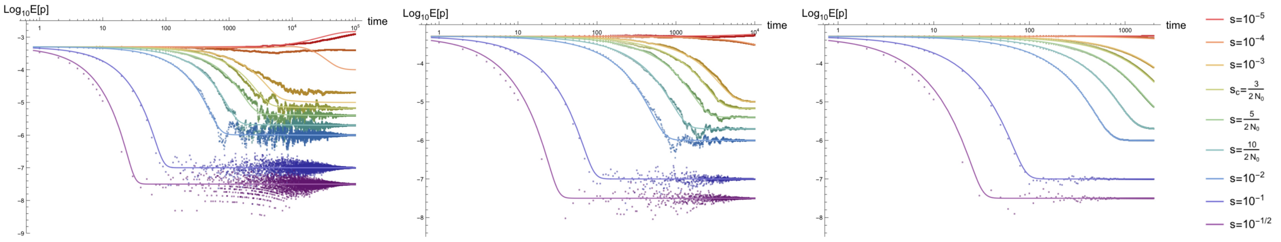

(g) $N_0 = 10^3, \Gamma = 10^{-4}, \mu = 10^{-8}$ (h) $N_0 = 10^3, \Gamma = 10^{-3}, \mu = 10^{-8}$ (i) $N_0 = 10^3, \Gamma = 10^{-2}, \mu = 10^{-8}$

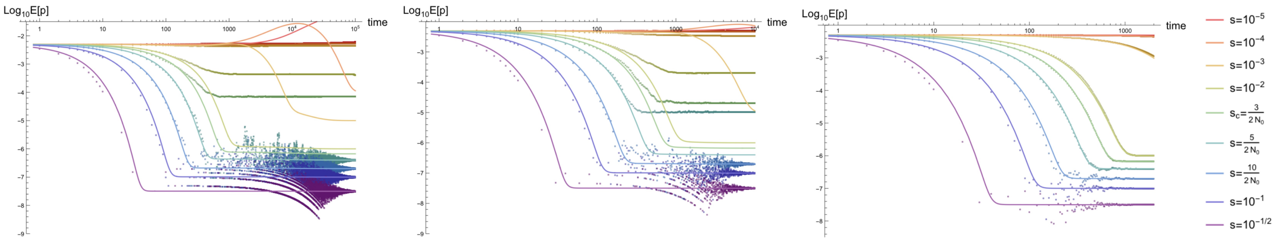

(j) $N_0 = 10^2, \Gamma = 10^{-4}, \mu = 10^{-8}$ (k) $N_0 = 10^2, \Gamma = 10^{-3}, \mu = 10^{-8}$ (l) $N_0 = 10^2, \Gamma = 10^{-2}, \mu = 10^{-8}$

Figure S15: *Time dependence of the mean frequency $\overline{p}(t)$ in an exponentially growing population including lowest order drift-based corrections.* Plots are shown for mutation rates $\mu = 10^{-6}$ (top row), $10^{-8}$ (all other rows) and $\mu_b = \mu/100$, initial size $N = 10^4$ (top two rows), $10^3$(third row), $10^2$ (fourth row), growth rate $\Gamma = 10^{-4}$(left), $10^{-3}$(center), $10^{-2}$ (right), and selection coefficients $s \in [10^{-5}, 10^{-1/2}]$.

##### S4.2.3 Exponential demography: variance $\kappa_2(t)$ to two-vertex order

For the variance, the second cumulant of the allele frequency probability distribution, the analytic approximation remains accurate beyond the strong initial population size regime ($2N_0 s' > 5$) due to rapid growth decreasing the effects of drift over time. This breaks down for slow exponential growth ($\Gamma = 10^{-4}$) for small initial population size. If the growth rate is sufficiently high ($\Gamma \gtrsim 10^{-2}$), the analytic approximations under strong selection apply for weak selection even in populations starting at very small sizes (e.g., $N_0 = 100$). For high mutation rates ($\mu = 10^{-6}$) and relatively large initial size ($N_0 = 10^4$), the analytic approximation for $E[p^2(t)]$ is extremely accurate independent of the strength of selection and the rate of population growth. The associated dynamics and regime of validity apply equally well to the homozygosity, with differences only in the initial and final states, which vanish for the variance.

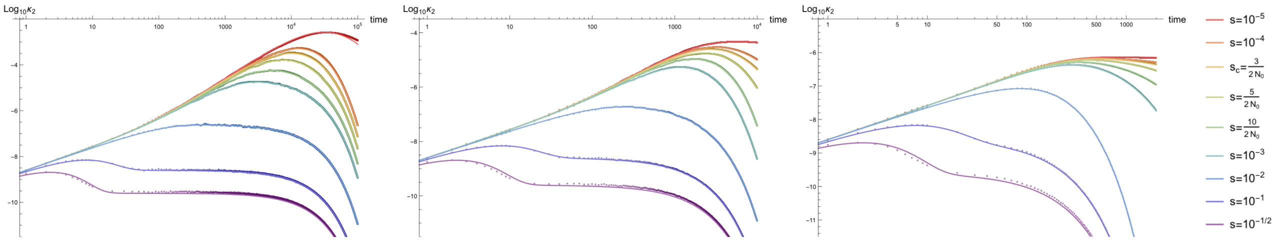

(a) $N_0 = 10^4, \Gamma = 10^{-4}, \mu = 10^{-6}$ (b) $N_0 = 10^4, \Gamma = 10^{-3}, \mu = 10^{-6}$ (c) $N_0 = 10^4, \Gamma = 10^{-2}, \mu = 10^{-6}$

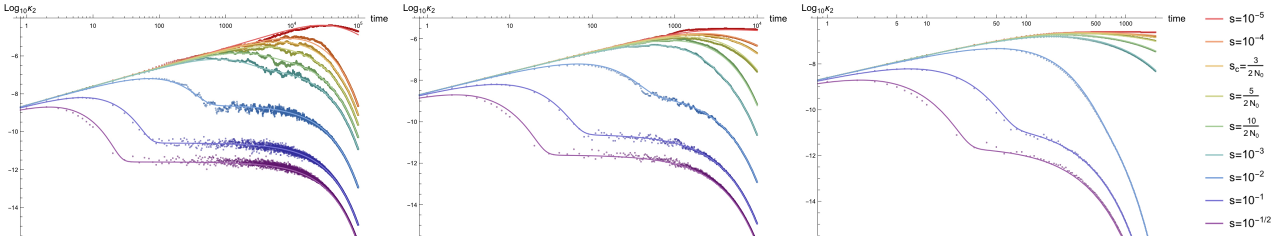

(d) $N_0 = 10^4, \Gamma = 10^{-4}, \mu = 10^{-8}$ (e) $N_0 = 10^4, \Gamma = 10^{-3}, \mu = 10^{-8}$ (f) $N_0 = 10^4, \Gamma = 10^{-2}, \mu = 10^{-8}$

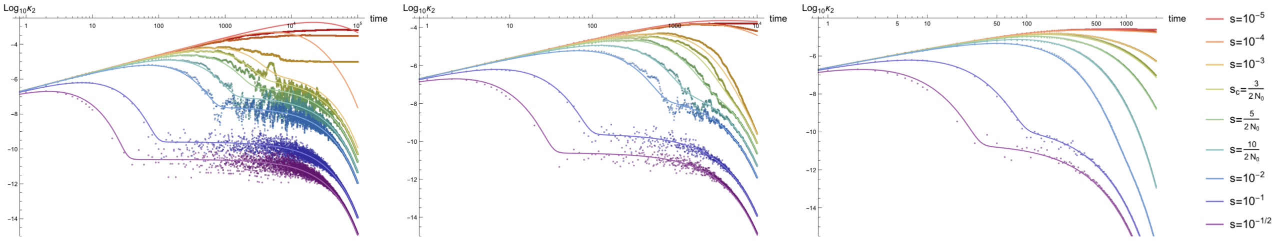

(g) $N_0 = 10^3, \Gamma = 10^{-4}, \mu = 10^{-8}$ (h) $N_0 = 10^3, \Gamma = 10^{-3}, \mu = 10^{-8}$ (i) $N_0 = 10^3, \Gamma = 10^{-2}, \mu = 10^{-8}$

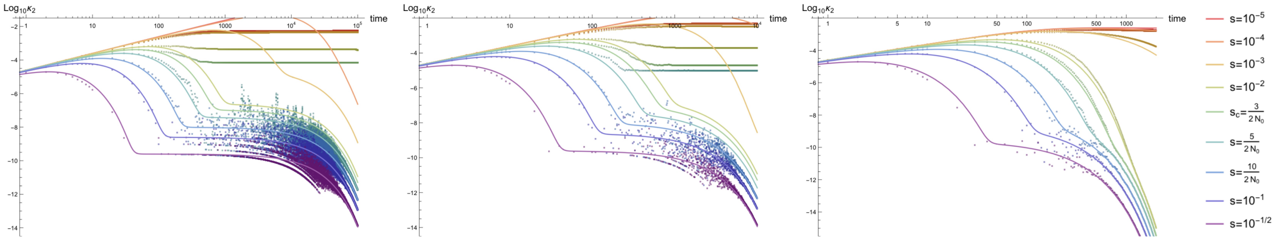

(j) $N_0 = 10^2, \Gamma = 10^{-4}, \mu = 10^{-8}$ (k) $N_0 = 10^2, \Gamma = 10^{-3}, \mu = 10^{-8}$ (l) $N_0 = 10^2, \Gamma = 10^{-2}, \mu = 10^{-8}$

Figure S16: *Time dependence of the variance of the allele frequency probability distribution $\kappa_2(t)$ in an exponentially growing population.* Parameter combinations correspond to those shown in Figure S15.

##### S4.2.4 Exponential demography: homozygosity $E[p^2(t)]$ to two-vertex order

The figure below depicts comparisons between the two-vertex approximation of the homozygosity, the second non-central moment of the allele frequency probability distribution, to simulations of the same moment. Like the variance, the homozygosity is drastically affected by rapid population growth and takes a highly non-monotonic path through time: initially the homozygosity takes on a value of $p_0^2$ dictated by the delta function initial condition, then reaches a maximum at early times (see discussion of $t_{\kappa_{2\max}}$ in the manuscript text), which is followed by reduction to a pseudo-equilibrated state (roughly until the e-folding time $t \sim 1/\Gamma$) prior to eventually reaching an equilibrium value of $(\frac{\mu}{s'})^2$ at asymptotically late times. The pseudo-equilibrated state is similar to the value of the homozygosity in a population of constant size $N = N_0$ (the initial size) and only appears for sufficiently strong selection such that the equivalent of the equilibration time in a constant population is faster than $t \sim 1/\Gamma$ (i.e., before the population size is dramatically changed).

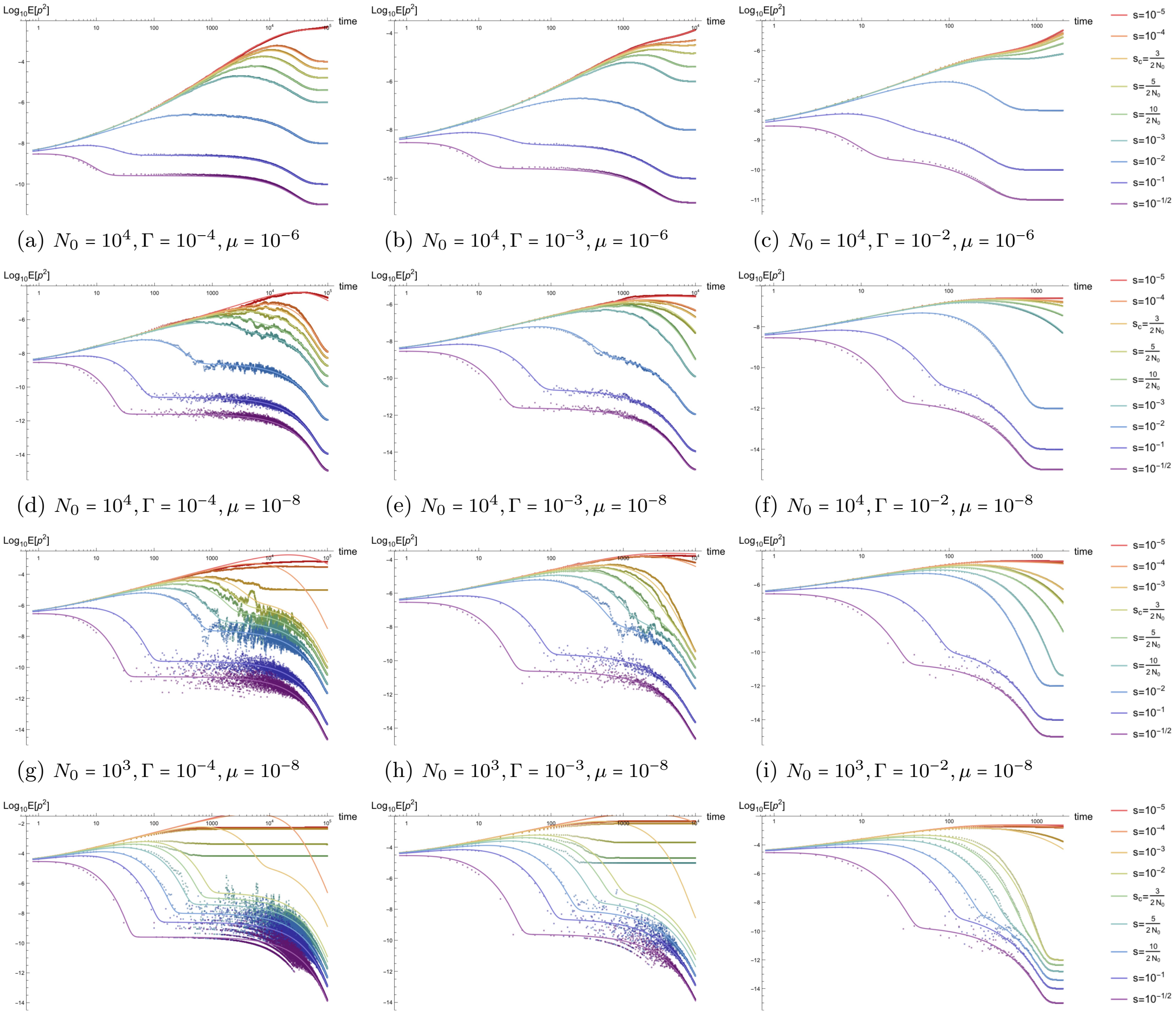


(a) $N_0 = 10^4, \Gamma = 10^{-4}, \mu = 10^{-6}$ (b) $N_0 = 10^4, \Gamma = 10^{-3}, \mu = 10^{-6}$ (c) $N_0 = 10^4, \Gamma = 10^{-2}, \mu = 10^{-6}$

(d) $N_0 = 10^4, \Gamma = 10^{-4}, \mu = 10^{-8}$ (e) $N_0 = 10^4, \Gamma = 10^{-3}, \mu = 10^{-8}$ (f) $N_0 = 10^4, \Gamma = 10^{-2}, \mu = 10^{-8}$

(g) $N_0 = 10^3, \Gamma = 10^{-4}, \mu = 10^{-8}$ (h) $N_0 = 10^3, \Gamma = 10^{-3}, \mu = 10^{-8}$ (i) $N_0 = 10^3, \Gamma = 10^{-2}, \mu = 10^{-8}$

(j) $N_0 = 10^2, \Gamma = 10^{-4}, \mu = 10^{-8}$ (k) $N_0 = 10^2, \Gamma = 10^{-3}, \mu = 10^{-8}$ (l) $N_0 = 10^2, \Gamma = 10^{-2}, \mu = 10^{-8}$

Figure S17: *Time dependence of the homozygosity $E[p^2(t)]$ in an exponentially growing population.* Parameter combinations correspond to those shown in Figure S15.

##### S4.2.5 Exponential demography: third cumulant $\kappa_3(t)$ to three-vertex order

Much of the dynamics of higher cumulants echo the variance. In plots of the third cumulant of the allele frequency probability distribution below, there is a clear maximum at early times, a pseudo-equilibrated state until the e-folding time at $t \sim 1/\Gamma$ when the growth rate overtakes this and rapidly drives the third cumulant to zero (corresponding to the deterministic delta function distribution at asymptotically late times). Note that the overall scale of the third cumulant is substantially smaller than that of the second cumulant, as expected. The analytic approximation is extremely accurate for high mutation rates, and applies to weaker selection coefficients due to the exponential decay of the drift constant as the population size grows.

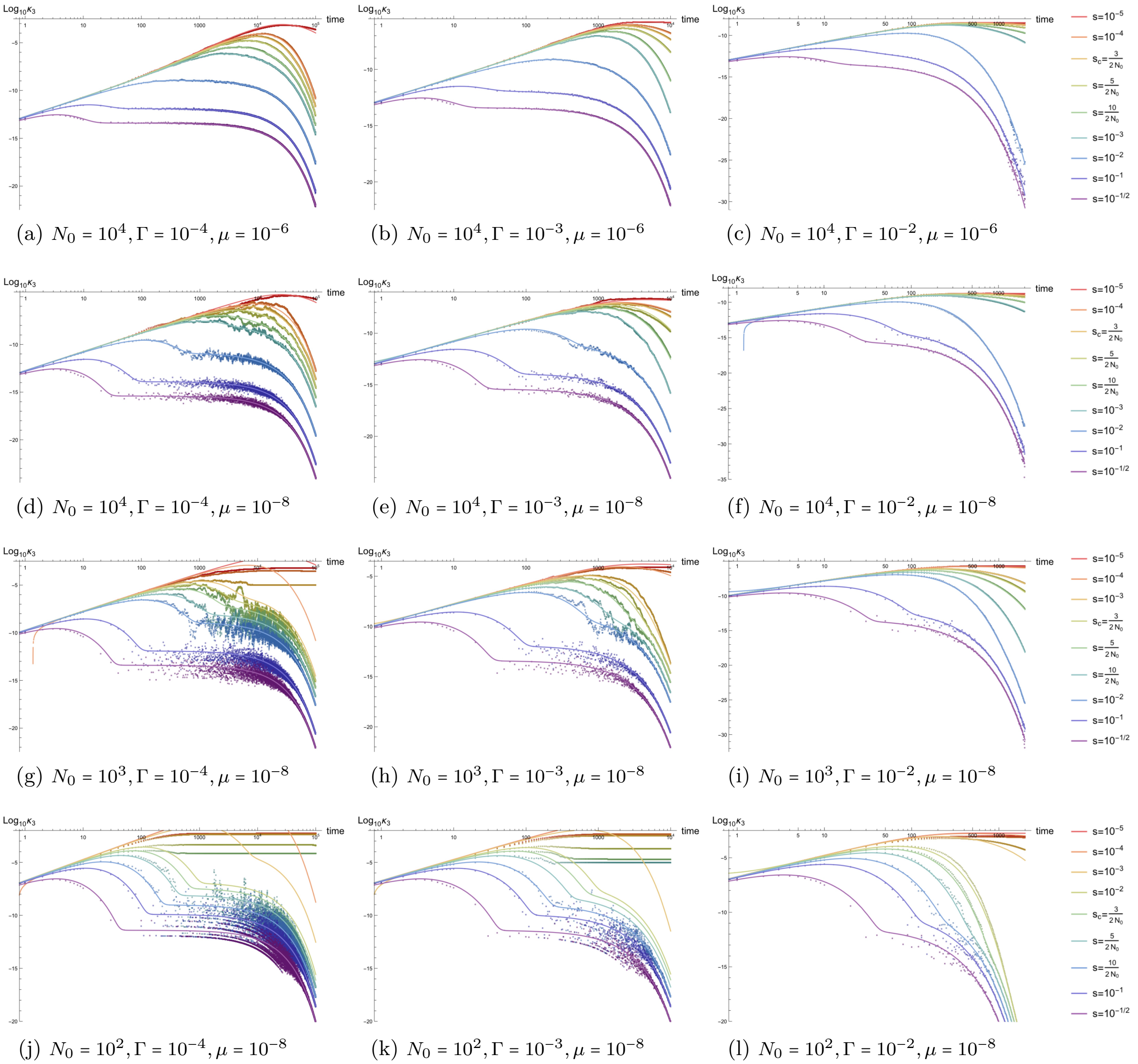


(a) $N_0 = 10^4, \Gamma = 10^{-4}, \mu = 10^{-6}$ (b) $N_0 = 10^4, \Gamma = 10^{-3}, \mu = 10^{-6}$ (c) $N_0 = 10^4, \Gamma = 10^{-2}, \mu = 10^{-6}$

(d) $N_0 = 10^4, \Gamma = 10^{-4}, \mu = 10^{-8}$ (e) $N_0 = 10^4, \Gamma = 10^{-3}, \mu = 10^{-8}$ (f) $N_0 = 10^4, \Gamma = 10^{-2}, \mu = 10^{-8}$

(g) $N_0 = 10^3, \Gamma = 10^{-4}, \mu = 10^{-8}$ (h) $N_0 = 10^3, \Gamma = 10^{-3}, \mu = 10^{-8}$ (i) $N_0 = 10^3, \Gamma = 10^{-2}, \mu = 10^{-8}$

(j) $N_0 = 10^2, \Gamma = 10^{-4}, \mu = 10^{-8}$ (k) $N_0 = 10^2, \Gamma = 10^{-3}, \mu = 10^{-8}$ (l) $N_0 = 10^2, \Gamma = 10^{-2}, \mu = 10^{-8}$

Figure S18: *Time dependence of the third cumulant $\kappa_3(t)$ in an exponentially growing population.* Parameter combinations correspond to those shown in Figure S15.

##### S4.2.6 Exponential demography: third moment $E[p^3(t)]$ to three-vertex order

The third non-central moment of the allele frequency probability distribution mimics the behavior of the homozygosity, analogous to the discussion of the third cumulant above. Like the second moment, the third moment under exponential growth asymptotes to a finite value, approaching the third moment of a delta function distribution at late times as the effects of drift are reduced.

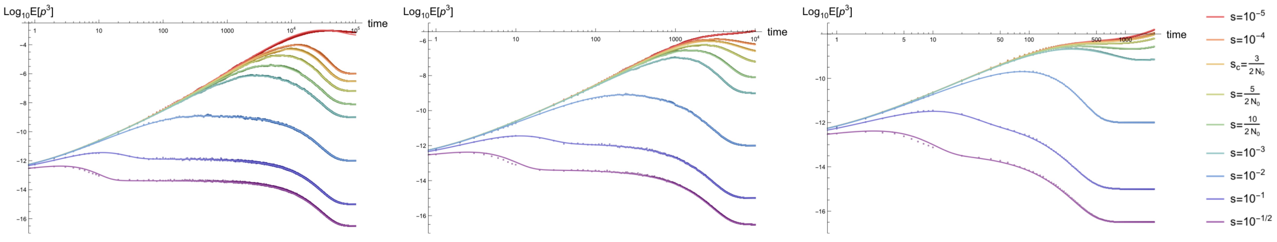

(a) $N_0 = 10^4, \Gamma = 10^{-4}, \mu = 10^{-6}$ (b) $N_0 = 10^4, \Gamma = 10^{-3}, \mu = 10^{-6}$ (c) $N_0 = 10^4, \Gamma = 10^{-2}, \mu = 10^{-6}$

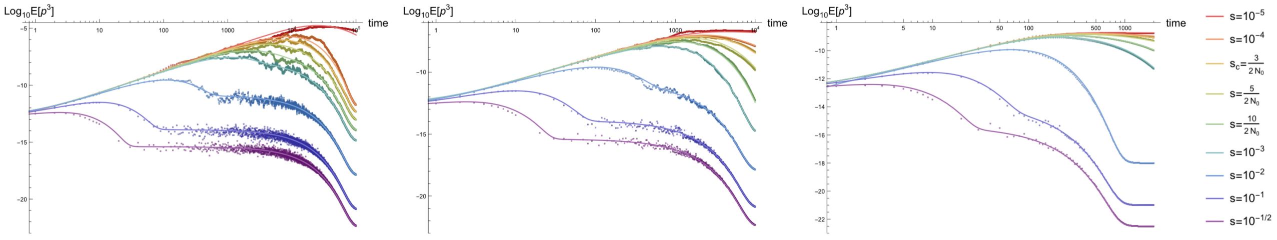

(d) $N_0 = 10^4, \Gamma = 10^{-4}, \mu = 10^{-8}$ (e) $N_0 = 10^4, \Gamma = 10^{-3}, \mu = 10^{-8}$ (f) $N_0 = 10^4, \Gamma = 10^{-2}, \mu = 10^{-8}$

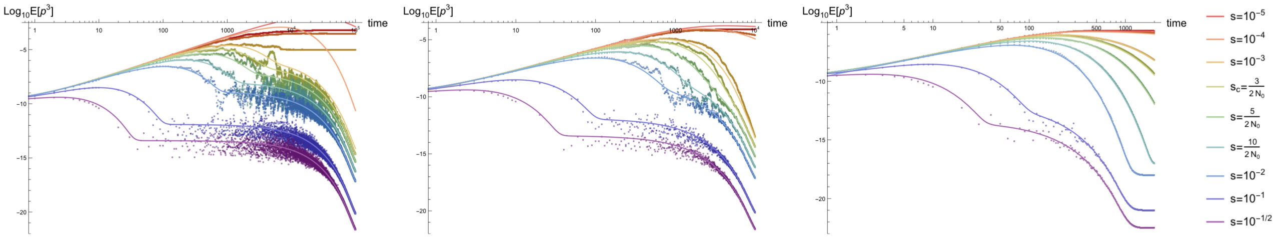

(g) $N_0 = 10^3, \Gamma = 10^{-4}, \mu = 10^{-8}$ (h) $N_0 = 10^3, \Gamma = 10^{-3}, \mu = 10^{-8}$ (i) $N_0 = 10^3, \Gamma = 10^{-2}, \mu = 10^{-8}$

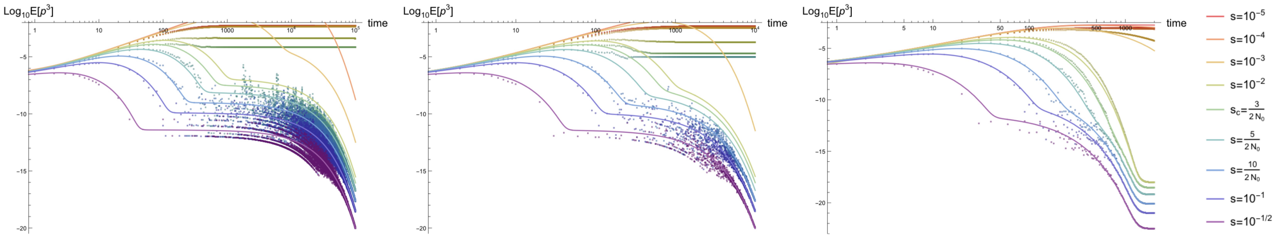

(j) $N_0 = 10^2, \Gamma = 10^{-4}, \mu = 10^{-8}$ (k) $N_0 = 10^2, \Gamma = 10^{-3}, \mu = 10^{-8}$ (l) $N_0 = 10^2, \Gamma = 10^{-2}, \mu = 10^{-8}$

Figure S19: *Time dependence of the third non-central moment of the allele frequency probability distribution $E[p^3(t)]$ in an exponentially growing population.* Parameter combinations correspond to those shown in Figure S15.

##### S4.2.7 Exponential demography: skew $K_3(t)$ to three-vertex order

The dynamics of the skew in an exponentially growing population drastically differ from the skew in a constant sized population (see Section S4.1.9). The pseudo-equilibrium regime is not quite flat here, indicating that even after rapidly equilibrating under strong selection, the skew slowly decays until the e-folding time, after which it rapidly vanishes. Under many of the demographies shown herein, the dynamics under weak selection decouple and reach a different pseudo-equilibrium at late times. At the end of the plots shown below, the population sizes are so large (e.g., roughly $10^{12}$ diploid individuals for demographies in the right column) that growth likely becomes unsustainable in most organisms; if the population size saturates, the pseudo-equilibrium shown for weak selection likely equilibrate in this state.

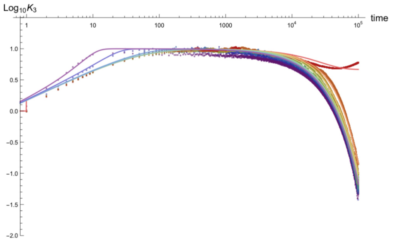


(a) $N_0 = 10^4, \Gamma = 10^{-4}, \mu = 10^{-6}$

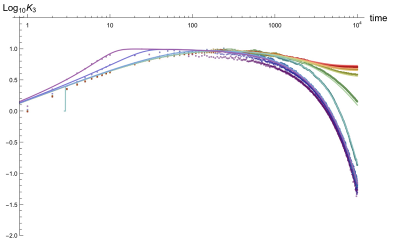


(b) $N_0 = 10^4, \Gamma = 10^{-3}, \mu = 10^{-6}$

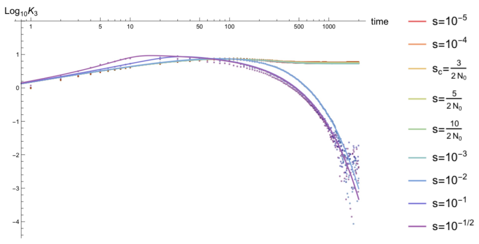


(c) $N_0 = 10^4, \Gamma = 10^{-2}, \mu = 10^{-6}$

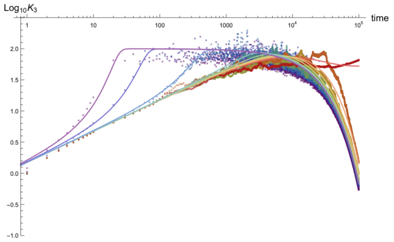


(d) $N_0 = 10^4, \Gamma = 10^{-4}, \mu = 10^{-8}$

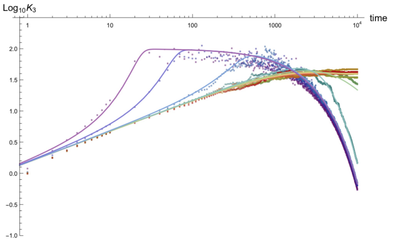


(e) $N_0 = 10^4, \Gamma = 10^{-3}, \mu = 10^{-8}$

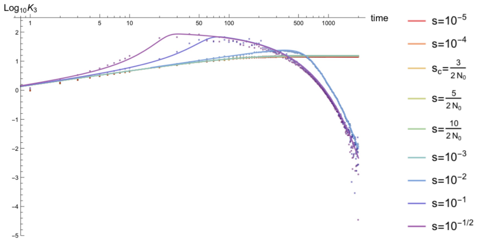


(f) $N_0 = 10^4, \Gamma = 10^{-2}, \mu = 10^{-8}$

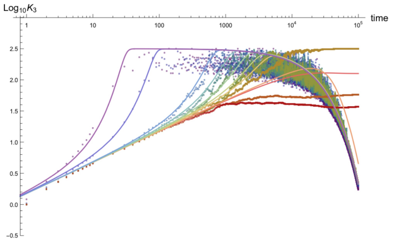


(g) $N_0 = 10^3, \Gamma = 10^{-4}, \mu = 10^{-8}$

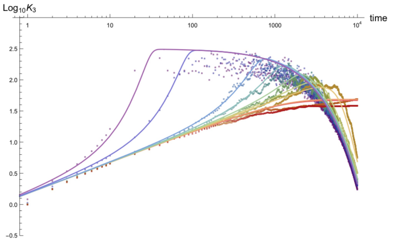


(h) $N_0 = 10^3, \Gamma = 10^{-3}, \mu = 10^{-8}$

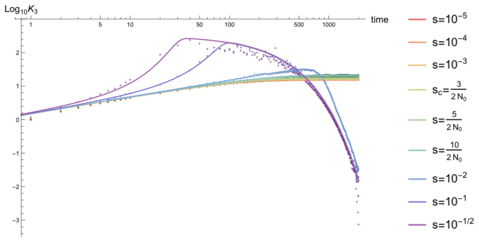


(i) $N_0 = 10^3, \Gamma = 10^{-2}, \mu = 10^{-8}$

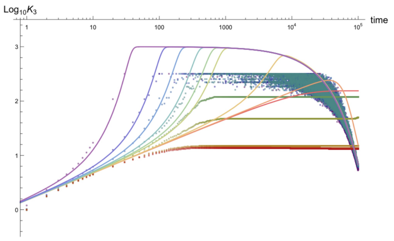


(j) $N_0 = 10^2, \Gamma = 10^{-4}, \mu = 10^{-8}$

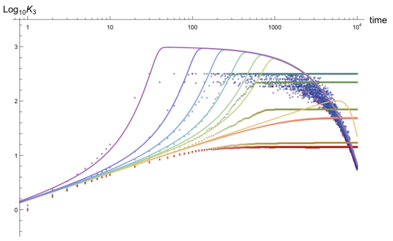


(k) $N_0 = 10^2, \Gamma = 10^{-3}, \mu = 10^{-8}$

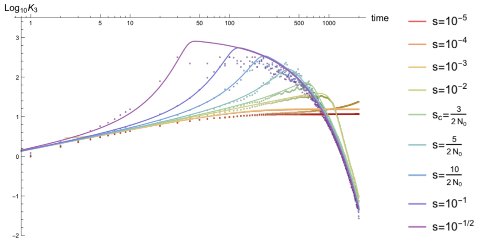


(l) $N_0 = 10^2, \Gamma = 10^{-2}, \mu = 10^{-8}$

Figure S20: *Time dependence of the skew of the allele frequency probability distribution $K_3(t)$ in an exponentially growing population.* Parameter combinations correspond to those shown in Figure S15.

##### S4.2.8 Exponential demography: fourth cumulant $\kappa_4(t)$ to four-vertex order

The dynamics of the fourth cumulant under exponential expansion are largely the same as for the third cumulant, mimicking the second moment, but on an overall reduced scale. See description in Section S4.2.7.

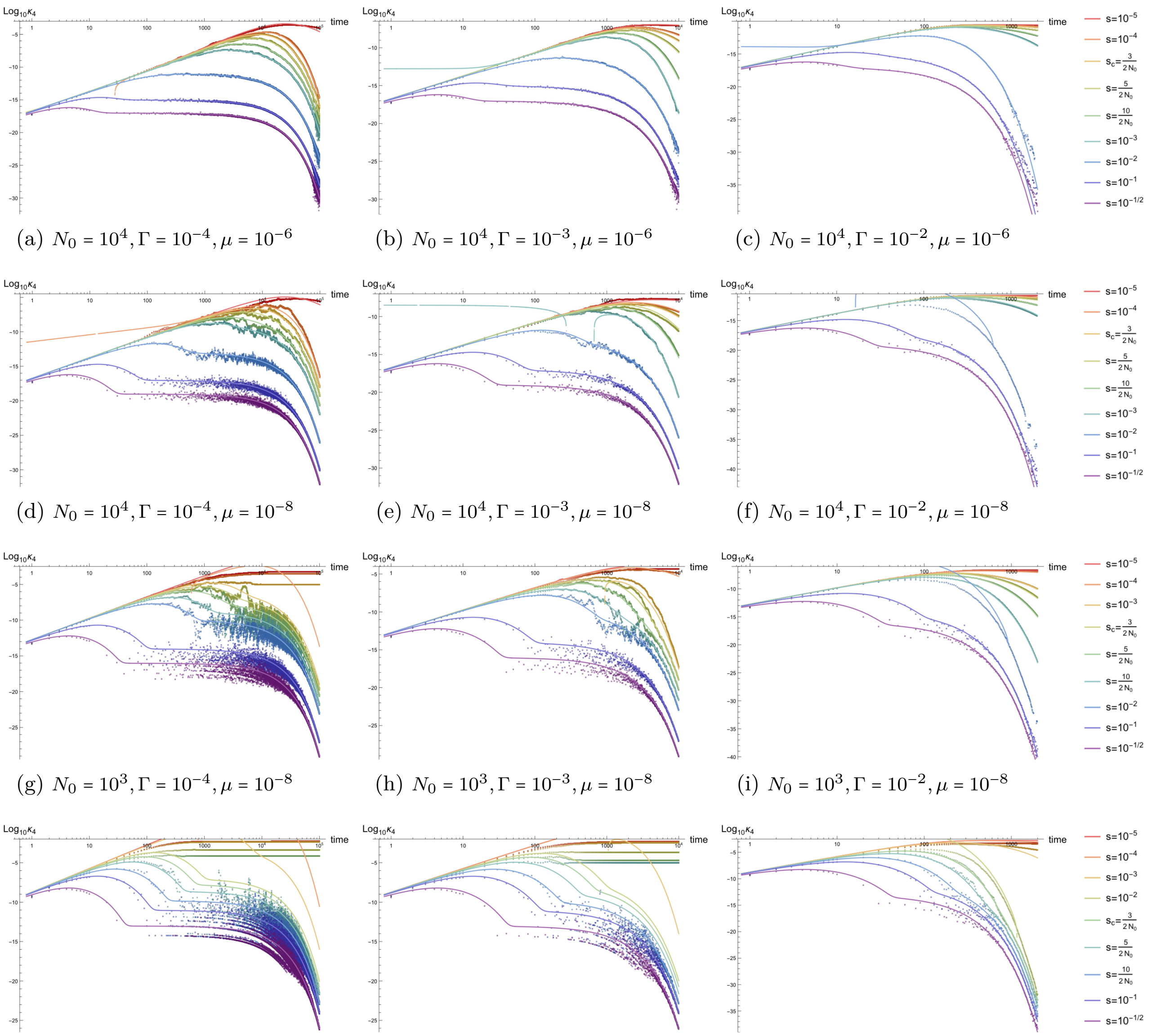


(a) $N_0 = 10^4, \Gamma = 10^{-4}, \mu = 10^{-6}$ (b) $N_0 = 10^4, \Gamma = 10^{-3}, \mu = 10^{-6}$ (c) $N_0 = 10^4, \Gamma = 10^{-2}, \mu = 10^{-6}$

(d) $N_0 = 10^4, \Gamma = 10^{-4}, \mu = 10^{-8}$ (e) $N_0 = 10^4, \Gamma = 10^{-3}, \mu = 10^{-8}$ (f) $N_0 = 10^4, \Gamma = 10^{-2}, \mu = 10^{-8}$

(g) $N_0 = 10^3, \Gamma = 10^{-4}, \mu = 10^{-8}$ (h) $N_0 = 10^3, \Gamma = 10^{-3}, \mu = 10^{-8}$ (i) $N_0 = 10^3, \Gamma = 10^{-2}, \mu = 10^{-8}$

(j) $N_0 = 10^2, \Gamma = 10^{-4}, \mu = 10^{-8}$ (k) $N_0 = 10^2, \Gamma = 10^{-3}, \mu = 10^{-8}$ (l) $N_0 = 10^2, \Gamma = 10^{-2}, \mu = 10^{-8}$

Figure S21: *Time dependence of the fourth cumulant $\kappa_4(t)$ in an exponentially growing population.* Parameter combinations correspond to those shown in Figure S15.

##### S4.2.9 Exponential demography: fourth moment $E[p^4(t)]$ to four-vertex order

As with the fourth cumulant, the fourth non-central moment of the allele frequency probability distribution under exponential expansion is largely controlled by the dynamics of the second moment, except in overall scale, which is reduced, and a much more rapid decay (see Figure S23 for comparison after standardization).

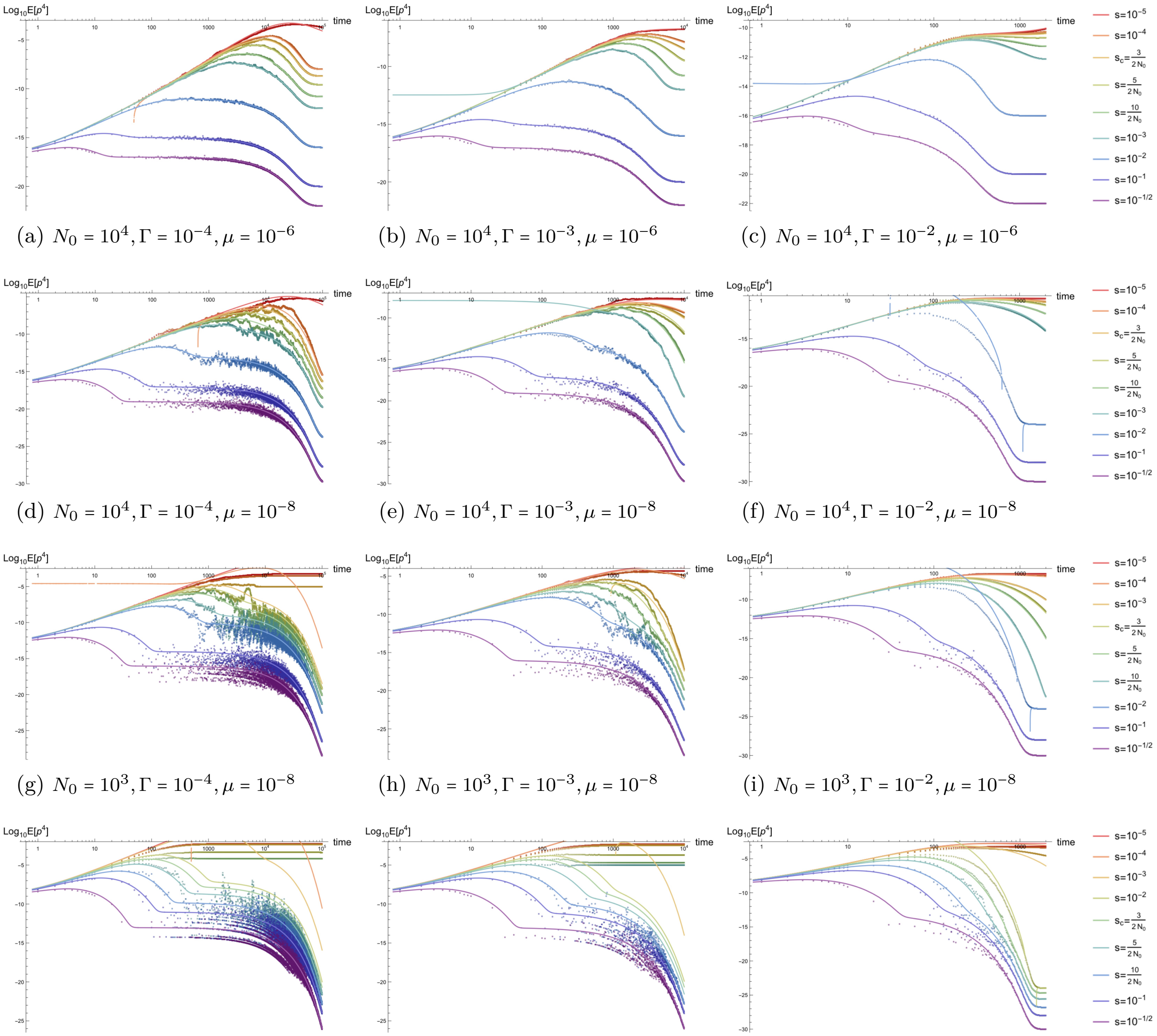


(a) $N_0 = 10^4, \Gamma = 10^{-4}, \mu = 10^{-6}$ (b) $N_0 = 10^4, \Gamma = 10^{-3}, \mu = 10^{-6}$ (c) $N_0 = 10^4, \Gamma = 10^{-2}, \mu = 10^{-6}$

(d) $N_0 = 10^4, \Gamma = 10^{-4}, \mu = 10^{-8}$ (e) $N_0 = 10^4, \Gamma = 10^{-3}, \mu = 10^{-8}$ (f) $N_0 = 10^4, \Gamma = 10^{-2}, \mu = 10^{-8}$

(g) $N_0 = 10^3, \Gamma = 10^{-4}, \mu = 10^{-8}$ (h) $N_0 = 10^3, \Gamma = 10^{-3}, \mu = 10^{-8}$ (i) $N_0 = 10^3, \Gamma = 10^{-2}, \mu = 10^{-8}$

(j) $N_0 = 10^2, \Gamma = 10^{-4}, \mu = 10^{-8}$ (k) $N_0 = 10^2, \Gamma = 10^{-3}, \mu = 10^{-8}$ (l) $N_0 = 10^2, \Gamma = 10^{-2}, \mu = 10^{-8}$

Figure S22: *Time dependence of the fourth moment of the allele frequency distribution $\kappa_4(t)$ in an exponentially growing population.* Parameter combinations correspond to those shown in Figure S15.

##### S4.2.10 Exponential demography: excess kurtosis $K_4(t)$ to four-vertex order

The dynamics of the excess kurtosis are similar to those of the skew under exponential growth. Note that, like the skew, an intermediate value of selection (e.g., $s = 10^{-2}$ in the top right plot) appears to interpolate between the short-term pseudo-equilibration for strong selection and the long-term equilibration for weak selection, maximizing at a later time when the strong selection curves have begun to decay. The weak selection curves are again well-described for sufficiently rapid exponential growth ($\Gamma = 10^{-2}$), including the late-time equilibration where the rate of decay of the fourth cumulant matches that of the square of the second cumulant. Note that very small initial population size can limit estimation of the skew and excess kurtosis in simulations due to discretization (also seen for constant population size in Figures S9 and S12); this effect is minimal under rapid growth ($\Gamma \geq 10^{-2}$).

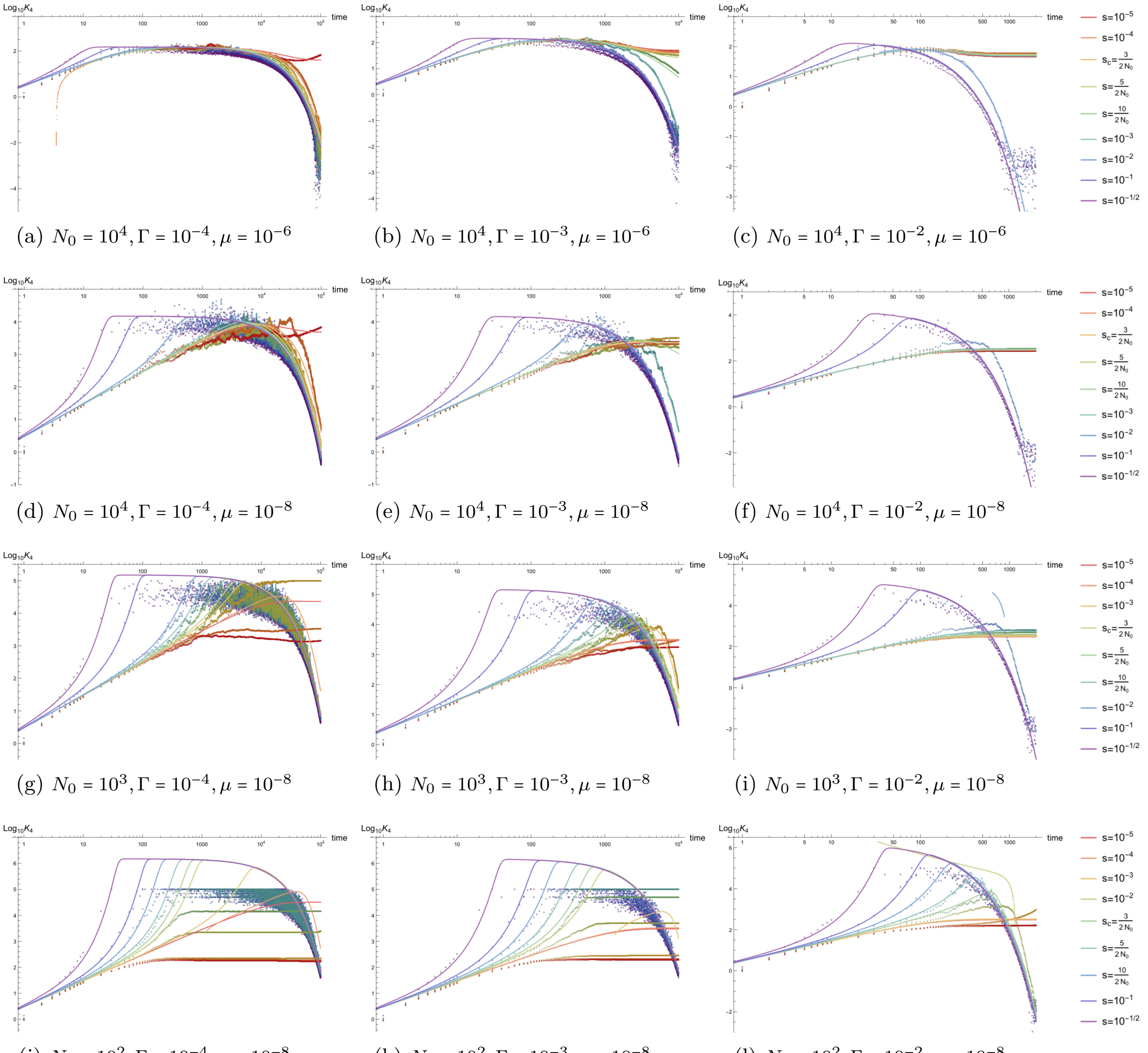


(a) $N_0 = 10^4, \Gamma = 10^{-4}, \mu = 10^{-6}$ (b) $N_0 = 10^4, \Gamma = 10^{-3}, \mu = 10^{-6}$ (c) $N_0 = 10^4, \Gamma = 10^{-2}, \mu = 10^{-6}$

(d) $N_0 = 10^4, \Gamma = 10^{-4}, \mu = 10^{-8}$ (e) $N_0 = 10^4, \Gamma = 10^{-3}, \mu = 10^{-8}$ (f) $N_0 = 10^4, \Gamma = 10^{-2}, \mu = 10^{-8}$

(g) $N_0 = 10^3, \Gamma = 10^{-4}, \mu = 10^{-8}$ (h) $N_0 = 10^3, \Gamma = 10^{-3}, \mu = 10^{-8}$ (i) $N_0 = 10^3, \Gamma = 10^{-2}, \mu = 10^{-8}$

(j) $N_0 = 10^2, \Gamma = 10^{-4}, \mu = 10^{-8}$ (k) $N_0 = 10^2, \Gamma = 10^{-3}, \mu = 10^{-8}$ (l) $N_0 = 10^2, \Gamma = 10^{-2}, \mu = 10^{-8}$

Figure S23: *Time dependence of the excess kurtosis of the allele frequency probability distribution $K_4(t)$ in an exponentially growing population.* Parameter combinations correspond to those shown in Figure S15.

#### S4.3 A population bottleneck and re-expansion

The effects of a temporary population bottleneck range from very subtle to extreme depending on the difference in drift due to the reduced population size. The parameter controlling the response of variation under selection to a bottleneck is the intensity of the bottleneck $I_b$, which is proportional to both the strength of drift during a bottleneck (i.e., the drift parameter $1/2N_b$) and the bottleneck duration $T_b$ (see, e.g., [5] for a discussion of the role of $I_b$ on the mean frequency).

$$I_b \equiv \frac{T_b}{2N_b} \tag{S79}$$

Bottlenecks with low intensity, i.e., short bottlenecks with a relatively small decrease in population size from the initial, pre-bottleneck size, represent a scenario where the bottleneck provides a *small* perturbation to the equilibrium frequency distribution [5]. On the other hand, if a bottleneck imposes a large change in population size (e.g., a two order of magnitude difference in population size between $N_b$ and the initial size $N_0$), and is prolonged relative to the drift constant (i.e., $T_b \gg 2N_b$ such that $I_b \gg 1$), the population has sufficient time (i.e., experiences sufficient drift) to equilibrate to the new population size $N_b$. Two examples of the latter are shown in the plots below, with $N_b = 10^2$ and $T_b = 10^3$ such that the bottleneck intensity $I_b = 10$. Additionally, four examples of a near-borderline case $I_b = 1/2$ for the well-behaved approximation appear below, with parameter combinations $N_b = 10^2, T_b = 10^2$ and $N_b = 10^3, T_b = 10^3$ to illustrate the regime of validity of the analytic expressions in the manuscript.

Since the analytic approximations provided in this manuscript require $2N(t)s' \gg 1$ to be valid descriptions of the non-equilibrium dynamics, selection must remain strong within the bottleneck (i.e., $2N_b s' \gg 1$) if the bottleneck intensity is very strong. As a result, some of the plots below (namely those with $N_b = 10^2$) show large deviations from the field theoretic expectations (including for the mean frequency) for many selection coefficients, especially when the bottleneck intensity is large $I_b \geq 1$. Additionally, even for small bottleneck sizes, if the intensity remains very low ($I_b \ll 1$ where $T_b \ll 2N_b$ even if $2N_b \ll 2N_0$), the approximations remain valid due to the perturbative effects of drift. Due to the subtlety of the effects on various moments, I have chosen to show all bottleneck plots with mutation rates $\mu = 10^{-6}$ and $\mu_b = 10^{-8}$ to limit the variance around each curve. As discussed above, decreasing the mutation rate (e.g., $\mu = 10^{-6} \to 10^{-8}$) increases the relative noise and reduces the equilibrium values (e.g., the mean $\mu/s'$ is reduced if $\mu$ is lower). The plots below, therefore, provide clearer expected values for each cumulant and moment. Since there are three demographic parameters for the bottleneck demography shown in S5, the plots below show all combinations of two values of the bottleneck population size $N_b \in \{10^2, 10^3\}$ (impacting the strength of drift relative to selection and mutation), the duration of the bottleneck $T_b \in \{10^2, 10^3\}$ (impacting the bottleneck intensity), and the time when the bottleneck begins $T_i \in \{10^3, 10^4\}$ (impacting where in the equilibration process of the initial population parameters the bottleneck begins), all while keeping the initial population size $N_0 = 10^4$ fixed. As with the previous plots, the selection coefficients shown range from $s = 10^{-1/2}$ to $s = 10^{-5}$, as well as demography-specific values $2N_b s = \{3, 5, 10\}$ to illustrate the breakdown of the strong selection approximation. Note that when $T_i = 10^3$ (left column of all plots below), the bottleneck occurs during the equilibration process of all but the strongest selection coefficients (i.e., for selection coefficients of roughly $s \gtrsim 10^{-2}$), which impacts both the equilibration itself and the response of the population to the bottleneck demography. In contrast, a bottleneck beginning at generation $T_i = 10^4$ occurs when a slightly larger range of selection coefficients have already reached equilibrium ($T_i > t_{\text{eq}}$). As with the previous simulations, moments and cumulants are computed as an average of $L = 10^5$ independent sites evolving through the same demography.

As a final note on the bottleneck plots in this section, some of the plots below show a breakdown in the numerical plotting software (Mathematica [6]) when plotting analytic expressions for cumulants or moments in the presence of a population bottleneck in logarithmic space; the breakdown occurs for a small number of generations immediately after the bottleneck. This is due to near-discontinuities associated with the sharp slope immediately after re-expansion from an *instantaneous* bottleneck (i.e., the square bottleneck) The nearly piecewise nature of the results can be difficult plot for strong selection coefficients, particularly when the bottleneck drift constant $1/2N_b$ is relatively large (e.g., on the order of $\frac{1}{2} \times 10^{-2}$ for $N_b = 10^2$).

However, this appears to only be an issue when plotting bottlenecks at late time $T_i = 10^4$ in logarithmic space. To show that the resulting expressions are indeed smooth, except at $t = T_i$ and $t = T_i + T_b$, and that this is only a plotting error, one can plot the temporal dependence of the same moment starting from a time near immediately before the beginning of the bottleneck. One example of such a discontinuity, appearing in Figure S24a for the time-dependence of the variance of the allele frequency probability distribution over time $t \in (0, 10^4]$ as a diagonal line immediately after re-expansion for $s = 10/2N_b$ (lighter blue curve), is shown below.

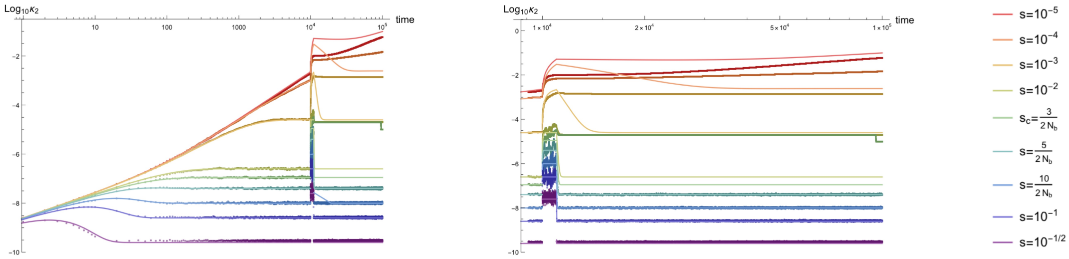


(a) Variance $\kappa_2$ for $N_b = 10^2, T_b = 10^3, T_i = 10^4$ (b) $\kappa_2$ with same parameters starting at $t = 9990$

Figure S24: *Numerical plotting error for strong selection in a severe bottleneck,* The variance of the allele frequency probability distribution is shown for a bottleneck demography with parameters (a) $N = 10^4, N_b = 10^2, T_b = 10^2, T_i = 10^4, \mu = 10^{-6}$ and (b) $\mu_b = 10^{-8}$. (a): The plotting error occurs immediately after re-expansion for the bottleneck; it appears as a diagonal line in the lighter blue curve in plot representing additive purifying selection with selection strength $s = 1/2N_b$. (b): This error is no longer visible if one plots time immediately before, during, and after the bottleneck.

The right plot, Figure S24b, shows the time dependence of the variance with the same parameters, but the plot begins 10 generations before the bottleneck (at $t = 9990$). From this plot, it is clear that the period after re-expansion is nearly flat for $s = 10/2N_b$ due to a swift equilibration, with an equilibrium value of $\kappa_2 = \frac{\lambda'}{2\gamma'}$. This indicates that the diagonal part of the left plot, Figure S24a is simply a numerical plotting error. The same kind of numerical error appears in several plots below for $N_b = 10^2, T_i = 10^4$, but all of these plots should be viewed as effectively flat after re-expansion due to the rapid re-equilibration of strong purifying selection.

##### S4.3.1 Bottleneck demography: time dependence of the population size

The demography associated with a range of parameters for $N_b$, $T_b$, and $T_i$ are shown below, where $N_0 = 10^4$, $\mu = 10^{-6}$, and $\mu_b = 10^{-8}$ remain constant. Results for these demographies are presented in this subsection in the same plot position as the corresponding demography.

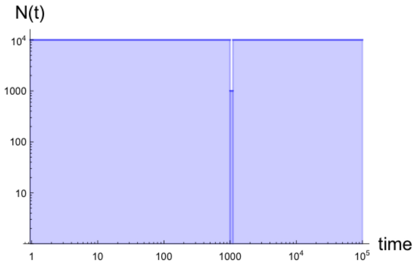


(a) $N_b = 10^3, T_b = 10^2, T_i = 10^3$

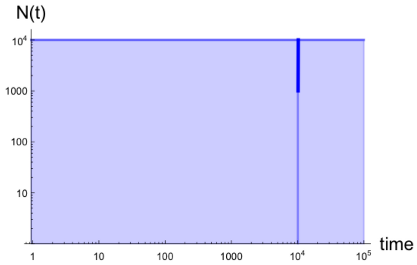


(b) $N_b = 10^3, T_b = 10^2, T_i = 10^4$

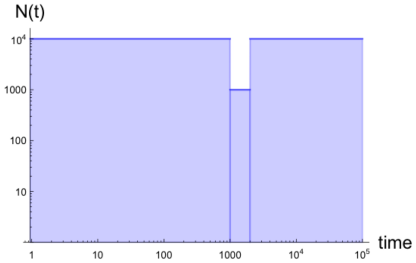


(c) $N_b = 10^3, T_b = 10^3, T_i = 10^3$

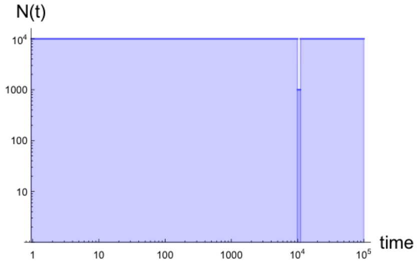


(d) $N_b = 10^3, T_b = 10^3, T_i = 10^4$

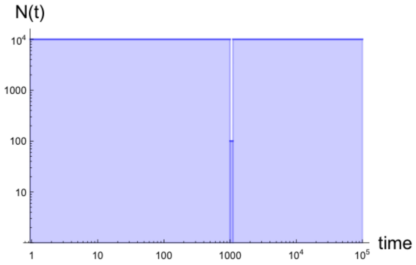


(e) $N_b = 10^2, T_b = 10^2, T_i = 10^3$

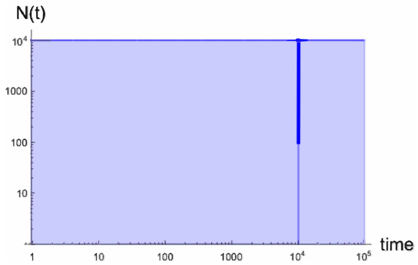


(f) $N_b = 10^2, T_b = 10^2, T_i = 10^4$

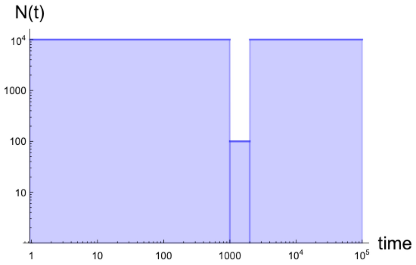


(g) $N_b = 10^2, T_b = 10^3, T_i = 10^3$

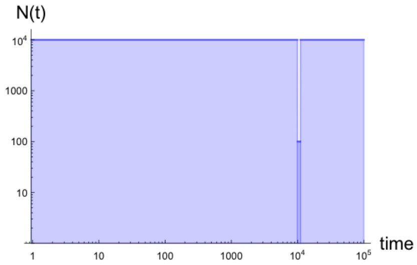


(h) $N_b = 10^2, T_b = 10^3, T_i = 10^4$

Figure S25: *Time dependence of the population size for simulated populations with a population bottleneck and subsequent re-expansion.* Demographies are presented for $N_0 = 10^4$, $N_b = 10^3$ (a–d), $10^2$ (e–h), $T_b = 10^2$ (a,b;e,f), $10^3$ (c,d;g,h), and $T_i = 10^3$ (left), $10^4$ (right). Some plots shown for bottleneck onset $T_i = 10^4$ are difficult to plot in detail in log-log space. Curves in plots (b) and (f) have been thickened to denote the initial time $T_i$ and population size $N_b$ of the bottleneck; the duration in both cases is $T_b = 10^2$.

##### S4.3.2 Bottleneck demography: mean frequency $\overline{p}(t)$ to three-vertex order

Below are plots of the mean of the allele frequency probability distribution $\overline{p}$, shown here with the three-vertex correction that accounts for a small amount of drift on the order of $1/2Ns'$, but breaks down when $1/2N_b \gtrsim 1$. Note that even the deterministic approximation of the mean breaks down at weak selection ($2N_b s \leq 3$) when the bottleneck intensity is large ($I_b \gg 1$ in Figures S26g and S26h).

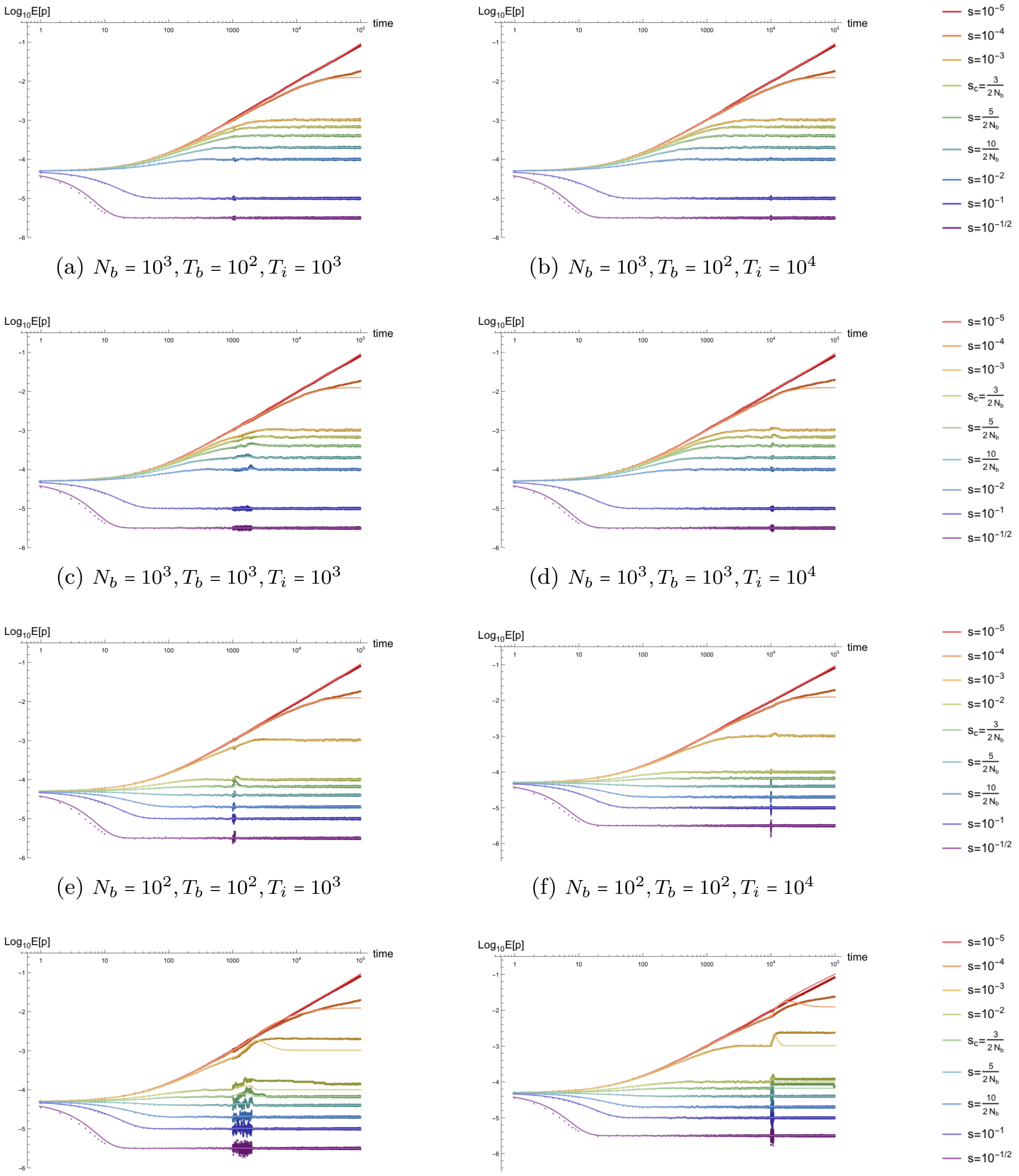


(a) $N_b = 10^3, T_b = 10^2, T_i = 10^3$ (b) $N_b = 10^3, T_b = 10^2, T_i = 10^4$

(c) $N_b = 10^3, T_b = 10^3, T_i = 10^3$ (d) $N_b = 10^3, T_b = 10^3, T_i = 10^4$

(e) $N_b = 10^2, T_b = 10^2, T_i = 10^3$ (f) $N_b = 10^2, T_b = 10^2, T_i = 10^4$

(g) $N_b = 10^2, T_b = 10^3, T_i = 10^3$ (h) $N_b = 10^2, T_b = 10^3, T_i = 10^4$

Figure S26: *Time dependence of the mean of the allele frequency probability distribution $\overline{p}$ for simulated populations with a population bottleneck and subsequent re-expansion.* Plots are shown for mutation rates $\mu = 10^{-6}$ and $\mu_b = \mu/100$, initial size $N_0 = 10^4$, bottleneck size $N_b = 10^3$ (top two rows), $10^2$ (bottom two rows), duration $T_b = 10^2$ (a,b;e,f), $10^3$ (c,d;g,h), bottleneck onset $T_i = 10^3$(left), $10^4$ (right), and selection coefficients $s \in [10^{-5}, 10^{-1/2}]$.

##### S4.3.3 Bottleneck demography: variance $\kappa_2(t)$ to two vertex order

Figure S27 shows plots of the two-vertex approximation of the variance of the allele frequency distribution $\kappa_2$. The variance breaks down for high intensity bottlenecks (Figures S27g and S27h) because the variance equilibrates to the bottleneck population size $N_b$ and does not immediately decay for selection coefficients at or below $2N_bs = 3$. Note that, even for bottlenecks with intensity $I_b = 1/2$, the analytic approximation remains accurate until $2N_0s \sim 3$, indicating that the bottleneck is a perturbative effect and the regime of validity is determined by the initial population size.

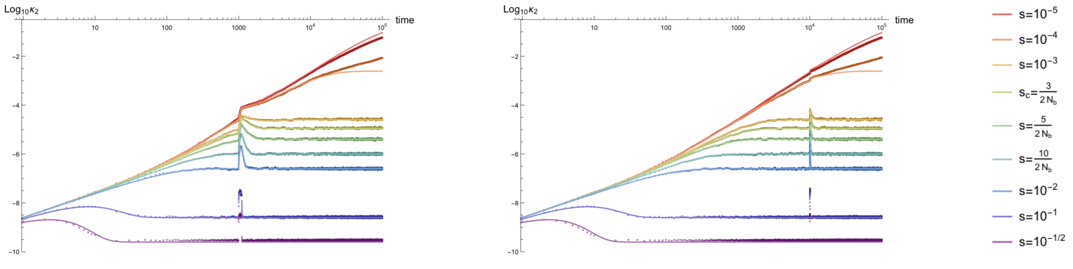


(a) $N_b = 10^3, T_b = 10^2, T_i = 10^3$ (b) $N_b = 10^3, T_b = 10^2, T_i = 10^4$

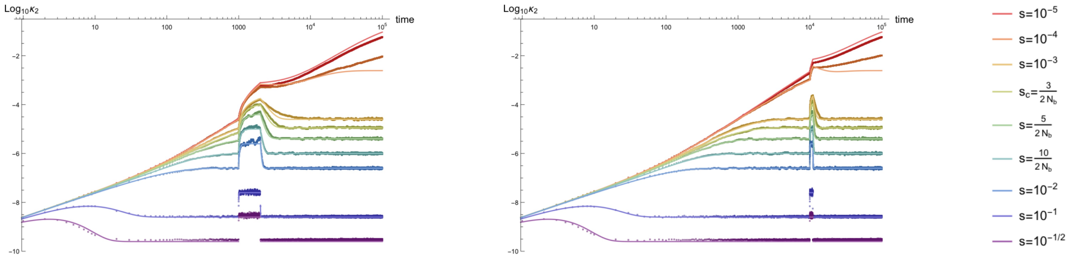


(c) $N_b = 10^3, T_b = 10^3, T_i = 10^3$ (d) $N_b = 10^3, T_b = 10^3, T_i = 10^4$

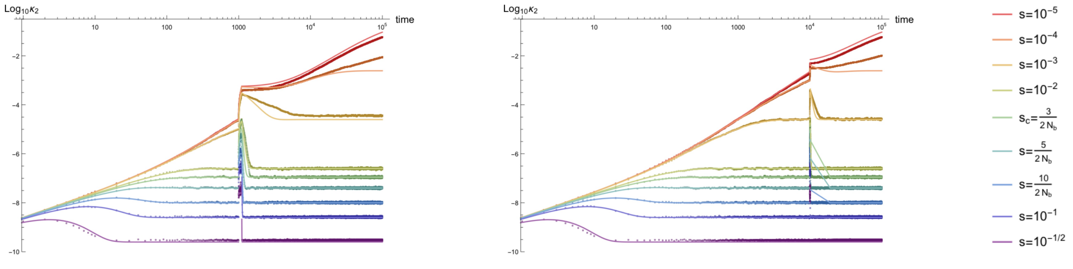


(e) $N_b = 10^2, T_b = 10^2, T_i = 10^3$ (f) $N_b = 10^2, T_b = 10^2, T_i = 10^4$

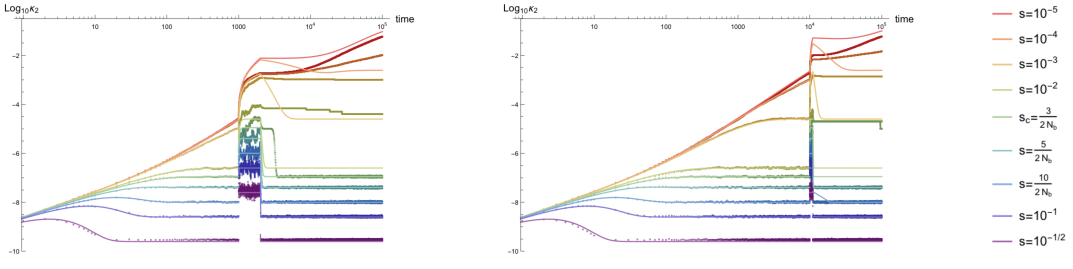


(g) $N_b = 10^2, T_b = 10^3, T_i = 10^3$ (h) $N_b = 10^2, T_b = 10^3, T_i = 10^4$

Figure S27: *Time dependence of variance of the allele frequency probability distribution $\kappa_2$for simulated populations with a population bottleneck and subsequent re-expansion.* Parameter combinations correspond to those shown in Figure S26.

##### S4.3.4 Bottleneck demography: homozygosity $E[p^2(t)]$ to two-vertex order

The two-vertex approximation for the homozygosity $E[p^2]$ is plotted below for an allele experiencing a population bottleneck and re-expansion. The dynamics are largely the same as for the variance, above, including the regime of validity of the analytic approximation. Note that, in both the variance and homozygosity, there is a numerical breakdown in the plotting software (appearing as diagonal lines after re-expansion) for plots with $N_b = 10^2, T_i = 10^4$ (Figures S28f and S28h) immediately after the bottleneck, but the analytic approximations match simulations before, during, and after the bottleneck (after numerical issues).

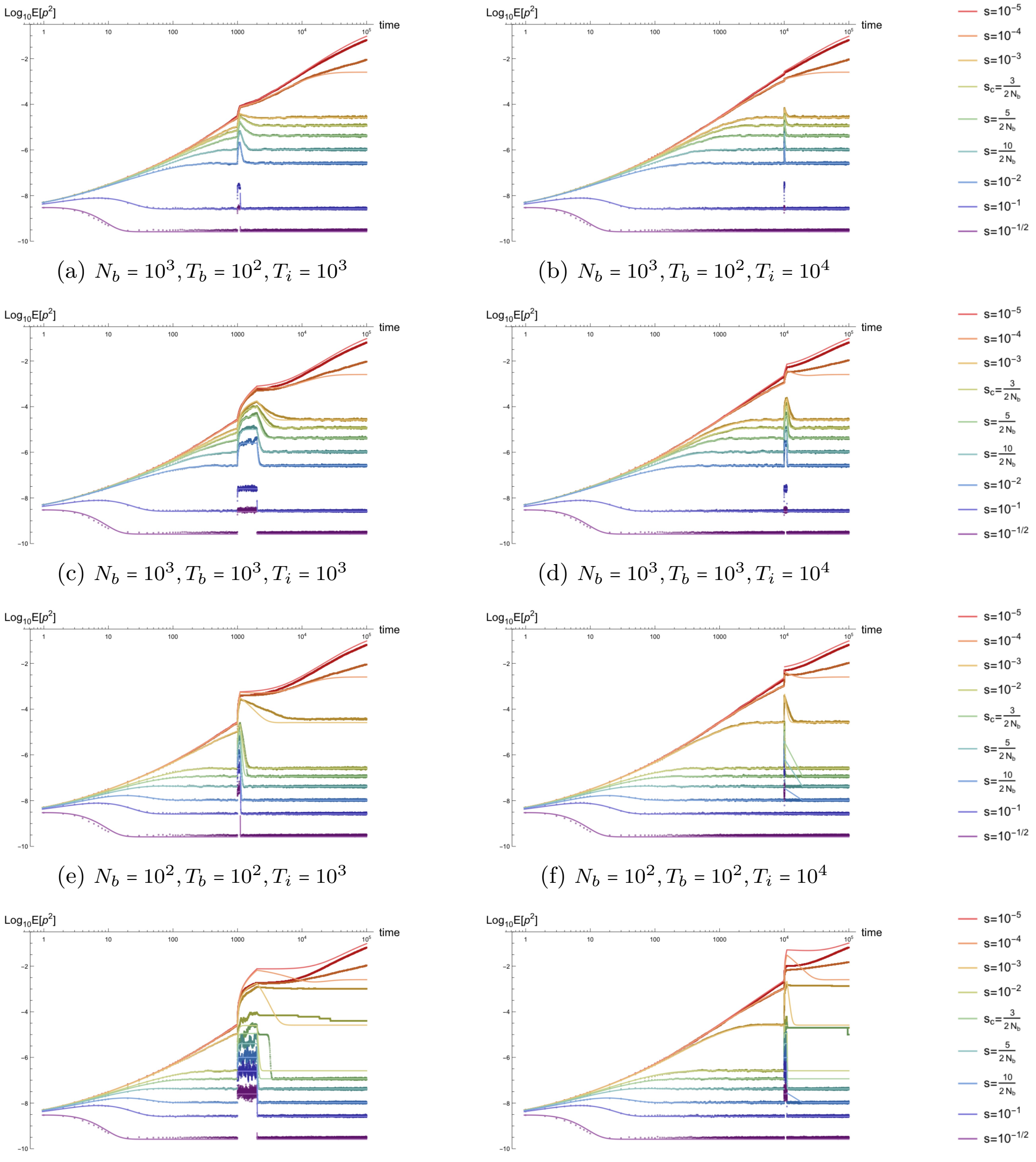


(a) $N_b = 10^3, T_b = 10^2, T_i = 10^3$

(b) $N_b = 10^3, T_b = 10^2, T_i = 10^4$

(c) $N_b = 10^3, T_b = 10^3, T_i = 10^3$

(d) $N_b = 10^3, T_b = 10^3, T_i = 10^4$

(e) $N_b = 10^2, T_b = 10^2, T_i = 10^3$

(f) $N_b = 10^2, T_b = 10^2, T_i = 10^4$

(g) $N_b = 10^2, T_b = 10^3, T_i = 10^3$

(h) $N_b = 10^2, T_b = 10^3, T_i = 10^4$

Figure S28: *Time dependence of the second non-central moment of the allele frequency probability distribution, the homozygosity $E[p^2]$, for simulated populations with a population bottleneck and subsequent re-expansion.* Parameter combinations correspond to those shown in Figure S26.

##### S4.3.5 Bottleneck demography: third cumulant $\kappa_3(t)$ to three-vertex order

The lowest-vertex order $(3V)$ approximation to the third cumulant of the allele frequency distribution $\kappa_3$ is plotted below. The dynamics are largely controlled by the variance and the regime of validity is the same. Note that, as for the second cumulant and moments, if the bottleneck has sufficiently large intensity, the population equilibrates within the bottleneck for all selection coefficients, and is slow to decay after re-expansion when selection is weak (roughly when $2N_0 s \lesssim 3$).

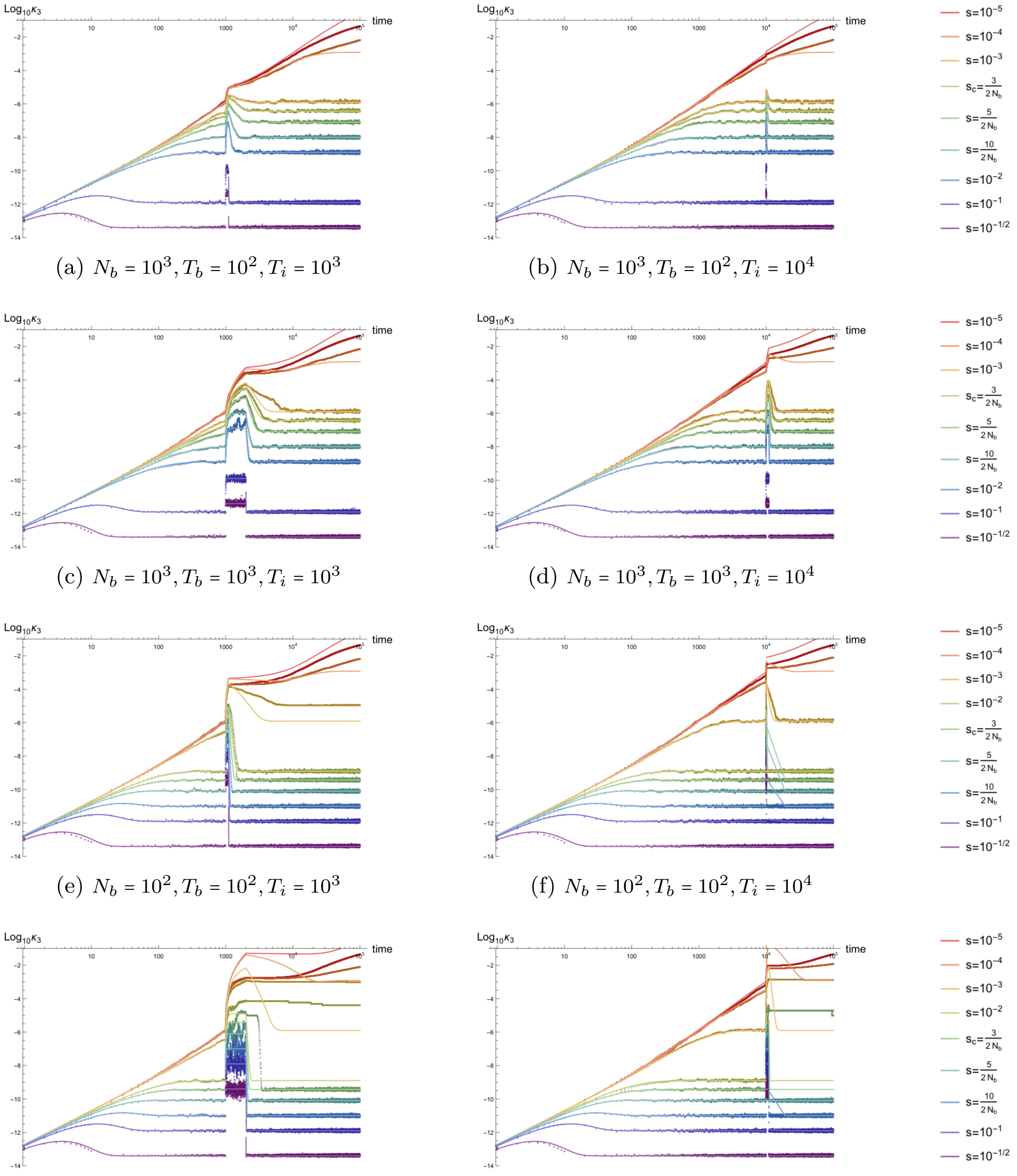


(a) $N_b = 10^3, T_b = 10^2, T_i = 10^3$ (b) $N_b = 10^3, T_b = 10^2, T_i = 10^4$

(c) $N_b = 10^3, T_b = 10^3, T_i = 10^3$ (d) $N_b = 10^3, T_b = 10^3, T_i = 10^4$

(e) $N_b = 10^2, T_b = 10^2, T_i = 10^3$ (f) $N_b = 10^2, T_b = 10^2, T_i = 10^4$

(g) $N_b = 10^2, T_b = 10^3, T_i = 10^3$ (h) $N_b = 10^2, T_b = 10^3, T_i = 10^4$

Figure S29: *Time dependence of the third cumulant of the allele frequency probability distribution $\kappa_3$ for simulated populations with a population bottleneck and subsequent re-expansion.* Parameter combinations correspond to those shown in Figure S26.

##### S4.3.6 Bottleneck demography: third non-central moment $E[p^3(t)]$ to three-vertex order

Plots of the third non-central moment of the allele frequency distribution $E[p^3]$ are shown below for the same range of bottleneck demographies. Again, the high intensity bottlenecks shown in the last row of plots deviate substantially from the analytic approximation for strong selection, but only when $2N_b s \leq 3$.

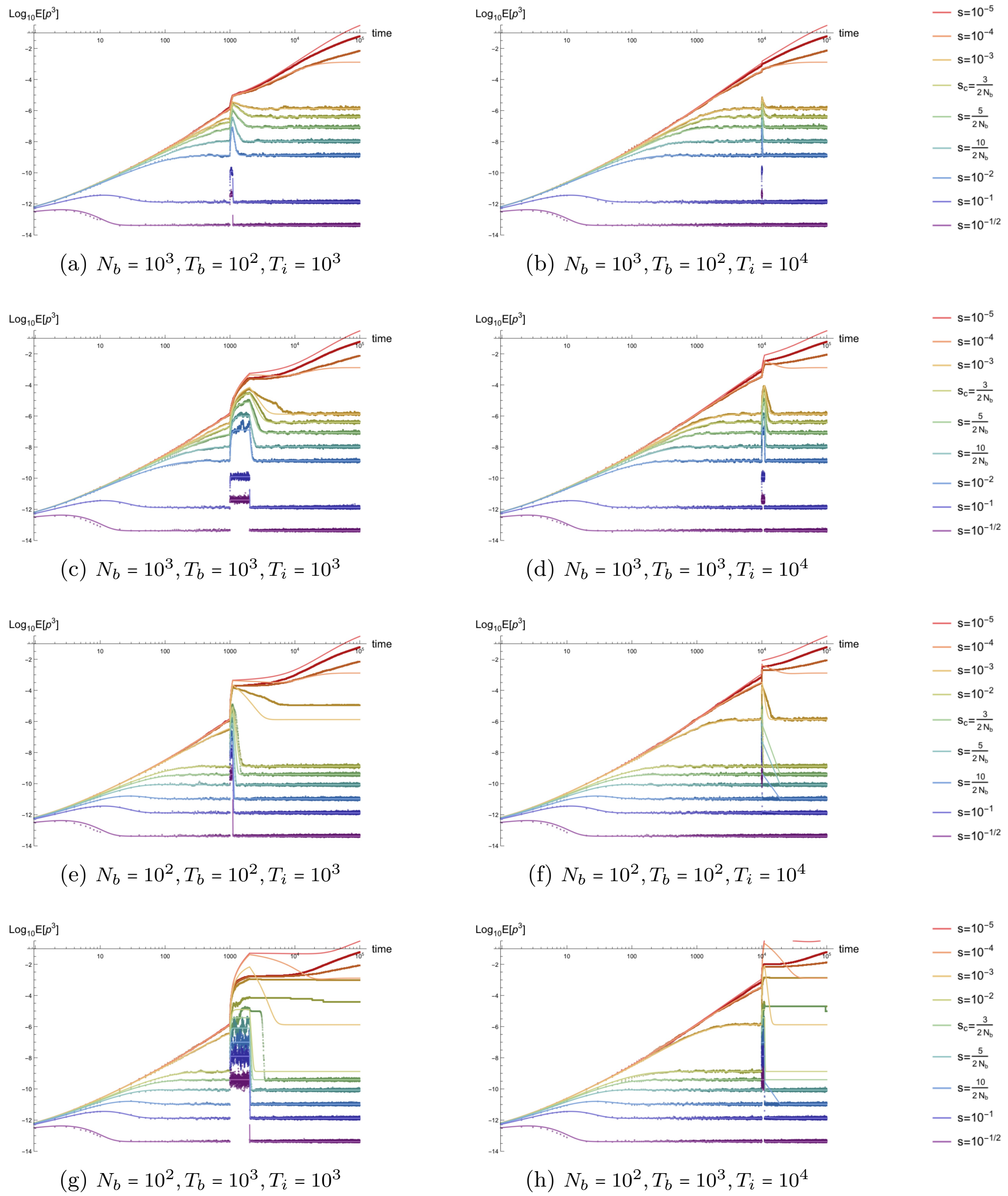


(a) $N_b = 10^3, T_b = 10^2, T_i = 10^3$

(b) $N_b = 10^3, T_b = 10^2, T_i = 10^4$

(c) $N_b = 10^3, T_b = 10^3, T_i = 10^3$

(d) $N_b = 10^3, T_b = 10^3, T_i = 10^4$

(e) $N_b = 10^2, T_b = 10^2, T_i = 10^3$

(f) $N_b = 10^2, T_b = 10^2, T_i = 10^4$

(g) $N_b = 10^2, T_b = 10^3, T_i = 10^3$

(h) $N_b = 10^2, T_b = 10^3, T_i = 10^4$

Figure S30: *Time dependence of the third non-central moment of the allele frequency probability distribution $E[p^3]$ for simulated populations with a population bottleneck and subsequent re-expansion.* Parameter combinations correspond to those shown in Figure S26.

##### S4.3.7 Bottleneck demography: skew $K_3(t)$ to three-vertex order

Unlike the previous moments, plotted above, the skew of the allele frequency probability distribution shows some deviation from the analytic expectation for less severe bottlenecks with $N_b = 10^3$. However, this happens at and below roughly $2N_b s = 3$, where the strong selection approximation is expected to fail. The contrast exists only because the previously plotted analytics aligned to simulated moments and cumulants for $N_b = 10^3$ beyond the expected range of validity. The effect of the bottleneck is to increase the skew to a new value of $1/\sqrt{N_b \mu}$, which is sustained for bottlenecks with high intensity.

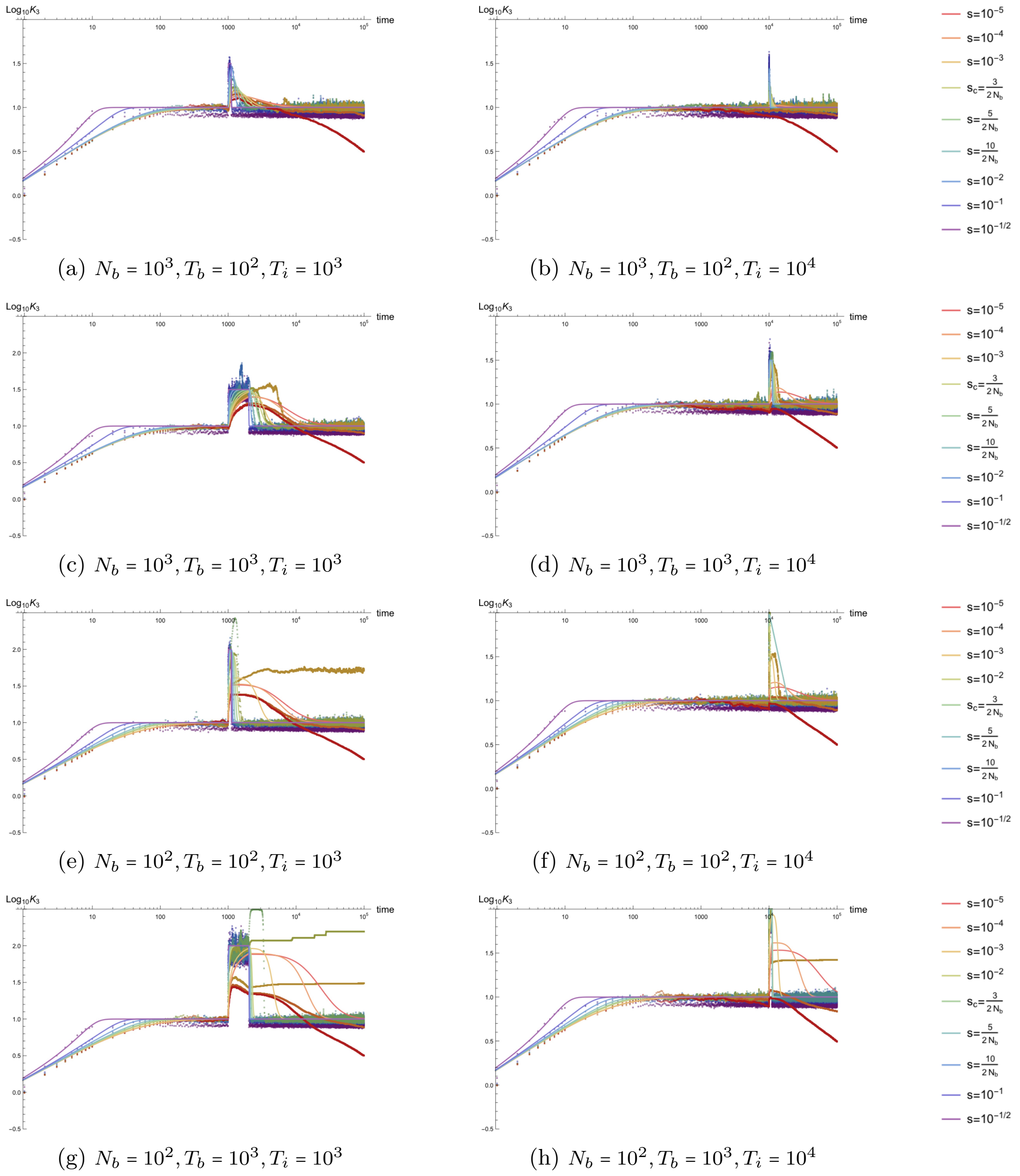


(a) $N_b = 10^3, T_b = 10^2, T_i = 10^3$ (b) $N_b = 10^3, T_b = 10^2, T_i = 10^4$

(c) $N_b = 10^3, T_b = 10^3, T_i = 10^3$ (d) $N_b = 10^3, T_b = 10^3, T_i = 10^4$

(e) $N_b = 10^2, T_b = 10^2, T_i = 10^3$ (f) $N_b = 10^2, T_b = 10^2, T_i = 10^4$

(g) $N_b = 10^2, T_b = 10^3, T_i = 10^3$ (h) $N_b = 10^2, T_b = 10^3, T_i = 10^4$

Figure S31: *Time dependence of the skew of the allele frequency probability distribution $K_4$ for simulated populations with a population bottleneck and subsequent re-expansion.* Parameter combinations correspond to those shown in Figure S26.

##### S4.3.8 Bottleneck demography: fourth cumulant $\kappa_4(t)$ to four-vertex order

The fourth cumulant of the allele frequency probability distribution $\kappa_4$ is plotted below. This is again well-described by the analytic approximations, sans numerical issues, in the first three rows of demographies. In the last row, the approximation again breaks down for selection coefficients below $2N_b s \sim 5$.

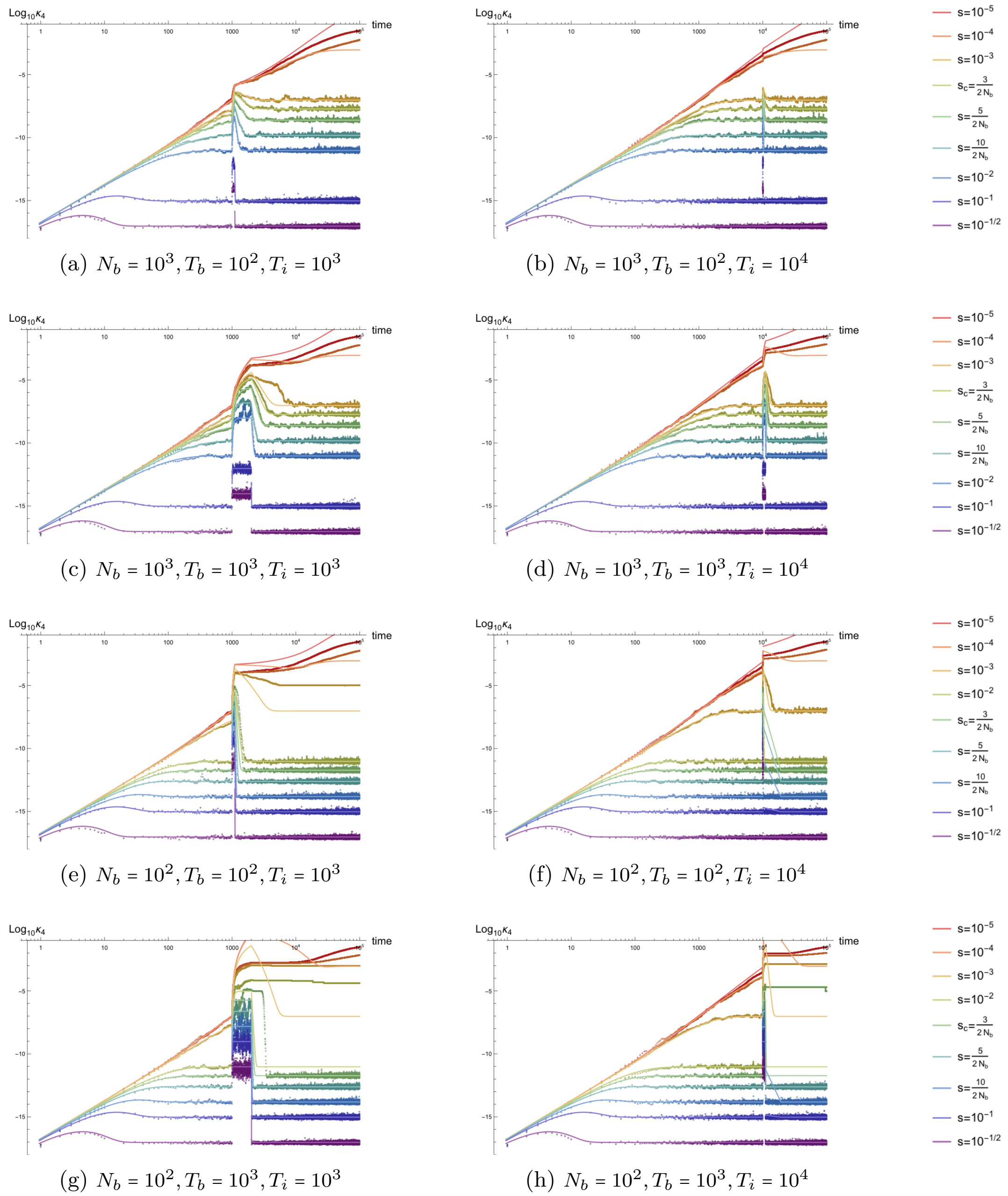


(a) $N_b = 10^3, T_b = 10^2, T_i = 10^3$

(b) $N_b = 10^3, T_b = 10^2, T_i = 10^4$

(c) $N_b = 10^3, T_b = 10^3, T_i = 10^3$

(d) $N_b = 10^3, T_b = 10^3, T_i = 10^4$

(e) $N_b = 10^2, T_b = 10^2, T_i = 10^3$

(f) $N_b = 10^2, T_b = 10^2, T_i = 10^4$

(g) $N_b = 10^2, T_b = 10^3, T_i = 10^3$

(h) $N_b = 10^2, T_b = 10^3, T_i = 10^4$

Figure S32: *Time dependence of the fourth cumulant of the allele frequency probability distribution $\kappa_4$ for simulated populations with a population bottleneck and subsequent re-expansion.* Parameter combinations correspond to those shown in Figure S26.

##### S4.3.9 Bottleneck demography: fourth non-central moment $E[p^4(t)]$ to four-vertex order

The fourth non-central moment $E[p^4]$ is again very similar to the corresponding cumulant, $\kappa_4$.

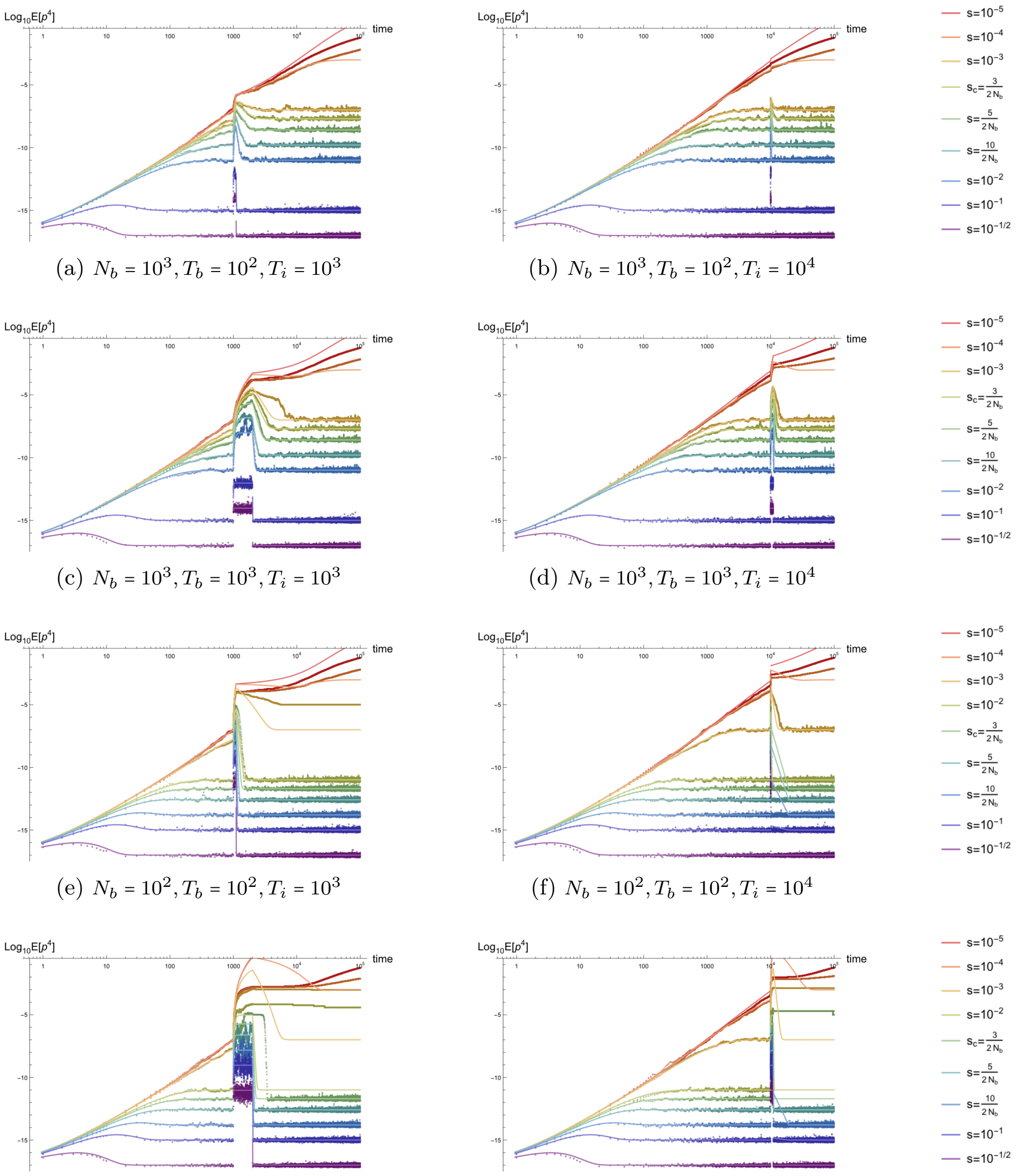


(a) $N_b = 10^3, T_b = 10^2, T_i = 10^3$

(b) $N_b = 10^3, T_b = 10^2, T_i = 10^4$

(c) $N_b = 10^3, T_b = 10^3, T_i = 10^3$

(d) $N_b = 10^3, T_b = 10^3, T_i = 10^4$

(e) $N_b = 10^2, T_b = 10^2, T_i = 10^3$

(f) $N_b = 10^2, T_b = 10^2, T_i = 10^4$

(g) $N_b = 10^2, T_b = 10^3, T_i = 10^3$

(h) $N_b = 10^2, T_b = 10^3, T_i = 10^4$

Figure S33: *Time dependence of the fourth non-central moment of the allele frequency probability distribution $\overline{p}$ for simulated populations with a population bottleneck and subsequent re-expansion.* Parameter combinations correspond to those shown in Figure S26.

##### S4.3.10 Bottleneck demography: excess kurtosis $K_4(t)$ to four-vertex order

The excess kurtosis of the allele frequency probability distribution, like the skew, approaches a new value of $3/2N_b\mu$ during the bottleneck, which independent of selection coefficients provided $2N_b s \gtrsim 5$. This indicates that the tails of the distribution temporarily have more mass such that, even for highly deleterious alleles, the distribution may harbor variation at higher frequency than in the initial population.

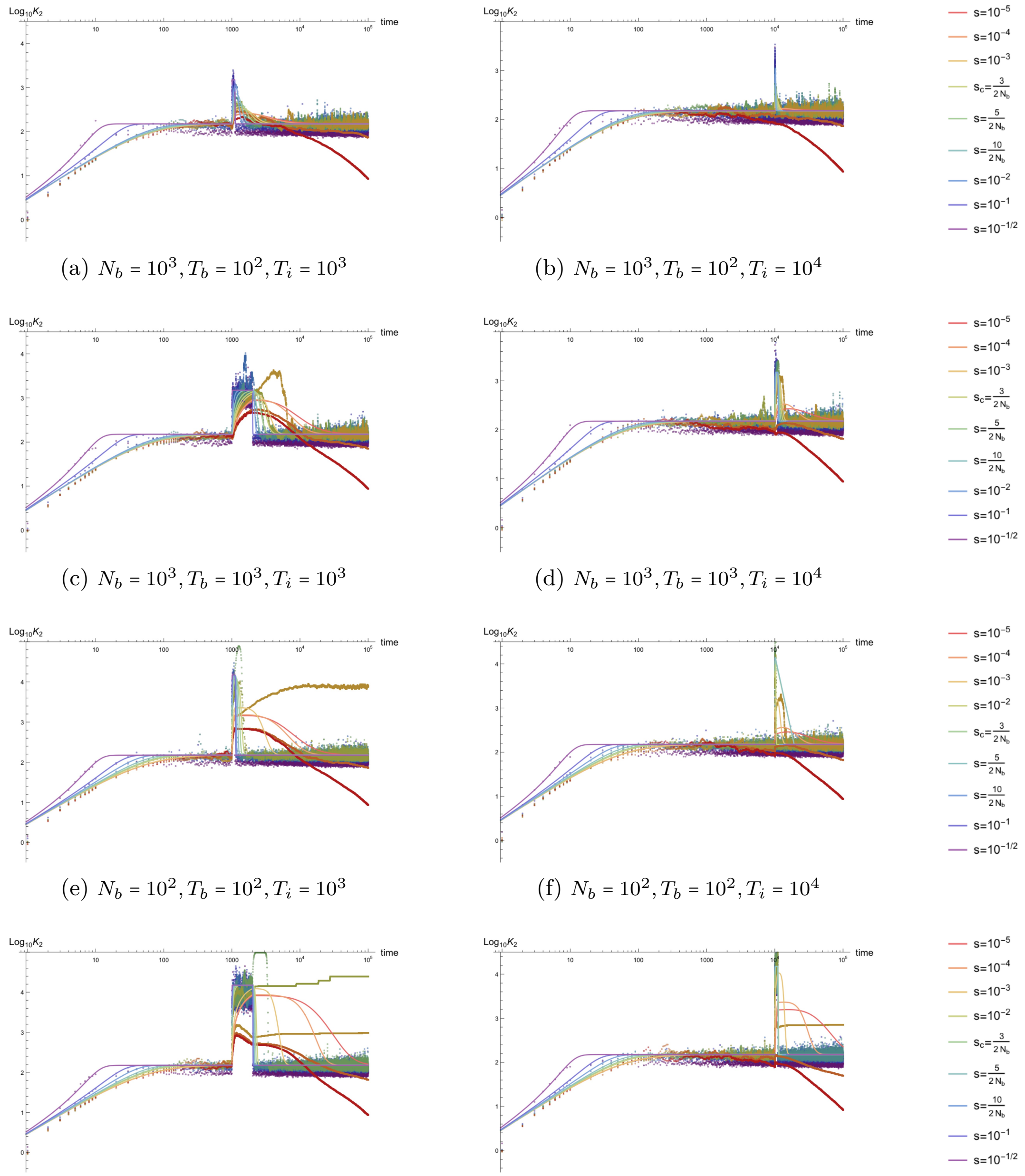


(a) $N_b = 10^3, T_b = 10^2, T_i = 10^3$
(b) $N_b = 10^3, T_b = 10^2, T_i = 10^4$
(c) $N_b = 10^3, T_b = 10^3, T_i = 10^3$
(d) $N_b = 10^3, T_b = 10^3, T_i = 10^4$
(e) $N_b = 10^2, T_b = 10^2, T_i = 10^3$
(f) $N_b = 10^2, T_b = 10^2, T_i = 10^4$
(g) $N_b = 10^2, T_b = 10^3, T_i = 10^3$
(h) $N_b = 10^2, T_b = 10^3, T_i = 10^4$

Figure S34: *Time dependence of the excess kurtosis of the allele frequency probability distribution $K_4$ for simulated populations with a population bottleneck and subsequent re-expansion.* Parameter combinations correspond to those shown in Figure S26.

#### S4.4 Cyclical demography with oscillatory population size

Cyclical populations are comparatively less well studied than more ubiquitous and transient features like exponential growth or population bottlenecks. However, such demographies are related to both rapid population growth and rapid decline; populations with regular, periodic oscillations in the number of individuals can be viewed as experiencing serial bottlenecks (see, e.g., [7–9]) at regular time intervals with subsequent re-expansion after each bottleneck. In this section, comparisons between simulations and the PGFT-derived closed form analytic approximations are shown for one class of cyclical demographies defined by the deterministic time-dependent oscillatory function in Equation S6 for $N_{\text{osc}}(t)$ diploid individuals. This smooth, continuous parameterization alleviates some minor lingering effects from the discontinuous Heaviside functions used to model a square bottleneck (see $N_{\text{BN}}(t)$ in Equation S5), such as the numerical plotting errors seen in Figure S24.

The cyclical diploid population size $N_{\text{osc}}(t)$ is parameterized by three constants: the maximum population size $N_{\min}$, the minimum size $N_{\min}$, and the period of oscillation $\tau$. The cosine dependence in $N_{\text{osc}}(t)$ describes populations that are initialized with the maximum number of individuals $N(t_0 = 0) = N_{\max}$ and return to this size every $\tau$ generations thereafter (i.e., $N(t = n\tau) = N_{\max}$ for all integer values of $n$) and reach the minimum population size $N_{\min}$ halfway between each maximum value (i.e., at each time point $t = \tau/2 + n\tau$ for all integer values of $n$). Importantly, this parameterization only meant to serve as one class of examples that aids in evaluating the performance of the analytic approximations; the choice of initial conditions is tunable and not intended to hold general biological significance (e.g., replacing the cosine with a sine function such that $N(0) = N_{\min}$ results in more intense drift at early times and potentially different transient dynamics) and, if desired, a phase parameter can be introduced by taking $\varphi(t) \to \varphi(t) + const.$ in Equation S6 to generalize the initialization condition. As with the square bottleneck, I have chosen to reduce the parameter space in this section by fixing the initial value of $N_{\max} = 10^4$, which limits the number of plots presented. In each figure, results are shown for two values of $N_{\min} = N_{\max}/2 = 5 \times 10^3$ and $N_{\min} = N_{\max}/10 = 10^3$, which describe moderate and significant oscillation amplitudes, respectively. The period is varied over three decades: results are shown for oscillations lasting $\tau = 10^2$, $10^3$, or $10^4$ generations. Additionally, results are shown for mutation rates $\mu = 10^{-8}$ and $10^{-6}$ (again fixing $\mu_b = \mu/100$ to limit the parameter space). Each moment and cumulant is computed by averaging over $L = 10^5$ simulated sites. In the figures below, each subfigure represents one parameter combination $(N_{\min}, \tau, \mu)$ with fixed values $(N_{\max} = 10^4, \mu_b = \frac{\mu}{100}, L = 10^5)$ and shows results for a range of deleterious selection coefficients $s \in [10^{-5}, 10^{-1/2}]$.

As discussed in the previous sections, a lower mutation rate introduces comparatively more noise for a fixed target size $L$. Accordingly, the $\mu = 10^{-6}$ simulations in the following figures provide the clearest comparisons between simulated and theoretical results. In this section, I have chosen to show one row of plots (three parameter combinations) with the lower mutation rate ($\mu = 10^{-8}$) because, unlike the constant size and exponential demographies, the subtlety of some of the oscillatory dynamics can be almost entirely masked when the noise is increased. This suggests that the detection of oscillations in the population size from empirical data may be power-limited in some parameter regimes, resulting in the potential misspecification of the underlying population model unless an observable can be found that is both sensitive to oscillations in the population size and robust to noise due to low target size (or summarizes data from $\gtrsim 10^6$ sites).

Finally, as detailed in the manuscript, the late time asymptotic dynamics of the allele frequency probability distribution under cyclical population size are not quite those of steady state mutation-selection-drift balance. instead, for some parameter combinations, the distribution can *oscillate* around a shape determined by $\overline{N}_{\text{osc}}$, the effective population size (harmonically) averaged over one period. As one might expect, the allele frequency probability distribution expands and contracts on a time-scale determined by $\tau$ in a non-equilibrium waltz, and all cumulants above the mean ($\kappa_n$ for $n \geq 2$) respond in turn. To identify the regime of validity of the analytic approximations for each moment, simulations were performed using selection coefficients $2\overline{N}_{\text{osc}}s = 3, 5, 10$ (in the regime where $s' \approx s$), as this is the appropriate scale that emerges from the asymptotic time-dependence of the variance (see manuscript text). Perhaps surprisingly, the speed of oscillations does not appear to affect the accuracy of the PGFT approximations (the analytic time-dependence fully describes the transition to a coarse-grained regime under rapid oscillations with $s'\tau \ll 1$), which faithfully track the dynamics while selection remains strong and break down roughly when $s \gtrsim 5/2\overline{N}_{\text{osc}}$. This provides a satisfying and intuitive analogy to the regime of validity for the dynamics of a population with constant size. Note that, as was the case for all previously discussed demographies, each cyclical demography plot is presented in logarithmic time. This may be particularly unintuitive for this demography, as the regular periodicity is obscured. However, this allows for better assessment of the impact of the oscillatory behavior on the transient dynamics at early times (i.e., prior to the waltz), which are much less intuitive than the resulting asymptotic dynamics, but appear to be theoretically well-described in the same regime of validity.

##### S4.4.1 Cyclical demography: time dependence of the population size

The following log-log plots show the time dependence of the population size in a cyclical populations spanning a range of parameters. These populations experience regular, periodic oscillations in size of varying amplitudes and periods. Analytic and simulation results in the following section are presented in the same order as the time-dependent population sizes in Figure S35. Each demography begins at an initial size of $N(0) = N_{\max} = 10^4$ diploid individuals and returns to this size at each multiple of $\tau$ generations; simulations are presented for periods of $\tau = 10^4$, $10^3$, and $10^2$ generations, shown from left to right. The minimum population size occurs at time $\tau/2$ (and thereafter at generations $t = n\tau + \tau/2$ for integer values of $n$) and is shown for two minimum population sizes: $N_{\min} = 5 \times 10^3$ ($= N_{\max}/2$) in the top two rows and $N_{\min} = 10^3$ ($N_{\max}/10$) in the bottom row of plots. The first and second rows from the top are identical in the figure below; the first row of all subsequent plots of cumulants and moments will be shown with mutation rates $\mu = 10^{-8}, \mu_b = 10^{-10}$ in the same demography as the second row with $\mu = 10^{-6}, \mu_b = 10^{-8}$. Lower mutation rates are shown for contrast, but obscure the performance of the analytic approximation, as the simulated stochastic noise partially masks the oscillatory behavior.

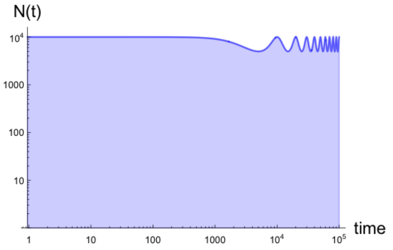


(a) $N_{\max} = 10^4, N_{\min} = 5 \times 10^3, \tau = 10^4$

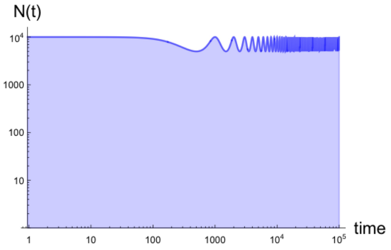


(b) $N_{\max} = 10^4, N_{\min} = 5 \times 10^3, \tau = 10^3$

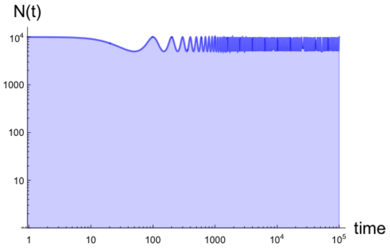


(c) $N_{\max} = 10^4, N_{\min} = 5 \times 10^3, \tau = 10^2$

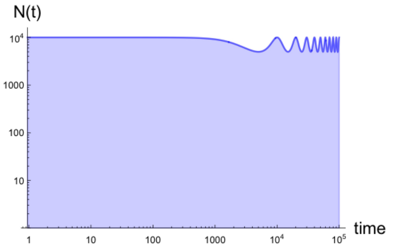


(d) $N_{\max} = 10^4, N_{\min} = 5 \times 10^3, \tau = 10^4$

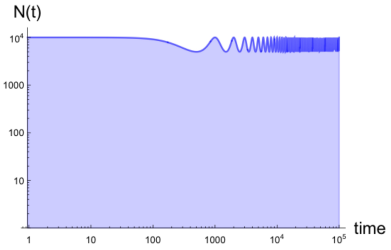


(e) $N_{\max} = 10^4, N_{\min} = 5 \times 10^3, \tau = 10^3$

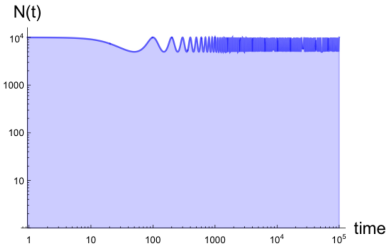


(f) $N_{\max} = 10^4, N_{\min} = 5 \times 10^3, \tau = 10^2$

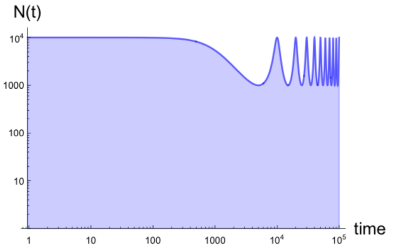


(g) $N_{\max} = 10^4, N_{\min} = 10^3, \tau = 10^4$

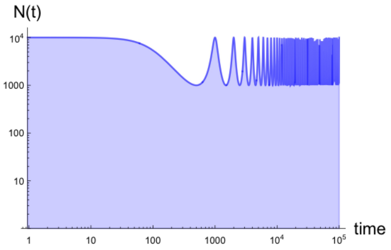


(h) $N_{\max} = 10^4, N_{\min} = 10^3, \tau = 10^3$

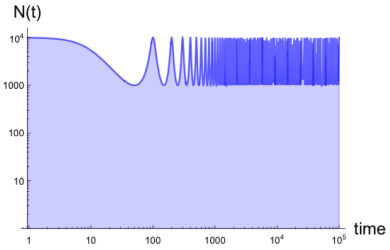


(i) $N_{\max} = 10^4, N_{\min} = 10^3, \tau = 10^2$

Figure S35: *Time dependence of the oscillatory population size for simulated cyclical populations.* Demographies are presented for maximum population size $N_{\max} = 10^4$, minimum size $N_{\min} = 5 \times 10^3$ (top two rows), $10^3$ (bottom row), and period $\tau = 10^4$ (left), $10^3$ (center), $10^2$ (right).

##### S4.4.2 Cyclical demography: mean frequency $\overline{p}(t)$ to three-vertex order

Plots of the first non-zero correction to the deterministic approximation for the mean of the allele frequency probability distribution $\overline{p}(t)$ are presented below for various cyclical demographies. The top row, representing the same demography as the middle row, but with lower mutation rate $\mu = 10^{-8}$, is substantially noisier, as discussed in Section S4.1. The higher order correction to the analytic approximation of the mean changes little; the deterministic approximation is highly accurate, despite oscillating population size. However, as can be seen in Figure S36g, the mean does mimic the oscillatory behavior and change appreciably for weaker selection coefficients; this effect is captured by the analytic approximation for selection coefficients just above and around the lower limit of the strong selection approximation (e.g., for $s = \frac{5}{2\overline{N}_{\rm osc}}$ and $\frac{10}{2\overline{N}_{\rm osc}}$ in Figure S36g). This effect is present in Figures S36h and S36i but is less visible due to the speed of oscillations. This magnitude of change in the mean frequency is minimal (this is a subdominant effect captured by the three-vertex corrections) and is completely masked by noise when mutation rates are reduced. Oscillations in population size also transiently increase the variance of the mean frequency, an effect most clearly visible in Figures S36g and S36h due to the more severe population size reduction. This effect is the same as can be seen during a bottleneck, but is more pronounced due to the repetitive population size changes.

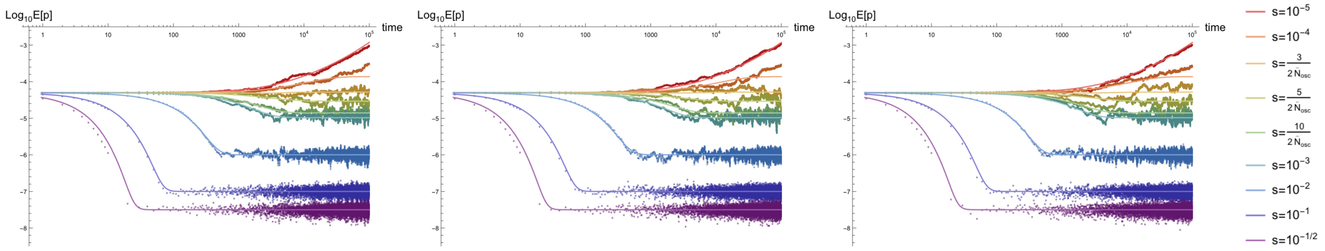


(a) $\frac{N_{\rm max}}{N_{\rm min}} = 2, \tau = 10^4, \mu = 10^{-8}$ (b) $\frac{N_{\rm max}}{N_{\rm min}} = 2, \tau = 10^3, \mu = 10^{-8}$ (c) $\frac{N_{\rm max}}{N_{\rm min}} = 2, \tau = 10^2, \mu = 10^{-8}$

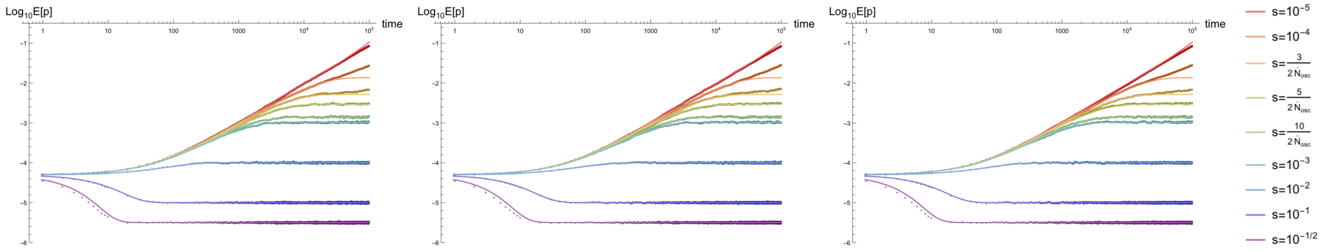


(d) $\frac{N_{\rm max}}{N_{\rm min}} = 2, \tau = 10^4, \mu = 10^{-6}$ (e) $\frac{N_{\rm max}}{N_{\rm min}} = 2, \tau = 10^3, \mu = 10^{-6}$ (f) $\frac{N_{\rm max}}{N_{\rm min}} = 2, \tau = 10^2, \mu = 10^{-6}$

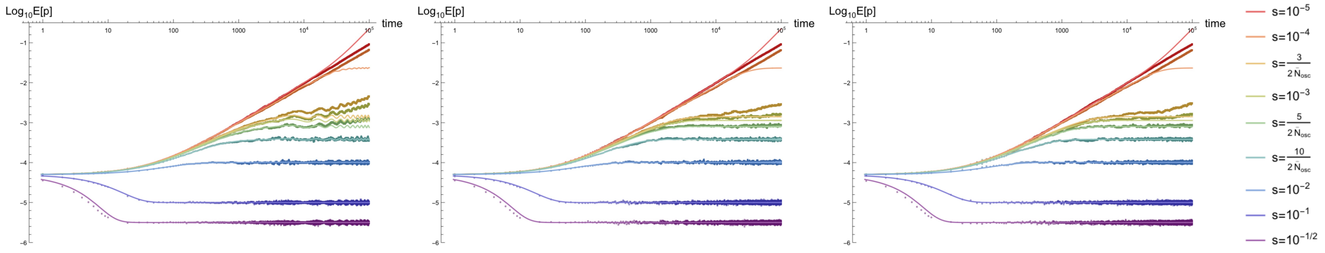


(g) $\frac{N_{\rm max}}{N_{\rm min}} = 10, \tau = 10^4, \mu = 10^{-6}$ (h) $\frac{N_{\rm max}}{N_{\rm min}} = 10, \tau = 10^3, \mu = 10^{-6}$ (i) $\frac{N_{\rm max}}{N_{\rm min}} = 10, \tau = 10^2, \mu = 10^{-6}$

Figure S36: *Time dependence of the mean of the allele frequency probability distribution $\overline{p}$ for populations with oscillating population size, including the lowest order drift-based corrections.* Plots are shown for mutation rates $\mu = 10^{-8}$ (top row), $10^{-6}$ (bottom two rows) and $\mu_b = \mu/100$, maximum population size $N_{\rm max} = 10^4$, minimum size $N_{\rm min} = N_{\rm max}/2 = 5 \times 10^3$ (top two rows), $N_{\rm max}/10 = 10^2$ (bottom row), and period $\tau = 10^4$ (left), $10^3$ (center), $10^2$ (right), and selection coefficients $s \in [10^{-5}, 10^{-1/2}]$.

##### S4.4.3 Cyclical demography: variance $\kappa_2(t)$ to two-vertex order

Plots of the lowest-order approximation to the variance, the second cumulant, of the allele frequency probability distribution $\kappa_2(t)$ are presented below for various cyclical demographies. As seen for constant and exponentially growing population sizes, the variance exhibits a transient increase and is maximized at early times. However, unlike these demographies, but similar to a bottleneck, this may not be the globally maximum variance if the reduction in population size is severe. In that case, the maximal variance seems to occur during a trough in population size one or more periods after the first minimum at $t = \tau/2$; this is more likely when $s$ is on the order of or less than than the inverse of the period, resulting in variance maximization once per period during the waltz. The dependence on $s'\tau$ is evident in the expression for the variance, but has a complicated functional form. In contrast to the mean frequency, the variance is driven by the oscillatory behavior, which is evident from the $\varphi(t)$ dependence at all times, including asymptotically, in the lowest order approximation. Note that, despite drift playing a dominant role in the dynamics of the variance, the perturbative approximation captures the oscillatory behavior impressively well–even at lowest order–lining up to simulations nearly exactly in the regime of validity.

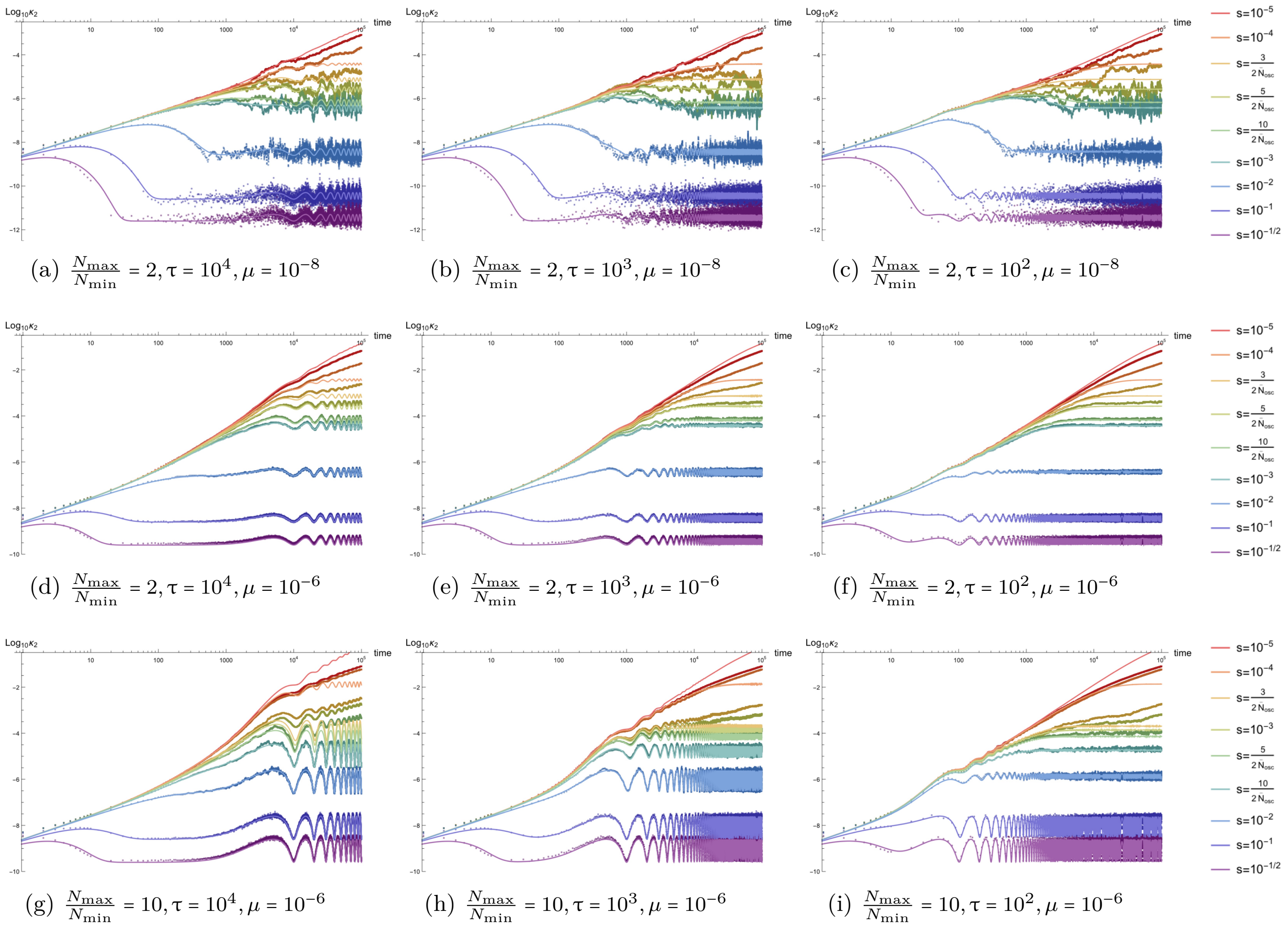


(a) $\frac{N_{\rm max}}{N_{\rm min}} = 2, \tau = 10^4, \mu = 10^{-8}$ (b) $\frac{N_{\rm max}}{N_{\rm min}} = 2, \tau = 10^3, \mu = 10^{-8}$ (c) $\frac{N_{\rm max}}{N_{\rm min}} = 2, \tau = 10^2, \mu = 10^{-8}$

(d) $\frac{N_{\rm max}}{N_{\rm min}} = 2, \tau = 10^4, \mu = 10^{-6}$ (e) $\frac{N_{\rm max}}{N_{\rm min}} = 2, \tau = 10^3, \mu = 10^{-6}$ (f) $\frac{N_{\rm max}}{N_{\rm min}} = 2, \tau = 10^2, \mu = 10^{-6}$

(g) $\frac{N_{\rm max}}{N_{\rm min}} = 10, \tau = 10^4, \mu = 10^{-6}$ (h) $\frac{N_{\rm max}}{N_{\rm min}} = 10, \tau = 10^3, \mu = 10^{-6}$ (i) $\frac{N_{\rm max}}{N_{\rm min}} = 10, \tau = 10^2, \mu = 10^{-6}$

Figure S37: *Time dependence of the variance of the allele frequency probability distribution $\kappa_2$ for populations with oscillating population size.* Parameter combinations correspond to those shown in Figure S36.

##### S4.4.4 Cyclical demography: homozygosity $E[p^2(t)]$ to two-vertex order

Plots of the lowest-order approximation to the homozygosity $E[p^2(t)]$, the second non-central moment of the allele frequency probability distribution, are presented below for various cyclical demographies. As seen previously, the homozygosity is largely driven by the dynamics of the variance. The notable difference is in the initial value, when the variance is zero due to the delta function describing the initial frequency distribution but the homozygosity is given by a constant value $p_0^2$. This moment thus exhibits a transition between the mean-driven and variance-driven regimes, which occurs during the transient period. Note that, while the oscillatory behavior is confounded with the noise for low mutation rates (see, e.g., Figure S38a for $\mu = 10^{-8}$, it is still visible. The variance of the homozygosity, dependent on the first four cumulants, also exhibits clear oscillations even when mutation is comparatively rare. Note that, if selection is sufficiently strong, the homozygosity (and higher moments) can briefly pseudo-equilibrate to a metastable value determined by the initial population size, but this only occurs if the timescale of selection is much less than the period (i.e., $s\tau \ll 1$); this is analogous to pseudo-equilibration for strong selection coefficients under exponential growth, taking the form of a relatively flat trough at early times in cyclical demographies. This is clearest in Figure S38g (with $\tau = 10^4$) for $s = 10^{-1/2}$ and $10^{-1}$, but vanishes as the period is reduced in Figure S38i (with for $\tau = 10^2$).

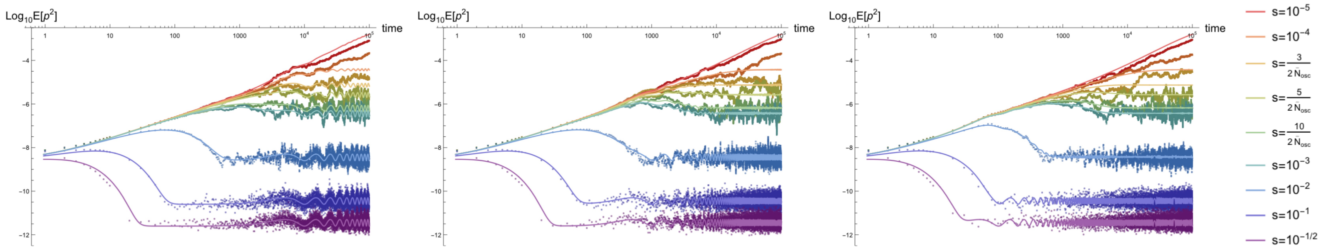


(a) $\frac{N_{\max}}{N_{\min}} = 2, \tau = 10^4, \mu = 10^{-8}$ (b) $\frac{N_{\max}}{N_{\min}} = 2, \tau = 10^3, \mu = 10^{-8}$ (c) $\frac{N_{\max}}{N_{\min}} = 2, \tau = 10^2, \mu = 10^{-8}$

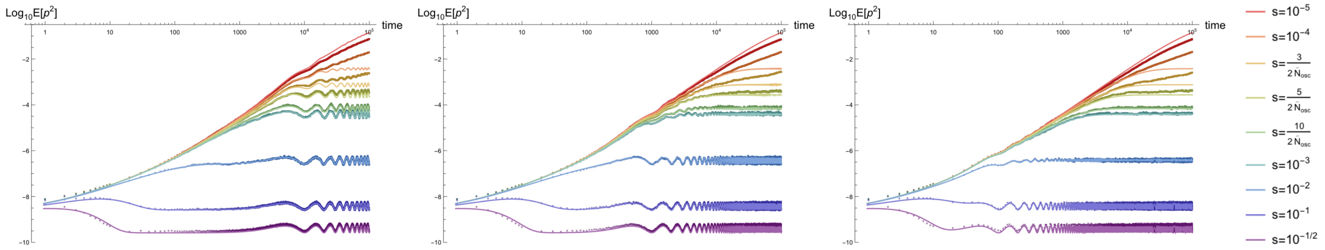


(d) $\frac{N_{\max}}{N_{\min}} = 2, \tau = 10^4, \mu = 10^{-6}$ (e) $\frac{N_{\max}}{N_{\min}} = 2, \tau = 10^3, \mu = 10^{-6}$ (f) $\frac{N_{\max}}{N_{\min}} = 2, \tau = 10^2, \mu = 10^{-6}$

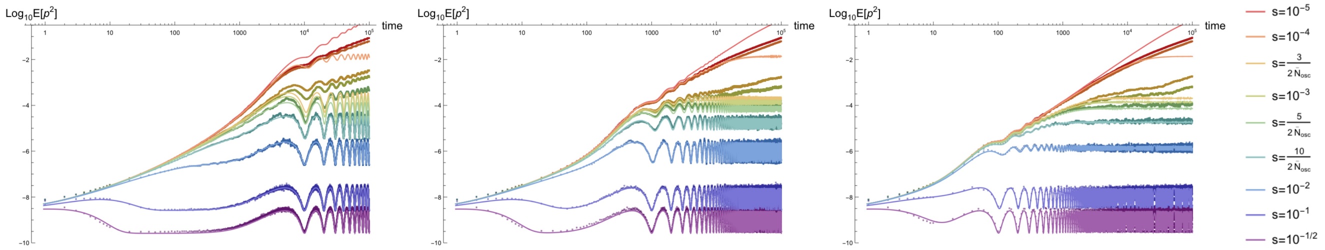


(g) $\frac{N_{\max}}{N_{\min}} = 10, \tau = 10^4, \mu = 10^{-6}$ (h) $\frac{N_{\max}}{N_{\min}} = 10, \tau = 10^3, \mu = 10^{-6}$ (i) $\frac{N_{\max}}{N_{\min}} = 10, \tau = 10^2, \mu = 10^{-6}$

Figure S38: *Time dependence of the homozygosity, the second non-central moment of the allele frequency probability distribution $E[p^2]$, for populations with oscillating population size.* Parameter combinations correspond to those shown in Figure S36.

##### S4.4.5 Cyclical demography: third cumulant $\kappa_3$ to three-vertex order

Plots of the lowest-order approximation to the third cumulant of the allele frequency probability distribution $\kappa_3(t)$ are presented below for various cyclical demographies. As discussed previously, the dynamics of the third cumulant are tied to those of the second cumulant such that the time dependence is highly similar to plots of the variance, but on an overall reduced scale. Again, the analytic approximations line up exceptionally well to simulated results, capturing both the transient and oscillatory dynamics for a large range of selection coefficients. The third cumulant is only maximized during the transient period for low mutation rate and very strong selection (see, e.g., selection coefficients $s \geq 10^{-2}$ in Figures S39a–S39c). As seen in the second cumulant and moment, pseudo-equilibration may occur at early times if $s\tau \ll 1$.

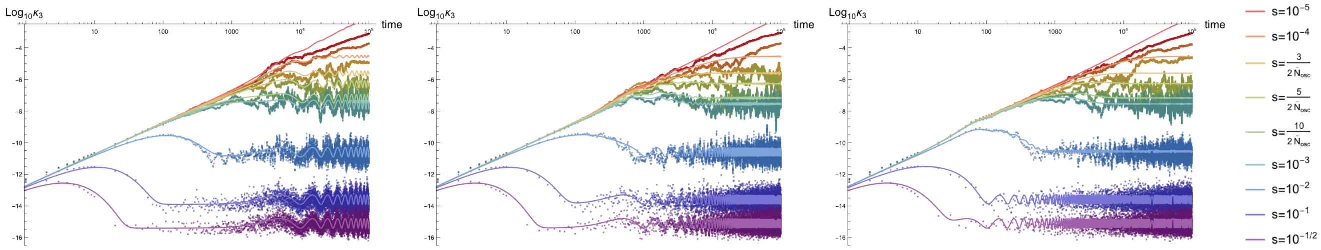


(a) $\frac{N_{\rm max}}{N_{\rm min}} = 2, \tau = 10^4, \mu = 10^{-8}$ (b) $\frac{N_{\rm max}}{N_{\rm min}} = 2, \tau = 10^3, \mu = 10^{-8}$ (c) $\frac{N_{\rm max}}{N_{\rm min}} = 2, \tau = 10^2, \mu = 10^{-8}$

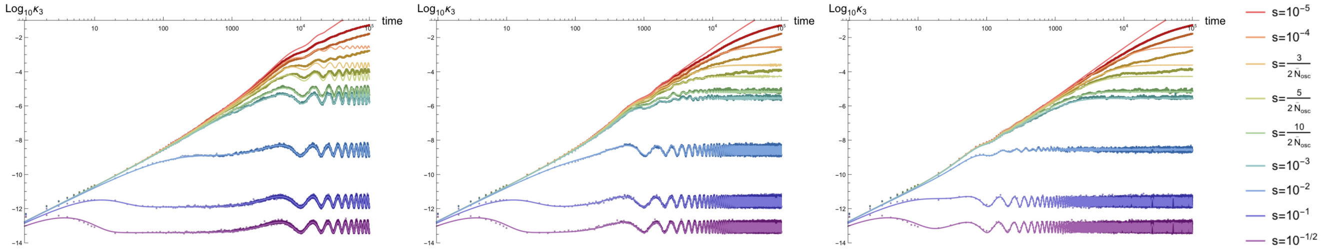


(d) $\frac{N_{\rm max}}{N_{\rm min}} = 2, \tau = 10^4, \mu = 10^{-6}$ (e) $\frac{N_{\rm max}}{N_{\rm min}} = 2, \tau = 10^3, \mu = 10^{-6}$ (f) $\frac{N_{\rm max}}{N_{\rm min}} = 2, \tau = 10^2, \mu = 10^{-6}$

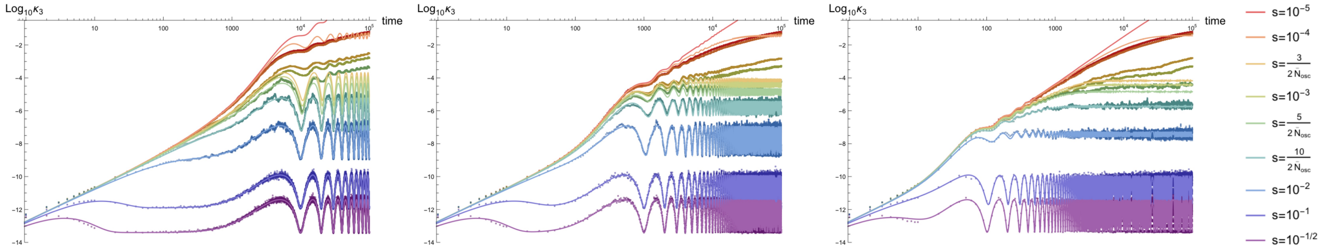


(g) $\frac{N_{\rm max}}{N_{\rm min}} = 10, \tau = 10^4, \mu = 10^{-6}$ (h) $\frac{N_{\rm max}}{N_{\rm min}} = 10, \tau = 10^3, \mu = 10^{-6}$ (i) $\frac{N_{\rm max}}{N_{\rm min}} = 10, \tau = 10^2, \mu = 10^{-6}$

Figure S39: *Time dependence of the third cumulant of the allele frequency probability distribution $\kappa_3$ for populations with oscillating population size.* Parameter combinations correspond to those shown in Figure S36.

##### S4.4.6 Cyclical demography: third non-central moment $E[p^3(t)]$ to three-vertex order

Plots of the lowest-order approximation to the third non-central moment of the allele frequency probability distribution $E[p^3(t)]$ are presented below for various cyclical demographies. The third moment mimics the behavior of the third cumulant, which is in turn driven by the dynamics of the variance, as discussed in the context of other demographies. Like the homozygosity, the third moment is initially at a fixed value of $p_0^3$ due to the delta function, after which it exhibits transient behavior and eventually reaches the asymptotic waltz around the value determined by the effective population size $\overline{N}_{\text{osc}}$. Despite comparatively noisy estimation of this moment relative to lower moments, the analytic approximation captures the oscillatory strong selection behavior extremely well, independent of the periodicity. Again, pseudo-equilibration can occur at early times for strong selection coefficients $s\tau \ll 1$.

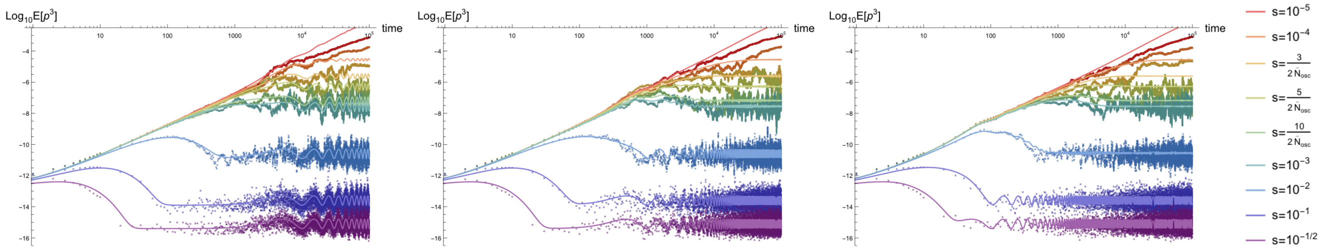


(a) $\frac{N_{\max}}{N_{\min}} = 2, \tau = 10^4, \mu = 10^{-8}$ (b) $\frac{N_{\max}}{N_{\min}} = 2, \tau = 10^3, \mu = 10^{-8}$ (c) $\frac{N_{\max}}{N_{\min}} = 2, \tau = 10^2, \mu = 10^{-8}$

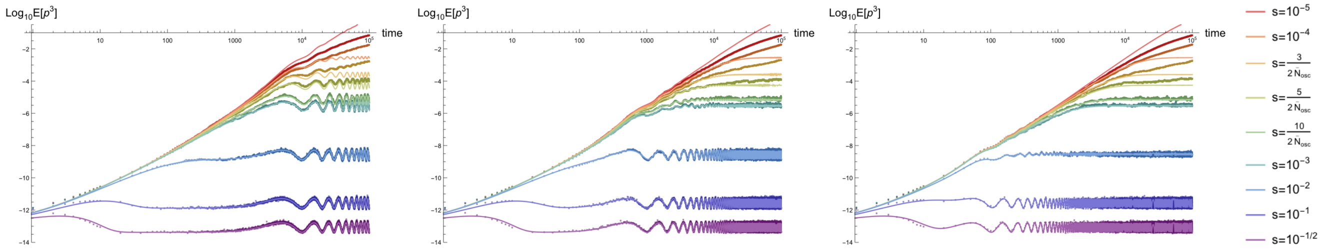


(d) $\frac{N_{\max}}{N_{\min}} = 2, \tau = 10^4, \mu = 10^{-6}$ (e) $\frac{N_{\max}}{N_{\min}} = 2, \tau = 10^3, \mu = 10^{-6}$ (f) $\frac{N_{\max}}{N_{\min}} = 2, \tau = 10^2, \mu = 10^{-6}$

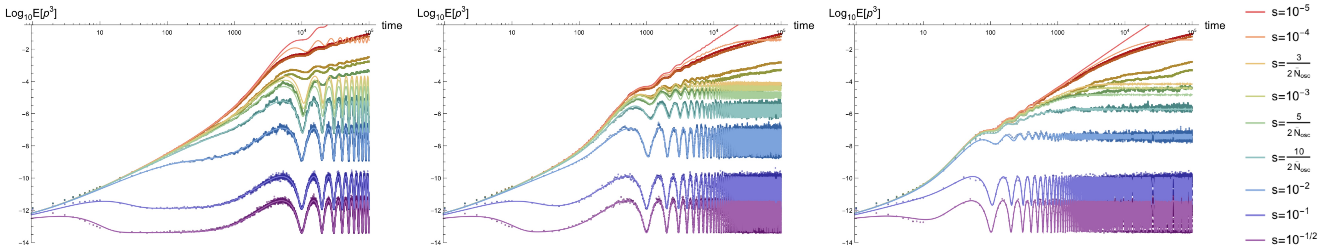


(g) $\frac{N_{\max}}{N_{\min}} = 10, \tau = 10^4, \mu = 10^{-6}$ (h) $\frac{N_{\max}}{N_{\min}} = 10, \tau = 10^3, \mu = 10^{-6}$ (i) $\frac{N_{\max}}{N_{\min}} = 10, \tau = 10^2, \mu = 10^{-6}$

Figure S40: *Time dependence of the third non-central moment of the allele frequency probability distribution $E[p^3]$ for populations with oscillating population size.* Parameter combinations correspond to those shown in Figure S36.

##### S4.4.7 Cyclical demography: skew $K_3(t)$ to three-vertex order

Plots of the lowest-order approximation to the skew, the standardized third cumulant, of the allele frequency probability distribution $K_3(t)$ are presented below for various cyclical demographies. This moment is particularly informative about the oscillatory dynamics of the distribution. Comparing the figures below to those for constant demography (see Figure S9), the skew approaches an oscillatory time-dependent value, rather than a constant, centered around the value that would be associated with $2\overline{N}_{\rm osc}\mu$ in a constant demography. This indicates that not all of the oscillatory behavior in the third moment is contained in the variance dependence; under strong selection, the skew rapidly pseudo-equilibrates as if in a constant population of size $N_{\rm max}(= N_{\rm osc}(0))$, but begins to oscillate on the timescale of the period $\tau$ due to the asymptotic waltzing of the distribution during the second period. Transient pseudo-equilibration only occurs for selection coefficients that are efficient in the initial population ($2N_{\rm max}s \gg 1$), but also requires sufficiently rapid adjustment in a regime dictated by the period ($s\tau \gg 1$); otherwise, the asymptotic waltz is reached uninterrupted. Asymptotically, the skew oscillates with an appreciable amplitude that is monotonically (and postively correlated) with both the period and the selection coefficient. This indicates that, while the variance of the skew may remain tight and looks unrelated to the period or selection coefficient, time-averaging the skew during the late-time dynamics (i.e., $\int_c^{c+\tau} dt K_3(t)$ beginning at some late time $c \gg \tau$) results in an observable sensitive to the amplitude of oscillations and thus correlated to both the selection period; the appreciable amplitude of oscillations over time will be converted into variance of the time-averaged skew. Also, see Section S4.4.10.

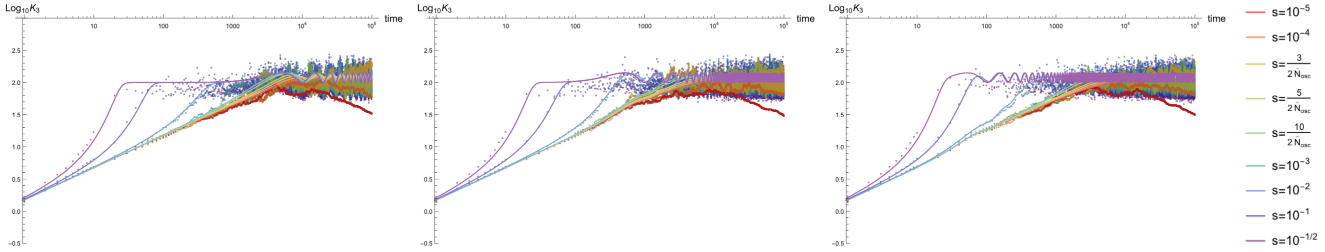

(a) $\frac{N_{\rm max}}{N_{\rm min}} = 2, \tau = 10^4, \mu = 10^{-8}$ (b) $\frac{N_{\rm max}}{N_{\rm min}} = 2, \tau = 10^3, \mu = 10^{-8}$ (c) $\frac{N_{\rm max}}{N_{\rm min}} = 2, \tau = 10^2, \mu = 10^{-8}$

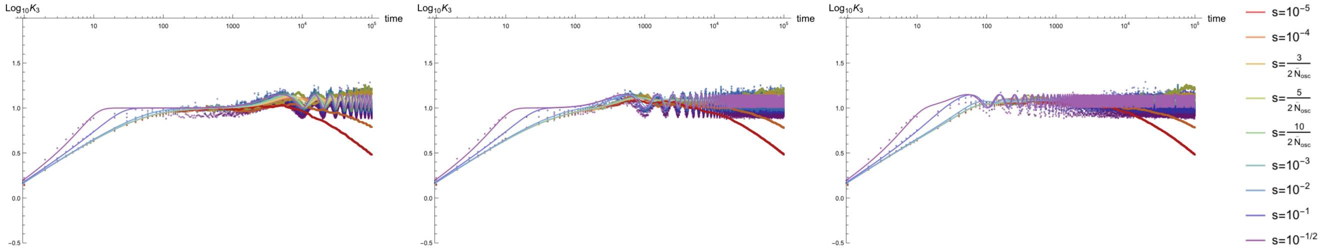

(d) $\frac{N_{\rm max}}{N_{\rm min}} = 2, \tau = 10^4, \mu = 10^{-6}$ (e) $\frac{N_{\rm max}}{N_{\rm min}} = 2, \tau = 10^3, \mu = 10^{-6}$ (f) $\frac{N_{\rm max}}{N_{\rm min}} = 2, \tau = 10^2, \mu = 10^{-6}$

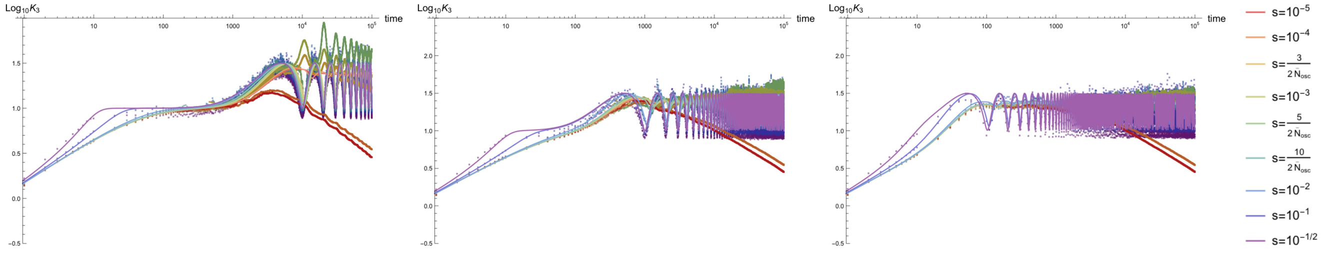

(g) $\frac{N_{\rm max}}{N_{\rm min}} = 10, \tau = 10^4, \mu = 10^{-6}$ (h) $\frac{N_{\rm max}}{N_{\rm min}} = 10, \tau = 10^3, \mu = 10^{-6}$ (i) $\frac{N_{\rm max}}{N_{\rm min}} = 10, \tau = 10^2, \mu = 10^{-6}$

Figure S41: *Time dependence of the skew of the allele frequency probability distribution $K_3$ for populations with oscillating population size.* Parameter combinations correspond to those shown in Figure S36.

##### S4.4.8 Cyclical demography: fourth cumulant $\kappa_4(t)$ to four-vertex order

Plots of the lowest-order approximation to the fourth cumulant of the allele frequency probability distribution $\kappa_4(t)$ are presented below for various cyclical demographies. As discussed previously, the dynamics of the fourth cumulant are tied to the second cumulant such that the time dependence is highly similar to plots of the variance, butt, like the third cumulant, on an overall reduced scale. Lowest-order approximations to the forth cumulant line up exceptionally well to simulated results, capturing both the transient and oscillatory dynamics for a large range of selection coefficients. The dynamics of the fourth cumulant are largely the same as those for the third cumulant. As seen in the lower cumulants, pseudo-equilibration may occur at early times if $s\tau \ll 1$.

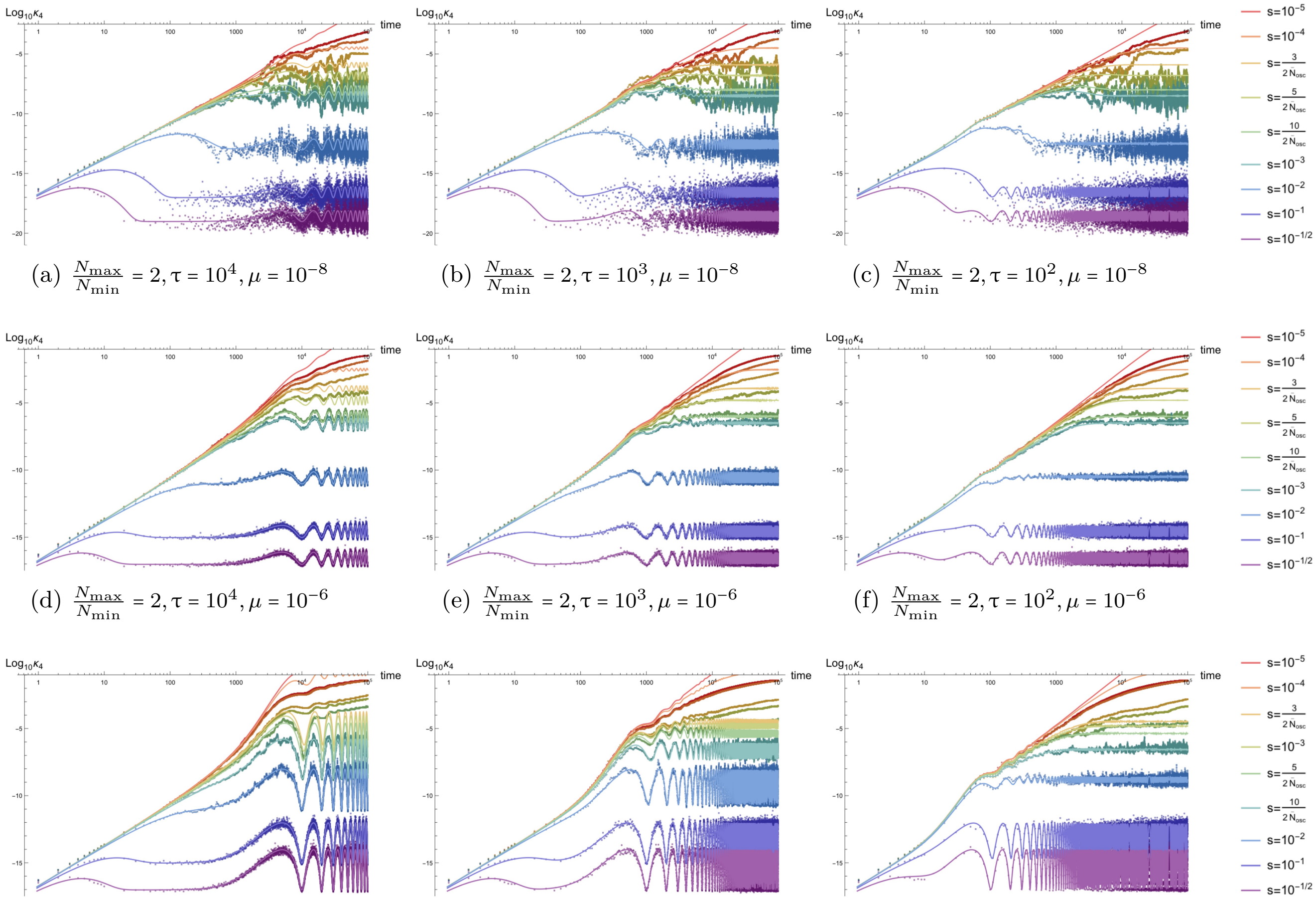


(a) $\frac{N_{\max}}{N_{\min}} = 2, \tau = 10^4, \mu = 10^{-8}$ (b) $\frac{N_{\max}}{N_{\min}} = 2, \tau = 10^3, \mu = 10^{-8}$ (c) $\frac{N_{\max}}{N_{\min}} = 2, \tau = 10^2, \mu = 10^{-8}$

(d) $\frac{N_{\max}}{N_{\min}} = 2, \tau = 10^4, \mu = 10^{-6}$ (e) $\frac{N_{\max}}{N_{\min}} = 2, \tau = 10^3, \mu = 10^{-6}$ (f) $\frac{N_{\max}}{N_{\min}} = 2, \tau = 10^2, \mu = 10^{-6}$

(g) $\frac{N_{\max}}{N_{\min}} = 10, \tau = 10^4, \mu = 10^{-6}$ (h) $\frac{N_{\max}}{N_{\min}} = 10, \tau = 10^3, \mu = 10^{-6}$ (i) $\frac{N_{\max}}{N_{\min}} = 10, \tau = 10^2, \mu = 10^{-6}$

Figure S42: *Time dependence of the fourth cumulant of the allele frequency probability distribution $\kappa_4$ for populations with oscillating population size.* Parameter combinations correspond to those shown in Figure S36.

##### S4.4.9 Cyclical demography: fourth non-central moment $E[p^4(t)]$ to four-vertex order

Plots of the lowest-order approximation to the fourth non-central moment of the allele frequency probability distribution $E[p^4(t)]$ are presented below for various cyclical demographies. The fourth moment mimics the behavior of the fourth cumulant, which is in turn driven by the dynamics of the variance, as described above. Like the second and third non-central moments, the fourth moment is initially at a fixed value of $p_0^4$ due to the delta function, exhibits transient behavior, and dances an asymptotic waltz at late times around the value determined by the effective population size $\overline{N}_{\text{osc}}$. Despite the additional noise inherent in estimating higher moments, the analytic approximation captures the oscillatory strong selection behavior extremely well, independent of the periodicity. Again, a brief pseudo-equilibration occurs in the fourth moment for parameters $s\tau \ll 1$.

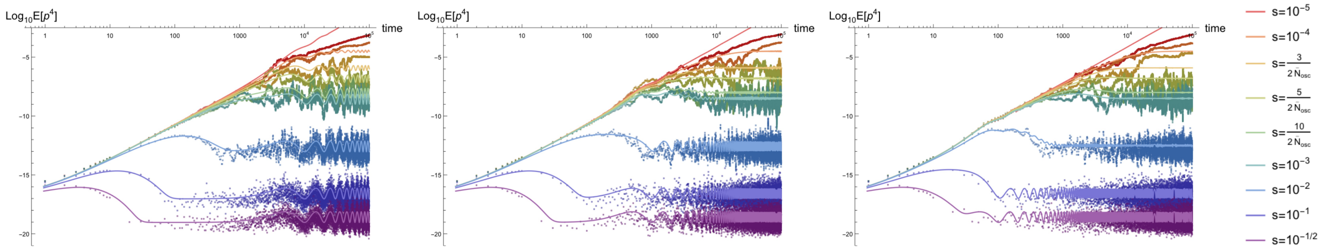


(a) $\frac{N_{\max}}{N_{\min}} = 2, \tau = 10^4, \mu = 10^{-8}$ (b) $\frac{N_{\max}}{N_{\min}} = 2, \tau = 10^3, \mu = 10^{-8}$ (c) $\frac{N_{\max}}{N_{\min}} = 2, \tau = 10^2, \mu = 10^{-8}$

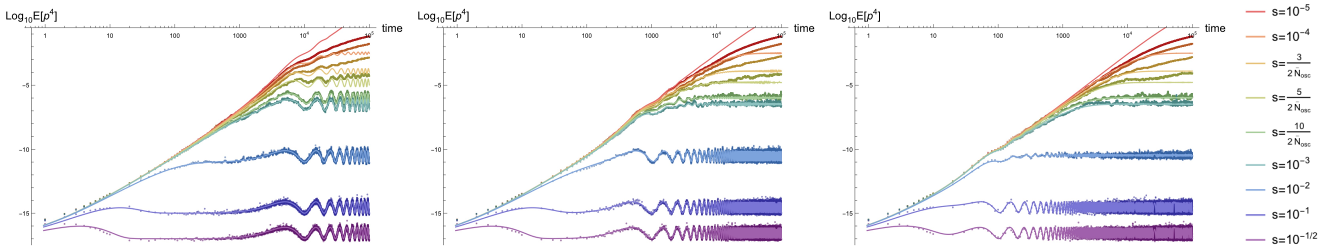


(d) $\frac{N_{\max}}{N_{\min}} = 2, \tau = 10^4, \mu = 10^{-6}$ (e) $\frac{N_{\max}}{N_{\min}} = 2, \tau = 10^3, \mu = 10^{-6}$ (f) $\frac{N_{\max}}{N_{\min}} = 2, \tau = 10^2, \mu = 10^{-6}$

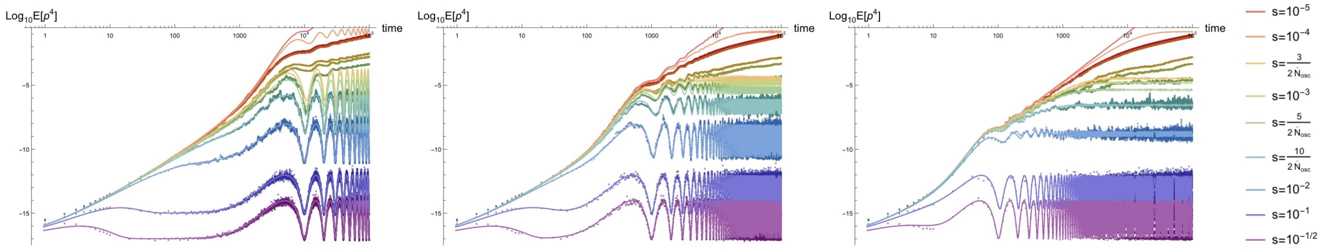


(g) $\frac{N_{\max}}{N_{\min}} = 10, \tau = 10^4, \mu = 10^{-6}$ (h) $\frac{N_{\max}}{N_{\min}} = 10, \tau = 10^3, \mu = 10^{-6}$ (i) $\frac{N_{\max}}{N_{\min}} = 10, \tau = 10^2, \mu = 10^{-6}$

Figure S43: *Time dependence of the fourth non-central moment of the allele frequency probability distribution $E[p^4]$ for populations with oscillating population size.* Parameter combinations correspond to those shown in Figure S36.

##### S4.4.10 Cyclical demography: excess kurtosis $K_4(t)$ to four-vertex order

Plots of the lowest-order approximation to the excess kurtosis, the standardized fourth cumulant, of the allele frequency probability distribution $\kappa_3(t)$ are presented below for various cyclical demographies. The qualitative dynamics of the excess kurtosis are similar to those for the skew, indicating that the discussion in Section S4.4.7) is applicable here (acknowledging that estimation of the excess kurtosis is inherently noisier). Comparing to the skew, the transient pseudo-equilibration (for selection coefficients $2N_{\text{osc}}(0)s \gg 1, s\tau \gg 1$) appears to be more pronounced in the excess kurtosis. In addition to the monotonic relations between the time-averaged excess kurtosis $\int_c^{c+\tau} dt K_4(t)$ (or time-averaged skew $\int_c^{c+\tau} dt K_3(t)$) and both the selection coefficient $s$ and period $\tau$, the asymptotic behavior of distinct selection coefficients appears to decouple for both moments. A phase emerges that is dependent on the selection coefficient and describes a temporal delay in oscillations of the excess kurtosis (and skew) relative to oscillations in the population size itself (this behavior is contained in the analytic expression for both moments). This phase can also be ignored, if desired, by time averaging; this allows sites under distinct selection coefficients to be pooled to estimate the average excess kurtosis (or skew). For organism with readily accessible temporal data, the variance of the time average of the excess kurtosis (or skew), after pooling deleterious variants to increase power, may aid in the identification of oscillations in the population size, as this quantity is generically inflated for cyclical populations relative to a constant population (all other parameters remaining equal).

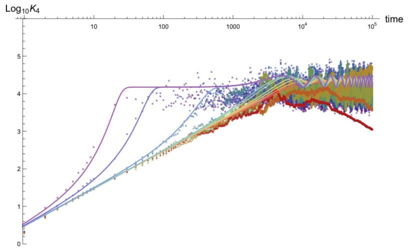


(a) $\frac{N_{\max}}{N_{\min}} = 2, \tau = 10^4, \mu = 10^{-8}$

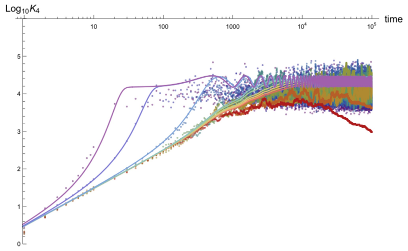


(b) $\frac{N_{\max}}{N_{\min}} = 2, \tau = 10^3, \mu = 10^{-8}$

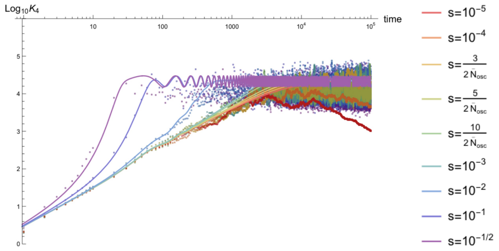


(c) $\frac{N_{\max}}{N_{\min}} = 2, \tau = 10^2, \mu = 10^{-8}$

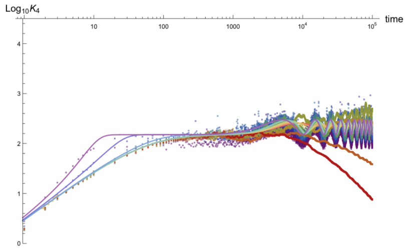


(d) $\frac{N_{\max}}{N_{\min}} = 2, \tau = 10^4, \mu = 10^{-6}$

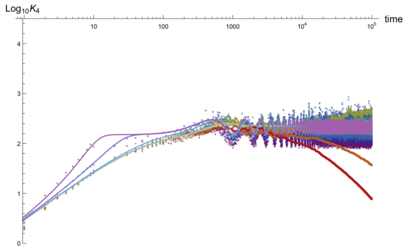


(e) $\frac{N_{\max}}{N_{\min}} = 2, \tau = 10^3, \mu = 10^{-6}$

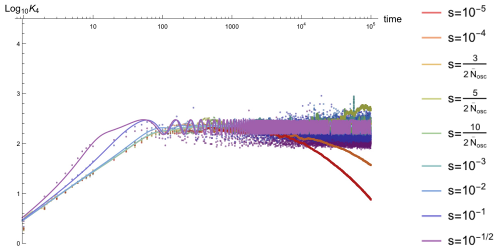


(f) $\frac{N_{\max}}{N_{\min}} = 2, \tau = 10^2, \mu = 10^{-6}$

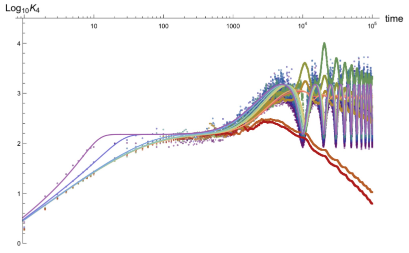


(g) $\frac{N_{\max}}{N_{\min}} = 10, \tau = 10^4, \mu = 10^{-6}$

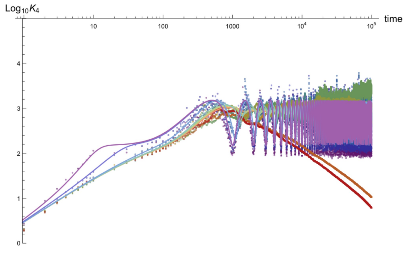


(h) $\frac{N_{\max}}{N_{\min}} = 10, \tau = 10^3, \mu = 10^{-6}$

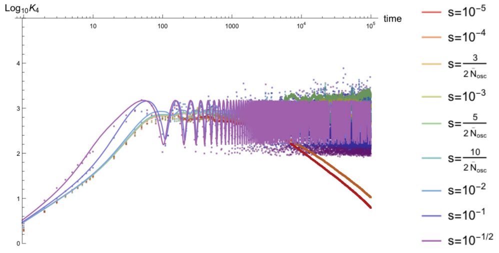


(i) $\frac{N_{\max}}{N_{\min}} = 10, \tau = 10^2, \mu = 10^{-6}$

Figure S44: *Time dependence of the excess kurtosis of the allele frequency probability distribution $K_4$ for populations with oscillating population size.* Parameter combinations correspond to those shown in Figure S36.

### S5 On the computation of the fourth and higher cumulants of the allele frequency distribution

Although higher moments can be computed using the same methodology described in the manuscript text, enumeration of the diagrams and evaluation of the resulting integrals becomes complicated. The fourth cumulant will be computed here as an example for a method of computation for general cumulants $\kappa_n$ (the main text contains a summary of the results for the fourth cumulant presented here). However, this requires the warning that the results here *only* apply to contributions represented by diagrams analogous to diagrams that appear as the lowest contributions to the fourth cumulant. Additional moments can be analyzed in a similar way, but require the identification of any missing diagrams, including those that do not have topological analogs to diagrams for the first four moments (e.g., there is one new type of diagram for the lowest order contributions to the fifth moment, etc.). The class of diagrams that has analogous contributions to those for both the the third and fourth cumulants are presented, along with an example of a diagram topology that appears in the fourth cumulant but has no analog for lower moments. Thus, this section is intended to provide some generalization of diagrammatic contributions that appear for the general cumulant $\kappa_n$, but cannot provide full expressions for higher cumulants; they must be constructed by adding the functional forms provided here to all additional diagrams that appear at that moment. Having said that, the presentation here may help in future applications that require analysis of specific higher moments or in an effort to find a general set of closed-form approximations for all moments, which is beyond the scope of this manuscript. Here, I will restrict my analysis to constant population size, but, in principle, the same methodology can be used in conjunction with integral expressions provided in Supplemental Material S3.4 to find generalized forms along the lines of expressions below (in practice, this is likely difficult for most demographies due to the complicated functional form of $N(t)$; the exponential demography may provide an example of where this is possible).

#### S5.1 Maximally connected diagrams

The fourth cumulant has four contributing diagrams at lowest order, each containing four vertices. The first two are analogous to the diagrams for $\kappa_3$ in Diagram D7. These diagrams have a single terminal vertex (one with coupling constant $p_0$ and another with coupling constant $\mu$) and three copies of the $N_3$ vertex in a *minimally connected* (or 'maximally long') chain of drift vertices.

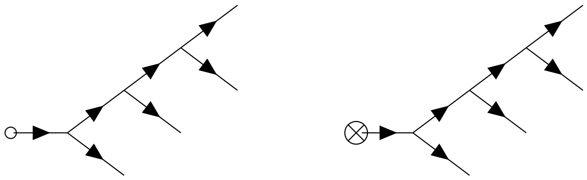

(D10)

The associated contributions sum to the following expression.

$$\kappa_{4D10} = \frac{2p_0}{\gamma'^3} e^{-s't} \left(1 - e^{-s't}\right)^3 + \frac{\lambda'}{2\gamma'^3} \left(1 - e^{-s't}\right)^4 \tag{S80}$$

For higher moments, all diagrams with this minimally connected topology have analogous contributions. The overall pre-factor is a product of $n!$ (the number of ways to connect to the external vertices), $(n-1)!/(n-1)!$ (the number of ways to choose each subsequent $N_3$ vertex divided by the repeated vertex factor), and a factor of $2^{n-2}$ (the number of $N_3$ vertices with two available $\tilde{p}$ connections that can be swapped), for an overall factor of $2^{n-2}n!$. The factor of $2^{n-2}$ partially cancels the overall factor of $1/2^{n-1}(2N)^{n-1}$ from the $n-1$ copies of the coupling constant $1/4N$, where each $1/2N$ becomes a factor of $\gamma'$ after each integral of an exponential produces a multiple of $1/s'$. Additionally, integration over $n-1$ vertex times $t_i$ for the diagram containing the $p_0$ results in an overall constant of $1/(n-1)!$. Similarly, the $n$ vertex time integrals in the diagram containing the $\mu$ vertex results in an overall constant of $1/n!$. Both of these cancel or partially cancel the $n$! dependence from the counting factor. Finally, integration of the two minimally connected diagrams contributing to the $n^{\text{th}}$ cumulant $\kappa_n$ results in the following time dependence and multiplicative factors.

$$\kappa_n{}^{\text{long}} = \frac{n\ p_0}{2(\gamma')^{n-1}} e^{-s't} \left(1 - e^{-s't}\right)^{n-1} + \frac{\lambda'}{2(\gamma')^{n-1}} \left(1 - e^{-s't}\right)^{n} \tag{S81}$$

Here, $1 < n \in \mathbb{Z}^+$ (i.e., any positive integer greater than one); contributions to the analogous diagrams for $n = 2$ and $n = 3$ are given in the equations for the variance and non-standardized skew in the main text, both of which follow the functional dependence in Equation S81 (also see results presented for Diagrams D4 and D7 in Supplemental Material S3.2 and S3.3, respectively). The contributions for the mean frequency (i.e., $n = 1$) in the manuscript text follow the same pattern (see results in Supplemental Material S3.1) with the caveat that the factor of $2^{n-2}$ requires $n$ to be greater than one to be valid.

#### S5.2 Minimally connected diagrams

The final two four-vertex diagrams for $\kappa_4$ have a topology with no analogous diagrams for cumulants $\kappa_n$ with $n \leq 3$.

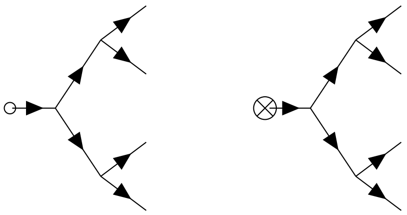

(D11)

These diagrams are a series of *maximally connected* $N_3$ drift vertices; higher moment analogs are diagrams with one terminal vertex ($p_0$ or $\mu$) and $n - 1$ drift vertices contributing to the moment $E[p^n]$ that are maximally connected or 'maximally wide' (i.e., the minimal average number of propagators between the terminal and external vertices in a connected diagram). Note that this is only well-defined for even cumulants $\kappa_{2^n}$ (evenly spaced branches, as with the diagrams above), but there are a variety of diagrams that can be considered maximally connected for both odd cumulants and for even cumulants that are not powers of two. This is the first example thus far of multiple diagrams contributing to a single cumulant with identical vertices and distinct topologies (e.g., the two leftmost or two rightmost in Diagrams D10 and D11). This becomes more relevant for higher moments, but can also occur in diagrams with, for example, multiple selection vertices (with coupling constant $s$) when computing high-vertex order contributions to the mean frequency. Calculating the appropriate integrals results in the following.

$$\kappa_{4D11} = \frac{p_0 e^{-s't}}{\gamma'^3} \left(1 - e^{-s't}\right)^3 + \frac{\lambda'}{4\gamma'^3} \left(1 - e^{-s't}\right)^4 \tag{S82}$$

These diagrams thus represent exactly half of the contributions from the minimally connected diagrams in Diagram D10, but are of the same order and thus non-negligible when computing $\kappa_4$. The additional factor of two found in Equation S80 can be attributed to a counting factor of $2^2 = 4$ associated with the choice of which $\tilde{p}$ exiting each of the $N_3$ vertices connects to a subsequent $N_3$ vertex (as opposed to an external vertex) and an additional factor of 1/2 from integration due to the difference in the propagators. The counting factor for maximally connected diagrams is one (see Appendix A2). This corresponds to the fact that, after laying down the first propagator to connect any two vertices in the above diagrams, the connections are fully determined for all remaining propagators (i.e., they either connect to the terminal vertex $G(t_i, 0)$, an external vertex $G(t, t_i)$, or the remaining connection between two $N_3$ vertices $G(t_{j+1}, t_j)$ a some specific vertex time $t_j$).

$$\kappa_{4D11} = \frac{n\ p_0 e^{-s't}}{2^{n-1}\gamma'^{n-1}} \left(1 - e^{-s't}\right)^{n-1} + \frac{\lambda'}{2^{n-1}\gamma'^{n-1}} \left(1 - e^{-s't}\right)^{n} \tag{S83}$$

Again, note that this expression is well-defined only for diagrams representing contributions to even cumulants, but it *may* also apply to some maximally connected diagrams for odd cumulants, though not necessarily for all (the author admits to not having proved the latter). It is clear from these generalized expressions that, as $n$ increases for successively higher moments, minimally connected diagrams rapidly starts to dominate over maximally connected diagrams corresponding to the same cumulant $\kappa_n$. This is due to a suppression by a relative factor of $1/2^{n-2}$ in the expression for maximally connected diagrams, which originates from (a lack of cancellation of) the counting factor for the number of allowed sets of Wick contractions (see Appendix A2). Interestingly, contributions from minimally connected diagrams can effectively be ignored (if desired) for cumulants $\kappa_{2^n}$ with sufficiently large $n$ (e.g., roughly for $n \geq 3$, where the relative contribution of maximally connected diagrams is $\leq 1/2^6 = 1/64$). However, contributions from a large number of additional diagrams without analogs for cumulants $n \leq 4$ tend to dominate due to the multiplicity of many diagrams with non-unity counting factors. This can be understood by comparing Equation S81 at asymptotically late times to moments of the equilibrium frequency distribution $\phi_{\text{eq}}(p)$ in the strong selection limit [1] by using the knowledge that minimally connected diagrams have the highest counting number of all contributing diagrams.

$$\int_0^1 dp\ p^n \phi_{\text{eq}}(p) \gg \lim_{t\to\infty} \kappa_n{}^{\text{long}} = \frac{\lambda'}{2^{n-1}\gamma'^{n-1}} \tag{S84}$$

#### S5.3 The sum of minimally and maximally connected diagrams

The time dependence of the fourth cumulant, the non-standardized excess kurtosis, is given by the sum of Diagrams S80 and S82, both of which have the same functional form.

$$\kappa_4 = \kappa_{4D10} + \kappa_{4D11} + \mathcal{O}[5V] \approx \frac{3p_0}{\gamma'^3} e^{-s't}\left(1 - e^{-s't}\right)^3 + \frac{3\lambda'}{4\gamma'^3}\left(1 - e^{-s't}\right)^4 \tag{S85}$$

Similarly, the subset of lowest-order contributions to the general cumulant $\kappa_n$ from directly analogous diagrams can be obtained by summing Equations S81 and S83. At the risk of repetition, these do not represent the *total* contribution to the lowest-order approximation for arbitrary cumulants; this presentation is intended to provide intuition for how general cumulants are computed.
